# Extensive HGT between soil-living hexapods contributes to functional convergence

**DOI:** 10.64898/2026.11.13.766446

**Authors:** Alexandra Jalaber de Dinechin, Maxime Policarpo, Clément Gilbert, Florian Maumus

## Abstract

Horizontal gene transfer (HGT) has been documented in diverse metazoan lineages, with most reported cases involving putative non-metazoan donors. Here, we combine protein taxonomic assignment with distance-based and phylogenetic approaches to infer 440 independent HGT events in sciarid flies (Sciaridae, Diptera). While 67 of these events involve genes of non-metazoan origin (mainly from bacteria and fungi), 352 likely occurred directly between sciarid flies and another group of soil-dwelling hexapods, the springtails (Collembola). Most of these HGTs (n=264) occurred in the springtail-to-sciarid orientation and involve genes that are expressed, evolve under functional constraints, and are enriched in carbohydrate and redox metabolism. Remarkably, genes springtails acquired independently from non-metazoans are enriched in the same functional categories. Consistent with the proposed role of these genes in processing plant- and fungal-derived polysaccharides and lignin compounds, their expression in a sciarid species is particularly high in larvae, which, like springtails, graze on plant and fungal resources. Because springtails were present in terrestrial habitats long before sciarid flies, our results suggest sciarid colonization of these habitats may have been facilitated by the horizontal acquisition of numerous springtail genes that were useful in soil and functional in a metazoan background. They also reveal that metazoan-to-metazoan HGT can be common in some lineages, calling for an in-depth investigation of the mechanisms underlying these transfers and for a systematic appraisal of their consequences on metazoan evolution.

## Introduction

Horizontal gene transfer (HGT) is the transmission of genes between different organisms through mechanisms other than reproduction. In prokaryotes, the patterns, mechanisms, and vectors of HGT are well understood, and these transfers are common and a significant source of innovation (1). HGT occurs less frequently in eukaryotes, but numerous well-documented cases exist across various lineages. Many of these transfers have likely facilitated rapid evolutionary changes by introducing new functions into recipient species (2–5). This is especially evident in insects, where multiple horizontally acquired genes influence traits such as reproductive behavior, dietary specialization, disease resistance, xenobiotic tolerance, and host range (6–11). Most horizontally acquired genes in insects originate from bacteria, including many from endosymbiotic bacteria (11, 12). A few genes of eukaryotic origin have also been reported in insects, including some from fungi, and plants, which likely contributed to phytophagy and xylophagy (10, 11, 13, 14). However, the contribution of eukaryotic genes, including genes originating from arthropods and other metazoans, to the repertoire of horizontally acquired genes in insects is poorly documented. Here, we performed a comprehensive analysis of HGT in a family of dipterans (Sciaridae) by detecting foreign genes of prokaryotic origin, but also those originating from other eukaryotes, including metazoans.

The family Sciaridae, commonly known as dark-winged fungus gnats, comprises small nematocerous flies with a cosmopolitan distribution and an ecological association with humid terrestrial habitats, where larvae predominantly inhabit soil, decaying organic matter, fungal substrates, and plant tissues (Figure S1). Larval feeding habits are primarily saprophagous or mycophagous, contributing to decomposition and nutrient cycling, although several species have evolved phytophagous habits and are recognized as significant pests in greenhouse and mushroom production systems (15, 16). We focused on fungus gnats because, in addition to harboring a tryptophan indole-lyase (TIL) gene of bacterial origin (17), some species of the *Bradysia* genus contain several cytochrome P450 (CYP) genes acquired from non-insect arthropods or fungi (18), as well as alternative oxidase (AOX) genes likely deriving from oomycetes (19). In addition, terpene synthase (TPS) genes likely acquired from acariform mites were described in multiple *Bradysia* and other Sciaridae species (20). Intriguingly, these sciarid genes also have close homologs in springtails, and they are generally absent in other metazoans (18–20). Furthermore, a recent study provided strong support for the direct transfer of a fatty acid desaturase gene from springtails to Sciaridae (21). Springtails are wingless hexapods phylogenetically positioned outside insects, forming part of the non-insect hexapod assemblage alongside Protura and Diplura (Figure S1) (22). They share many habitats and resources with sciarid larvae, often grazing on fungi and being abundant in organic litter and decaying wood. Previous analysis of HGT events in two springtail species, *Folsomia candida* and *Orchesella cincta*, unveiled hundreds of foreign genes, mostly acquired from bacterial and fungal donors (23, 24). These genes are enriched for carbohydrate-active enzymes (CAZymes), among other features, and only 30% are conserved between the two species, suggesting that they play distinct roles in their adaptation (24). Together, the indication of relatively high HGT permeability in Sciaridae and springtails, combined with their promiscuous overlap in specific ecological niches, prompted us to perform a comprehensive analysis of HGT in Sciaridae, with particular attention to genes shared with springtails. Our study reveals dozens of genes of bacterial and fungal origin in sciarids, as well as extensive HGT between sciarids and springtails, which likely contributed to the adaptation of both arthropod groups to their shared ecological niche.

## Results

### Extensive HGT events in Sciaridae

We used a multipronged approach to detect genes acquired from non-metazoan sources and from springtails in two sciarid genomes (*Pseudolycoriella hygida* and *Bradysia coprophila*) (Table S1.A) (Figure 1). We first assigned a last common ancestor (LCA) to all predicted proteins from the two species using a pipeline largely inspired from (13) (see methods). We clustered all proteins with a non-metazoan LCA at 50% identity and submitted them for phylogenetic analysis. This yielded 525 trees whose topologies suggest potential HGT events, as these proteins are more closely related to non-insect homologs and lack insect homologs in the local phylogenetic context. Interestingly, many of the predicted non-metazoan proteins were in fact most closely related to springtail homologs. We therefore performed the same search on the predicted proteins from four annotated springtail genomes (*Folsomia candida*, *Orchesella dallaii*, *Orchesella cincta*, and *Allacma fusca*) (Table S1.A), which allowed us to discover 36 new clusters whose phylogenies support HGT events between sciarids and springtails. Finally, a best-hit approach using sciarid and springtail proteins as queries against Uniref90, followed by protein clustering, enabled us to detect another 149 clusters whose phylogenetic trees indicate HGT between the two taxa (Figure 1).

**Figure 1:**
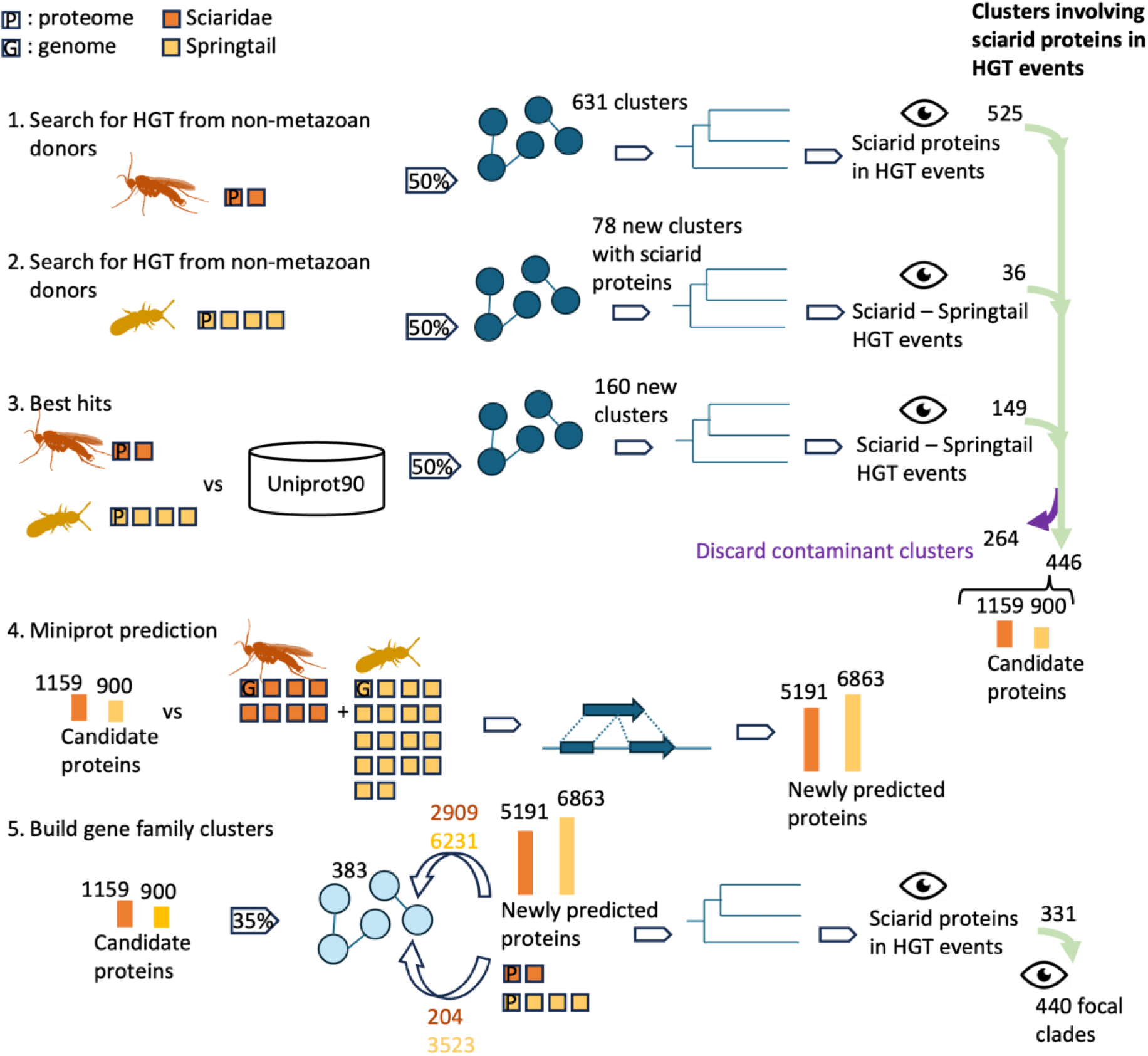
Detection of HGT events in sciarids. The figure illustrates the main methodological steps used to identify HGT candidates and validate them (steps 1-3), annotate homologous sequences using Miniprot (step 4), and build gene-family-level protein clusters and identify independent HGT events (step 5). Dark and light orange silhouettes, values, and datasets correspond to sciarid and springtail features, respectively. Each square represents an independent dataset, either a proteome (P) or a genome (G). Percentages in arrows indicate the threshold used for clustering in percent identity. Network cartoons represent protein clusters. Histograms represent a collection of proteins (not to scale). Eye silhouettes indicate manual examination of phylogenetic trees.

Inspection of the distribution of candidate transferred genes along sciarid genome scaffolds led us to exclude 264 clusters whose genes were concentrated in a 3.14 Mb region at the beginning of the *P. hygida* chromosome CM055403.1, as multiple lines of evidence suggest this region derives from contaminating bacterial DNA (Figure S2, Table S2). The remaining genes are unlikely to result from contamination because they are scattered across all genome scaffolds, and many are present in the two sciarid species (see below). In total, we thus selected 446 trees showing at least one HGT event involving sciarid proteins, representing 1159 candidate proteins from the two Sciaridae genomes and 900 from the four springtail genomes (Figure 1).

Although the tree topologies strongly suggest HGT, they also revealed complex relationships between sciarid and springtail candidate proteins in many instances, hampering straightforward inference of HGT scenarios. To place these events in a broader evolutionary context, we searched for homologs of the candidate proteins in the genomes of an additional 10 Sciaridae species and 18 springtail species with no publicly available gene annotations (Table S1.B). We then grouped the previously identified candidate proteins into clusters at an approximate gene-family level (35% identity threshold), yielding 383 clusters, to which we appended newly found homologs, previously unselected reference springtail and sciarid proteins, and representatives of their diversity among UniProt homologs. Finally, we submitted each cluster to phylogenetic analysis, and manual examination of the 383 resulting trees revealed that 331 involve Sciaridae proteins in at least one HGT event. Among many of these trees, we found the transferred proteins distributed across several subclades, indicating that some gene families were transferred multiple times independently. In total, we counted as many as 440 subclades, which we refer to as “focal clades” and which reflect independent HGT events involving sciarid genes (Figure 1).

While incongruences between gene and species phylogenies may result from HGT, they may also arise from long-term vertical inheritance of genes, alongside multiple losses across many lineages, including all non-dipteran insect orders and all non-sciarid dipteran lineages (Figure S1) (25–27). This second scenario would imply that the genes interpreted as resulting from HGT were in fact present in the common ancestor of springtails and sciarids, and that their genetic distance should be similar to that of vertically inherited, core arthropod genes. We thus compared the distribution of synonymous distances (dS) calculated for each focal clade to that of BUSCO genes. In agreement with the HGT scenario, the vast majority of candidate transfers (331 out of 352) showed pairwise dS values (mean = 1.75) that were well below the mean dS of BUSCO genes, which in most instances could not be accurately calculated because of saturation (dS = 9.9) (Figure S3). The dS of the remaining 21 candidates was also saturated and fell within the distribution of BUSCO gene dS. While this result may indicate vertical inheritance of these genes from a springtail-sciarid ancestor (and loss from many insect lineages), it may also result from more ancient HGT events, or from accelerated evolution of these genes after transfer. Given that the local phylogenetic context of these genes lacks metazoan genes other than those of springtails, we included them in our count of HGT events.

### Non-metazoan-to-Sciaridae HGT

We next inferred a refined scenario for each of the 440 HGT events based on a detailed inspection of each phylogenetic tree topology. We first focused on 67 focal clades that lack springtail or any metazoan proteins other than Sciaridae in their local phylogenetic context. Applying stringent HGT validation criteria, we found that 51 out of these 67 focal clades most likely derive from a non-metazoan donor (Figure 2.B, Table S3, Figure S4). These 51 focal clades fall into 51 distinct trees. In these trees, the sister branch of the focal clade is strongly supported (UF bootstrap >=90%) as a non-metazoan taxon, which is confirmed by a second highly-supported node when traversing the tree towards the root, while maintaining high purity of the candidate donor taxon (Figure 2.A). 18 HGT events involve a bacterial donor, whereas 33 involve eukaryotic donors distributed across fungi (31), SAR (1), and Viridiplantae (1) (Figure 2.B, Table S3, Figure S4). In addition to these clear non-metazoan-to-sciarid HGT events, we found 16 trees in which the sister branch of the focal clade is highly supported as non-metazoan. Yet, the candidate donor taxon is not validated by upward nodes. However, the larger clades in which these focal clades are embedded lack insect or metazoan sequences, suggesting they derive from non-metazoan-to-sciarid HGT events, hence their selection for subsequent analysis as HGT events of unclear non-metazoan origin (Figure 2.A, Table S3, Figure S4). In total, the 67 selected focal clades resulting from non-metazoan-to-sciarid HGT comprise 478 proteins from 12 sciarid species. Importantly, we found that 62 of these clades comprise members from at least two distinct species (and up to 11), further arguing against contamination artifacts (Figure 2.C). In addition, most focal clades span several sciarid subfamilies, including for about half of them (n=34) the most basal sciarid subfamily (Sciarinae). This suggests that the underlying HGT events occurred in a sciarid common ancestor, at least 82 million years ago (MYA) (16). However, we cannot rule out that independent HGT events occurred. In addition, several focal clades include more than one protein from a single species, suggesting multiple post-HGT gene duplications, although unequal genome assembly completeness and gene-prediction efficiency across species likely affect these patterns.

**Figure 2:**
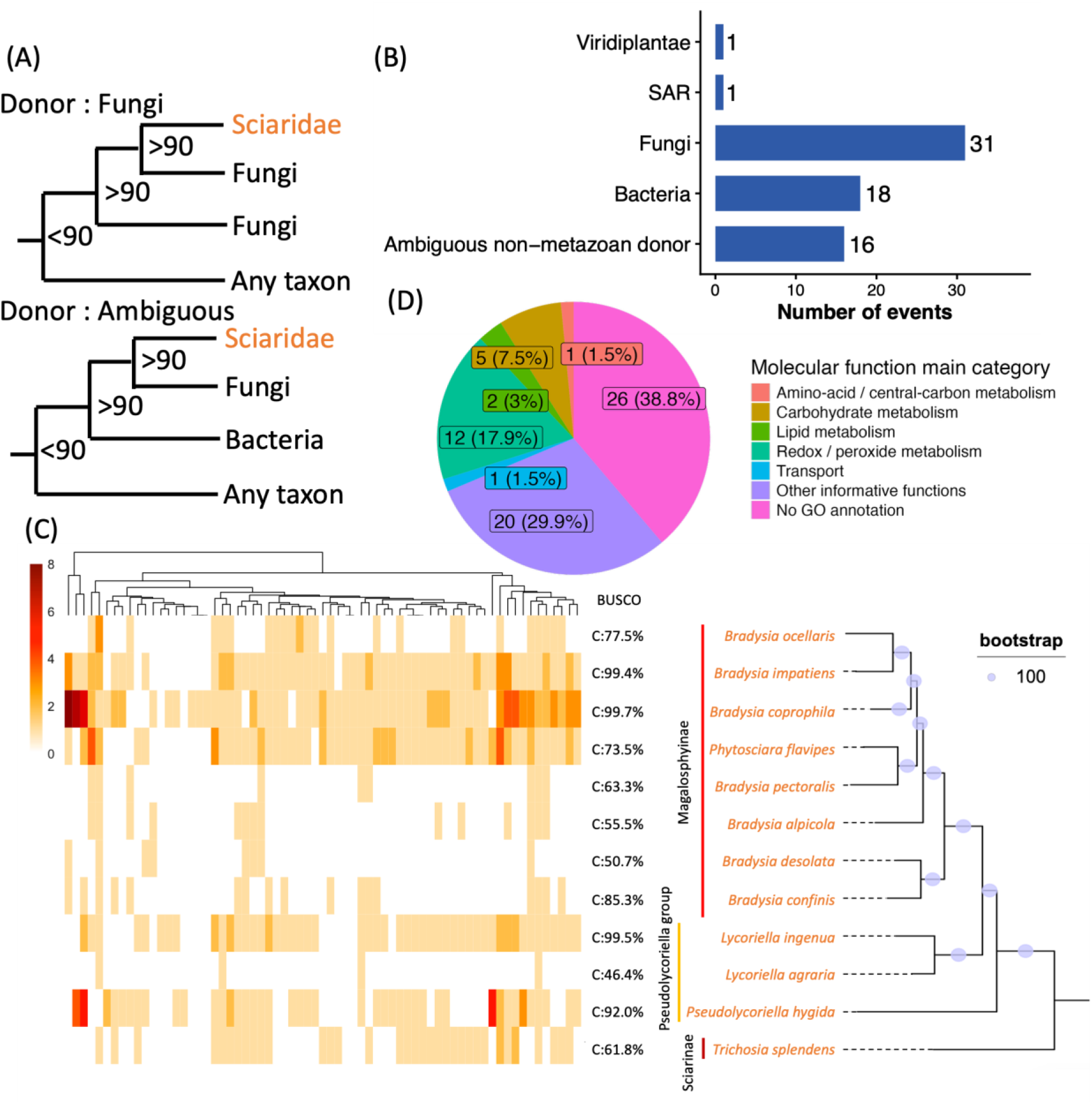
Non-metazoan genes in sciarids. (A) Schematic topological categories indicating validated donor (top, fungi taken as exemple) and ambiguous non-metazon donor (bottom). (B) Distribution of HGT events per donor taxon. (C) Gene copy number in sciarid genomes per HGT event (left), BUSCO genome scores (middle), and sciarid species tree (right). (D) GO distribution into molecular function categories per HGT event

Finally, we examined the putative functions of the non-metazoan-derived focal clades. We found that their gene ontologies were primarily associated with metabolic functions, including carbohydrate, lipid, amino acid, and central carbon metabolism, although many remained unannotated (Figure 2.D).

### Multiple, successive, and parallel transfers between Sciaridae, springtails and other metazoans

We next analyzed the remaining 373 focal clades of sciarid proteins that contain springtail or other metazoan proteins in their local phylogenetic context. Using the same donor-taxon inference criteria as above, we identified 266 focal clades that are embedded in a clade of springtail homologs, likely reflecting a springtail-to-sciarid HGT. In contrast, we inferred an HGT in the opposite direction for another 20 cases, in which a clade of springtail proteins is embedded in a larger clade of sciarid proteins (Figure 3.A, Figure 3.B). Remarkably, while 207 trees support a single HGT event between sciarids and springtails, 59 instead show evidence of multiple HGT events between the two taxa. In some of these trees, the focal clades were nested, suggesting successive HGT events of the same gene in the same or opposite direction. In the other trees, focal clades are distantly related, suggesting parallel HGT events, once again in the same or opposite direction (Table S3, Figure S4). In total, we inferred up to 11 putative HGT events between sciarids and springtails within a single tree (Table S3, Figure S4). In addition to these clear, oriented HGT events, 66 focal clades showed a highly supported springtail sister clade, suggesting HGT between the two taxa, but the rest of the tree could not resolve the direction (Figure 3.A, Table S3, Figure S4).

**Figure 3:**
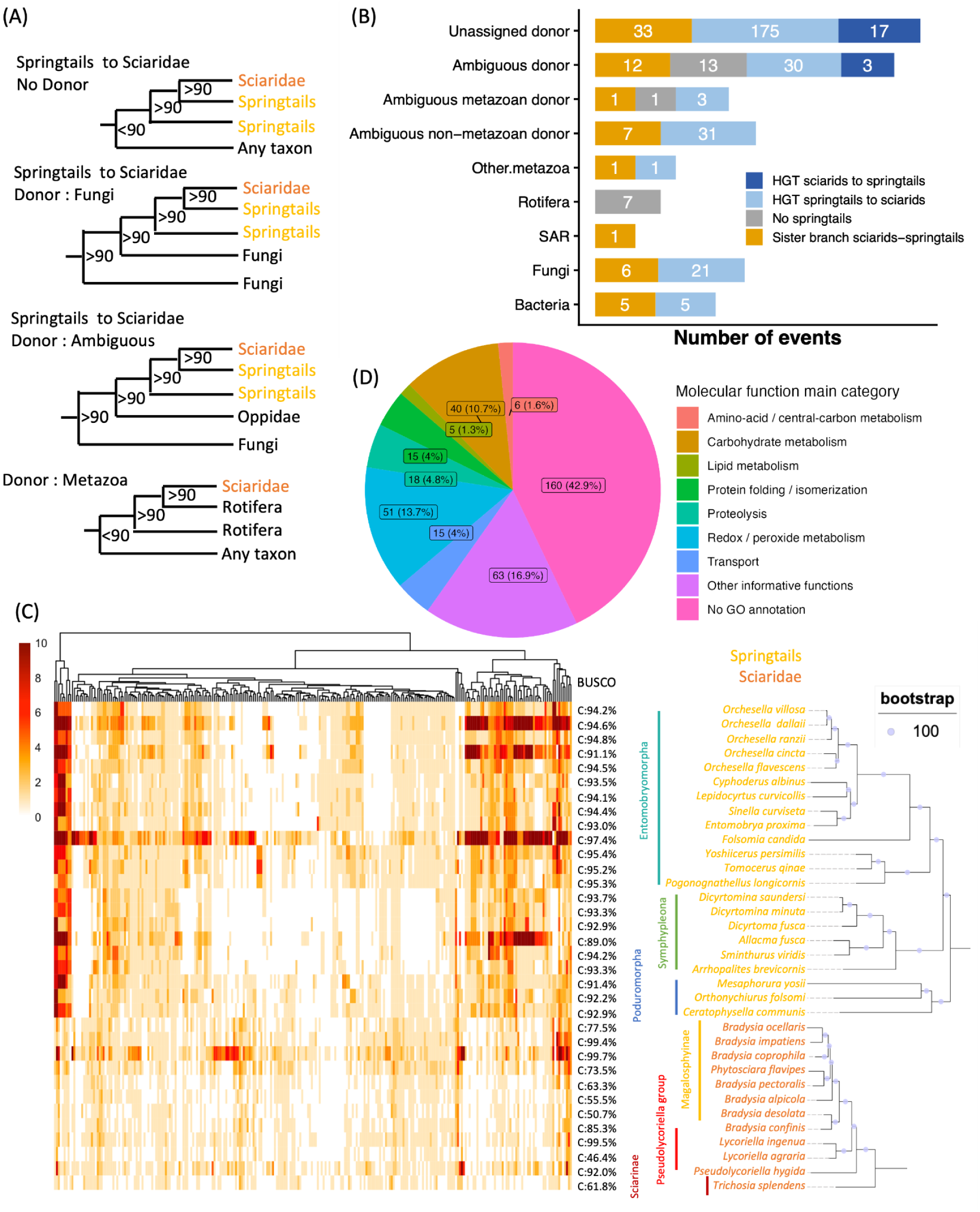
Metazoan-to-sciarids HGT events. (A) Schematic topologies illustrating various categories of HGT events. (B) Distribution of HGT events per donor taxon and per type of transfer between sciarids and springtails (not to scale). All categories represent two successive HGT events except for “Unassigned donor” and “No springtails”. (C) Gene copy number in springtail and sciarid genomes for each springtail-to-sciarid HGT event (left), BUSCO genome scores (middle), and springtails and sciarid species tree (right). (D) GO distribution into molecular function categories per HGT event.

In total, we selected 352 putative HGT events between sciarids and springtails, involving 3502 sciarid proteins and 7160 springtail proteins. Importantly, 28 of these events involve at least two sciarid species, 30 involve at least two springtail species, and 293 involve both, arguing against contamination in both taxa. For several oriented HGT events, we observed multicopy representation in recipient-taxon species, suggesting many post-HGT gene duplications (Figure 3.C, Figure S5). As the most extreme case, we found as many as 25 copies of a springtail-derived fatty acid desaturase protein in *B. coprophila*. Similar to the distribution of non-metazoan genes across sciarids, a large number (149 out of the 352) of the springtail-derived genes were shared between Sciarinae and non-Sciarinae, suggesting that the underlying HGT events occurred in a sciarid common ancestor. We also measured the identities between sciarid and springtail sequences within each focal clade. Consistent with ancient transfers, the mean of the highest identities was 67%. However, the values ranged from 38,34% to 88,81%, suggesting that more recent transfers also occurred (Figure S6, Table S4).

Remarkably, we found that at least 76 of the 352 selected clades involving an HGT event between sciarids and springtails (Figure 3.B) showed local embeddings depleted of arthropod and other metazoan proteins. Instead, they contain homologous proteins from more distant lineages. These trees provide evidence for successive HGT events, including a primary HGT event from non-metazoan donors to springtails, followed by secondary HGT between springtails and sciarids. Among these cases, 10 involved a bacterial donor, 28 involved a eukaryotic donor (1 SAR and 27 Fungi), and 38 involved an ambiguous non-metazoan donor (bacteria, fungi, Viridiplantae, or SAR). We also identified 2 events involving a metazoan donor (Crustacea and Mites), 5 events involving an ambiguous metazoan donor and 45 with an ambiguous origin (metazoan or non-metazoan) (Table S3, Figure S4).

Remarkably, we also identified 2 trees that support yet another scenario in which a non-metazoan-to-sciarid HGT and a sciarid-springtail HGT concern distinct focal clades that are paraphyletic (Figure S4). Such topologies strongly suggest that distinct genes from the same family were transferred through parallel HGT events from non-metazoans to sciarids and between sciarids and springtails.

In addition to HGT events involving sciarids and springtails, we identified 7 focal clades resulting from rotifer-to-sciarid HGT. We also detected 14 focal clades with an ambiguous donor that could be either a metazoan (Rotifera, Sarcoptiformes and Trombidiformes (Mites), Ostracoda, Tardigrada) or a non-metazoan (bacteria, fungi, and SAR) (Figure 3.B, Table S3, Figure S4).

Finally, analysis of the putative functions of the focal clades of metazoan origin revealed that, as for genes of non-metazoan origin reported in the previous section, their gene ontologies were primarily associated with metabolic functions, including carbohydrate, lipid, amino acid, and redox/peroxide metabolism, although about half of the clades remained unannotated (Figure 3.D).

### Springtail-derived genes in sciarids’ germline-restricted chromosomes

In addition to the chromosomes assembled in reference genome assemblies, which are sexual or autosomal, many Sciaridae species harbor germline-restricted chromosomes (GRCs) that are eliminated in somatic cells. Recently, the two GRCs of *B. coprophila* were assembled and found to contain more than fifteen thousand genes (28). The functional status of these genes remains enigmatic: although they evolve under strong selective constraints, most have yet to show transcriptional evidence (29). Many GRC genes have homologs on other chromosomes, yet thousands are restricted to GRCs (28). To fully address the extent of HGT between sciarids and springtails and to assess the involvement of GRCs in these events, we independently searched for HGT between the two taxa in the *B. coprophila* GRCs using the same best-hit approach as above. Out of 23 049 GRC proteins we predicted, 449 had a springtail best hit, including 299 that did not group with any of the clusters established above from the somatic HGT set. Applying the same criteria as above, we selected 110 that clustered with springtail proteins at 50% identity, corresponding to 82 clusters that we subsequently merged using a 35% identity threshold, resulting in 79 clusters. Phylogenetic trees were constructed as above for the 70 clusters that comprised proteins from taxa other than springtails and sciarids. Among the 70 phylogenetic trees, 50 showed strong support for HGT between sciarids and springtails: 28 showed a clear HGT from springtails to sciarids, while 4 showed a transfer from sciarids to springtails. The remaining 18 trees did not allow us to infer the direction of these events (Table S5, Figure S7). As observed for HGT in the somatic chromosomes and in agreement with the HGT scenario, the vast majority of GRC genes involved in horizontal transfer had a springtail-sciarid dS falling well below that of BUSCO genes (Figure S3).

### Functional analysis

To assess whether the transferred elements represent functional, selectively maintained genes rather than neutral sequences, we estimated dN/dS (ω) for each focal clade. For this, we retrieved the coding sequences from each focal clade along with their respective springtail homologs and computed codon alignments to calculate branch-specific ω values. Across all focal clades, mean ω was well below 1 in both the Sciaridae and springtail lineages (0.159 and 0.161, respectively). The vast majority of focal clades had a mean ω below 0.3 in both lineages (94.4% in Sciaridae and 94.2% in springtails) and nearly all were below 0.5 (99.4% and 98.4%, respectively). Among focal clades with at least four branches in each lineage (n = 208), the ω distributions did not differ significantly between sciarids and springtails in the large majority of cases (195/208, 93.8%; Wilcoxon rank-sum test, FDR > 0.05); only nine clades showed significantly higher ω in Sciaridae and four in Springtails (FDR < 0.05). Thus, HGT events were apparently not associated with a change in selection regime. Candidate transferred genes located on GRCs showed similarly low ω, with mean values of 0.188 in Sciaridae and 0.111 in springtails, and with 88.2% and 95.6% of focal clades below 0.5, respectively. Overall, these results indicate that transferred genes evolve predominantly under purifying selection in both lineages, and thus are functionally constrained (Figure S8).

We also searched for transcriptional evidence of the non-GRC HGT-derived sciarid genes. For this, we used NCBI transcriptome shotgun assemblies (TSA) of *Pseudolycoriella hygida* and *Bradysia coprophila* to search for transcripts with high identity (>95%) to each coding sequence involved in HGT over at least 30% of their length (qcovs). Among the 440 HGT events, 416 involve at least one gene from one of these two species, the remaining involving genes from other species for which TSA data is not available. 402 of the 416 events involve at least one gene matching at least one transcript. Furthermore, 279 out of the 402 HGT events involve both species, and 259 of them are supported by at least one transcript in both species. Overall, these findings are consistent with the vast majority of horizontally acquired proteins being expressed and functional in sciarids (Table S6, Table S7).

We further used RNA-seq libraries available for *P. hygida* to examine the transcriptional abundance, expressed as transcripts per million (TPM), of horizontally transferred genes across developmental stages and larval tissues (30). HGT-derived functions displayed marked differences in their expression profiles across biological contexts. The clearest pattern involved carbohydrate-active HGTs, which were strongly expressed in whole larvae and particularly in the two larval gut datasets, with median TPM values of 31.0 in whole larvae, 61.9 in 12-day-old gut, and 29.4 in fourth-instar gut. In contrast, their median expression was close to zero in salivary glands, brain, fat body, and embryos, and remained comparatively low in pupae and adults (Figure 4). Peptidase HGTs showed a related, although less pronounced, larval and gut-associated expression profile. By contrast, redox-associated and protein-folding/isomerase HGTs were more broadly expressed across developmental stages and larval tissues, consistent with their deployment in a wider range of physiological contexts.

**Figure 4.**
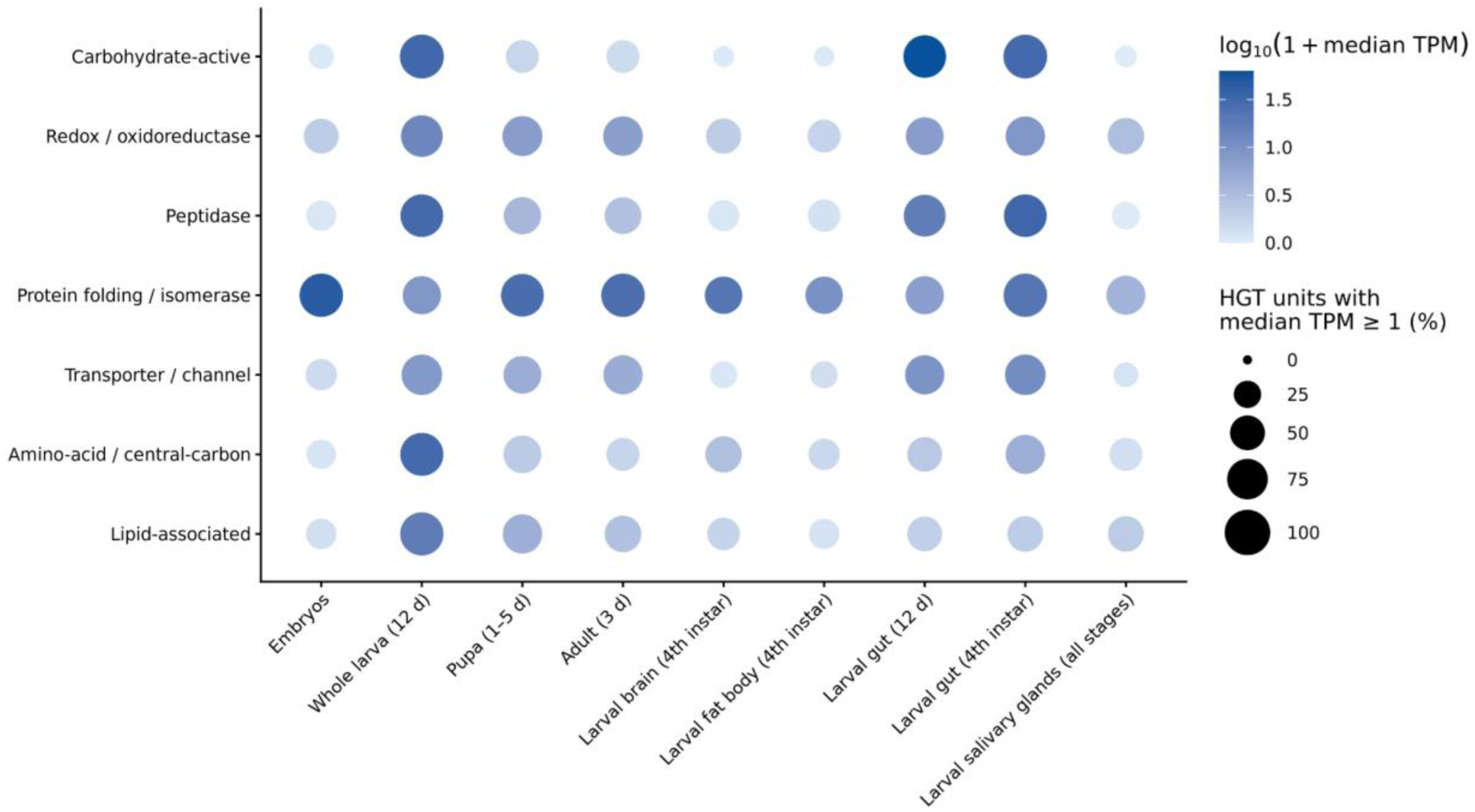
Tissue- and developmental-context expression of HGT-derived functional categories in *Pseudolycoriella hygida*. Expression is shown across whole-body developmental stages and larval tissues. Bubble plots summarize TPM expression of HGT genes assigned to the main molecular-function categories across whole-body developmental samples and larval tissues. Bubble color indicates log10(1 + median TPM) across HGT genes within each functional category, from pale to dark blue for lower to higher expression, while bubble size represents the percentage of HGT genes with a context-median TPM ≥1.

We specifically examined the expression profile of two *P. hygida* HGT-derived proteins (KAJ6634914.1 and KAJ6620840.1) that cluster with a non-metazoan-derived carbohydrate-active enzyme from the springtail *F. candida*, whose capacity to degrade an analog of L-arabinofuranose, a major component of hemicellulose and pectin, was experimentally validated (31). Interestingly, the associated phylogenetic tree (XP_037025833.1) indicates that the two *P. hygida* genes derive from independent springtail-to-sciarid HGT events. Nevertheless, both show a pronounced transcriptional bias in larvae, particularly in the gut (Figure S9), suggesting that gut-specific regulatory elements may have been co-transferred with the coding sequence. Because this molecular function likely involves degrading plant tissues in the gut in both taxa, it further supports the idea that sciarid and springtail HGT proteins can fulfill similar functions linked to adaptation to their ecological niche.

We conducted GO enrichment analyses separately for *Bradysia coprophila* and *Pseudolycoriella hygida*. For each species, we used the complete predicted proteome as the reference set, and the HGT candidate set was used as the foreground. The analysis identified 45 Biological Process and 55 Molecular Function terms passing the enrichment filter in *B. coprophila*, and 16 and 28 terms, respectively, in *P. hygida*. The principal functional categories comprised carbohydrate and polysaccharide metabolism, oxidoreductase and redox cofactor functions, proteolysis, protein folding, amino acid and central carbon metabolism, lipid metabolism, and transport. Carbohydrate-associated terms included glucan, xylan, and arabinan catabolism, together with β-glucosidase, xylan β-xylosidase, α-L-arabinofuranosidase, and polygalacturonase activities, indicating an overrepresentation of enzymes capable of hydrolyzing structurally diverse polysaccharides, including major components of plant cell walls. Redox-associated terms included oxidoreductase and catalase activities, hydrogen peroxide metabolism, and FAD-, FMN-, and NAD(P)-binding functions, indicating an overrepresentation of oxidation–reduction reactions and peroxide-control functions. Enrichment for peptidase activities, peptidyl-prolyl isomerization, ATP-dependent protein folding, lipid metabolism, and membrane transport further identified functions related to protein turnover, proteostasis, membrane-associated metabolism, and solute transport. Together, these results show that the sciarid HGT candidate repertoires are enriched for biochemical functions involved in degrading and transforming complex organic substrates and in cellular redox and protein homeostasis (Figure S10).

## Discussion

While HGT events have been reported in many metazoans, most correspond to inter-domain HGT (6, 11, 12, 32). It is unclear whether this pattern has biological underpinnings or simply reflects our limited ability to distinguish HGT from vertical inheritance as phylogenetic distances decrease (33). Here, the combination of protein taxonomic assignment, distance-based, and phylogenetic approaches allows us to infer a total of 440 HGT events in sciarid flies, most of which likely correspond to intra-domain transfers, that is, direct HGT between sciarids and springtails (Figure S11).

First, consistent with earlier studies showing that insects can acquire genes from non-metazoan sources, 67 of the 440 HGT events we report imply genes from fungi (n=31), bacteria (n=18), and other non-metazoans (n=18). Most of these transfers are relatively ancient, as 62 of them involve genes found in at least two sciarid species. Compared to a set of 218 insects that underwent an average of 3.4 HGT events from non-metazoans (11), sciarid flies have been particularly prone to acquiring such genes, being surpassed only by the whitefly *Bemisia tabaci*.

Next, we provide multiple lines of evidence supporting 352 HGT events between sciarid flies and springtails. We show that sciarid proteins involved in these HGT events are most closely related to springtail homologs, and that such homologs are virtually absent in other metazoans. This topology could in principle result from incomplete lineage sorting (ILS), differential gene loss, or HGT (26, 27). However, protein databases include a wide diversity of insect species, arguing against ILS as the main explanation. Furthermore, differential gene loss would imply conservation of ancestral genes from Hexapoda to the emergence of Sciaridae (about 82 MYA, (16)), in parallel with loss in all major insect lineages, including Diptera except Sciaridae. We consider this coincidence with springtail homologs unlikely, especially for so many genes. In addition, the vast majority of genes horizontally transferred between springtails and sciarids show low dS values compared to vertically inherited ones (BUSCO set), indicating they began diverging well after the sciarid-springtail split. Altogether, we thus consider HGT the most plausible explanation reconciling all these observations. While our phylogenetic analyses are consistent with direct transfer between springtails and sciarids, one could invoke independent horizontal acquisition of members of the same gene families in springtails and sciarids from extinct or undersampled non-metazoan (ghost) lineages (34) (Figure S12). This scenario is difficult to rule out for the 66 non-oriented springtail-sciarid HGT events. Given that direct HGT between metazoans has so far rarely been documented, two non-metazoan-to-metazoan HGT events may be seen as less unlikely than a single metazoan-to-metazoan transfer. However, we consider the scenario involving ghost lineages unlikely to explain the remaining 272 oriented events, as it would require at least one metazoan-to-non-metazoan transfer in addition to two non-metazoan-to-metazoan HGT. Our study, together with a previous report of multiple mantid-to-nematomorph HGTs (35), suggests that HGT between metazoans may be relatively common in some clades and calls for a broader appraisal of this type of transfer.

The finding of massive HGT between springtails and sciarids raises questions about the molecular mechanisms underlying these transfers. In this regard, previous work interpreted the presence of 454 genes originating from Cecidomyiidae flies in the GRCs of *B. coprophila* as resulting from introgression of entire cecidomyiid chromosomes into a sciarid ancestor through a single hybridization event. This scenario was judged puzzling because, although cecidomyiids and sciarids fall within the same dipteran superfamily (Sciaroidea), they diverged several dozens of MYA, and hybridization between such divergent insects has never been reported (28). However, the alternative scenario involving recurrent HGT events was deemed less likely because it would require a high number of transfer events. Here, we favor HGT over hybridization because the latter would have had to occur between sciarids and springtails, which diverged 450 MYA (36), i.e., much more deeply than sciarids and cecidomyiids. In addition, hybridization would not easily account for the fact that genes acquired from springtails are scattered throughout the sciarid genomes (GRCs, sex chromosomes and autosomes), and that HGT occurred in both directions between sciarids and springtails, sometimes through successive events involving the same gene family. The detailed HGT mechanisms remain however mysterious. Several hypotheses have been proposed to explain HGT between metazoans, some involving vectors such as viruses or extracellular vesicles, but all lack experimental support (37–39). Here, 149 out of the 352 HGT between springtails and sciarids involve gene families for which at least one member is present in the two most distantly related lineages of Sciaridae (Sciarinae and non-Sciarinae). Only 36 HGT events are restricted to a single species. Thus, while the heterogeneous completeness of the sciarid genomes included in this study warrants caution in interpreting the age of HGT events, many likely occurred in the last common ancestor of Sciaridae. Whatever transfer mechanism(s) are at play here, the presence of springtails and sciarids within shared ecological niches has likely favored HGT between them, as shown for HGT among bacteria (40) and for TE transfers among metazoans (41).

The finding that most horizontally acquired genes are expressed and evolve under strong purifying selection strongly suggests that they are functional and that HGT has likely had a substantial impact on sciarid biology. Remarkably, genes acquired directly from non-metazoan sources, especially bacteria and fungi, are enriched in the same functional categories previously reported for non-metazoan genes in springtails, namely enzymes involved in carbohydrate and redox metabolism (23, 24). Our phylogenies lack springtail proteins for these genes, indicating that the two groups independently acquired similar biochemical capacities through HGT. Furthermore, we found that genes sciarids acquired from springtails, often through recruitment of the springtails’ own HGT repertoire, are also mainly associated with the same metabolic categories.

As proposed for springtails, carbohydrate-active enzymes could facilitate the utilization of plant- and fungal-derived polysaccharides, including lignocellulose, whereas oxidoreductases could participate in transforming lignin-derived phenolics, quinones, and other aromatic compounds (42, 43). Although the physiological and ecological roles of these genes remain largely untested (30), their annotations suggest that HGT repeatedly provided biochemical functions that may have conferred selective advantages for adaptation to fungus-rich, decomposing plant substrates in both taxa. Together, these findings suggest that springtails and sciarids may illustrate the intersection of the recently proposed concepts of xenotypes and epaktovars (44), whereby horizontally acquired genetic similarities coexist with independently evolved phenotypic similarities. Indeed, the independent acquisition of genes with overlapping predicted metabolic functions, together with the horizontal exchange of genes between the two groups, provides a potential genomic basis for such an evolutionary pattern. However, whether these overlapping metabolic repertoires translate into convergent physiological capabilities remains to be established.

Most transfers between the two groups occurred from springtails to sciarids (n=266), with only a few in the opposite direction (n=20). This asymmetry could reflect either a greater permeability of sciarids to foreign DNA or a greater propensity for springtail-derived genes to be beneficial and retained in sciarids than for sciarid-derived genes to be retained in springtails. The latter possibility is consistent with the earlier appearance of springtails in soil habitats (≈296 MYA; (45)) relative to sciarids (≈82 MYA; (16)). Springtails may therefore have donated genes that were both useful in this ecological niche and already adapted to a metazoan genomic background. This pattern could reflect how eco-evolutionary dynamics may shape HGT donor-recipient dynamics (5).

Interestingly, the ecological overlap between springtails and sciarids occurs primarily during the larval stage for sciarids, when both groups inhabit soil, leaf litter and rotten wood and graze on similar fungal resources. Consistent with a role for horizontally acquired genes in processing compounds found in these environments, expression patterns in *P. hygida* show that carbohydrate-processing and, to a lesser extent, proteolytic functions are particularly associated with the larval digestive system. These patterns support the hypothesis that HGT directly contributed to the digestive capabilities of *P. hygida* larvae in soil environments.

The recurrent acquisition and retention of similar HGT-derived functions in distantly related detritivorous hexapods suggest that comparable ecological conditions favored genes encoding similar metabolic capacities. Altogether, our findings indicate that HGT, operating repeatedly and through different forms of transfer, has driven functional convergence among ecologically overlapping detritivorous soil animals.

## Materials and Methods

### Identifying HGT candidates (steps 1-3 in **Figure 1**)

#### Search for HGT from non-metazoan donors

We searched for HGT events from non-metazoan donors independently in the proteomes of two sciarid species (*P. hygida* and *B. coprophila*) and four springtail species (*Folsomia candida*, *Orchesella dallaii*, *Orchesella cincta*, and *Allacma fusca*), retrieved from NCBI (Table S1). Predicted protein sequences were first clustered with MMseqs2 (46) at 70% sequence identity and 70% coverage (--cov-mode 1). We then compared cluster representative proteins against the UniRef90 database (release 2024_05), excluding Dipteran proteins, and inferred taxonomic origin using the MMseqs2 taxonomy module majorityLCA (47) with a 50% majority threshold. Sequences assigned to non-metazoan origins and for which ancestry could be resolved deeper than genus level were retained as preliminary HGT candidates. We then compared these candidates against the NCBI nr database using a stricter majorityLCA threshold of 80% to confirm taxonomic assignments.

We removed potential transposable elements by comparing candidate sequences against the TransposonPSI library (https://transposonpsi.sourceforge.net/) using BLASTp from the BLAST+ suite (48). We then clustered the remaining proteins with MMseqs2 using relaxed thresholds (sequence identity ≥ 50% and coverage ≥ 50%) and selected representative sequences from each resulting cluster.

#### Search for best hits between springtail and sciarid proteins

We also directly searched candidate sciarid genes involved in HGT events between sciarids and springtails using a best-hit approach. For this, we independently compared the two sciarid proteomes and the four springtail proteomes to the UniRef90 database using MMseqs2. We retained the sciarid proteins with best UniProt hit against springtails and the springtail proteins with best hit against sciarids. We then clustered all selected sciarid and springtail proteins using MMseqs2 with a minimum sequence identity of 50% and a minimum coverage of 50% (--cov-mode 1), and retained proteins belonging to clusters containing at least one sciarid and one springtail protein for further analyses. We used the same approach to identify the genes involved in HGT events between sciarids and springtails in the two *B. coprophila* GRCs. As the *B. coprophila* GRC protein sequences annotated in previous work (28) were not publicly available, we firstly performed gene prediction on the GRC1 and GRC2 chromosomes with Helixer (v0.3.6) using the “invertebrate” mode (49).

#### Construction of phylogenetic trees

We used representative sequences from each selected cluster as queries in MMseqs2 similarity searches against multiple datasets, including UniRef90 proteins from the focal taxon, UniRef90 proteins from putative donor taxa, UniRef90 eukaryotic proteins excluding both the donor and query clades, and comprehensive prokaryotic and viral protein databases. From each search, we retained the ten best hits, together with twenty sequences selected to maximize taxonomic diversity (filterresult --diff 20 & filterdb --extract-lines 10), thereby ensuring broad representation of potential homologs across lineages.

We retrieved all selected sequences in FASTA format and aligned them using MAFFT (v7.475) (50) in local alignment mode. Poorly aligned sequences were removed with trimAl (v1.4) (51) using parameters resoverlap = 0.6 and seqoverlap = 60, followed by a second alignment step with MAFFT. Poorly informative positions were subsequently removed from the alignments using the gappyout option in trimAl. We then inferred the phylogenetic relationships among each cluster with IQ-TREE (v3.1.3) (52) using the optimal model selected by ModelFinder (53) and ultrafast bootstrap with 1,000 replicates, and we manually examined each tree to identify topologies that supported probable HGT events from non-metazoans to sciarids or between sciarids and springtails. We first selected the candidate clusters identified through the majority LCA approach applied on sciarids (n=525), followed by those involving other sciarid proteins identified through the same approach applied to springtails (n=36), and last, those involving yet other sciarid proteins identified with the best-hit approach (n=149). In total, these clusters comprised 1159 candidate proteins from sciarids and 900 from springtails.

#### Identification of likely contaminant genomic regions

The genome of *Pseudolycoriella hygida* was analyzed in R using the Biostrings package. The sequence corresponding to scaffold CM055403.1 was extracted from the reference genome and divided into consecutive 20,000-bp windows. For each window, the G+C content was calculated using the letterFrequency() function from the Biostrings package, with as.prob = TRUE to obtain the frequency of G and C nucleotides for each windows and then visualisation was obtained using ggplot2.

The genomic distribution of 525 candidate proteins identified as potential horizontal gene transfer (HGT) candidates was assessed using the *Pseudolycoriella hygida* genome. The GFF annotation was imported using rtracklayer, and genomic coordinates of the candidate proteins were retrieved from the annotation. Data manipulation and filtering were performed using dplyr and stringr. Scaffold lengths were obtained from the corresponding genomic features in the GFF file. To allow comparison among scaffolds of different lengths, the genomic position of each candidate protein was normalized to the length of its corresponding scaffold by dividing its start coordinate by the total scaffold length. Normalized positions therefore ranged from 0 (scaffold start) to 1 (scaffold end). The distribution of candidate proteins across scaffolds was visualized using ggplot2, with relative genomic position plotted against scaffold identity.

Transcriptomic analyses were performed using one TSA master assembly of *Pseudolycoriella hygida* (accession GICH00000000.1). Protein sequences were searched against the corresponding TSA datasets using tBLASTn from the BLAST+ suite.

Illumina paired-end reads from sequencing run SRR18645815 and SRR18645814 (FASTQ files) were aligned to the reference genome using BWA-MEM (https://arxiv.org/abs/1303.3997) and the long-read PacBio Sequel data (SRR18684862) were aligned to the reference genome using minimap2 (54) with the map-pb preset. Prior to alignment, the reference genome was indexed using BWA (55) and SAMtools (56), and a sequence dictionary was generated using GATK (57). The resulting SAM file was converted to BAM format using SAMtools, sorted by genomic coordinates, and indexed for efficient access to genomic regions. The final coordinate-sorted and indexed BAM files were used for downstream analyses and visualization in Integrative Genomics Viewer (IGV) (58).

### Identification of homologous proteins in additional genomes (step 4 in Figure 1)

To identify sequences homologous to candidate horizontally transferred proteins in additional sciarid and springtail genomes that lack protein annotation, we first compiled a dataset containing all 2,059 sciarid and springtail candidate proteins recovered from the clusters selected through steps 1-3. We then screened genome assemblies from eight sciarid and 18 springtail species (Figure S1) with Miniprot (59) using protein-to-genome alignments generated in translated mode (--trans). We subsequently processed predicted gene models with AGAT (https://github.com/NBISweden/AGAT) to resolve overlapping gene predictions and retain only the longest isoform for each locus. We then extracted protein sequences from the curated annotations using gffread (60) and removed redundant protein sequences with SeqKit (61), yielding a final set of 5,191 newly predicted proteins from sciarids and 6,863 from springtails.

### Building gene-family clusters and final phylogenetic trees (step 5 in Figure 1)

To analyze HGT events at an approximate gene-family level, we first grouped candidate proteins from reference genomes using MMseqs2 with a 35% sequence-identity threshold and 50% minimum coverage, yielding 383 clusters. We next enriched these clusters with the ClusterUpdate module of MMseqs2, using the same parameters to include sequences from the newly predicted proteins (2,909 from sciarids and 6,231 from springtails) as well as proteins from the six reference proteomes that met the clustering conditions but were not selected in the initial detection steps (204 from sciarids and 3,523 from springtails). We then retrieved the diversity-representative UniRef90 proteins collected above to analyze the clusters established at 50% identity and, after removing redundancy as well as sciarid and springtail proteins, we appended them to the respective 35% identity cluster of their representative protein. We finally aligned and trimmed the sequences from each consolidated 35% identity cluster as described above, and used the resulting alignments to build phylogenetic trees as described above.

### Species tree

We inferred the species phylogeny using a phylogenomic approach based on conserved single-copy orthologs. Genome assemblies were first screened with BUSCO (62) using the Arthropoda dataset (arthropoda_odb12) in genome mode to identify conserved single-copy orthologous genes. Orthologous BUSCO genes were subsequently extracted and processed using BUSCO_phylogenomics.py (63) to generate a concatenated supermatrix and its corresponding partition file, retaining loci with a completeness threshold of 90%. Phylogenetic inference was performed in IQ-TREE 3 using the partitioned supermatrix approach. The best-fitting substitution models were selected using ModelFinder Plus with partition merging enabled (MFP+MERGE). Node support was assessed using 1,000 ultrafast bootstrap replicates.

### Copy-number analysis

We processed and visualized the gene copy number matrices per HGT event in R. We imported and rooted species phylogenies using ape, and ordered species according to their phylogenetic relationships using ggtree (64). The matrices were displayed as rows, with HGT events represented as columns. We generated heatmaps using pheatmap (https://github.com/raivokolde/pheatmap), with hierarchical clustering of HGT events, while keeping species in phylogenetic order without clustering. For visualization purposes, copy numbers greater than 10 were capped at 10. Gene copy numbers were represented using a continuous color gradient ranging from white to dark red, with increasing color intensity corresponding to higher copy numbers.

### Transcriptomic analysis

Transcriptomic analyses were performed using one TSA (Transcriptome Shotgun Assembly) master assemblies per species, *Bradysia coprophila* and *Pseudolycoriella hygida*. Protein sequences were searched against the corresponding TSA datasets using BLASTn from the BLAST+ suite. Hits with a minimum sequence identity of 95% and an alignment coverage of at least 30% were retained for downstream analyses.

For TPM analysis, we obtained public paired-end RNA-seq data for *Pseudolycoriella hygida* from BioProject PRJNA575761, including 32 RNA-seq libraries representing embryos, whole larvae, pupae, adults, larval brain, fat body, gut, and salivary glands. One salivary-gland library (SG_D12_R1) was excluded because of evidence for duplication/mislabeling. Read quality was assessed with FastQC v0.12.1 (https://www.bioinformatics.babraham.ac.uk/projects/fastqc/) and MultiQC v1.35 (65), and adapters were removed with fastp v1.3.6 (66). Transcript abundance was quantified with Salmon v1.12.0 (67) against a harmonized annotated CDS reference containing 17,881 CDS. Salmon was run with library type ISR, --validateMappings, --minScoreFraction 0.65, --seqBias and --gcBias. Salmon collapsed exact duplicate CDS sequences during indexing, resulting in 17,690 distinguishable sequence targets. Among 443 HGT candidate loci, eight belonged to four pairs of sequence-identical HGT CDS, yielding 439 RNA-seq-distinguishable expression units. We used TPM values for descriptive comparisons among tissues and developmental contexts. For contexts represented by multiple libraries, expression was summarized for each HGT unit as the median TPM across libraries. We complemented GO-based functional categories with manually curated InterPro annotations for HGT genes lacking informative GO assignments. Informative MF classification increased from 293/443 HGT loci (66.1%) using informative GO alone to 377/443 (85.1%) after curated InterPro integration.

### Best hits analysis between springtails and sciarids within HGT events

Protein sequence similarities within each event involving sciarids and springtails were assessed using BLASTp with an E-value threshold of (10e-5). Sciarid proteins were used as queries and those from springtails as the reference database. Results were exported in tabular format, including query and subject sequence identifiers, percentage identity, alignment length, number of mismatches and gap openings, alignment coordinates, E-value, bit score, and query coverage (*qcovs*). For each event, the retained best hit was defined as the alignment with the highest percentage identity applying a minimum query coverage threshold at 30%.

### Gene ontology enrichment analysis

Gene ontology annotation of the complete *B. coprophila* and *P. hygida* proteomes was performed with InterProScan version 5.73-104.0 (68). For each species, we used the complete predicted proteome as the reference population and the HGT-derived protein set as the foreground. We performed Biological Process (BP) and Molecular Function (MF) enrichment analyses separately with topGO v2.62.0 (https://bioconductor.org/packages/release/bioc/html/topGO.html). We obtained GO terms and hierarchical relationships from GO.db v3.22.0, corresponding to the Gene Ontology release dated July 22, 2025. We propagated direct protein annotations to ancestral terms through the GO directed acyclic graph. We evaluated enrichment using one-sided Fisher tests with the elim, weight01, and classic algorithms. A term was retained when it was associated with at least three HGT candidates, showed at least twofold enrichment relative to the ontology-specific annotated proteome, and had both elim and weight01 P<0.05. We reported Benjamini– Hochberg-adjusted values from the classic test as complementary statistics but did not include them in the primary selection criterion. For interpretation, we grouped enriched terms into predefined functional categories using “is_a” and “part_of” relationships in the same GO release.

### dN/dS Analysis

To characterize the selective regime acting on horizontally transferred genes, we estimated branch-specific dN/dS ratios (= ω) for every focal clade containing both Sciaridae and Collembola members. For each such focal clade, the coding sequences (CDS) corresponding to sciarid and springtail proteins were aligned using the OMM_MACSE pipeline v10.02 (69). Briefly, this pipeline comprises multiple steps, including: (i) a prefiltering step to remove non-homologous sequence fragments, (ii) amino-acid alignment with MUSCLE (70), (iii) cleaning of the resulting alignment with HMMcleaner (71), and (iv) reverse translation into a codon alignment. For each alignment, we inferred a maximum-likelihood gene tree from the amino-acid alignment with IQ-TREE v3.0.1, using the optimal model selected by ModelFinder and 1,000 ultrafast bootstrap replicates. Branch-specific ω values were estimated on each codon alignment and its tree under the MG94×REV codon-substitution model (= free-ratio model, (72)) as implemented in HyPhy FitMG94 (73).

Maximum-likelihood estimates of branch-specific ω were filtered to exclude poorly estimated values, specifically branches with too few synonymous substitutions (dS < 0.01) or substitutions saturation (dS and/or dN > 1). Internal branches were assigned to Sciaridae or the Collembola lineages only if all descendant tips belonged to only one of these lineages. For each gene, we then averaged ω separately over the retained Sciaridae branches and the retained Collembola branches, yielding one mean ω value per gene per lineage. Finally, for genes with at least four retained branches in each lineage, we compared the distributions of ω values between Sciaridae and Springtails using a Wilcoxon rank-sum test, and corrected P values with the Benjamini-Hochberg procedure (FDR < 0.05).

### dS analysis

To test whether candidate transferred genes diverged more recently than vertically inherited genes, we compared their synonymous divergence (dS) with that of conserved single-copy genes. For each candidate transferred gene, located either on somatic chromosomes or on GRCs, we computed dS between all pairs of sequences in the codon alignment obtained above (see dN/dS analysis), as implemented in the kaks function of the seqinr R package (74). For each gene, we retained the minimum dS between a springtail and a sciarid sequence. As a control for the expected dS distribution of vertically inherited genes, we used BUSCO genes [hexapoda_odb12] found as complete and single-copy in both *B. coprophila* and *F. candida* (n = 2,383). For each shared BUSCO gene, the two CDS were aligned with MACSE as described above, and their dS was computed in the same way. When dS cannot be estimated because of substitution saturation, seqinr sets it to a ceiling value of 9.99. These values were kept and interpreted as saturated. Finally, we compared the distributions of minimum dS of candidate transferred genes on somatic chromosomes and on GRCs with the distribution of BUSCO dS.

## Author contributions

CG and FM designed the research. AJdD, MP, and FM performed the analyses. All authors interpreted the data. AJdD, CG, and FM drafted the manuscript. All authors contributed to the final manuscript.

## Acknowledgements

AJdD was funded by an INRAe PhD fellowship. AJdD and FM are supported by Saclay Plant Sciences-SPS (ANR-17-EUR-0007) and the PlantBioinfoPF platform. FM and CG are supported by the French national research agency (ANR), project HORIGIN ANR-26-CE02-2878. CG is also supported by ANR project VIRHOZFER (ANR-24-CE02-1004). The funders had no role in study design, data collection and analysis, decision to publish, or preparation of the manuscript.

## Supplementary information text

Multiple lines of evidence suggest that the 3.14 Mb region at the beginning of the *P. hygida* chromosome CM055403.1 represents contaminating bacterial DNA erroneously incorporated into the genome assembly. This region showed no evidence of gene transcription (Table S2), exhibited an atypical G+C content (Figure S2), and contained no genes with an inferred metazoan last common ancestor (data not shown). Furthermore, neither long-nor short-read mapping supported its physical continuity with the remainder of the chromosome (data not shown), consistent with published Hi-C data (30).

**Figure S1:**
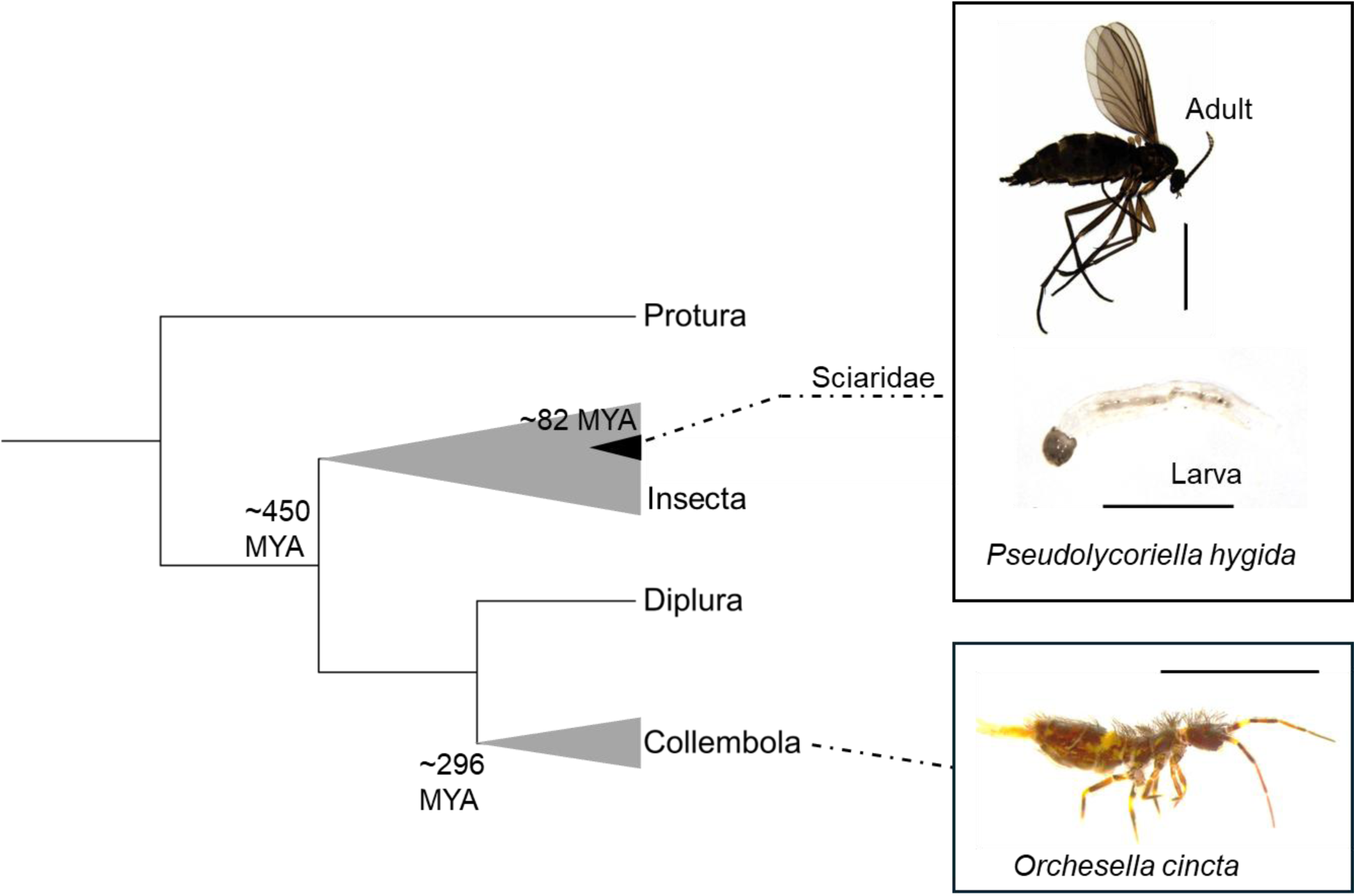
Phylogeny and morphology of Sciaridae and collembola (springtails). Left: cladogram of the four clades of hexapods, adapted from (22). Approximate datations were taken from (16, 36, 45). Right: images of representative species from the insect family Sciaridae (*P. hygida*, top) and from collembola (*O. cincta*, bottom). *P. hygida* (Sauaia & Alves): adapted from Menzel et al. (2024), Insects 15, 118, doi:10.3390/insects15020118, under CC BY 4.0; Adult scale bar: 2 mm, larva scale bar: 500 μm. *O. cincta*: photograph by Donald Hobern (2018), scale bar: 2 mm, adapted from Wikimedia Commons under the Creative Commons Attribution 2.0 license (CC BY 2.0). https://commons.wikimedia.org/wiki/File:Orchesella_cincta_(39474705284).jpg

**Figure S2.**
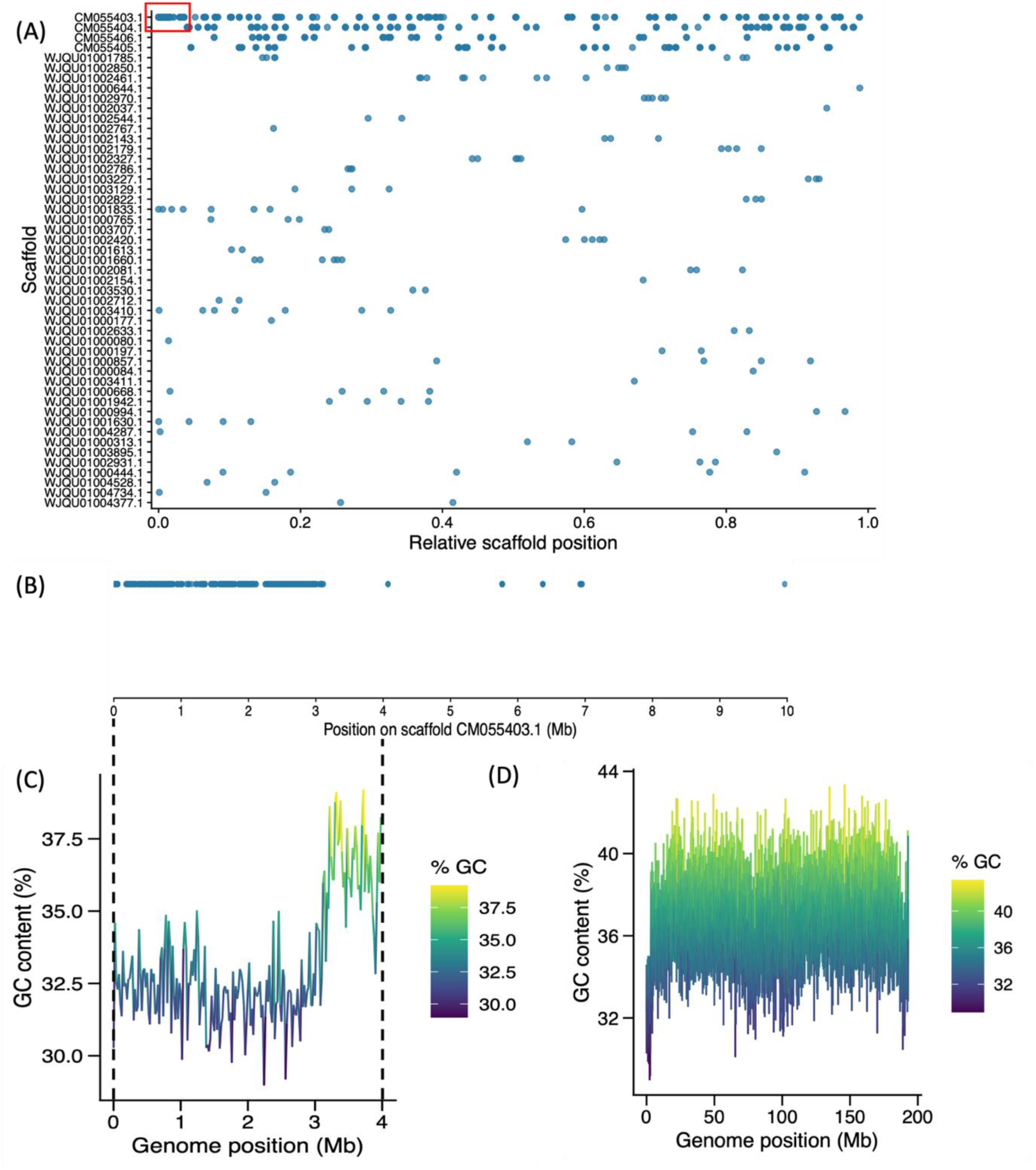
Genomic distribution of putative transferred genes and G+C content of scaffold CM055403.1 support the identification of a candidate bacterial contamination in *Pseudolycoriella hygida*. (A) Distribution of HGT candidates across the *P. hygida* genomic scaffolds. Each point represents a candidate HGT-derived protein selected after manual examination of phylogenetic trees. The x-axis indicates the relative position of each protein along its scaffold, and scaffolds are ordered by decreasing length. The red box indicates the hotspot of candidates. (B) Enlarged view of the candidate protein distribution in the first 10 Mb of the scaffold CM055403.1. (C) G+C content distribution along the first 4 Mb of scaffold CM055403.1 of *P. hygida*. The colour gradient represents the G+C content of each window. Dashed lines are cast at the same positions as in (B). (D) G+C content along the whole scaffold CM055403.1.

**Figure S3.**
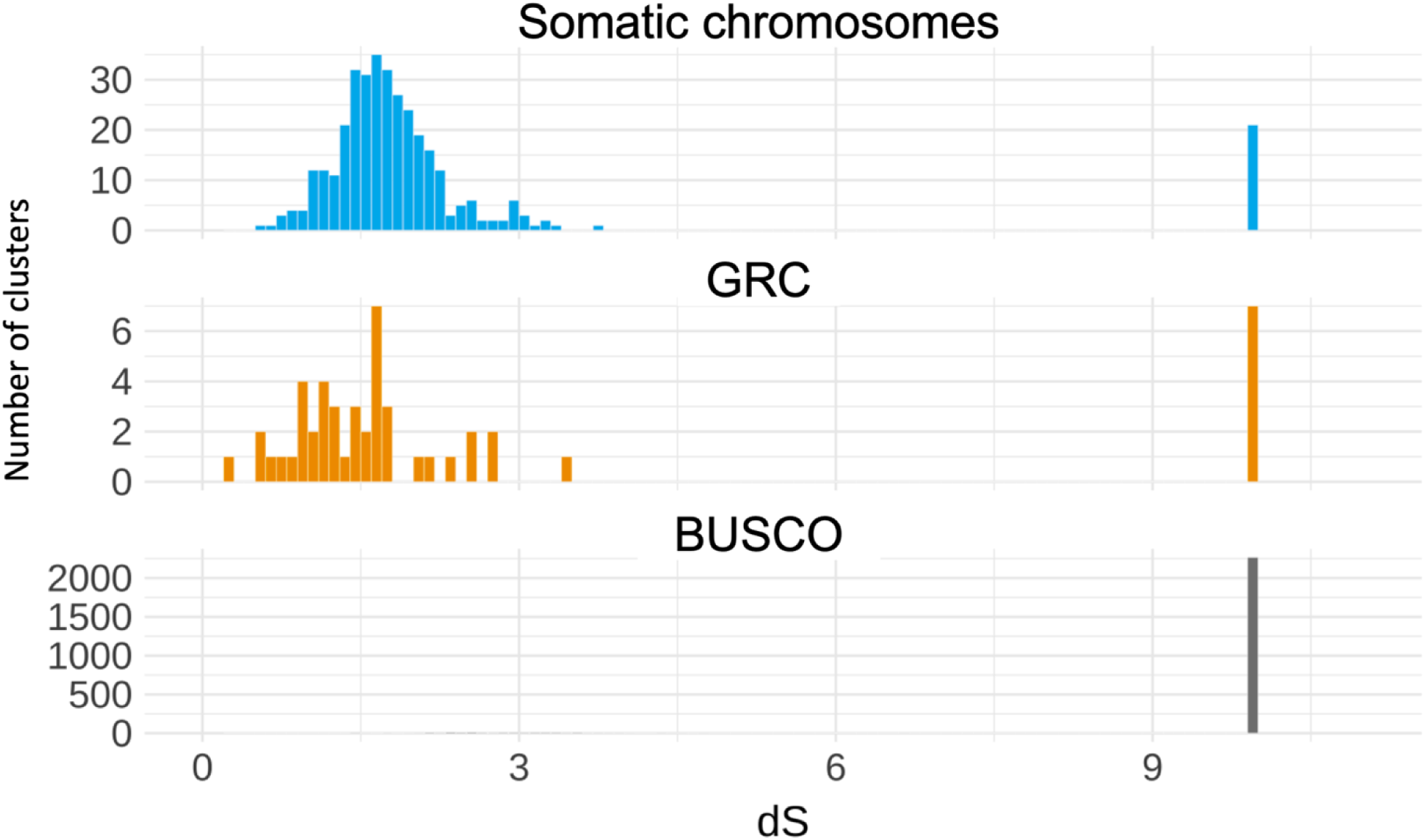
Distribution of minimum pairwise dS between springtail and sciarid sequences. Histograms showing the number of focal clades per pairwise dS estimate for HGT genes found on sciarid somatic chromosomes (top) or on *B. coprophila* GRCs only (middle), and the number of BUSCO genes per pairwise dS estimate (bottom).

**Figure S4. Phylogenetic trees with horizontal gene transfer (HGT) events in somatic chromosomes.** The Figure compiles 331 phylogenetic trees, from which inferred HGT events are listed in Table S3. For each tree, “Tree name” embeds the name of the representative protein of the corresponding final cluster. For instance, “bra_cop_dipt_XP_037024267.1_treefile” refers to the representative protein “XP_037024267.1”. “Protein length” indicates the size (in aa) of the cluster representative protein. “Alignment length” indicates the size (in aa) of the multiple sequence alignment before trimming less informative columns. “Aligned AA count” refers to the size (in aa) of the final alignment used to build the phylogenetic tree, after trimming less informative columns. Red dots indicate nodes with bootstrap support values >90%, and leaf colors represent the different taxa. Numbers at the nodes identify inferred HGT events and correspond to the event names listed in Table S3. Black nodes represent upper nodes, which identify individual HGT events or the overall event associated with multiple parallel occurrences. Blue nodes represent deep nodes, which identify the focal clades associated with individual parallel occurrences of an HGT event. An upper node without an associated deep node indicates an individual HGT event, whereas an upper node with an associated deep node indicates multiple parallel occurrences. Note: Due to the large number of trees, the complete high-resolution figure is provided as a separate, standalone PDF file.

**Figure S5.**
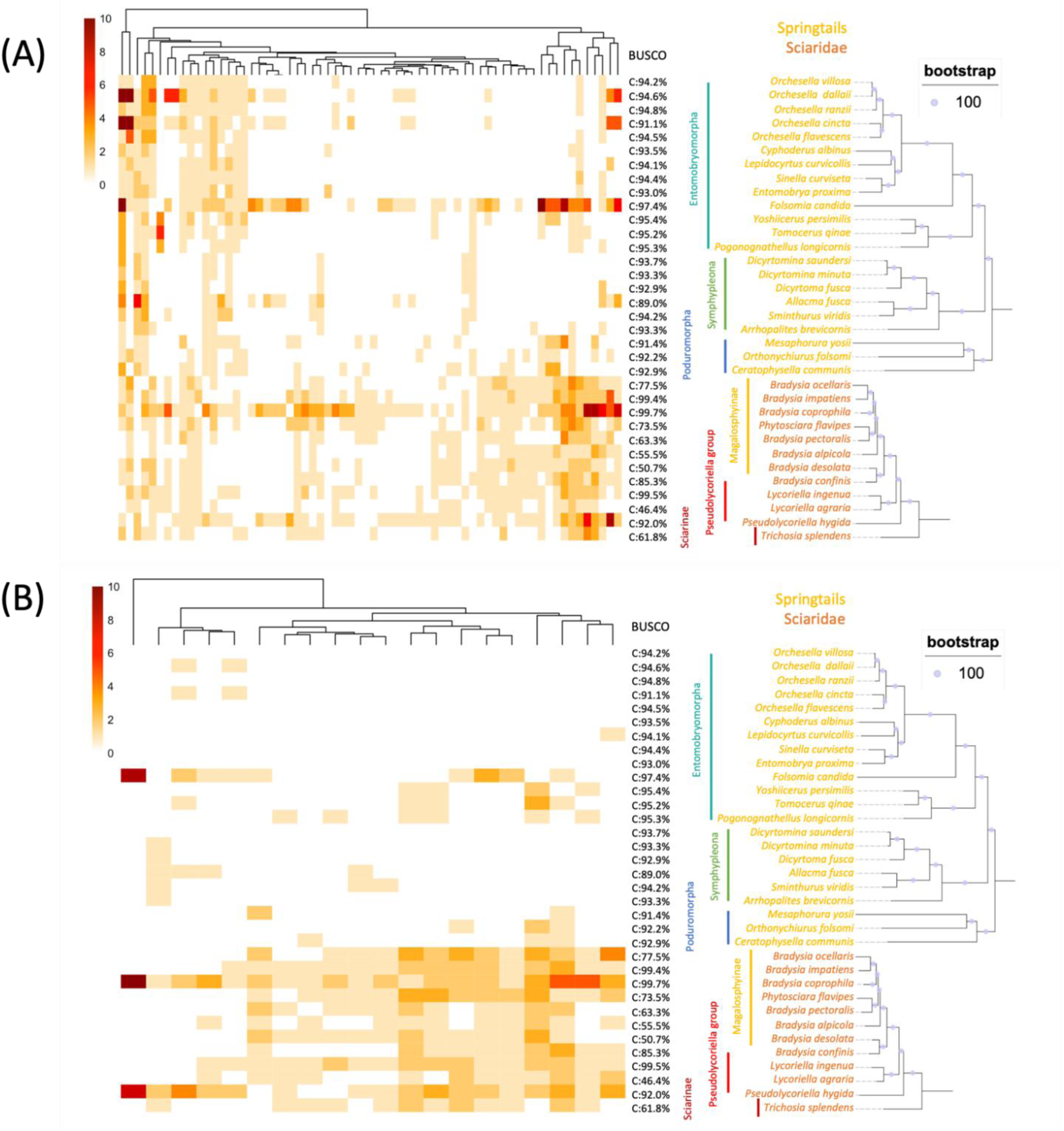
Gene copy number in springtail and sciarid genomes per HGT event (left), BUSCO genome scores (middle), and springtails and sciarid species tree (right). (A) Sciarids-to-springtails HGT. (B) Sciarid-springtail sistership.

**Figure S6.**
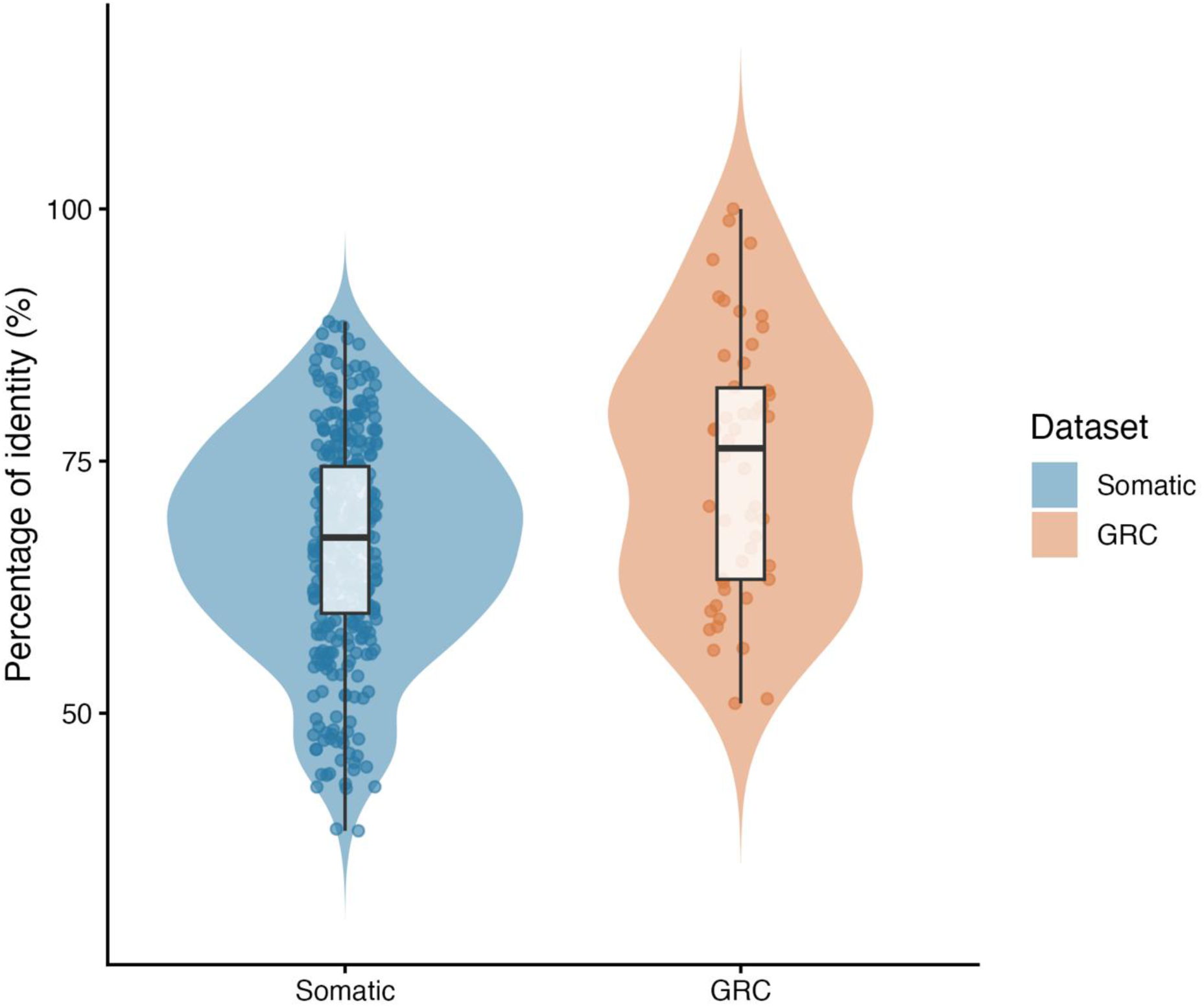
Distribution of the highest sequence identity value (%) among sciarid versus springtail proteins within each HGT event. All values are available in Table S4. The violin represents the distribution density of percentage identity values across protein pairs. Each point represents the protein pair with the highest percentage identity for a given event, and the white box indicates the median and interquartile range.

**Figure S7. Phylogenetic trees with horizontal gene transfer (HGT) events in GRCs only.** The Figure compiles 50 phylogenetic trees, from which inferred HGT events are listed in Table S5. For each tree, “Tree name” embeds the name of the representative protein of the corresponding final cluster. For instance, “all_fus_coll_CAG7630510.1_treefile” refers to the representative protein “CAG7630510.1”. “Protein length” indicates the size (in aa) of the cluster representative protein. “Alignment length” indicates the size (in aa) of the multiple sequence alignment before trimming less informative columns. “Aligned AA count” refers to the size (in aa) of the final alignment used to build the phylogenetic tree, after trimming less informative columns. Red dots indicate nodes with bootstrap support values >90%, and leaf colors represent the different taxa. Numbers at the nodes identify inferred HGT events and correspond to the event names listed in Table S5. Black nodes represent upper nodes, which identify individual HGT events or the overall event associated with multiple parallel occurrences. Blue nodes represent deep nodes, which identify the focal clades associated with individual parallel occurrences of an HGT event. An upper node without an associated deep node indicates an individual HGT event, whereas an upper node with an associated deep node indicates multiple parallel occurrences. Note: Due to the large number of trees, the complete high-resolution figure is provided as a separate, standalone PDF file.

**Figure S8.**
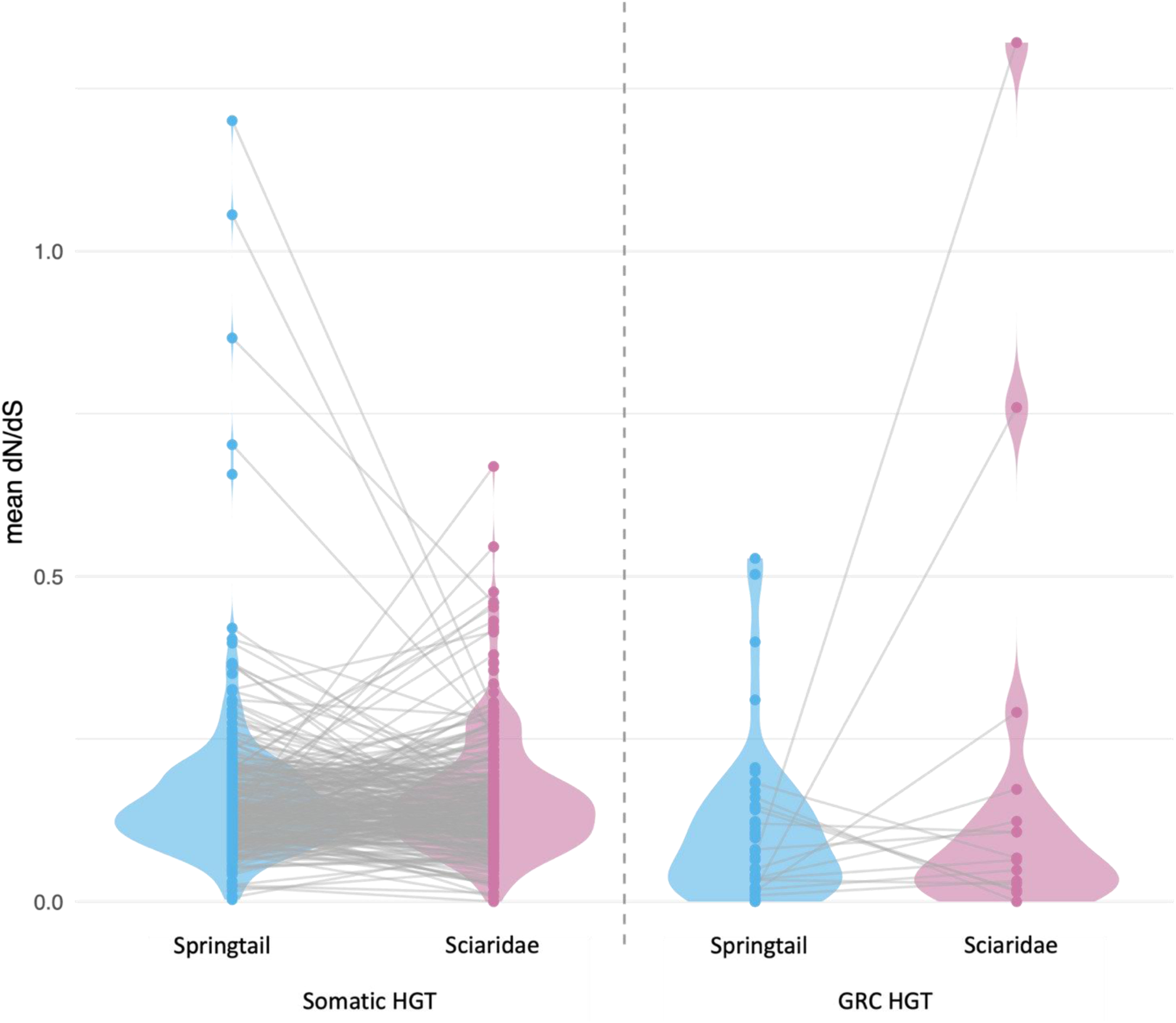
Mean dN/dS of HGT candidates in Springtails and Sciaridae for the proteins found on sciarid somatic chromosomes (left) and *B. coprophila* GRCs only (right). Each point represents the mean ω across all branches of one species clade for one HGT event. Grey lines connect the two values of the same gene in each taxon, so each line compares the mean selective regime of the springtail and sciarid copies. Violins show the density distribution of all gene means.

**Figure S9.**
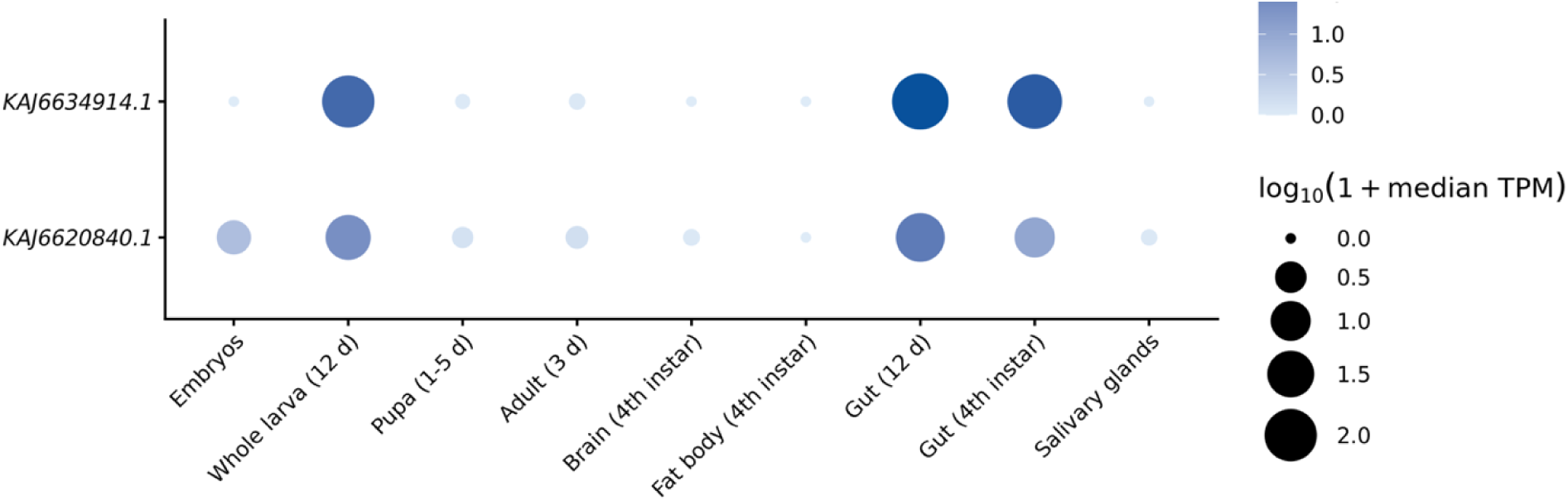
Expression profiles of two horizontally transferred genes across developmental stages and tissues of *Pseudolycoriella hygida*. Bubble heatmap showing the expression levels of two HGT-derived genes (KAJ6634914.1 and KAJ6620840.1) across nine developmental and tissue contexts. Expression was quantified as transcripts per million (TPM) using Salmon. Bubble size and colour intensity both represent log₁₀(1 + median TPM), with larger and darker bubbles indicating higher expression. Median TPM values were calculated across six embryonic libraries and 18 salivary gland libraries, whereas the remaining contexts were each represented by a single library. Developmental and tissue contexts include embryos, whole larvae (12 days), pupae (1–5 days), adults (3 days), fourth-instar brains and fat bodies, and guts from 12-day-old larvae and fourth-instar larvae. One suspected duplicate or mislabeled salivary gland library (SG_D12_R1) was excluded from the analysis.

**Figure S10.**
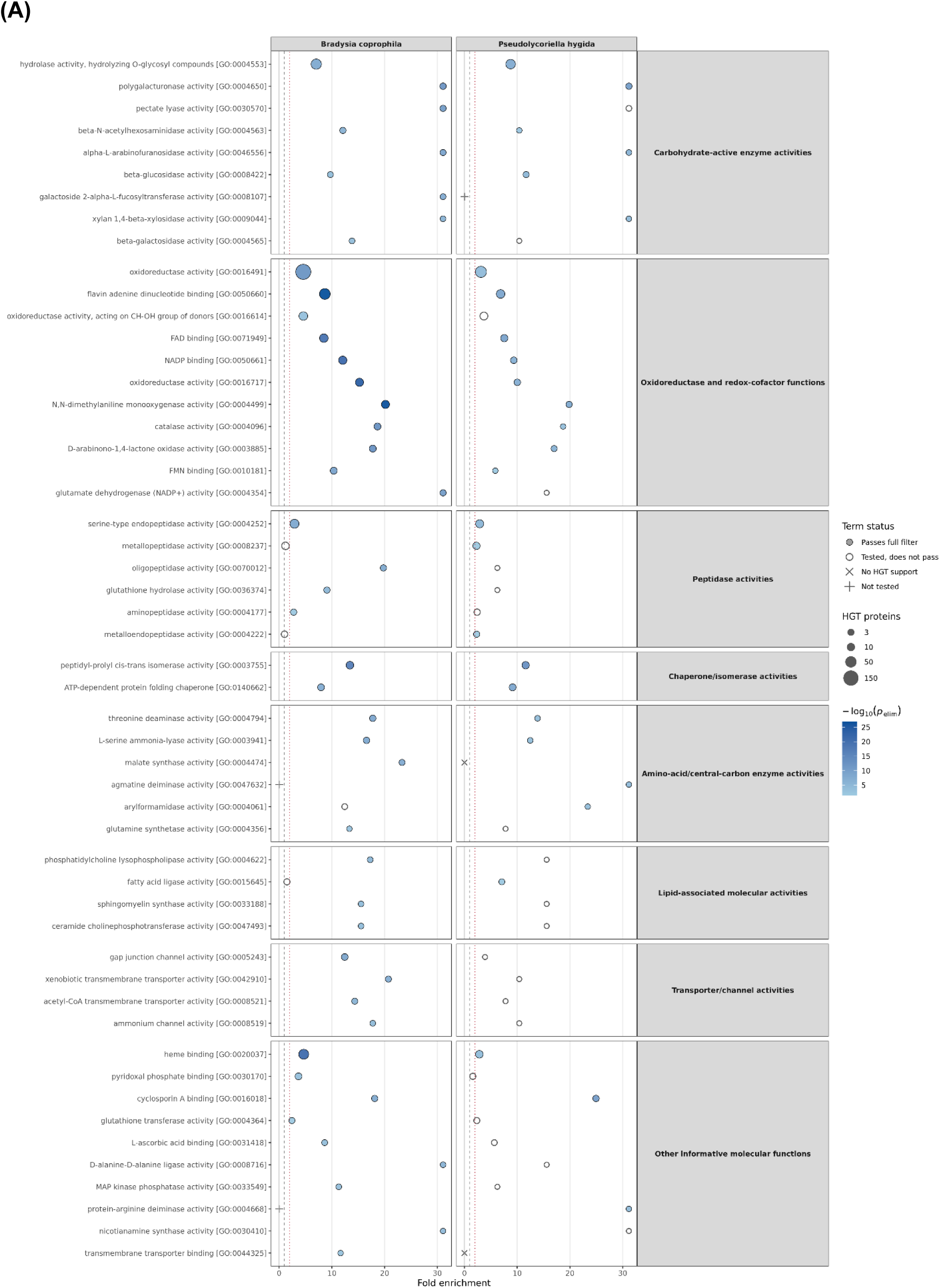
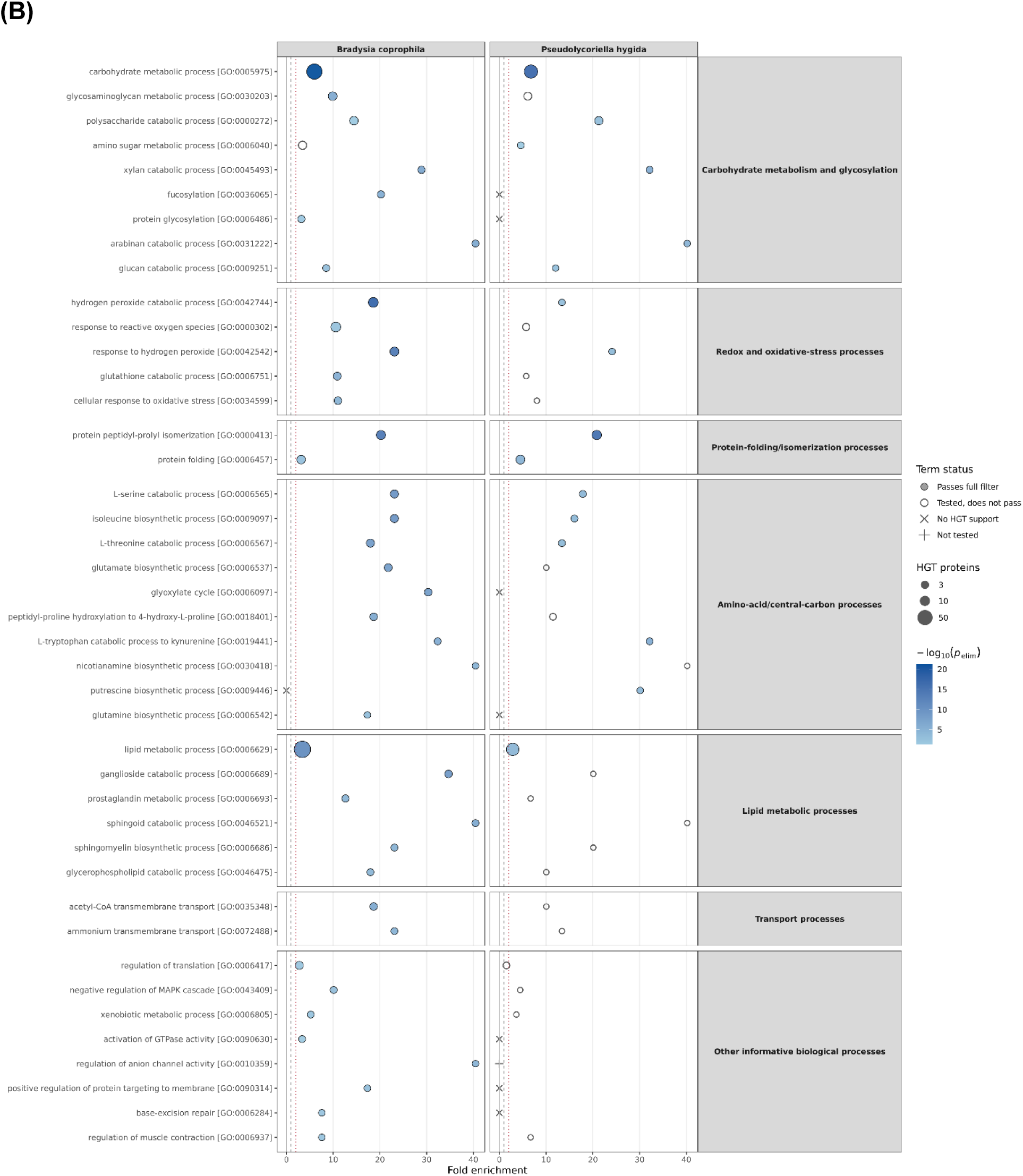
Gene Ontology enrichment among sciarid HGT candidates. Enriched GO terms are shown for (A) Molecular Function and (B) Biological Process in *Bradysia coprophila* and *Pseudolycoriella hygida*. Point size indicates the number of HGT candidates associated with a term, and point fill represents −log⁡10 of the topGO elim Fisher P value. Filled circles indicate terms passing the complete filter in that species: at least three annotated HGT candidates, at least twofold enrichment, elim Fisher P<0.05, and weight01 Fisher P<0.05. Open circles indicate tested terms that did not meet all four criteria; crosses indicate terms with no HGT support, and plus signs indicate terms not tested in that species. The vertical dashed line marks twofold enrichment. Categories containing no more than 10 qualifying terms are shown in full; for larger categories, the union of the five terms with the largest HGT-candidate counts and the five with the greatest fold enrichment is displayed.

**Figure S11.**
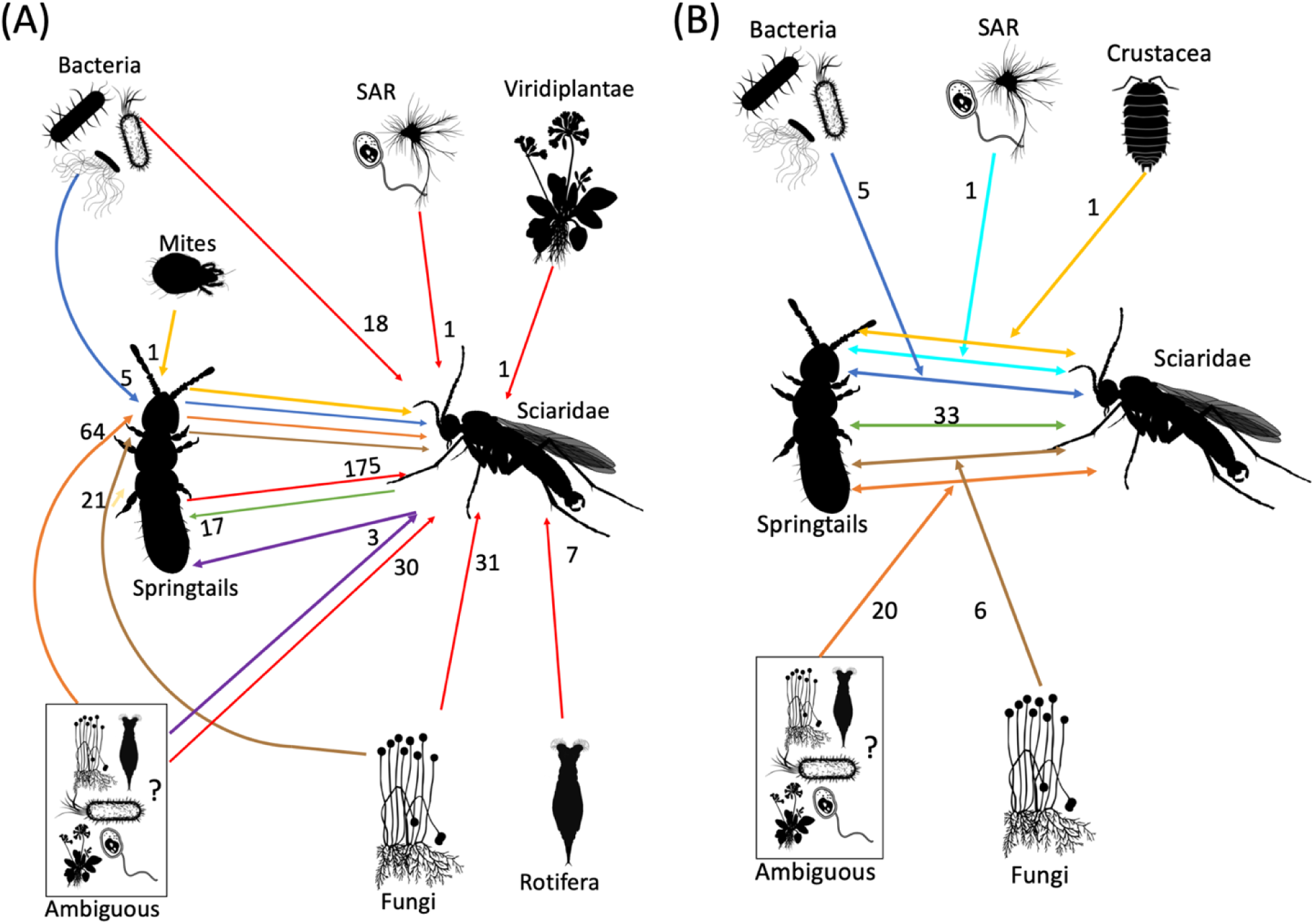
Summary of the horizontal gene transfer (HGT) events in somatic chromosomes. (A) All inferred HGT events, excluding those involving sister relationships between springtails and sciarids. (B) HGT events associated with sister relationships between springtails and sciarids.

**Figure S12.**
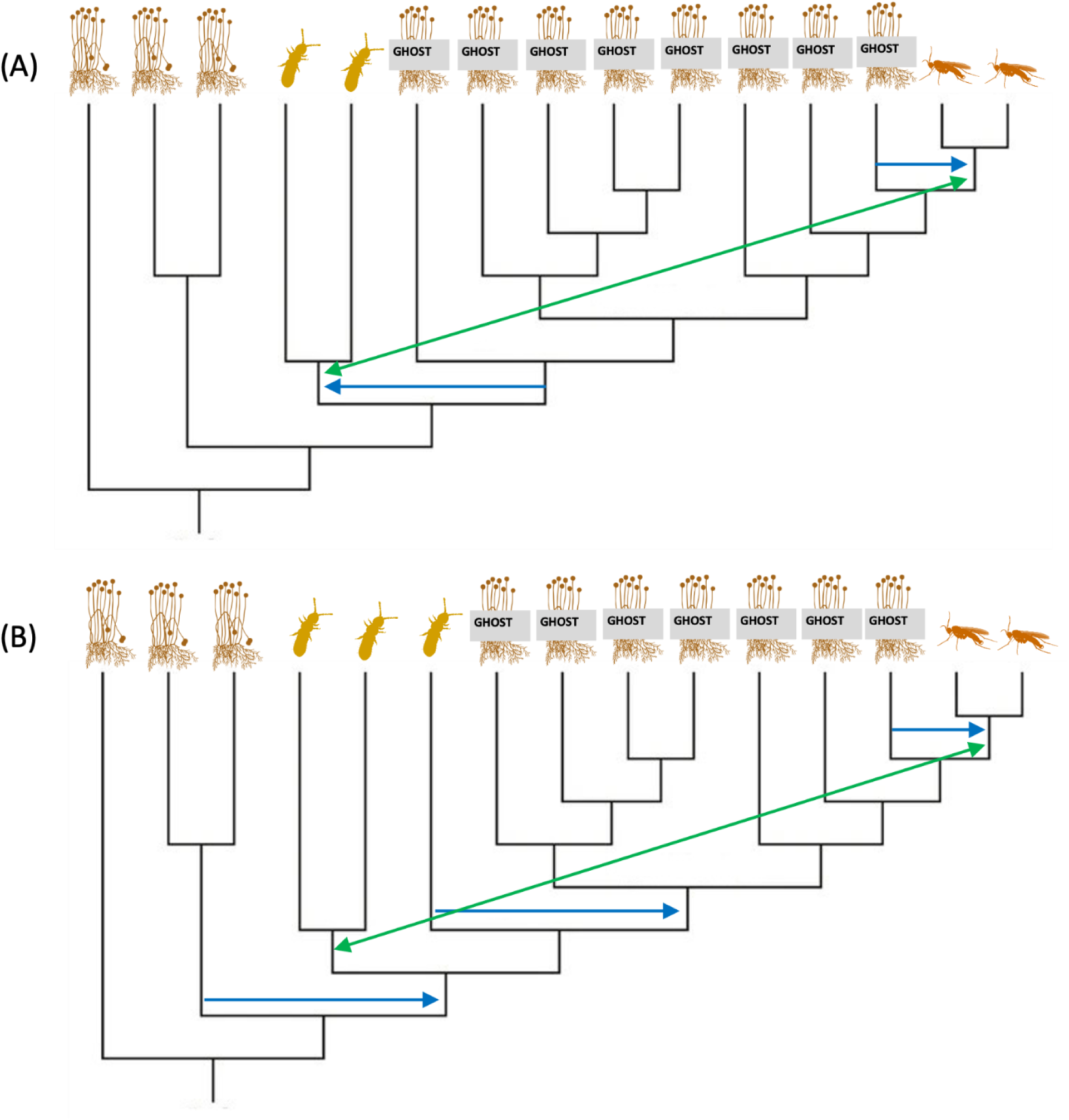
Alternative scenarios involving ghost lineages to explain the phylogenetic relationships observed between springtail and sciarid proteins. Green arrows follow a scenario involving direct HGT between sciarids and springtails, and blue arrows depict successive transfers involving a putative ghost lineage. (A) Situation in which the direction of HGT between springtails and sciarids remains unresolved (sistership). (B) Supported springtails-to-sciarids or sciarids-to-springtails HGT.

**Table S1.**
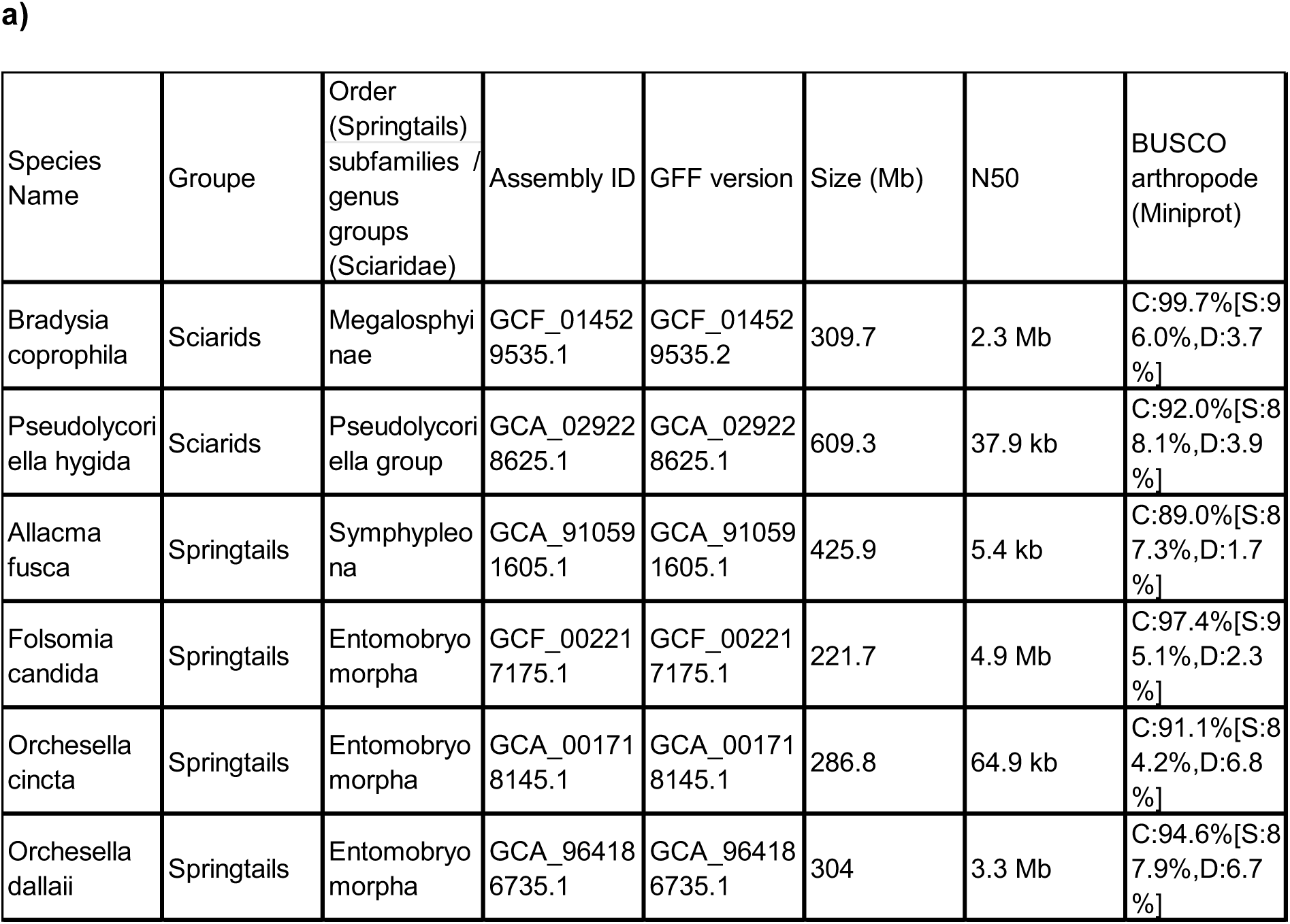
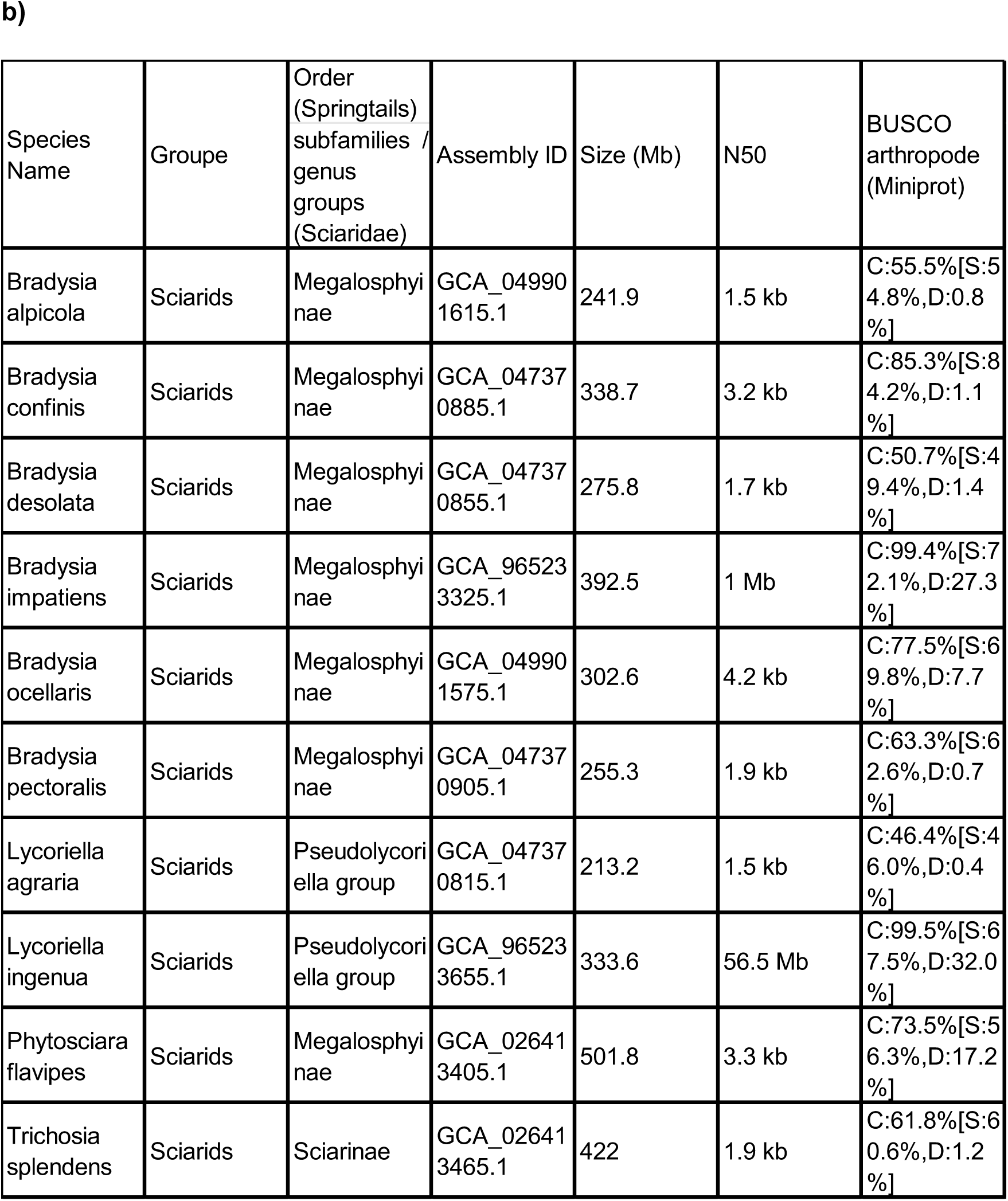
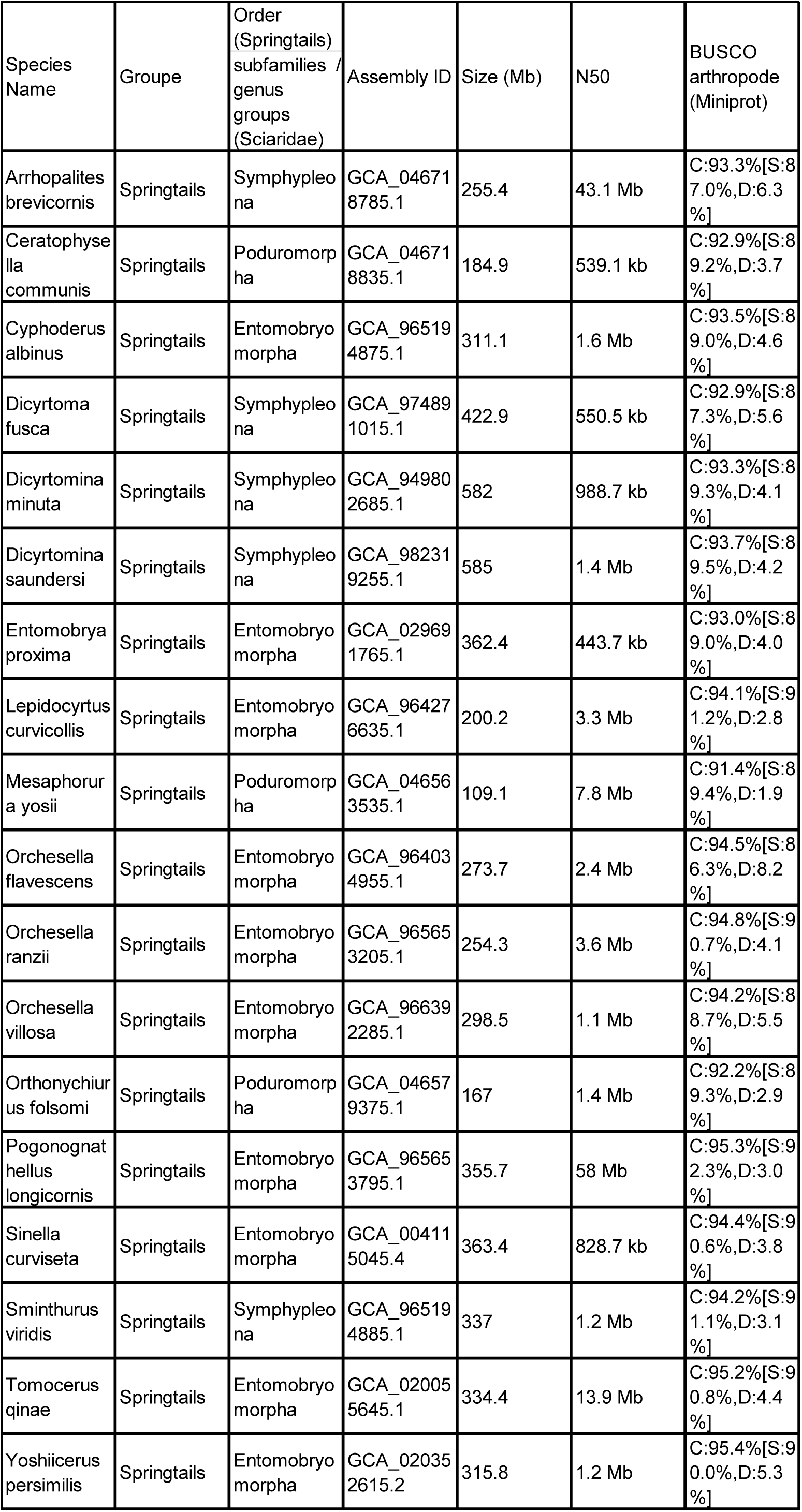
Genomic assembly statistics of the selected springtail and sciarid species. The table shows the species, taxonomic group, subfamily or genus group, assembly accession, GFF annotation version, assembly size, N50, and BUSCO completeness assessed using Miniprot with the arthropod lineage dataset. BUSCO scores are reported as complete (C), single-copy (S), and duplicated (D) orthologs. a) NCBI genomes and proteoms used for Taxtrack b) NCBI genomes used for Miniprot

| Species Name | Groupe | Order (Springtails) subfamilies / genus groups (Sciaridae) | Assembly ID | GFF version | Size (Mb) | N50 | BUSCO arthropode (Miniprot) |
|---|---|---|---|---|---|---|---|
| Bradysia coprophila | Sciarids | Megalosphyinae | GCF_014529535.1 | GCF_014529535.2 | 309.7 | 2.3 Mb | C:99.7%[S:96.0%,D:3.7%] |
| Pseudolycoriella hygida | Sciarids | Pseudolycoriella group | GCA_029228625.1 | GCA_029228625.1 | 609.3 | 37.9 kb | C:92.0%[S:88.1%,D:3.9%] |
| Allacma fusca | Springtails | Symphypleona | GCA_910591605.1 | GCA_910591605.1 | 425.9 | 5.4 kb | C:89.0%[S:87.3%,D:1.7%] |
| Folsomia candida | Springtails | Entomobryomorpha | GCF_002217175.1 | GCF_002217175.1 | 221.7 | 4.9 Mb | C:97.4%[S:95.1%,D:2.3%] |
| Orchesella cincta | Springtails | Entomobryomorpha | GCA_001718145.1 | GCA_001718145.1 | 286.8 | 64.9 kb | C:91.1%[S:84.2%,D:6.8%] |
| Orchesella dallaii | Springtails | Entomobryomorpha | GCA_964186735.1 | GCA_964186735.1 | 304 | 3.3 Mb | C:94.6%[S:87.9%,D:6.7%] |
| Species Name | Groupe | Order (Springtails) subfamilies / genus groups (Sciaridae) | Assembly ID | Size (Mb) | N50 | BUSCO arthropode (Miniprot) |  |
| Bradysia alpicola | Sciarids | Megalosphyinae | GCA_049901615.1 | 241.9 | 1.5 kb | C:55.5%[S:54.8%,D:0.8%] |  |
| Bradysia confinis | Sciarids | Megalosphyinae | GCA_047370885.1 | 338.7 | 3.2 kb | C:85.3%[S:84.2%,D:1.1%] |  |
| Bradysia desolata | Sciarids | Megalosphyinae | GCA_047370855.1 | 275.8 | 1.7 kb | C:50.7%[S:49.4%,D:1.4%] |  |
| Bradysia impatiens | Sciarids | Megalosphyinae | GCA_965233325.1 | 392.5 | 1 Mb | C:99.4%[S:72.1%,D:27.3%] |  |
| Bradysia ocellaris | Sciarids | Megalosphyinae | GCA_049901575.1 | 302.6 | 4.2 kb | C:77.5%[S:69.8%,D:7.7%] |  |
| Bradysia pectoralis | Sciarids | Megalosphyinae | GCA_047370905.1 | 255.3 | 1.9 kb | C:63.3%[S:62.6%,D:0.7%] |  |
| Lycoriella agraria | Sciarids | Pseudolycoriella group | GCA_047370815.1 | 213.2 | 1.5 kb | C:46.4%[S:46.0%,D:0.4%] |  |
| Lycoriella ingenua | Sciarids | Pseudolycoriella group | GCA_965233655.1 | 333.6 | 56.5 Mb | C:99.5%[S:67.5%,D:32.0%] |  |
| Phytosciara flavipes | Sciarids | Megalosphyinae | GCA_026413405.1 | 501.8 | 3.3 kb | C:73.5%[S:56.3%,D:17.2%] |  |
| Trichosia splendens | Sciarids | Sciarinae | GCA_026413465.1 | 422 | 1.9 kb | C:61.8%[S:60.6%,D:1.2%] |  |
| Species Name | Groupe | Order (Springtails) subfamilies / genus groups (Sciaridae) | Assembly ID | Size (Mb) | N50 | BUSCO arthropode (Miniprot) |  |
| Arrhopalites brevicornis | Springtails | Symphypleona | GCA_046718785.1 | 255.4 | 43.1 Mb | C:93.3%[S:87.0%,D:6.3%] |  |
| Ceratophysella communis | Springtails | Poduromorpha | GCA_046718835.1 | 184.9 | 539.1 kb | C:92.9%[S:89.2%,D:3.7%] |  |
| Cyphoderus albinus | Springtails | Entomobryomorpha | GCA_965194875.1 | 311.1 | 1.6 Mb | C:93.5%[S:89.0%,D:4.6%] |  |
| Dicyrtoma fusca | Springtails | Symphypleona | GCA_974891015.1 | 422.9 | 550.5 kb | C:92.9%[S:87.3%,D:5.6%] |  |
| Dicyrtomina minuta | Springtails | Symphypleona | GCA_949802685.1 | 582 | 988.7 kb | C:93.3%[S:89.3%,D:4.1%] |  |
| Dicyrtomina saundersi | Springtails | Symphypleona | GCA_982319255.1 | 585 | 1.4 Mb | C:93.7%[S:89.5%,D:4.2%] |  |
| Entomobrya proxima | Springtails | Entomobryomorpha | GCA_029691765.1 | 362.4 | 443.7 kb | C:93.0%[S:89.0%,D:4.0%] |  |
| Lepidocyrtus curvicollis | Springtails | Entomobryomorpha | GCA_964276635.1 | 200.2 | 3.3 Mb | C:94.1%[S:91.2%,D:2.8%] |  |
| Mesaphorura yosii | Springtails | Poduromorpha | GCA_046563535.1 | 109.1 | 7.8 Mb | C:91.4%[S:89.4%,D:1.9%] |  |
| Orchesella flavescens | Springtails | Entomobryomorpha | GCA_964034955.1 | 273.7 | 2.4 Mb | C:94.5%[S:86.3%,D:8.2%] |  |
| Orchesella ranzii | Springtails | Entomobryomorpha | GCA_965653205.1 | 254.3 | 3.6 Mb | C:94.8%[S:90.7%,D:4.1%] |  |
| Orchesella villosa | Springtails | Entomobryomorpha | GCA_966392285.1 | 298.5 | 1.1 Mb | C:94.2%[S:88.7%,D:5.5%] |  |
| Orthonychiurus folsomi | Springtails | Poduromorpha | GCA_046579375.1 | 167 | 1.4 Mb | C:92.2%[S:89.3%,D:2.9%] |  |
| Pogonognathellus longicornis | Springtails | Entomobryomorpha | GCA_965653795.1 | 355.7 | 58 Mb | C:95.3%[S:92.3%,D:3.0%] |  |
| Sinella curviseta | Springtails | Entomobryomorpha | GCA_004115045.4 | 363.4 | 828.7 kb | C:94.4%[S:90.6%,D:3.8%] |  |
| Sminthurus viridis | Springtails | Symphypleona | GCA_965194885.1 | 337 | 1.2 Mb | C:94.2%[S:91.1%,D:3.1%] |  |
| Tomocerus qinae | Springtails | Entomobryomorpha | GCA_020055645.1 | 334.4 | 13.9 Mb | C:95.2%[S:90.8%,D:4.4%] |  |
| Yoshiicerus persimilis | Springtails | Entomobryomorpha | GCA_020352615.2 | 315.8 | 1.2 Mb | C:95.4%[S:90.0%,D:5.3%] |  |

**Table S2. Best BLAST hit results for the candidate contaminant sequences identified on scaffold CM055403.1 of *Pseudolycoriella hygida*.** The dataset reports the best TBLASTN hit for each protein query, including the protein and transcript identifiers, percentage identity, alignment length, number of mismatches and gap openings, query and subject alignment coordinates, E-value, and alignment score. Note: Due to its extensive length and to allow for data sorting, this table is provided as a separate standalone Excel file.

**Table S3. Summary information of the HGT events in somatic chromosomes inferred from manual examination of the phylogenetic trees provided in Figure S4.** The table summarizes information associated with each inferred HGT event, including the phylogenetic tree file name, the upper node (black node in Figure S4), the deep node (blue node in Figure S4), the event name, the inferred donor, and the inferred direction of transfer between springtails. Upper nodes without an associated deep node represent individual HGT events, whereas upper nodes associated with one or more deep nodes represent multiple parallel HGT events involving distinct focal clades. The upper node identifies the overall event, while each deep node identifies a focal clade associated with an individual parallel occurrence. Note: Due to its extensive length and to allow for data sorting, this table is provided as a separate standalone Excel file.

**Table S4. Summary of best protein hits identified by sequence similarity searches**. The file indicates the query protein/tree file, “protein_A” and “protein_B” identify the two proteins compared, ID_percent represents the percentage of identical amino acid residues, bitscore is the alignment bit score, hit_length is the alignment length in amino acids, and qcovs indicates query coverage (%). Note: Due to its extensive length and to allow for data sorting, this table is provided as a separate standalone Excel file.

**Table S5. Summary information of the HGT events found in GRCs only inferred from manual examination of the phylogenetic trees provided in Figure S7.** The table summarizes the information associated with each inferred HGT event, including the phylogenetic tree file name, the event node (Figure S7), the event name, the inferred donor, and the inferred direction of transfer between springtails. Note: Due to its extensive length and to allow for data sorting, this table is provided as a separate standalone Excel file.

**Table S6. Summary of protein and transcript representation in inferred HGT events in *Bradysia coprophila* and *Pseudolycoriella hygida*.** For each event, the table reports the presence or absence of proteins from *B. coprophila* and *P. hygida*, the presence or absence of corresponding transcript hits (ID>95% and coverage>30%), and the number of transcripts identified for each species. TRUE and FALSE indicate presence and absence, respectively. Note: Due to its extensive length and to allow for data sorting, this table is provided as a separate standalone Excel file.

**Table S7. Best BLASTn transcript hits associated with inferred HGT events in *Bradysia coprophila* and *Pseudolycoriella hygida*.** For each event, the table reports the query sequence identifier (qseqid), subject sequence identifier (sseqid), percentage sequence identity, alignment length, number of mismatches and gap openings, query and subject alignment coordinates, E-value, bit score, and query coverage. The best transcript hit is reported for each species when available. Note: Due to its extensive length and to allow for data sorting, this table is provided as a separate standalone Excel file.

**Dataset S1. Initial FASTA sequence files used for phylogenetic analyses.** (A) Somatic chromosomes dataset. (B) GRC dataset. Sequences were selected based on clustering results and included the 10 best hits and the 20 most divergent hits for each cluster representative.

**Dataset S2. Final multiple sequence alignments (MSAs) used for phylogenetic analyses.** (A) Somatic chromosomes dataset. (B) GRC dataset. The alignments were generated from the initial FASTA sequences (Dataset S1) and subsequently trimmed prior to phylogenetic inference.

**Dataset S3. Phylogenetic trees inferred using IQ-TREE 3.** (A) Somatic chromosomes dataset. (B) GRC dataset. Trees were inferred from the final trimmed multiple sequence alignments provided in Dataset S2.

## Data availability

For each final cluster containing HGT proteins, the initial sequences (Dataset S1), their final multiple sequence alignment (Dataset S2), and the associated phylogenetic tree (Dataset S3) are available on Figshare (https://figshare.com/s/1e3f35fa3f16d7709fc0).

