## Supplementary material for "Extensive HGT between soil-living hexapods contributes to functional convergence": Figure S7

**Figure S7. Phylogenetic trees with horizontal gene transfer (HGT) events in GRCs only.** The Figure compiles 50 phylogenetic trees, from which inferred HGT events are listed in Table S5. For each tree, “Tree name” embeds the name of the representative protein of the corresponding final cluster. For instance, “all_fus_coll_CAG7630510.1_treefile” refers to the representative protein “CAG7630510.1”. “Protein length” indicates the size (in aa) of the cluster representative protein. “Alignment length” indicates the size (in aa) of the multiple sequence alignment before trimming less informative columns. “Aligned AA count” refers to the size (in aa) of the final alignment used to build the phylogenetic tree, after trimming less informative columns. Red dots indicate nodes with bootstrap support values >90%, and leaf colors represent the different taxa. Numbers at the nodes identify inferred HGT events and correspond to the event names listed in Table S5. Black nodes represent upper nodes, which identify individual HGT events or the overall event associated with multiple parallel occurrences. Blue nodes represent deep nodes, which identify the focal clades associated with individual parallel occurrences of an HGT event. An upper node without an associated deep node indicates an individual HGT event, whereas an upper node with an associated deep node indicates multiple parallel occurrences.

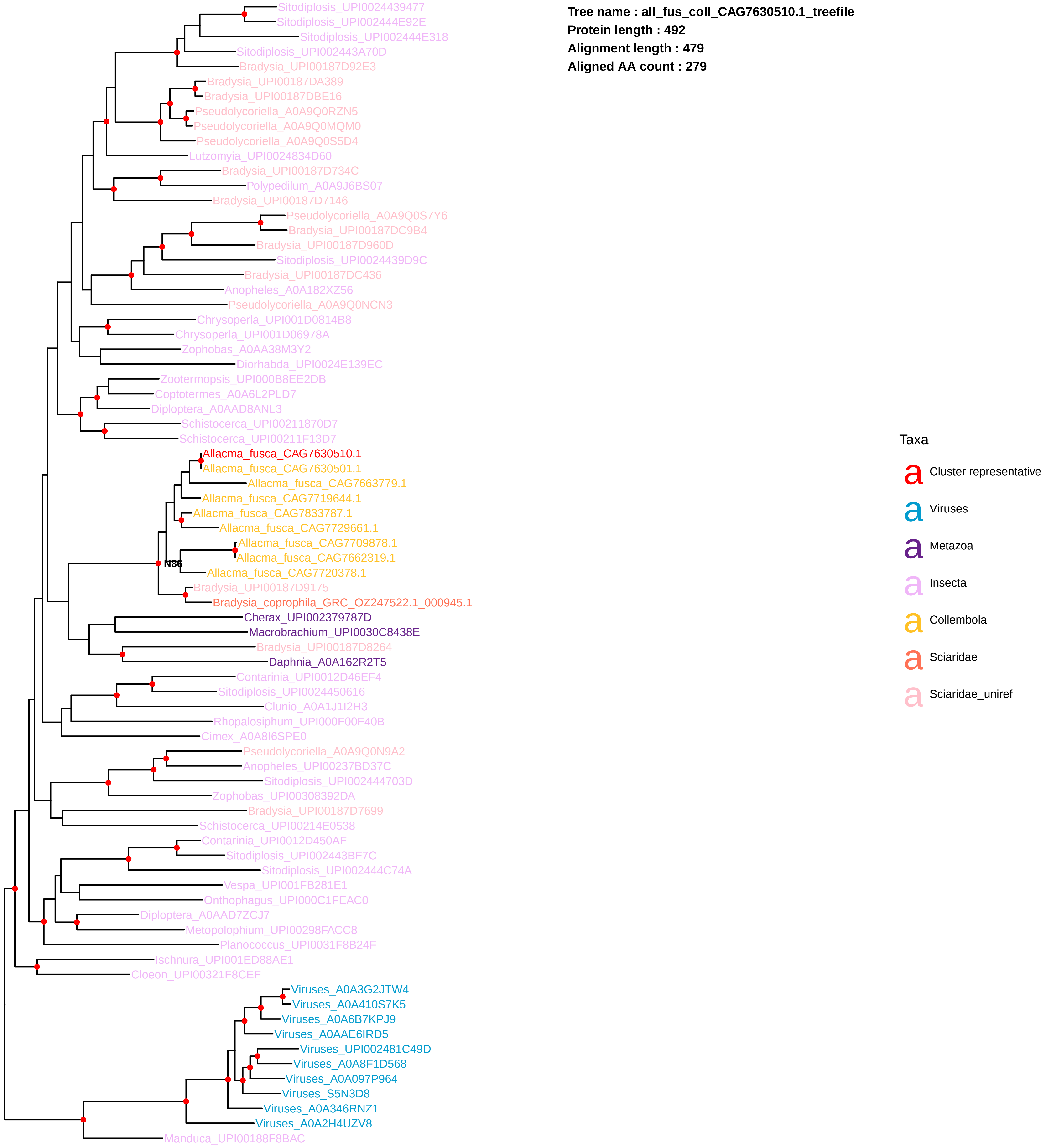
Tree name : all_fus_coll_CAG7630510.1_treefile
Protein length : 492
Alignment length : 479
Aligned AA count : 279
Sitodiplosis_UPI0024439477
Sitodiplosis_UPI002444E92E
Sitodiplosis_UPI002444E318
Sitodiplosis_UPI002443A70D
Bradysia_UPI00187D92E3
Bradysia_UPI00187DA389
Bradysia_UPI00187DBE16
Pseudolycoriella_A0A9Q0RZN5
Pseudolycoriella_A0A9Q0MQM0
Pseudolycoriella_A0A9Q0S5D4
Lutzomyia_UPI0024834D60
Bradysia_UPI00187D734C
Polypedilum_A0A9J6BS07
Bradysia_UPI00187D7146
Pseudolycoriella_A0A9Q0S7Y6
Bradysia_UPI00187DC9B4
Bradysia_UPI00187D960D
Sitodiplosis_UPI0024439D9C
Bradysia_UPI00187DC436
Anopheles_A0A182XZ56
Pseudolycoriella_A0A9Q0NCN3
Chrysoperla_UPI001D0814B8
Chrysoperla_UPI001D06978A
Zophobas_A0AA38M3Y2
Diorhabda_UPI0024E139EC
Zootermopsis_UPI000B8EE2DB
Coptotermes_A0A6L2PLD7
Diploptera_A0AAD8ANL3
Schistocerca_UPI00211870D7
Schistocerca_UPI00211F13D7
Allacma_fusca_CAG7630510.1
Allacma_fusca_CAG7630501.1
Allacma_fusca_CAG7663779.1
Allacma_fusca_CAG7719644.1
Allacma_fusca_CAG7833787.1
Allacma_fusca_CAG7729661.1
Allacma_fusca_CAG7709878.1
Allacma_fusca_CAG7662319.1
N86
Allacma_fusca_CAG7720378.1
Bradysia_UPI00187D9175
Bradysia_coprophila_GRC_OZ247522.1_000945.1
Cherax_UPI002379787D
Macrobrachium_UPI0030C8438E
Bradysia_UPI00187D8264
Daphnia_A0A162R2T5
Contarinia_UPI0012D46EF4
Sitodiplosis_UPI0024450616
Clunio_A0A1J1I2H3
Rhopalosiphum_UPI000F00F40B
Cimex_A0A8I6SPE0
Pseudolycoriella_A0A9Q0N9A2
Anopheles_UPI00237BD37C
Sitodiplosis_UPI002444703D
Zophobas_UPI00308392DA
Bradysia_UPI00187D7699
Schistocerca_UPI00214E0538
Contarinia_UPI0012D450AF
Sitodiplosis_UPI002443BF7C
Sitodiplosis_UPI002444C74A
Vespa_UPI001FB281E1
Onthophagus_UPI000C1FEAC0
Diploptera_A0AAD7ZCJ7
Metopolophium_UPI00298FACC8
Planococcus_UPI0031F8B24F
Ischnura_UPI001ED88AE1
Cloeon_UPI00321F8CEF
Viruses_A0A3G2JTW4
Viruses_A0A410S7K5
Viruses_A0A6B7KPJ9
Viruses_A0AAE6IRD5
Viruses_UPI002481C49D
Viruses_A0A8F1D568
Viruses_A0A097P964
Viruses_S5N3D8
Viruses_A0A346RNZ1
Viruses_A0A2H4UZV8
Manduca_UPI00188F8BAC
Taxa
a Cluster representative
a Viruses
a Metazoa
a Insecta
a Collembola
a Sciaridae
a Sciaridae_uniref

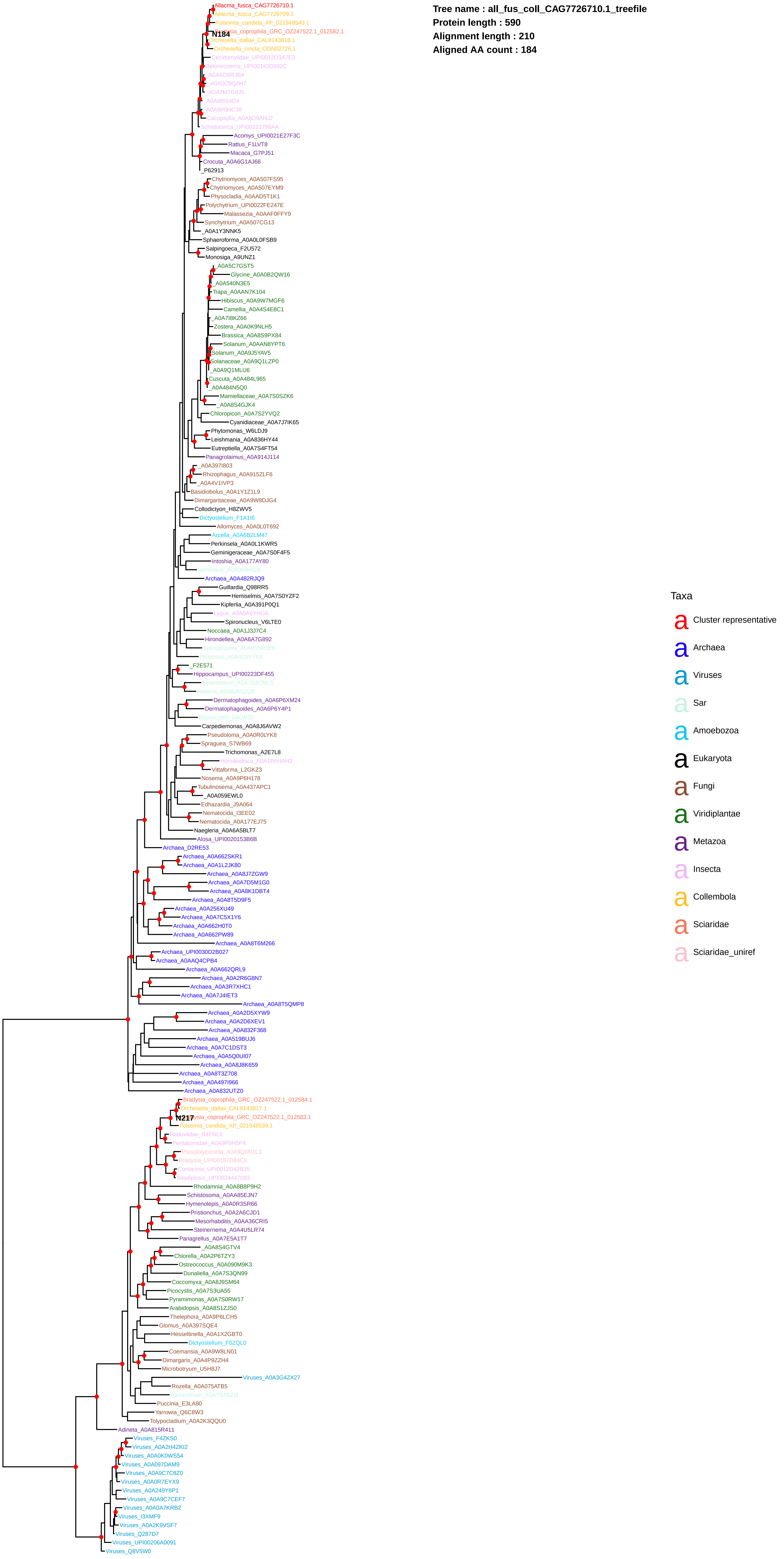
Tree name : all_fus_coll_CAG7726710.1_treefile
Protein length : 590
Alignment length : 210
Aligned AA count : 184
Allacma_fusca_CAG7726710.1
Allacma_fusca_CAG7726709.1
Folsomia_candida_XP_021948540.1
Bradysia_coprophila_GRC_OZ247522.1_012582.1
N184
Orchesella_dallaii_CAL8143818.1
Orchesella_cincta_ODN02726.1
Cecidomyiidae_UPI0012D3A7E3
Belonocnema_UPI00143D592C
_A0AAD9RJB4
_A0A0C9Q4H7
_A0A7M7G8J5
_A0A8I6S4D4
_A0A9P0HC38
Cacopsylla_A0A8D9AHJ2
Schistocerca_UPI00211798AA
Acomys_UPI0021E27F3C
Rattus_F1LVT8
Macaca_G7PJ51
Crocuta_A0A6G1AJ66
_P62913
Chytriomyces_A0A507FS95
Chytriomyces_A0A507EYM9
Physocladia_A0AAD5T1K1
Polychytrium_UPI0022FE247E
Malassezia_A0AAF0FFY9
Synchytrium_A0A507CG13
_A0A1Y3NNK5
Sphaeroforma_A0A0L0FSB9
Salpingoeca_F2U572
Monosiga_A9UNZ1
_A0A5C7GST5
Glycine_A0A0B2QW16
_A0A540N3E5
Trapa_A0AAN7K104
Hibiscus_A0A9W7MGF6
Camellia_A0A4S4E8C1
_A0A7I8KZ66
Zostera_A0A0K9NLH5
Brassica_A0A8S9PX84
Solanum_A0AAN8YPT6
Solanum_A0A9J5YAV5
Solanaceae_A0A9Q1LZP0
_A0A9Q1MLU6
Cuscuta_A0A484L965
_A0A484N5Q0
Mamiellaceae_A0A7S0SZK6
_A0A8S4GJK4
Chloropicon_A0A7S2YVQ2
Cyanidiaceae_A0A7J7IK65
Phytomonas_W6LDJ9
Leishmania_A0A836HY44
Eutreptiella_A0A7S4FT54
Panagrolaimus_A0A914J114
_A0A397I803
Rhizophagus_A0A915ZLF6
_A0A4V1IVP3
Basidiobolus_A0A1Y1Z1L9
Dimargaritaceae_A0A9W8DJG4
Collodictyon_H8ZWV5
Dictyostelium_F1A1I6
Allomyces_A0A0L0T692
Arcella_A0A6B2LM47
Perkinsela_A0A0L1KWR5
Geminigeraceae_A0A7S0F4F5
Intoshia_A0A177AY80
Seminavis_A0A9N8HI28
Archaea_A0A482RJQ9
Guillardia_Q98RR5
Hemiselmis_A0A7S0YZF2
Kipferlia_A0A391P0Q1
Lygus_A0A0A9YHG6
Spironucleus_V6LTE0
Noccaea_A0A1J3J7C4
Hirondellea_A0A6A7G892
Spongospora_A0A0H5R2E6
Perkinsus_A0A4D8Y7B4
_F2E571
Hippocampus_UPI00223DF455
Strombidium_A0A7S3CML5
Halteria_A0A8J8SZQ6
Dermatophagoides_A0A6P6XM24
Dermatophagoides_A0A6P6Y4P1
Blastocystis_D8LW32
Carpediemonas_A0A8J6AVW2
Pseudoloma_A0A0R0LYK8
Spraguea_S7WB69
Trichomonas_A2E7L8
Homalodisca_A0A1B6HAH2
Vittaforma_L2GKZ3
Nosema_A0A9P6H178
Tubulinosema_A0A437APC1
_A0A059EWL0
Edhazardia_J9A064
Nematocida_I3EE02
Nematocida_A0A177EJ75
Naegleria_A0A6A5BLT7
Alosa_UPI0020153B6B
Archaea_D2RE53
Archaea_A0A662SKR1
Archaea_A0A1L2JK80
Archaea_A0A8J7ZGW9
Archaea_A0A7D5M1G0
Archaea_A0A8K1DBT4
Archaea_A0A8T5D9F5
Archaea_A0A256XU49
Archaea_A0A7C5X1Y6
Archaea_A0A662H0T0
Archaea_A0A662PW89
Archaea_A0A8T6M266
Archaea_UPI0030D2B027
Archaea_A0AAQ4CPB4
Archaea_A0A662QRL9
Archaea_A0A2R6G8N7
Archaea_A0A3R7XHC1
Archaea_A0A7J4IET3
Archaea_A0A8T5QMP8
Archaea_A0A2D5XYW9
Archaea_A0A2D6XEV1
Archaea_A0A832F368
Archaea_A0A519BUJ6
Archaea_A0A7C1DST3
Archaea_A0A5Q0UI07
Archaea_A0A8J8K659
Archaea_A0A8T3Z708
Archaea_A0A497I966
Archaea_A0A832UTZ0
Bradysia_coprophila_GRC_OZ247522.1_012584.1
Orchesella_dallaii_CAL8143817.1
Bradysia_coprophila_GRC_OZ247522.1_012583.1
N217
Folsomia_candida_XP_021948539.1
Reduviidae_R4FNL6
Pentatomidae_A0A9P0H5P4
Pseudolycoriella_A0A9Q0N1L3
Bradysia_UPI00187D84C8
Contarinia_UPI0012D42B35
Sitodiplosis_UPI0024447093
Rhodamnia_A0A8B8P9H2
Schistosoma_A0AA85EJN7
Hymenolepis_A0A0R3SR66
Pristionchus_A0A2A6CJD1
Mesorhabditis_A0AA36CRI5
Steinernema_A0A4U5LR74
Panagrellus_A0A7E5A1T7
_A0A8S4GTV4
Chlorella_A0A2P6TZY3
Ostreococcus_A0A090M9K3
Dunaliella_A0A7S3QN99
Coccomyxa_A0A8J9SM64
Picocystis_A0A7S3UA55
Pyramimonas_A0A7S0RW17
Arabidopsis_A0A8S1ZJS0
Thelephora_A0A9P6LCH5
Glomus_A0A397SQE4
Hesseltinella_A0A1X2GBT0
Dictyostelium_F0ZQL0
Coemansia_A0A9W8LN01
Dimargaris_A0A4P9ZZH4
Microbotryum_U5H8J7
Viruses_A0A3G4ZX27
Rozella_A0A075ATB5
Alexandrium_A0A7S4S231
Puccinia_E3LA80
Yarrowia_Q6C8W3
Tolypocladium_A0A2K3QQU0
Adineta_A0A815R411
Viruses_F4ZKS0
Viruses_A0A2H4ZKI2
Viruses_A0A0K0WS54
Viruses_A0A097DAM9
Viruses_A0A9C7C8Z0
Viruses_A0A0R7EYX9
Viruses_A0A249Y6P1
Viruses_A0A9C7CEF7
Viruses_A0A0A7KRB2
Viruses_I3XMF9
Viruses_A0A2K9VSF7
Viruses_Q287D7
Viruses_UPI00206A0091
Viruses_Q8V5W0
Taxa
a Cluster representative
a Archaea
a Viruses
a Sar
a Amoebozoa
a Eukaryota
a Fungi
a Viridiplantae
a Metazoa
a Insecta
a Collembola
a Sciaridae
a Sciaridae_uniref

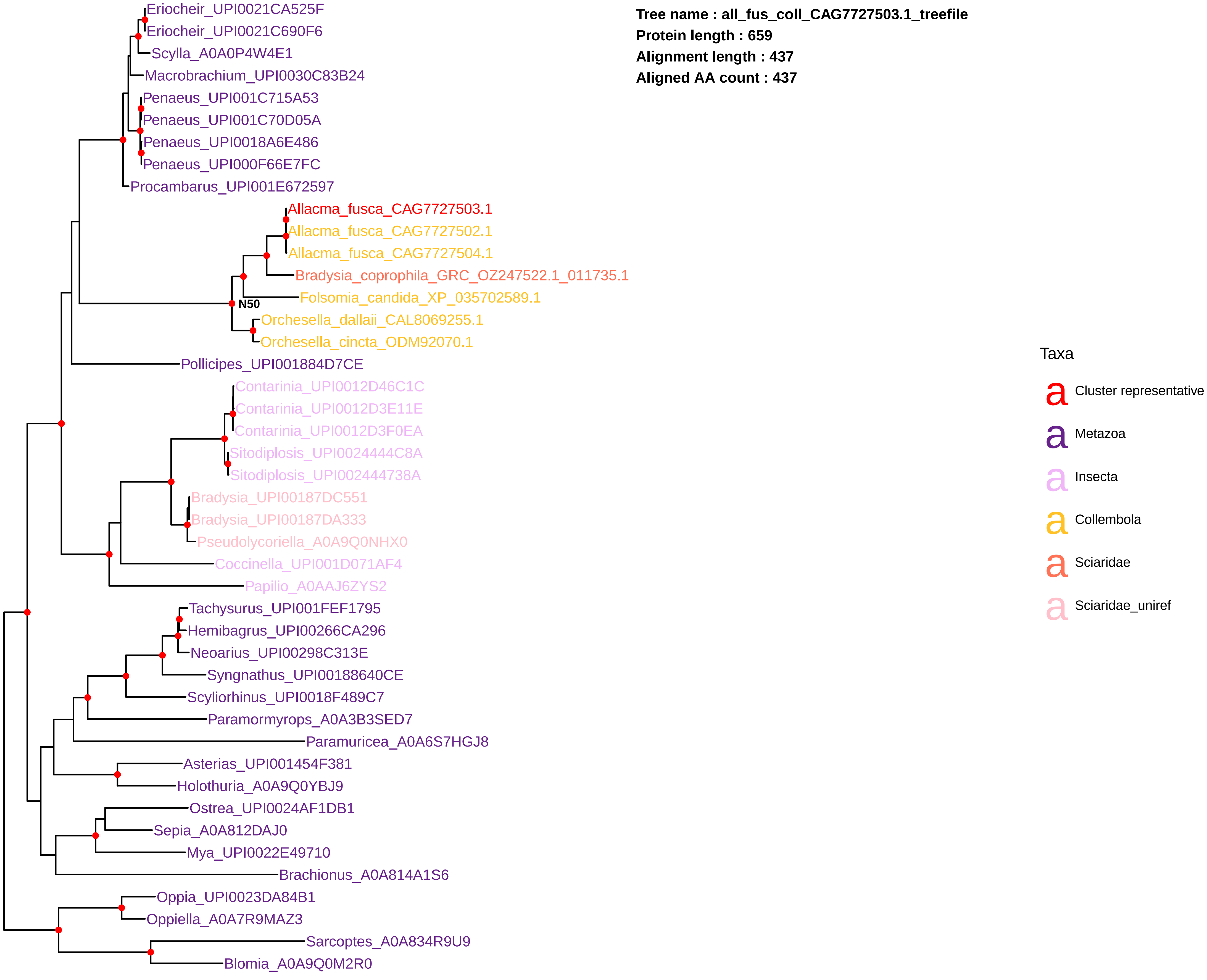
Tree name : all_fus_coll_CAG7727503.1_treefile
Protein length : 659
Alignment length : 437
Aligned AA count : 437
Eriocheir_UPI0021CA525F
Eriocheir_UPI0021C690F6
Scylla_A0A0P4W4E1
Macrobrachium_UPI0030C83B24
Penaeus_UPI001C715A53
Penaeus_UPI001C70D05A
Penaeus_UPI0018A6E486
Penaeus_UPI000F66E7FC
Procambarus_UPI001E672597
Allacma_fusca_CAG7727503.1
Allacma_fusca_CAG7727502.1
Allacma_fusca_CAG7727504.1
Bradysia_coprophila_GRC_OZ247522.1_011735.1
Folsomia_candida_XP_035702589.1
N50
Orchesella_dallaii_CAL8069255.1
Orchesella_cincta_ODM92070.1
Pollicipes_UPI001884D7CE
Contarinia_UPI0012D46C1C
Contarinia_UPI0012D3E11E
Contarinia_UPI0012D3F0EA
Sitodiplosis_UPI0024444C8A
Sitodiplosis_UPI002444738A
Bradysia_UPI00187DC551
Bradysia_UPI00187DA333
Pseudolycoriella_A0A9Q0NHX0
Coccinella_UPI001D071AF4
Papilio_A0AAJ6ZYS2
Tachysurus_UPI001FEF1795
Hemibagrus_UPI00266CA296
Neoarius_UPI00298C313E
Syngnathus_UPI00188640CE
Scyliorhinus_UPI0018F489C7
Paramormyrops_A0A3B3SED7
Paramuricea_A0A6S7HGJ8
Asterias_UPI001454F381
Holothuria_A0A9Q0YBJ9
Ostrea_UPI0024AF1DB1
Sepia_A0A812DAJ0
Mya_UPI0022E49710
Brachionus_A0A814A1S6
Oppia_UPI0023DA84B1
Oppiella_A0A7R9MAZ3
Sarcoptes_A0A834R9U9
Blomia_A0A9Q0M2R0
Taxa
Cluster representative
Metazoa
Insecta
Collembola
Sciaridae
Sciaridae_uniref

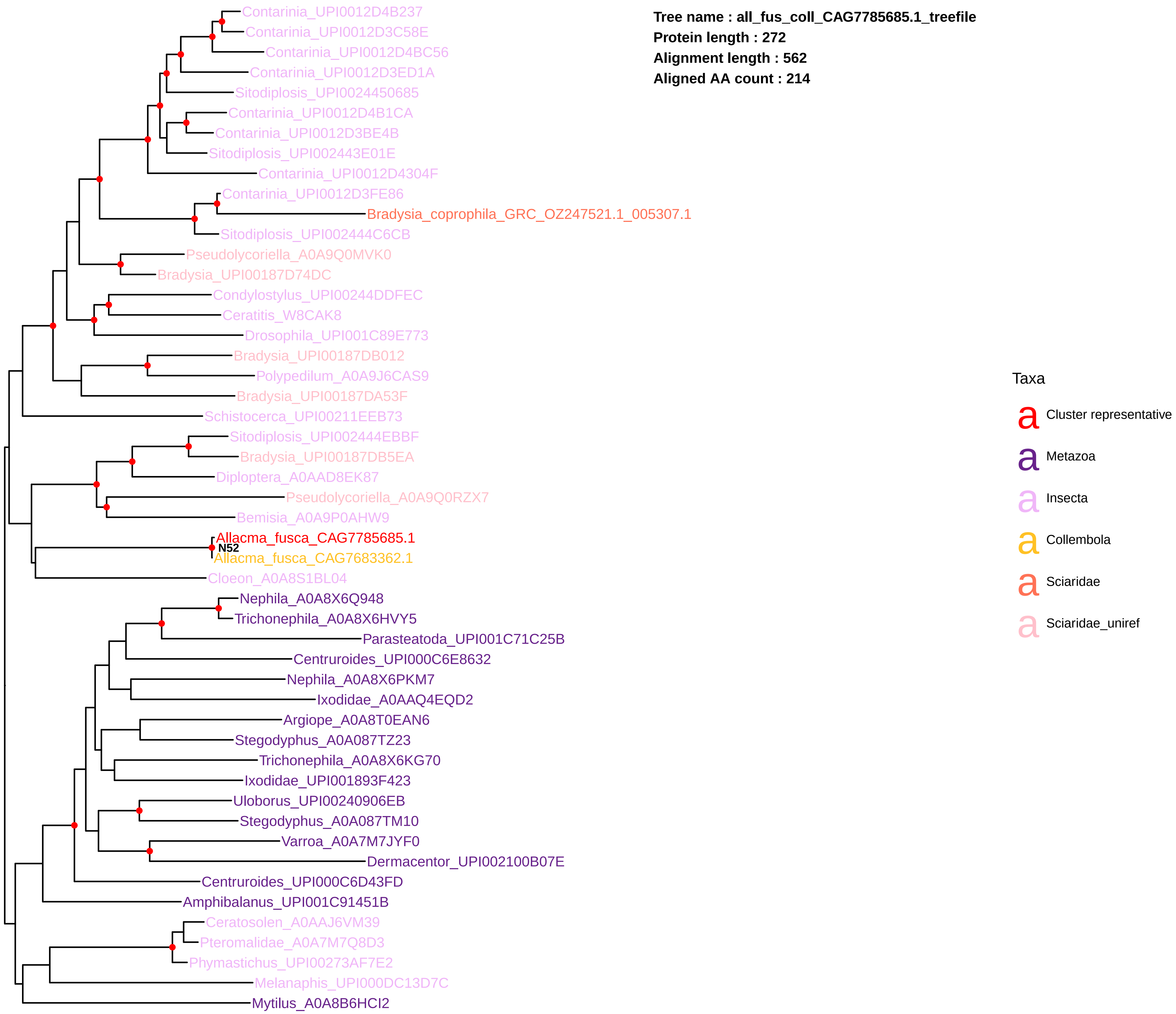
Tree name : all_fus_coll_CAG7785685.1_treefile
Protein length : 272
Alignment length : 562
Aligned AA count : 214
Contarinia_UPI0012D4B237
Contarinia_UPI0012D3C58E
Contarinia_UPI0012D4BC56
Contarinia_UPI0012D3ED1A
Sitodiplosis_UPI0024450685
Contarinia_UPI0012D4B1CA
Contarinia_UPI0012D3BE4B
Sitodiplosis_UPI002443E01E
Contarinia_UPI0012D4304F
Contarinia_UPI0012D3FE86
Bradysia_coprophila_GRC_OZ247521.1_005307.1
Sitodiplosis_UPI002444C6CB
Pseudolycoriella_A0A9Q0MVK0
Bradysia_UPI00187D74DC
Condylostylus_UPI00244DDFEC
Ceratitis_W8CAK8
Drosophila_UPI001C89E773
Bradysia_UPI00187DB012
Polypedilum_A0A9J6CAS9
Bradysia_UPI00187DA53F
Schistocerca_UPI00211EEB73
Sitodiplosis_UPI002444EBBF
Bradysia_UPI00187DB5EA
Diploptera_A0AAD8EK87
Pseudolycoriella_A0A9Q0RZX7
Bemisia_A0A9P0AHW9
Allacma_fusca_CAG7785685.1
N52
Allacma_fusca_CAG7683362.1
Cloeon_A0A8S1BL04
Nephila_A0A8X6Q948
Trichonephila_A0A8X6HVY5
Parasteatoda_UPI001C71C25B
Centruroides_UPI000C6E8632
Nephila_A0A8X6PKM7
Ixodidae_A0AAQ4EQD2
Argiope_A0A8T0EAN6
Stegodyphus_A0A087TZ23
Trichonephila_A0A8X6KG70
Ixodidae_UPI001893F423
Uloborus_UPI00240906EB
Stegodyphus_A0A087TM10
Varroa_A0A7M7JYF0
Dermacentor_UPI002100B07E
Centruroides_UPI000C6D43FD
Amphibalanus_UPI001C91451B
Ceratosolen_A0AAJ6VM39
Pteromalidae_A0A7M7Q8D3
Phymastichus_UPI00273AF7E2
Melanaphis_UPI000DC13D7C
Mytilus_A0A8B6HCI2
Taxa
Cluster representative
Metazoa
Insecta
Collembola
Sciaridae
Sciaridae_uniref

**Tree name : all_fus_coll_CAG7815295.1_treefile**
**Protein length : 733**
**Alignment length : 493**
**Aligned AA count : 471**

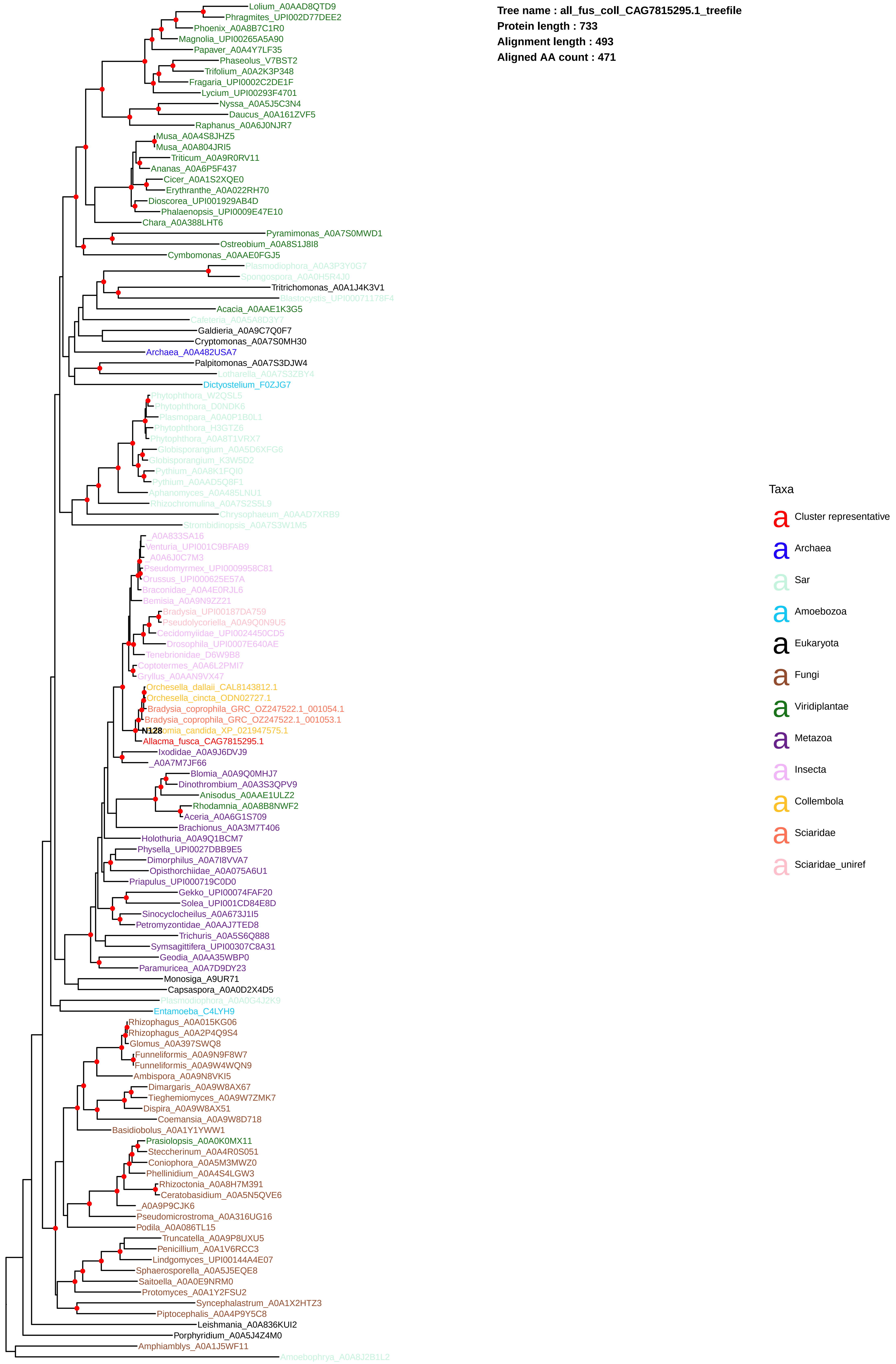

**Tree name : all_fus_coll_CAG7819770.1_treefile**
**Protein length : 522**
**Alignment length : 232**
**Aligned AA count : 232**

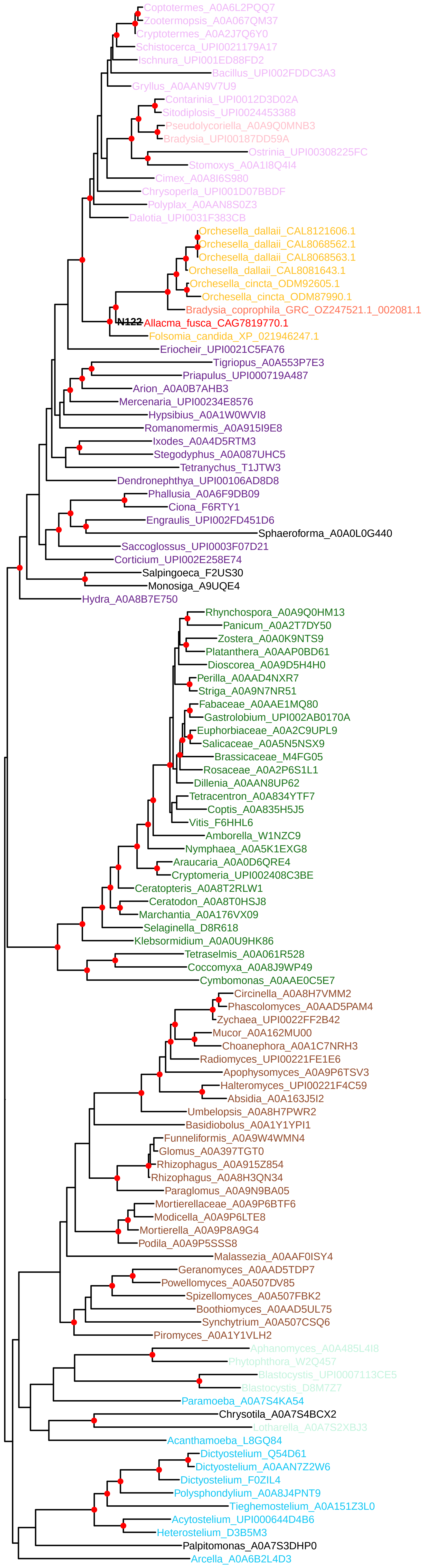


Taxa

- a Cluster representative
- a Sar
- a Amoebozoa
- a Eukaryota
- a Fungi
- a Viridiplantae
- a Metazoa
- a Insecta
- a Collembola
- a Sciaridae
- a Sciaridae_uniref

**Tree name : all_fus_coll_CAG7828195.1_treefile**
**Protein length : 558**
**Alignment length : 416**
**Aligned AA count : 416**

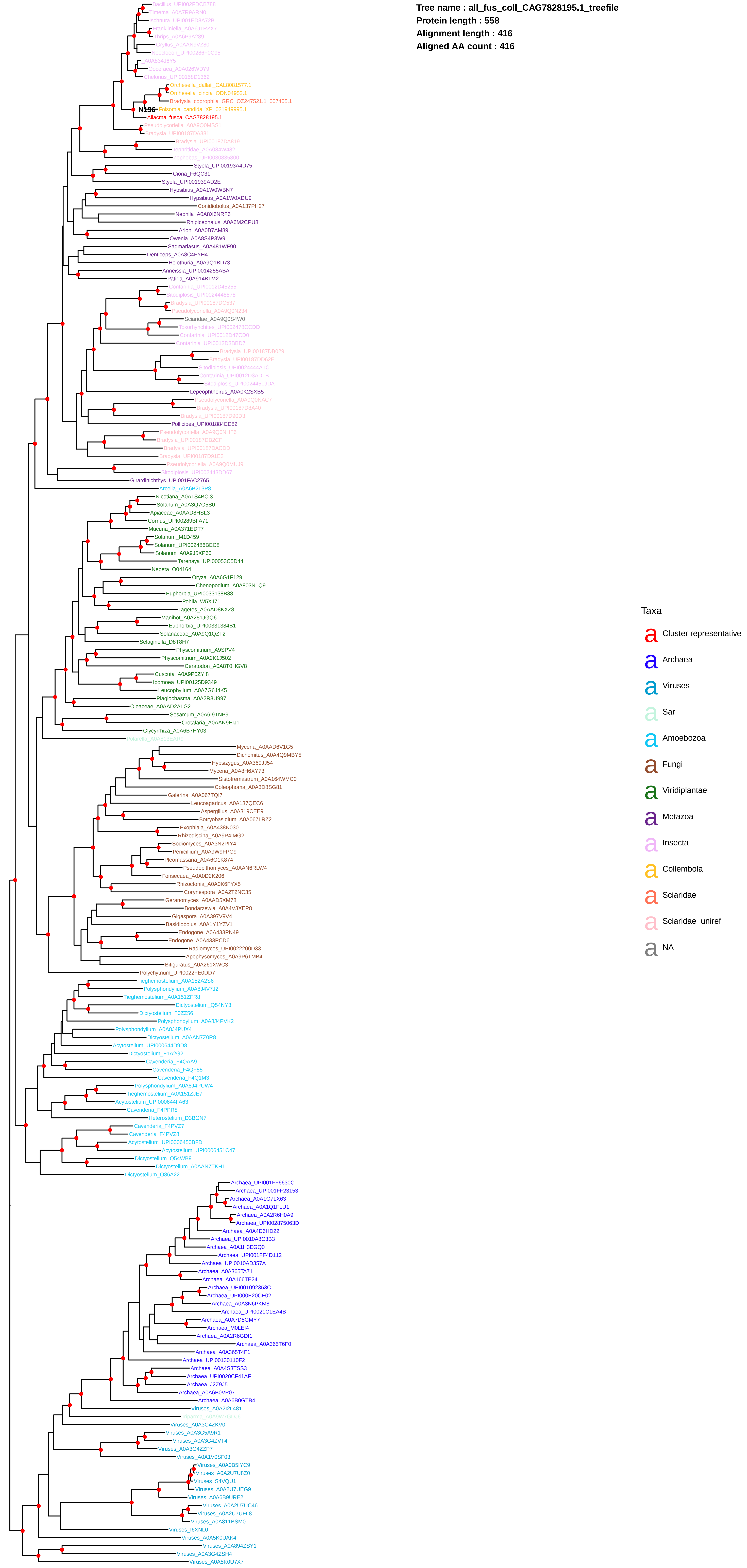

**Tree name : all_fus_coll_CAG7836705.1_treefile**
**Protein length : 212**
**Alignment length : 167**
**Aligned AA count : 156**

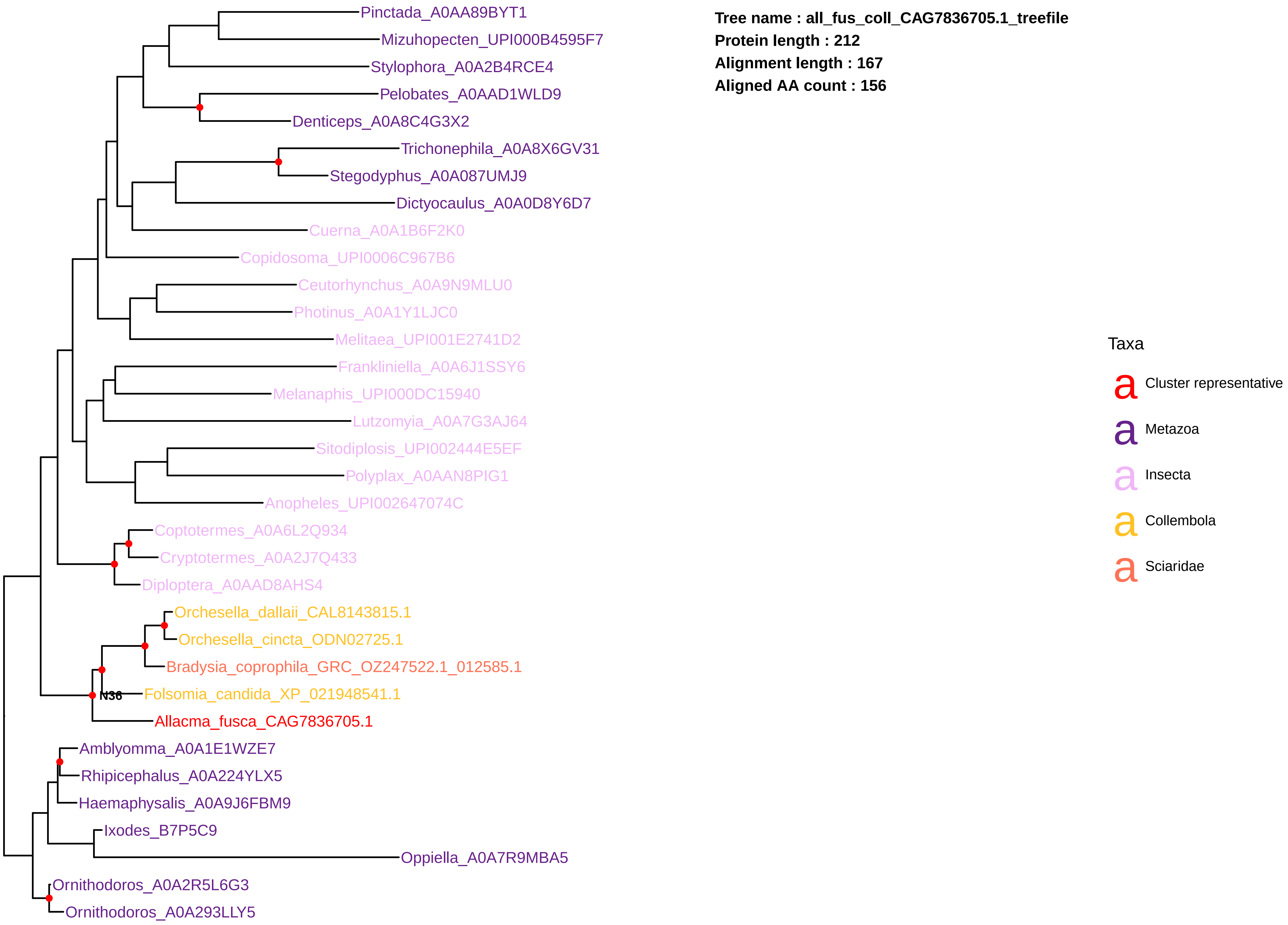

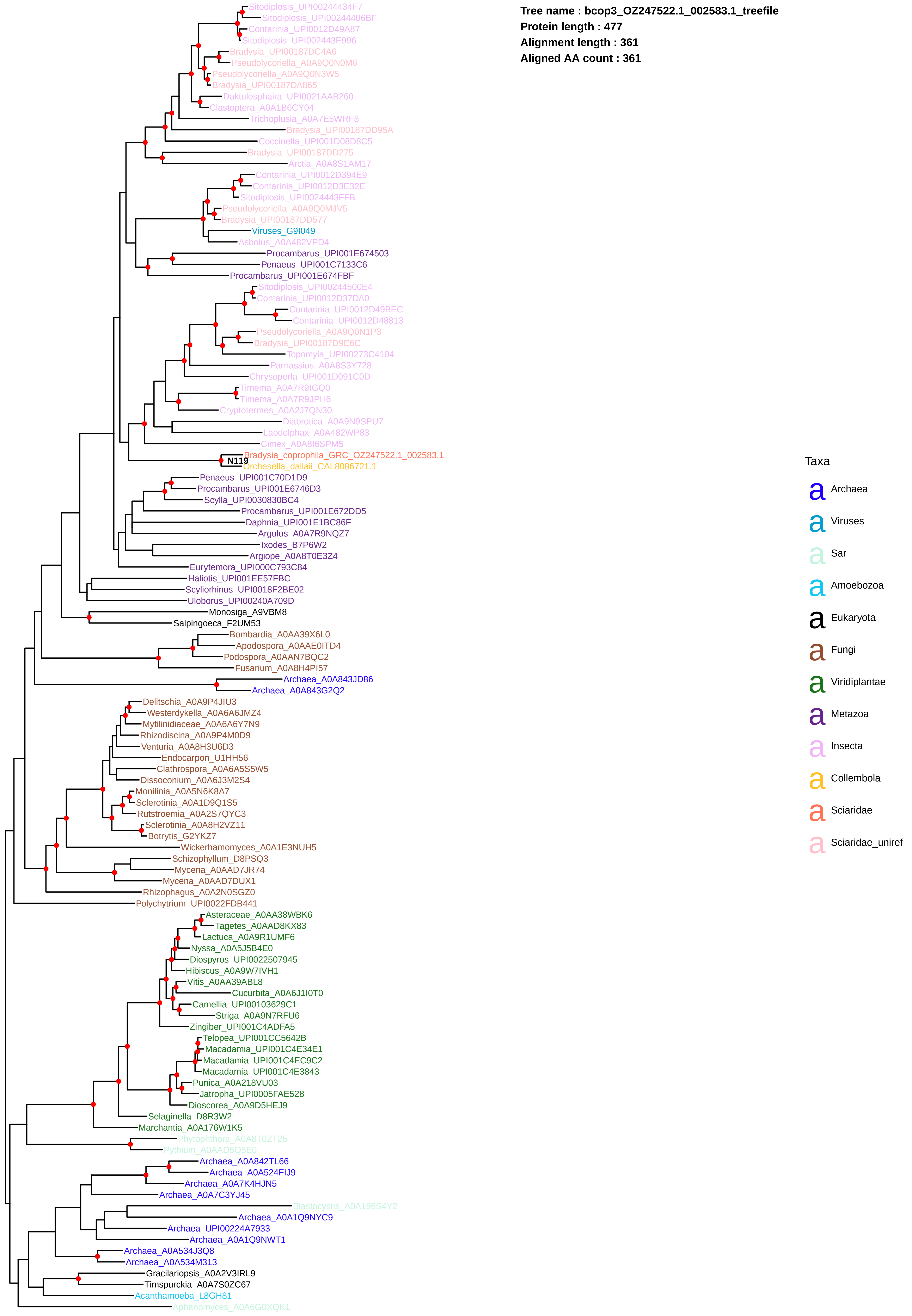
Tree name : bcop3_OZ247522.1_002583.1_treefile
Protein length : 477
Alignment length : 361
Aligned AA count : 361
Sitodiplosis_UPI00244434F7
Sitodiplosis_UPI00244406BF
Contarinia_UPI0012D49A87
Sitodiplosis_UPI002443E996
Bradysia_UPI00187DC4A6
Pseudolycoriella_A0A9Q0N0M6
Pseudolycoriella_A0A9Q0N3W5
Bradysia_UPI00187DA865
Daktulosphaira_UPI0021AAB260
Clastoptera_A0A1B6CY04
Trichoplusia_A0A7E5WRF8
Bradysia_UPI00187DD95A
Coccinella_UPI001D08D8C5
Bradysia_UPI00187DD275
Arctia_A0A8S1AM17
Contarinia_UPI0012D394E9
Contarinia_UPI0012D3E32E
Sitodiplosis_UPI0024443FFB
Pseudolycoriella_A0A9Q0MJV5
Bradysia_UPI00187DD577
Viruses_G9I049
Asbolus_A0A482VPD4
Procambarus_UPI001E674503
Penaeus_UPI001C7133C6
Procambarus_UPI001E674FBF
Sitodiplosis_UPI00244500E4
Contarinia_UPI0012D37DA0
Contarinia_UPI0012D49BEC
Contarinia_UPI0012D48813
Pseudolycoriella_A0A9Q0N1P3
Bradysia_UPI00187D9E6C
Topomyia_UPI00273C4104
Parnassius_A0A8S3Y728
Chrysoperla_UPI001D091C0D
Timema_A0A7R9IGQ0
Timema_A0A7R9JPH6
Cryptotermes_A0A2J7QN30
Diabrotica_A0A9N9SPU7
Laodelphax_A0A482WP83
Cimex_A0A8I6SPM5
Bradysia_coprophila_GRC_OZ247522.1_002583.1
N119
Orchesella_dallaii_CAL8086721.1
Penaeus_UPI001C70D1D9
Procambarus_UPI001E6746D3
Scylla_UPI0030830BC4
Procambarus_UPI001E672DD5
Daphnia_UPI001E1BC86F
Argulus_A0A7R9NQZ7
Ixodes_B7P6W2
Argiope_A0A8T0E3Z4
Eurytemora_UPI000C793C84
Haliotis_UPI001EE57FBC
Scyliorhinus_UPI0018F2BE02
Uloborus_UPI00240A709D
Monosiga_A9VBM8
Salpingoeca_F2UM53
Bombardia_A0AA39X6L0
Apodospora_A0AAE0ITD4
Podospora_A0AAN7BQC2
Fusarium_A0A8H4PI57
Archaea_A0A843JD86
Archaea_A0A843G2Q2
Delitschia_A0A9P4JIU3
Westerdykella_A0A6A6JMZ4
Mytilinidiaceae_A0A6A6Y7N9
Rhizodiscina_A0A9P4M0D9
Venturia_A0A8H3U6D3
Endocarpon_U1HH56
Clathrospora_A0A6A5S5W5
Dissoconium_A0A6J3M2S4
Monilinia_A0A5N6K8A7
Sclerotinia_A0A1D9Q1S5
Rutstroemia_A0A2S7QYC3
Sclerotinia_A0A8H2VZ11
Botrytis_G2YKZ7
Wickerhamomyces_A0A1E3NUH5
Schizophyllum_D8PSQ3
Mycena_A0AAD7JR74
Mycena_A0AAD7DUX1
Rhizophagus_A0A2N0SGZ0
Polychytrium_UPI0022FDB441
Asteraceae_A0AA38WBK6
Tagetes_A0AAD8KX83
Lactuca_A0A9R1UMF6
Nyssa_A0A5J5B4E0
Diospyros_UPI0022507945
Hibiscus_A0A9W7IVH1
Vitis_A0AA39ABL8
Cucurbita_A0A6J1I0T0
Camellia_UPI00103629C1
Striga_A0A9N7RFU6
Zingiber_UPI001C4ADFA5
Telopea_UPI001CC5642B
Macadamia_UPI001C4E34E1
Macadamia_UPI001C4EC9C2
Macadamia_UPI001C4E3843
Punica_A0A218VU03
Jatropha_UPI0005FAE528
Dioscorea_A0A9D5HEJ9
Selaginella_D8R3W2
Marchantia_A0A176W1K5
Phytophthora_A0A8T0ZT25
Pythium_A0AAD5Q5E0
Archaea_A0A842TL66
Archaea_A0A524FIJ9
Archaea_A0A7K4HJN5
Archaea_A0A7C3YJ45
Blastocystis_A0A196S4Y2
Archaea_A0A1Q9NYC9
Archaea_UPI00224A7933
Archaea_A0A1Q9NWT1
Archaea_A0A534J3Q8
Archaea_A0A534M313
Gracilariopsis_A0A2V3IRL9
Timspurckia_A0A7S0ZC67
Acanthamoeba_L8GH81
Aphanomyces_A0A6G0XQK1
Taxa
Archaea
Viruses
Sar
Amoebozoa
Eukaryota
Fungi
Viridiplantae
Metazoa
Insecta
Collembola
Sciaridae
Sciaridae_uniref

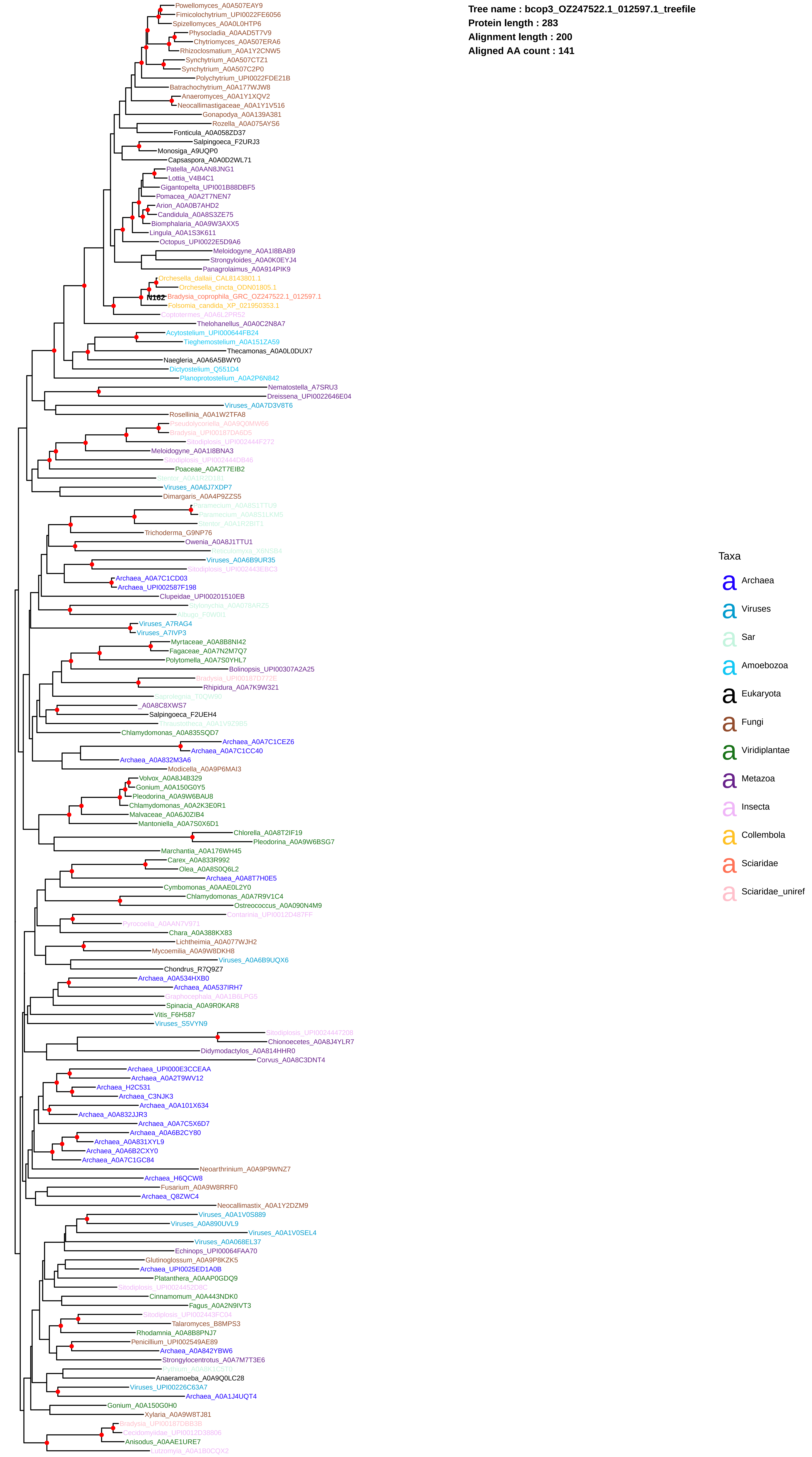
Tree name : bcop3_OZ247522.1_012597.1_treefile
Protein length : 283
Alignment length : 200
Aligned AA count : 141
Powellomyces_A0A507EAY9
Fimicolochytrium_UPI0022FE6056
Spizellomyces_A0A0L0HTP6
Physocladia_A0AAD5T7V9
Chytriomyces_A0A507ERA6
Rhizoclosmatium_A0A1Y2CNW5
Synchytrium_A0A507CTZ1
Synchytrium_A0A507C2P0
Polychytrium_UPI0022FDE21B
Batrachochytrium_A0A177WJW8
Anaeromyces_A0A1Y1XQV2
Neocallimastigaceae_A0A1Y1V516
Gonapodya_A0A139A381
Rozella_A0A075AYS6
Fonticula_A0A058ZD37
Salpingoeca_F2URJ3
Monosiga_A9UQP0
Capsaspora_A0A0D2WL71
Patella_A0AAN8JNG1
Lottia_V4B4C1
Gigantopelta_UPI001B88DBF5
Pomacea_A0A2T7NEN7
Arion_A0A0B7AHD2
Candidula_A0A8S3ZE75
Biomphalaria_A0A9W3AXX5
Lingula_A0A1S3K611
Octopus_UPI0022E5D9A6
Meloidogyne_A0A1I8BAB9
Strongyloides_A0A0K0EYJ4
Panagrolaimus_A0A914PIK9
Orchesella_dallaii_CAL8143801.1
Orchesella_cincta_ODN01805.1
N162
Bradysia_coprophila_GRC_OZ247522.1_012597.1
Folsomia_candida_XP_021950353.1
Coptotermes_A0A6L2PR52
Thelohanellus_A0A0C2N8A7
Acytostelium_UPI000644FB24
Tieghemostelium_A0A151ZA59
Thecamonas_A0A0L0DUX7
Naegleria_A0A6A5BWY0
Dictyostelium_Q551D4
Planoprotostelium_A0A2P6N842
Nematostella_A7SRU3
Dreissena_UPI0022646E04
Viruses_A0A7D3V8T6
Rosellinia_A0A1W2TFA8
Pseudolycoriella_A0A9Q0MW66
Bradysia_UPI00187DA6D5
Sitodiplosis_UPI002444F272
Meloidogyne_A0A1I8BNA3
Sitodiplosis_UPI002444DB46
Poaceae_A0A2T7EIB2
Stentor_A0A1R2D181
Viruses_A0A6J7XDP7
Dimargaris_A0A4P9ZZS5
Paramecium_A0A8S1TTU9
Paramecium_A0A8S1LKM5
Stentor_A0A1R2BIT1
Trichoderma_G9NP76
Owenia_A0A8J1TTU1
Reticulomyxa_X6NSB4
Viruses_A0A6B9UR35
Sitodiplosis_UPI002443EBC3
Archaea_A0A7C1CD03
Archaea_UPI002587F198
Clupeidae_UPI00201510EB
Stylonychia_A0A078ARZ5
Albugo_F0W0I1
Viruses_A7RAG4
Viruses_A7IVP3
Myrtaceae_A0A8B8NI42
Fagaceae_A0A7N2M7Q7
Polytomella_A0A7S0YHL7
Bolinopsis_UPI00307A2A25
Bradysia_UPI00187D772E
Rhipidura_A0A7K9W321
Saprolegnia_T0QW90
_A0A8C8XWS7
Salpingoeca_F2UEH4
Thraustotheca_A0A1V9Z9B5
Chlamydomonas_A0A835SQD7
Archaea_A0A7C1CEZ6
Archaea_A0A7C1CC40
Archaea_A0A832M3A6
Modicella_A0A9P6MAI3
Volvox_A0A8J4B329
Gonium_A0A150G0Y5
Pleodorina_A0A9W6BAU8
Chlamydomonas_A0A2K3E0R1
Malvaceae_A0A6J0ZIB4
Mantoniella_A0A7S0X6D1
Chlorella_A0A8T2IF19
Pleodorina_A0A9W6BSG7
Marchantia_A0A176WH45
Carex_A0A833R992
Olea_A0A8S0Q6L2
Archaea_A0A8T7H0E5
Cymbomonas_A0AAE0L2Y0
Chlamydomonas_A0A7R9V1C4
Ostreococcus_A0A090N4M9
Contarinia_UPI0012D487FF
Pyrocoelia_A0AAN7V971
Chara_A0A388KX83
Lichtheimia_A0A077WJH2
Mycoemilia_A0A9W8DKH8
Viruses_A0A6B9UQX6
Chondrus_R7Q9Z7
Archaea_A0A534HXB0
Archaea_A0A537IRH7
Graphocephala_A0A1B6LPG5
Spinacia_A0A9R0KAR8
Vitis_F6H587
Viruses_S5VYN9
Sitodiplosis_UPI0024447208
Chionoecetes_A0A8J4YLR7
Didymodactylos_A0A814HHR0
Corvus_A0A8C3DNT4
Archaea_UPI000E3CCEAA
Archaea_A0A2T9WV12
Archaea_H2C531
Archaea_C3NJK3
Archaea_A0A101X634
Archaea_A0A832JJR3
Archaea_A0A7C5X6D7
Archaea_A0A6B2CY80
Archaea_A0A831XYL9
Archaea_A0A6B2CXY0
Archaea_A0A7C1GC84
Neoarthrinium_A0A9P9WNZ7
Archaea_H6QCW8
Fusarium_A0A9W8RRF0
Archaea_Q8ZWC4
Neocallimastix_A0A1Y2DZM9
Viruses_A0A1V0S889
Viruses_A0A890UVL9
Viruses_A0A1V0SEL4
Viruses_A0A068EL37
Echinops_UPI00064FAA70
Glutinoglossum_A0A9P8KZK5
Archaea_UPI0025ED1A0B
Platanthera_A0AAP0GDQ9
Sitodiplosis_UPI0024452D8C
Cinnamomum_A0A443NDK0
Fagus_A0A2N9IVT3
Sitodiplosis_UPI002443FC04
Talaromyces_B8MPS3
Rhodamnia_A0A8B8PNJ7
Penicillium_UPI002549AE89
Archaea_A0A842YBW6
Strongylocentrotus_A0A7M7T3E6
Pythium_A0A8K1C5T0
Anaeramoeba_A0A9Q0LC28
Viruses_UPI00226C63A7
Archaea_A0A1J4UQT4
Gonium_A0A150G0H0
Xylaria_A0A9W8TJ81
Bradysia_UPI00187DBB3B
Cecidomyiidae_UPI0012D38806
Anisodus_A0AAE1URE7
Lutzomyia_A0A1B0CQX2
Taxa
a Archaea
a Viruses
a Sar
a Amoebozoa
a Eukaryota
a Fungi
a Viridiplantae
a Metazoa
a Insecta
a Collembola
a Sciaridae
a Sciaridae_uniref

**Tree name : fol_can_coll_XP_021948097.1_treefile**
**Protein length : 188**
**Alignment length : 129**
**Aligned AA count : 129**

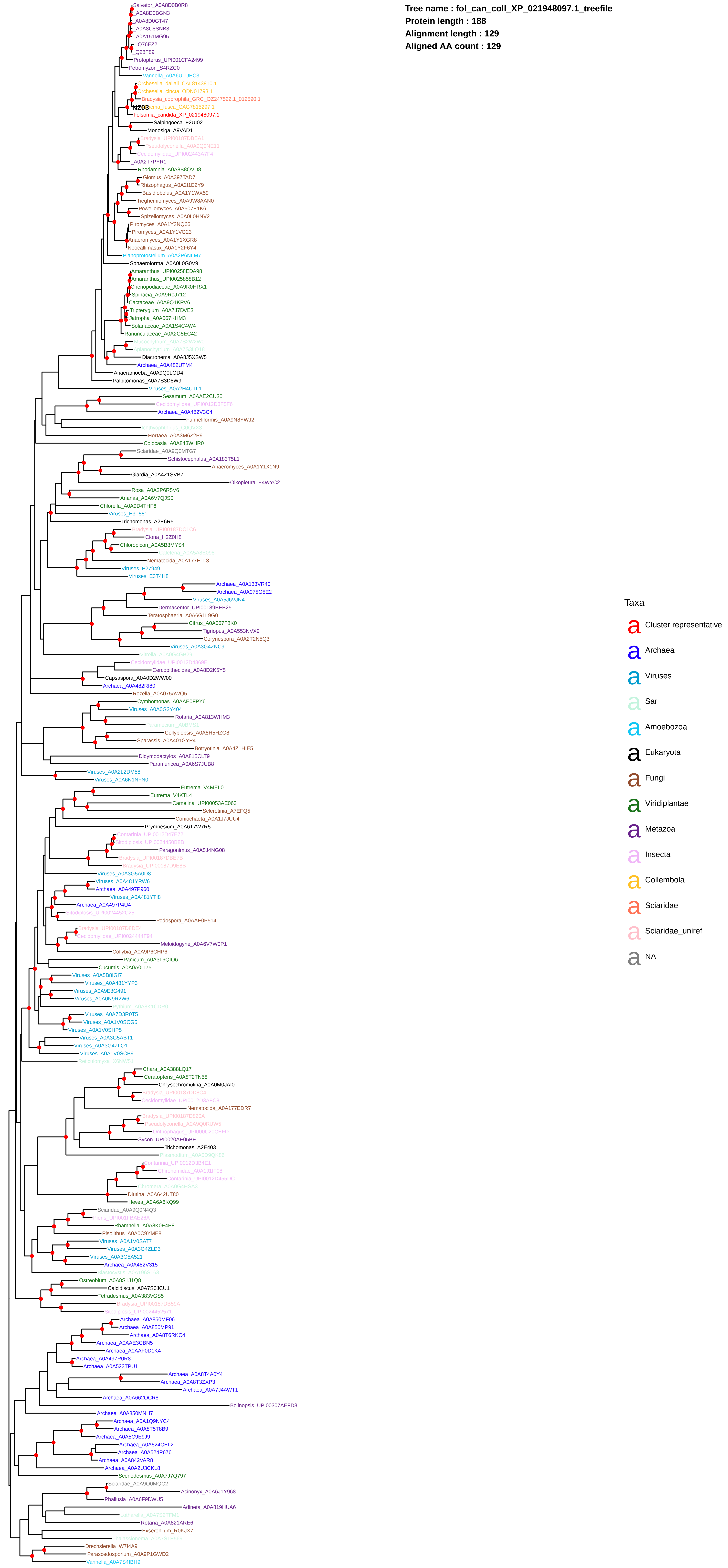

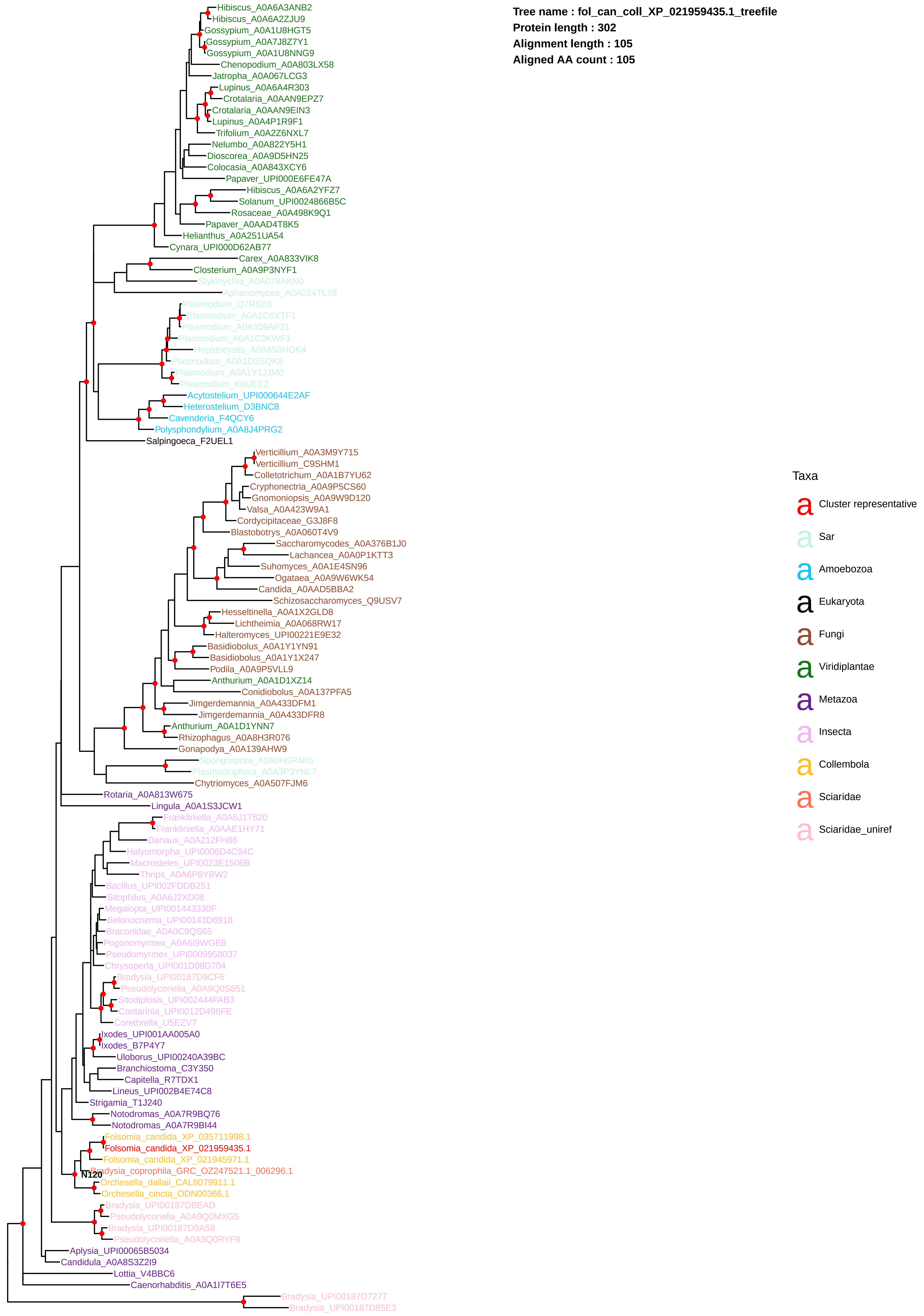

Tree name : fol_can_coll_XP_021959435.1_treefile
Protein length : 302
Alignment length : 105
Aligned AA count : 105
Hibiscus_A0A6A3ANB2
Hibiscus_A0A6A2ZJU9
Gossypium_A0A1U8HGT5
Gossypium_A0A7J8Z7Y1
Gossypium_A0A1U8NNG9
Chenopodium_A0A803LX58
Jatropha_A0A067LCG3
Lupinus_A0A6A4R303
Crotalaria_A0AAN9EPZ7
Crotalaria_A0AAN9EIN3
Lupinus_A0A4P1R9F1
Trifolium_A0A2Z6NXL7
Nelumbo_A0A822Y5H1
Dioscorea_A0A9D5HN25
Colocasia_A0A843XCY6
Papaver_UPI000E6FE47A
Hibiscus_A0A6A2YFZ7
Solanum_UPI0024866B5C
Rosaceae_A0A498K9Q1
Papaver_A0AAD4T8K5
Helianthus_A0A251UA54
Cynara_UPI000D62AB77
Carex_A0A833VIK8
Closterium_A0A9P3NYF1
Stylonychia_A0A078AKN0
Aphanomyces_A0A024TL78
Plasmodium_Q7RSE6
Plasmodium_A0A1C6XTF1
Plasmodium_A0A509AP31
Plasmodium_A0A1C3KWF1
Hepatocystis_A0A653HGK4
Plasmodium_A0A1D3SQK9
Plasmodium_A0A1Y1JJM0
Plasmodium_K6UEE2
Acytostelium_UPI000644E2AF
Heterostelium_D3BNC8
Cavenderia_F4QCY6
Polysphondylium_A0A8J4PRG2
Salpingoeca_F2UEL1
Verticillium_A0A3M9Y715
Verticillium_C9SHM1
Colletotrichum_A0A1B7YU62
Cryphonectria_A0A9P5CS60
Gnomoniopsis_A0A9W9D120
Valsa_A0A423W9A1
Cordycipitaceae_G3J8F8
Blastobotrys_A0A060T4V9
Saccharomycodes_A0A376B1J0
Lachancea_A0A0P1KTT3
Suhomyces_A0A1E4SN96
Ogataea_A0A9W6WK54
Candida_A0AAD5BBA2
Schizosaccharomyces_Q9USV7
Hesseltinella_A0A1X2GLD8
Lichtheimia_A0A068RW17
Halteromyces_UPI00221E9E32
Basidiobolus_A0A1Y1YN91
Basidiobolus_A0A1Y1X247
Podila_A0A9P5VLL9
Anthurium_A0A1D1XZ14
Conidiobolus_A0A137PFA5
Jimgerdemannia_A0A433DFM1
Jimgerdemannia_A0A433DFR8
Anthurium_A0A1D1YNN7
Rhizophagus_A0A8H3R076
Gonapodya_A0A139AHW9
Spongospora_A0A0H5R4K6
Plasmodiophora_A0A3P3YNL7
Chytriomyces_A0A507FJM6
Rotaria_A0A813W675
Lingula_A0A1S3JCW1
Frankliniella_A0A6J1T620
Frankliniella_A0AAE1HY71
Danaus_A0A212FH86
Halyomorpha_UPI0006D4C94C
Macrosteles_UPI0023E1506B
Thrips_A0A6P8YBW2
Bacillus_UPI002FDDB251
Sitophilus_A0A6J2XD08
Megalopta_UPI001443330F
Belonocnema_UPI00143D6918
Braconidae_A0A0C9QS65
Pogonomyrmex_A0A6I9WGE8
Pseudomyrmex_UPI0009958037
Chrysoperla_UPI001D08D704
Bradysia_UPI00187D9CF6
Pseudolycoriella_A0A9Q0S851
Sitodiplosis_UPI002444FAB3
Contarinia_UPI0012D498FE
Corethrella_U5EZV7
Ixodes_UPI001AA005A0
Ixodes_B7P4Y7
Uloborus_UPI00240A39BC
Branchiostoma_C3Y350
Capitella_R7TDX1
Lineus_UPI002B4E74C8
Strigamia_T1J240
Notodromas_A0A7R9BQ76
Notodromas_A0A7R9BI44
Folsomia_candida_XP_035711998.1
Folsomia_candida_XP_021959435.1
Folsomia_candida_XP_021945971.1
Bradysia_coprophila_GRC_OZ247521.1_006296.1
N120
Orchesella_dallaii_CAL8079911.1
Orchesella_cincta_ODN00366.1
Bradysia_UPI00187DBEAD
Pseudolycoriella_A0A9Q0MXG5
Bradysia_UPI00187D9A58
Pseudolycoriella_A0A9Q0RYF9
Aplysia_UPI00065B5034
Candidula_A0A8S3Z2I9
Lottia_V4BBC6
Caenorhabditis_A0A1I7T6E5
Bradysia_UPI00187D7277
Bradysia_UPI00187D85E3
Taxa
Cluster representative
Sar
Amoebozoa
Eukaryota
Fungi
Viridiplantae
Metazoa
Insecta
Collembola
Sciaridae
Sciaridae_uniref

**Tree name : fol_can_coll_XP_021959935.1_treefile**
**Protein length : 921**
**Alignment length : 331**
**Aligned AA count : 331**

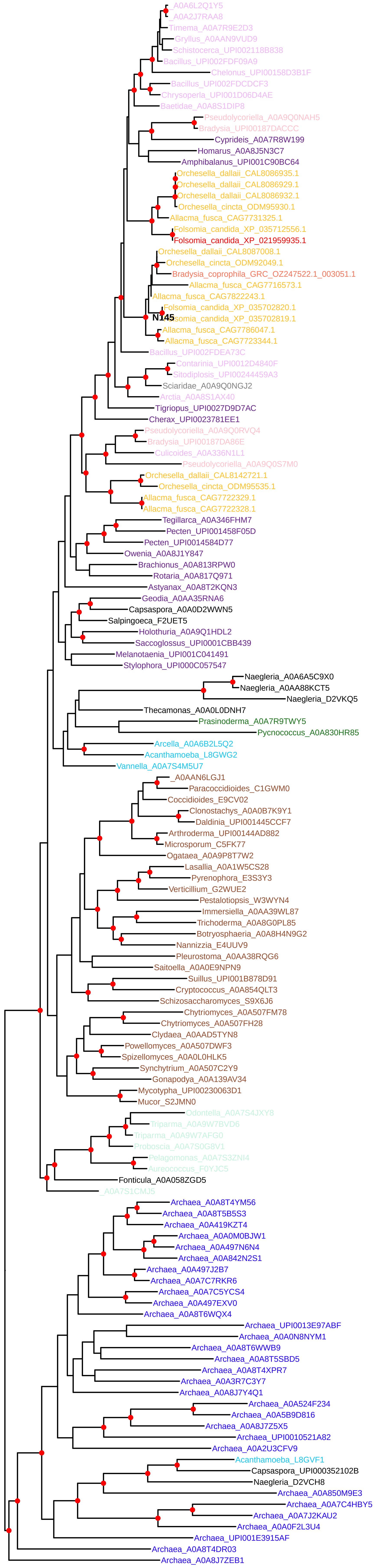


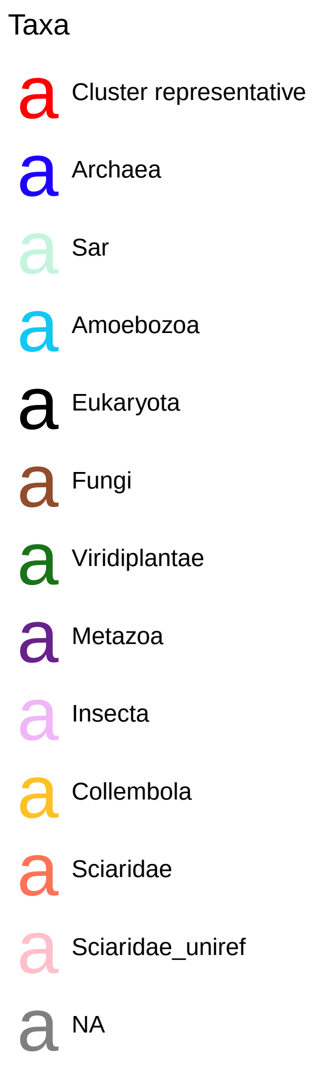

**Tree name : fol_can_coll_XP_021961662.1_treefile**
**Protein length : 1474**
**Alignment length : 475**
**Aligned AA count : 475**

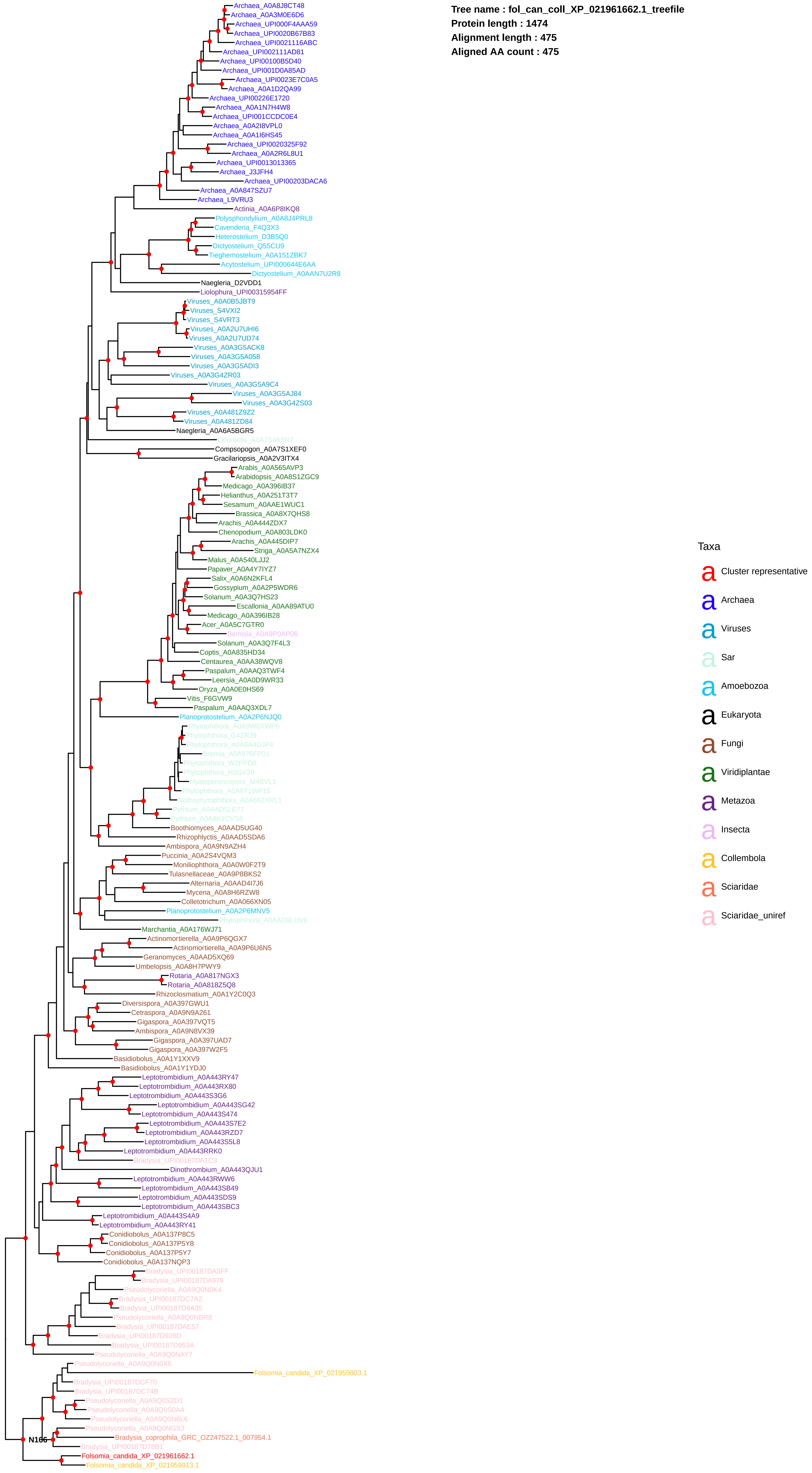

**Tree name : fol_can_coll_XP_021966621.1_treefile**
**Protein length : 753**
**Alignment length : 370**
**Aligned AA count : 370**

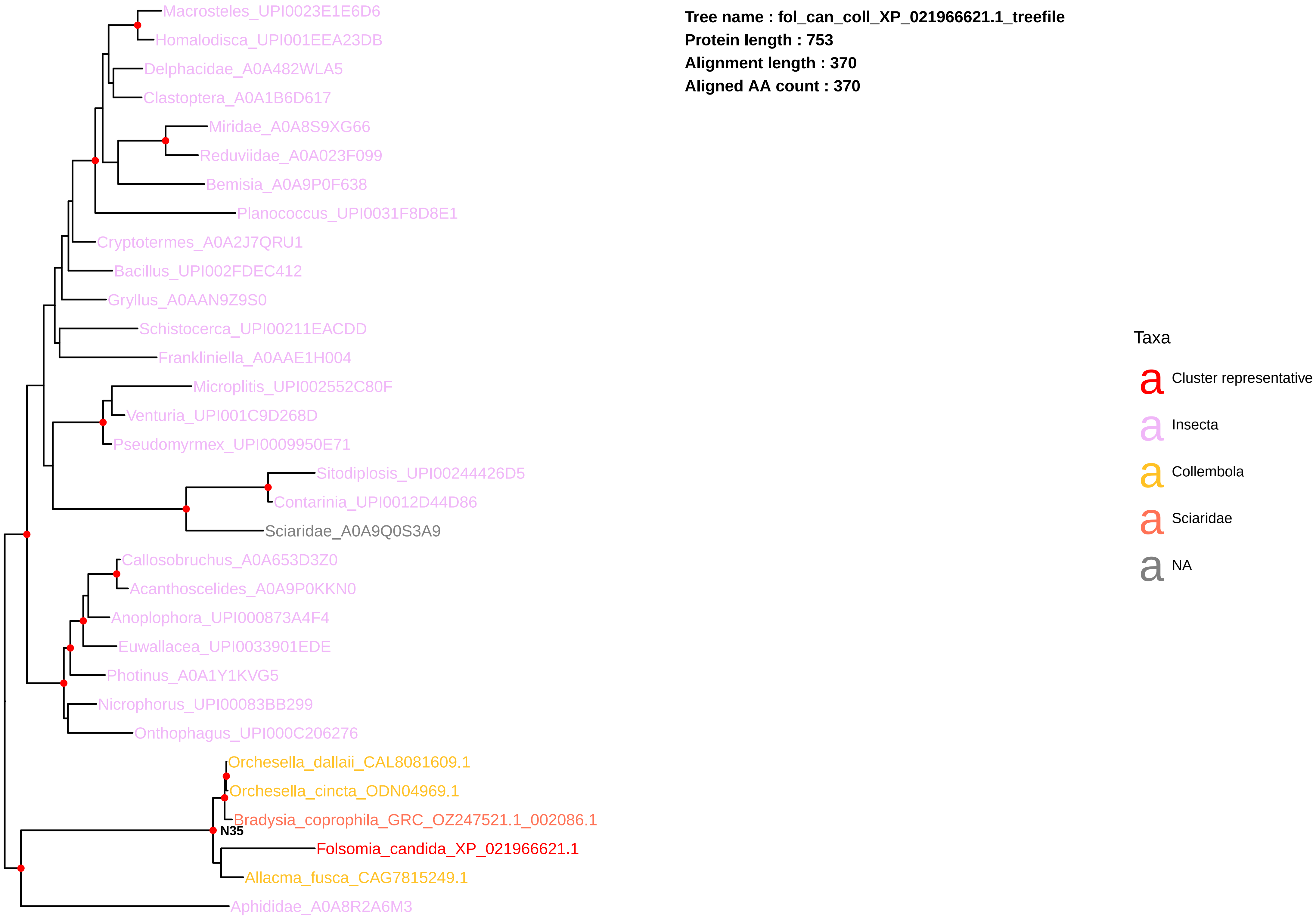

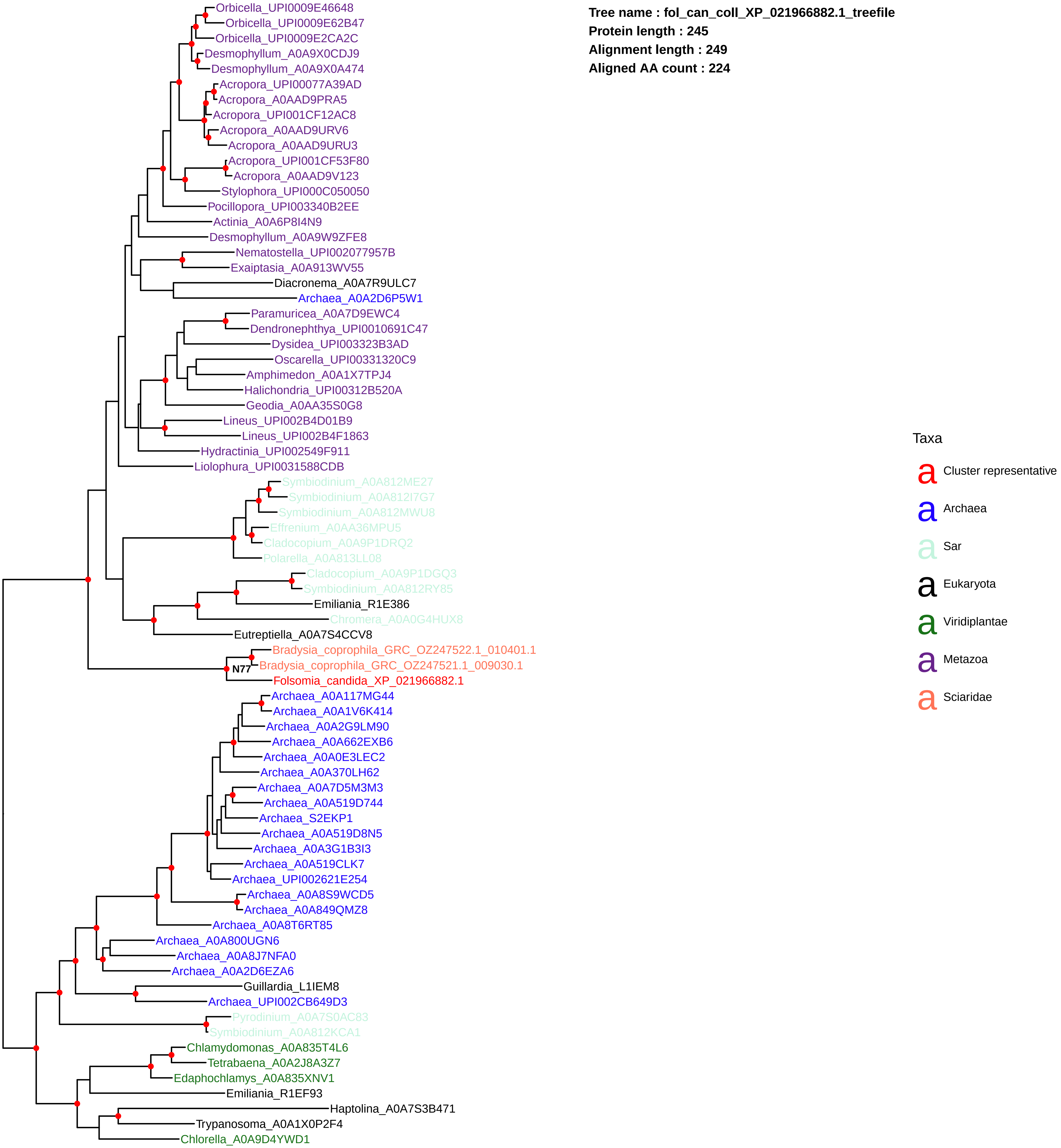

Tree name : fol_can_coll_XP_021966882.1_treefile
Protein length : 245
Alignment length : 249
Aligned AA count : 224
Orbicella_UPI0009E46648
Orbicella_UPI0009E62B47
Orbicella_UPI0009E2CA2C
Desmophyllum_A0A9X0CDJ9
Desmophyllum_A0A9X0A474
Acropora_UPI00077A39AD
Acropora_A0AAD9PRA5
Acropora_UPI001CF12AC8
Acropora_A0AAD9URV6
Acropora_A0AAD9URU3
Acropora_UPI001CF53F80
Acropora_A0AAD9V123
Stylophora_UPI000C050050
Pocillopora_UPI003340B2EE
Actinia_A0A6P8I4N9
Desmophyllum_A0A9W9ZFE8
Nematostella_UPI002077957B
Exaiptasia_A0A913WV55
Diacronema_A0A7R9ULC7
Archaea_A0A2D6P5W1
Paramuricea_A0A7D9EWC4
Dendronephthya_UPI0010691C47
Dysidea_UPI003323B3AD
Oscarella_UPI00331320C9
Amphimedon_A0A1X7TPJ4
Halichondria_UPI00312B520A
Geodia_A0AA35S0G8
Lineus_UPI002B4D01B9
Lineus_UPI002B4F1863
Hydractinia_UPI002549F911
Liolophura_UPI0031588CDB
Symbiodinium_A0A812ME27
Symbiodinium_A0A812I7G7
Symbiodinium_A0A812MWU8
Effrenium_A0AA36MPU5
Cladocopium_A0A9P1DRQ2
Polarella_A0A813LL08
Cladocopium_A0A9P1DGQ3
Symbiodinium_A0A812RY85
Emiliania_R1E386
Chromera_A0A0G4HUX8
Eutreptiella_A0A7S4CCV8
Bradysia_coprophila_GRC_OZ247522.1_010401.1
N77
Bradysia_coprophila_GRC_OZ247521.1_009030.1
Folsomia_candida_XP_021966882.1
Archaea_A0A117MG44
Archaea_A0A1V6K414
Archaea_A0A2G9LM90
Archaea_A0A662EXB6
Archaea_A0A0E3LEC2
Archaea_A0A370LH62
Archaea_A0A7D5M3M3
Archaea_A0A519D744
Archaea_S2EKP1
Archaea_A0A519D8N5
Archaea_A0A3G1B3I3
Archaea_A0A519CLK7
Archaea_UPI002621E254
Archaea_A0A8S9WCD5
Archaea_A0A849QMZ8
Archaea_A0A8T6RT85
Archaea_A0A800UGN6
Archaea_A0A8J7NFA0
Archaea_A0A2D6EZA6
Guillardia_L1IEM8
Archaea_UPI002CB649D3
Pyrodinium_A0A7S0AC83
Symbiodinium_A0A812KCA1
Chlamydomonas_A0A835T4L6
Tetrabaena_A0A2J8A3Z7
Edaphochlamys_A0A835XNV1
Emiliania_R1EF93
Haptolina_A0A7S3B471
Trypanosoma_A0A1X0P2F4
Chlorella_A0A9D4YWD1
Taxa
a Cluster representative
a Archaea
a Sar
a Eukaryota
a Viridiplantae
a Metazoa
a Sciaridae

**Tree name : fol_can_coll_XP_035707195.1_treefile**
**Protein length : 375**
**Alignment length : 223**
**Aligned AA count : 223**

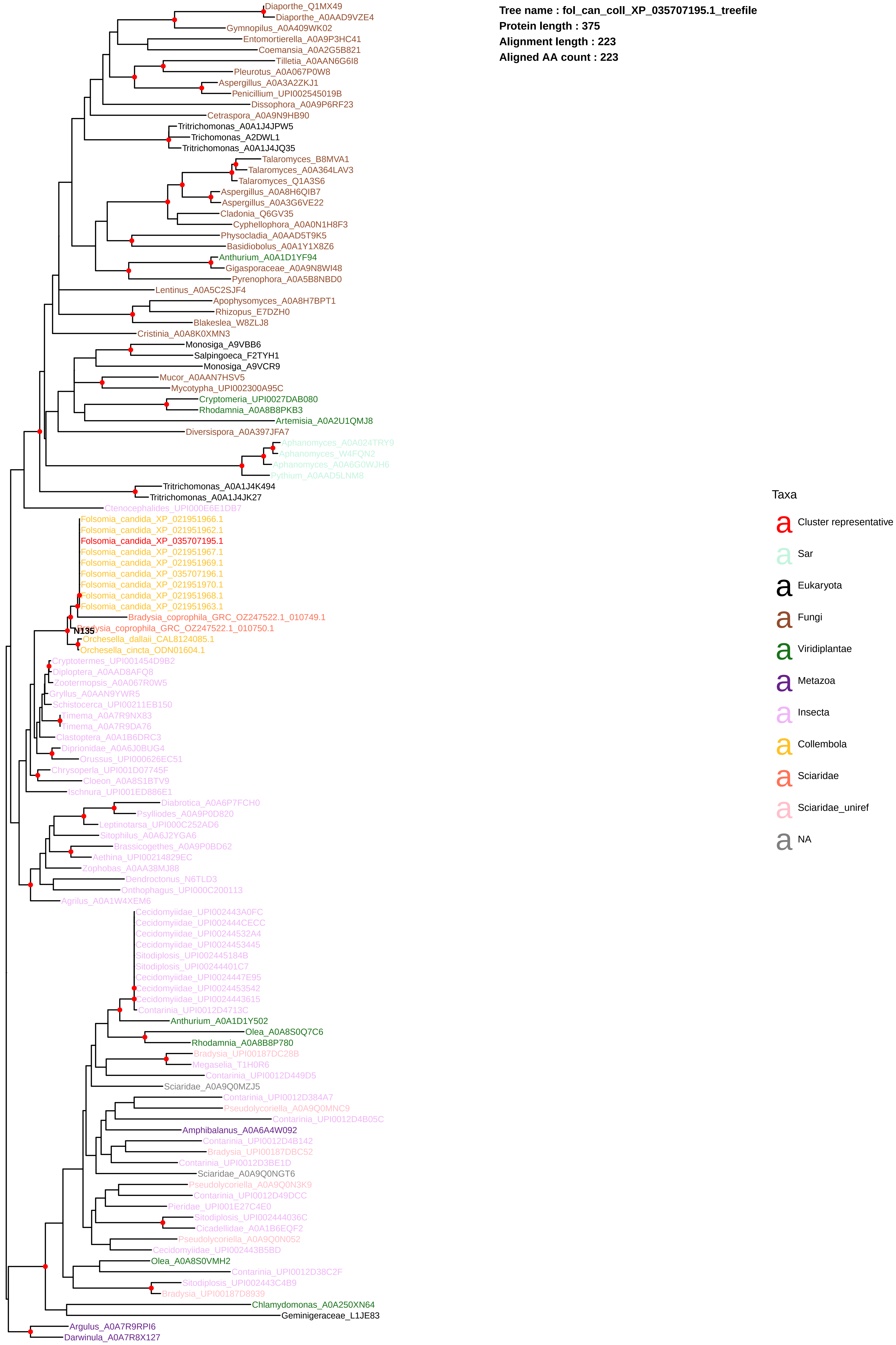

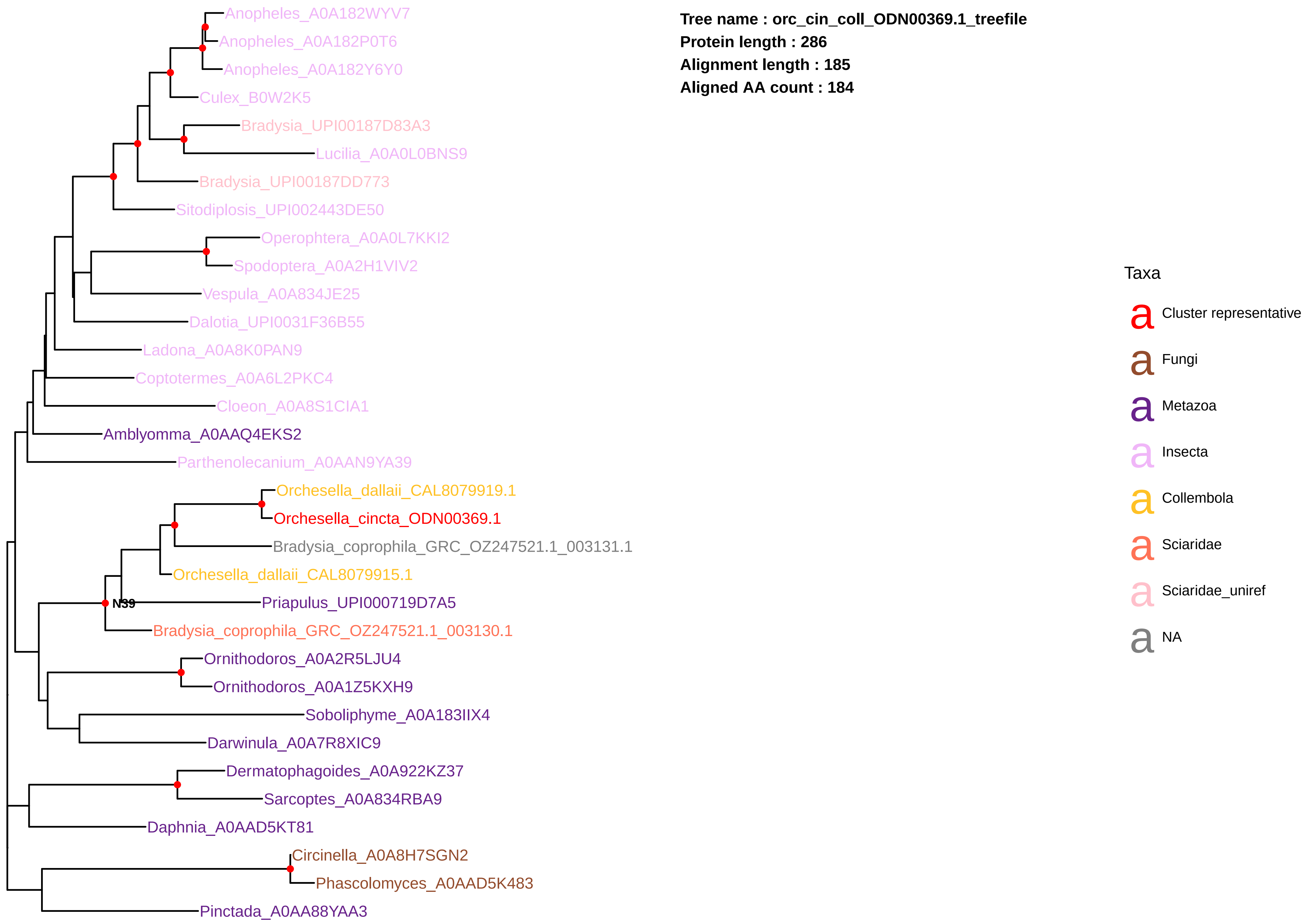

Tree name : orc_cin_coll_ODN00369.1_treefile
Protein length : 286
Alignment length : 185
Aligned AA count : 184
Anopheles_A0A182WYV7
Anopheles_A0A182P0T6
Anopheles_A0A182Y6Y0
Culex_B0W2K5
Bradysia_UPI00187D83A3
Lucilia_A0A0L0BNS9
Bradysia_UPI00187DD773
Sitodiplosis_UPI002443DE50
Operophtera_A0A0L7KKI2
Spodoptera_A0A2H1VIV2
Vespula_A0A834JE25
Dalotia_UPI0031F36B55
Ladona_A0A8K0PAN9
Coptotermes_A0A6L2PKC4
Cloeon_A0A8S1CIA1
Amblyomma_A0AAQ4EKS2
Parthenolecanium_A0AAN9YA39
Orchesella_dallaii_CAL8079919.1
Orchesella_cincta_ODN00369.1
Bradysia_coprophila_GRC_OZ247521.1_003131.1
Orchesella_dallaii_CAL8079915.1
N39
Priapulus_UPI000719D7A5
Bradysia_coprophila_GRC_OZ247521.1_003130.1
Ornithodoros_A0A2R5LJU4
Ornithodoros_A0A1Z5KXH9
Soboliphyme_A0A183IIX4
Darwinula_A0A7R8XIC9
Dermatophagoides_A0A922KZ37
Sarcoptes_A0A834RBA9
Daphnia_A0AAD5KT81
Circinella_A0A8H7SGN2
Phascolomyces_A0AAD5K483
Pinctada_A0AA88YAA3
Taxa
Cluster representative
Fungi
Metazoa
Insecta
Collembola
Sciaridae
Sciaridae_uniref
NA

**Tree name : orc_cin_coll_ODN01798.1_treefile**
**Protein length : 418**
**Alignment length : 204**
**Aligned AA count : 204**

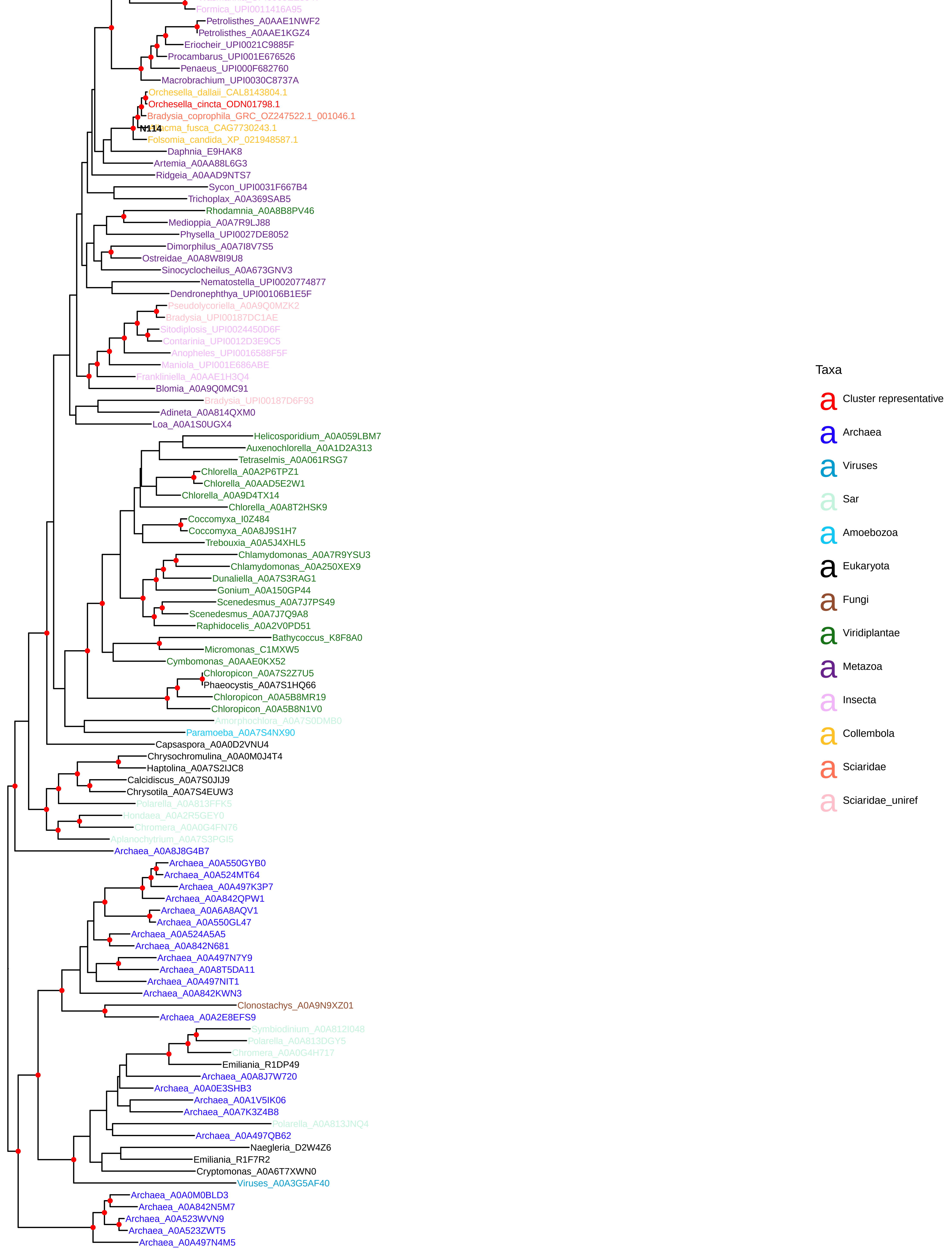

**Tree name : orc_dal_coll_CAL8071491.1_treefile**
**Protein length : 1319**
**Alignment length : 281**
**Aligned AA count : 281**

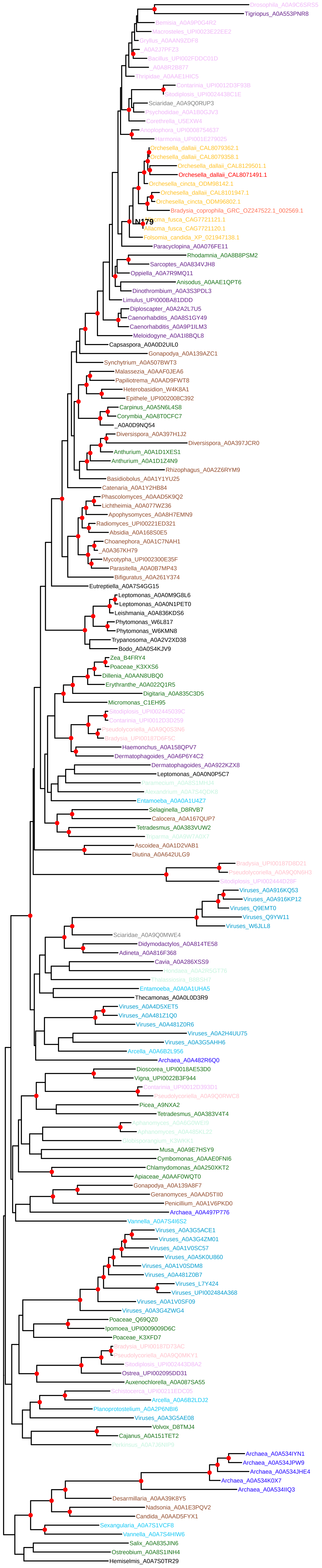


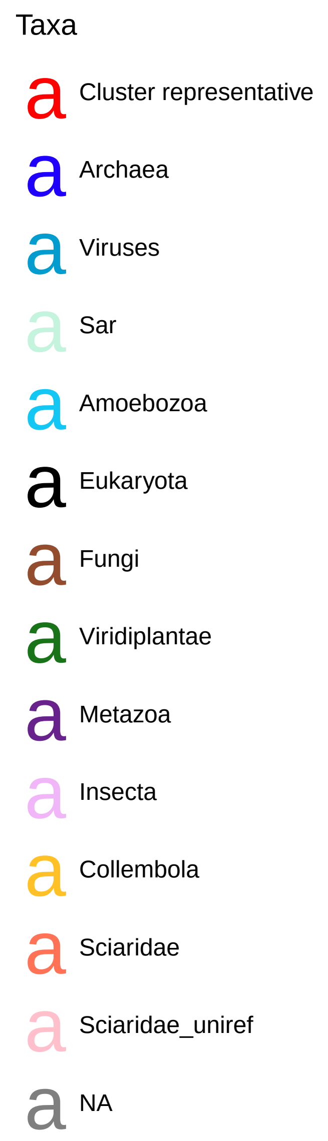

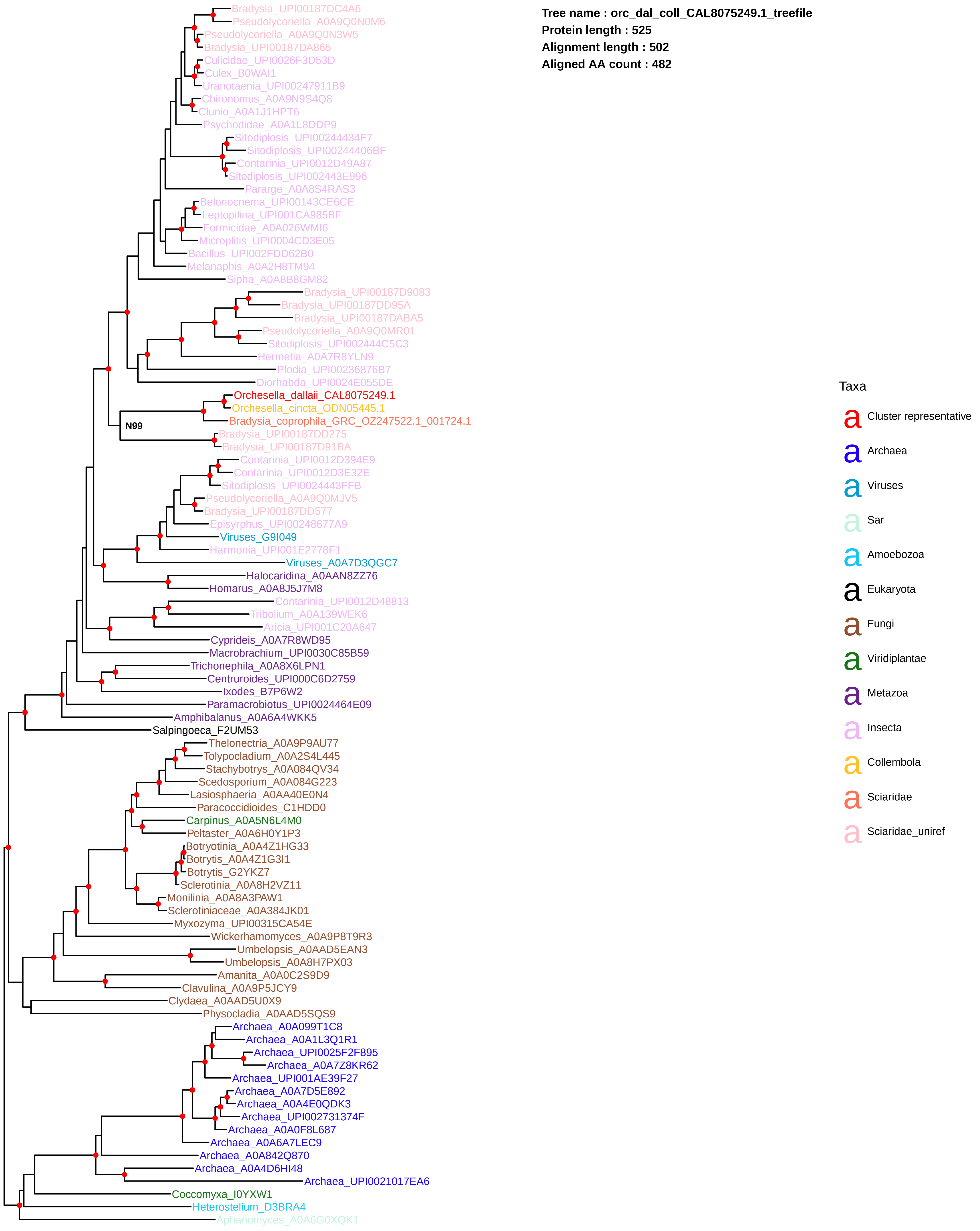

Tree name : orc_dal_coll_CAL8075249.1_treefile
Protein length : 525
Alignment length : 502
Aligned AA count : 482
Bradysia_UPI00187DC4A6
Pseudolycoriella_A0A9Q0N0M6
Pseudolycoriella_A0A9Q0N3W5
Bradysia_UPI00187DA865
Culicidae_UPI0026F3D53D
Culex_B0WAI1
Uranotaenia_UPI00247911B9
Chironomus_A0A9N9S4Q8
Clunio_A0A1J1HPT6
Psychodidae_A0A1L8DDP9
Sitodiplosis_UPI00244434F7
Sitodiplosis_UPI00244406BF
Contarinia_UPI0012D49A87
Sitodiplosis_UPI002443E996
Pararge_A0A8S4RAS3
Belonocnema_UPI00143CE6CE
Leptopilina_UPI001CA985BF
Formicidae_A0A026WMI6
Microplitis_UPI0004CD3E05
Bacillus_UPI002FDD62B0
Melanaphis_A0A2H8TM94
Sipha_A0A8B8GM82
Bradysia_UPI00187D9083
Bradysia_UPI00187DD95A
Bradysia_UPI00187DABA5
Pseudolycoriella_A0A9Q0MR01
Sitodiplosis_UPI002444C5C3
Hermetia_A0A7R8YLN9
Plodia_UPI00236876B7
Diorhabda_UPI0024E055DE
Orchesella_dallaii_CAL8075249.1
Orchesella_cincta_ODN05445.1
N99
Bradysia_coprophila_GRC_OZ247522.1_001724.1
Bradysia_UPI00187DD275
Bradysia_UPI00187D91BA
Contarinia_UPI0012D394E9
Contarinia_UPI0012D3E32E
Sitodiplosis_UPI0024443FFB
Pseudolycoriella_A0A9Q0MJV5
Bradysia_UPI00187DD577
Episyrphus_UPI00248677A9
Viruses_G9I049
Harmonia_UPI001E2778F1
Viruses_A0A7D3QGC7
Halocaridina_A0AAN8ZZ76
Homarus_A0A8J5J7M8
Contarinia_UPI0012D48813
Tribolium_A0A139WEK6
Aricia_UPI001C20A647
Cyprideis_A0A7R8WD95
Macrobrachium_UPI0030C85B59
Trichonephila_A0A8X6LPN1
Centruroides_UPI000C6D2759
Ixodes_B7P6W2
Paramacrobiotus_UPI0024464E09
Amphibalanus_A0A6A4WKK5
Salpingoeca_F2UM53
Thelonectria_A0A9P9AU77
Tolypocladium_A0A2S4L445
Stachybotrys_A0A084QV34
Scedosporium_A0A084G223
Lasiosphaeria_A0AA40E0N4
Paracoccidioides_C1HDD0
Carpinus_A0A5N6L4M0
Peltaster_A0A6H0Y1P3
Botryotinia_A0A4Z1HG33
Botrytis_A0A4Z1G3I1
Botrytis_G2YKZ7
Sclerotinia_A0A8H2VZ11
Monilinia_A0A8A3PAW1
Sclerotiniaceae_A0A384JK01
Myxozyma_UPI00315CA54E
Wickerhamomyces_A0A9P8T9R3
Umbelopsis_A0AAD5EAN3
Umbelopsis_A0A8H7PX03
Amanita_A0A0C2S9D9
Clavulina_A0A9P5JCY9
Clydaea_A0AAD5U0X9
Physocladia_A0AAD5SQS9
Archaea_A0A099T1C8
Archaea_A0A1L3Q1R1
Archaea_UPI0025F2F895
Archaea_A0A7Z8KR62
Archaea_UPI001AE39F27
Archaea_A0A7D5E892
Archaea_A0A4E0QDK3
Archaea_UPI002731374F
Archaea_A0A0F8L687
Archaea_A0A6A7LEC9
Archaea_A0A842Q870
Archaea_A0A4D6HI48
Archaea_UPI0021017EA6
Coccomyxa_I0YXW1
Heterostelium_D3BRA4
Aphanomyces_A0A6G0XQK1
Taxa
Cluster representative
Archaea
Viruses
Sar
Amoebozoa
Eukaryota
Fungi
Viridiplantae
Metazoa
Insecta
Collembola
Sciaridae
Sciaridae_uniref

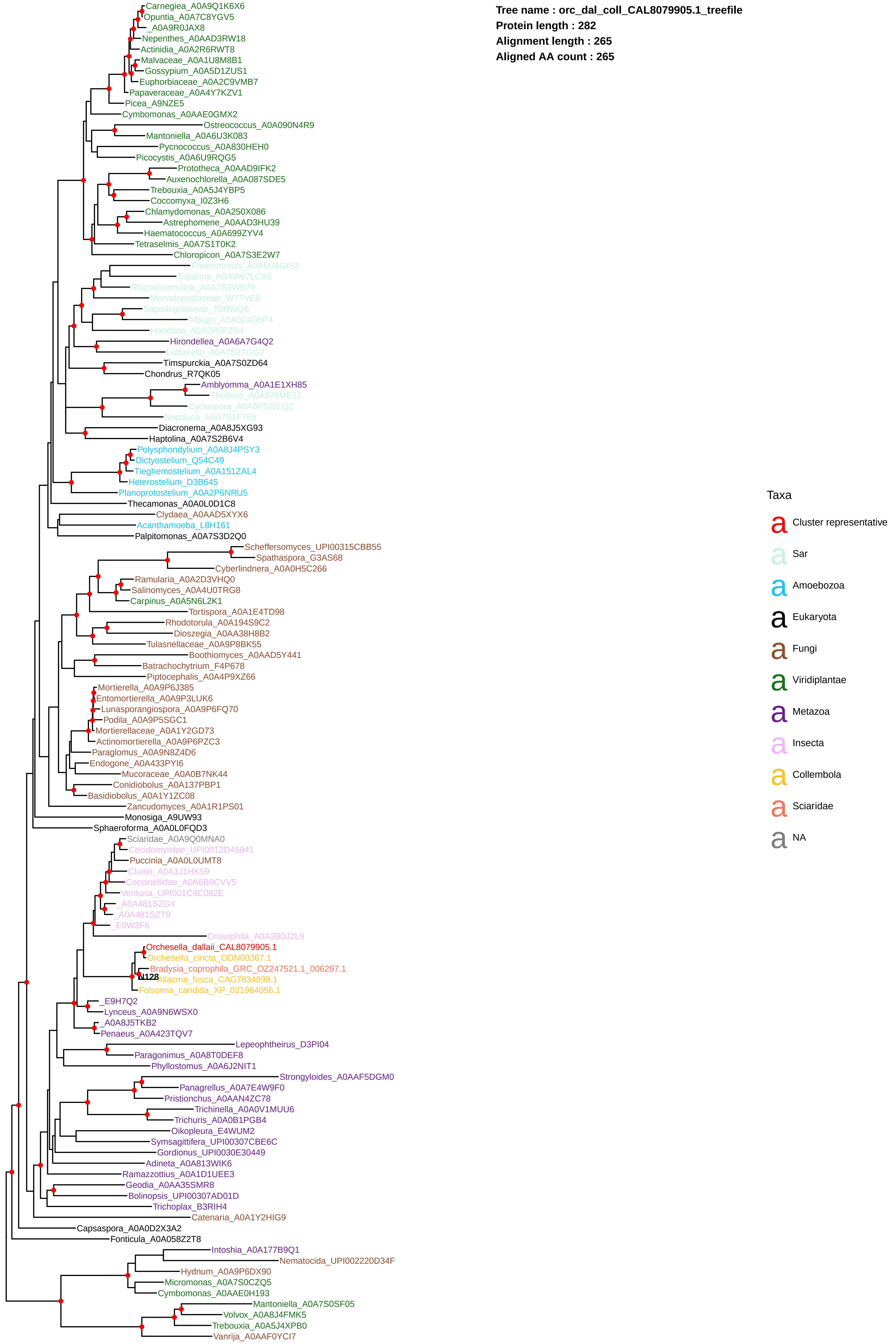

Tree name : orc_dal_coll_CAL8079905.1_treefile
Protein length : 282
Alignment length : 265
Aligned AA count : 265
Carnegiea_A0A9Q1K6X6
Opuntia_A0A7C8YGV5
_A0A9R0JAX8
Nepenthes_A0AAD3RW18
Actinidia_A0A2R6RWT8
Malvaceae_A0A1U8M8B1
Gossypium_A0A5D1ZUS1
Euphorbiaceae_A0A2C9VMB7
Papaveraceae_A0A4Y7KZV1
Picea_A9NZE5
Cymbomonas_A0AAE0GMX2
Ostreococcus_A0A090N4R9
Mantoniella_A0A6U3K083
Pycnococcus_A0A830HEH0
Picocystis_A0A6U9RQG5
Prototheca_A0AAD9IFK2
Auxenochlorella_A0A087SDE5
Trebouxia_A0A5J4YBP5
Coccomyxa_I0Z3H6
Chlamydomonas_A0A250X086
Astrephomene_A0AAD3HU39
Haematococcus_A0A699ZYV4
Tetraselmis_A0A7S1T0K2
Chloropicon_A0A7S3E2W7
Phaeomonas_A0A6U4GIS3
Triparma_A0A9W7LCX6
Rhizochromulina_A0A7S2WB79
Monodopsidaceae_W7TVE8
Saprolegniaceae_T0RWQ6
Albugo_A0A024G6P4
Hondaea_A0A2R5FZB4
Hirondellea_A0A6A7G4Q2
Lotharella_A0A7S2TGG7
Timspurckia_A0A7S0ZD64
Chondrus_R7QK05
Amblyomma_A0A1E1XH85
Theileria_A0A976ME11
Cyclospora_A0A6P5WEQ2
Noctiluca_A0A7S1F769
Diacronema_A0A8J5XG93
Haptolina_A0A7S2B6V4
Polysphondylium_A0A8J4PSY3
Dictyostelium_Q54C49
Tieghemostelium_A0A151ZAL4
Heterostelium_D3B645
Planoprotostelium_A0A2P6NRU5
Thecamonas_A0A0L0D1C8
Clydaea_A0AAD5XYX6
Acanthamoeba_L8H161
Palpitomonas_A0A7S3D2Q0
Scheffersomyces_UPI00315CBB55
Spathaspora_G3AS68
Cyberlindnera_A0A0H5C266
Ramularia_A0A2D3VHQ0
Salinomyces_A0A4U0TRG8
Carpinus_A0A5N6L2K1
Tortispora_A0A1E4TD98
Rhodotorula_A0A194S9C2
Dioszegia_A0AA38H8B2
Tulasnellaceae_A0A9P8BK55
Boothiomyces_A0AAD5Y441
Batrachochytrium_F4P678
Piptocephalis_A0A4P9XZ66
Mortierella_A0A9P6J385
Entomortierella_A0A9P3LUK6
Lunasporangiospora_A0A9P6FQ70
Podila_A0A9P5SGC1
Mortierellaceae_A0A1Y2GD73
Actinomortierella_A0A9P6PZC3
Paraglomus_A0A9N8Z4D6
Endogone_A0A433PYI6
Mucoraceae_A0A0B7NK44
Conidiobolus_A0A137PBP1
Basidiobolus_A0A1Y1ZC08
Zancudomyces_A0A1R1PS01
Monosiga_A9UW93
Sphaeroforma_A0A0L0FQD3
Sciaridae_A0A9Q0MNA0
Cecidomyiidae_UPI0012D45841
Puccinia_A0A0L0UMT8
Clunio_A0A1J1HK59
Coccinellidae_A0A6B9CVV5
Venturia_UPI001C9C082E
_A0A481SZG4
_A0A481SZT9
_E0W3F6
Drosophila_A0A3B0J2L9
Orchesella_dallaii_CAL8079905.1
Orchesella_cincta_ODN00367.1
Bradysia_coprophila_GRC_OZ247521.1_006297.1
N128
Allacma_fusca_CAG7834898.1
Folsomia_candida_XP_021964056.1
_E9H7Q2
Lynceus_A0A9N6WSX0
_A0A8J5TKB2
Penaeus_A0A423TQV7
Lepeophtheirus_D3PI04
Paragonimus_A0A8T0DEF8
Phyllostomus_A0A6J2NIT1
Strongyloides_A0AAF5DGM0
Panagrellus_A0A7E4W9F0
Pristionchus_A0AAN4ZC78
Trichinella_A0A0V1MUU6
Trichuris_A0A0B1PGB4
Oikopleura_E4WUM2
Symsagittifera_UPI00307CBE6C
Gordionus_UPI0030E30449
Adineta_A0A813WIK6
Ramazzottius_A0A1D1UEE3
Geodia_A0AA35SMR8
Bolinopsis_UPI00307AD01D
Trichoplax_B3RIH4
Catenaria_A0A1Y2HIG9
Capsaspora_A0A0D2X3A2
Fonticula_A0A058Z2T8
Intoshia_A0A177B9Q1
Nematocida_UPI002220D34F
Hydnum_A0A9P6DX90
Micromonas_A0A7S0CZQ5
Cymbomonas_A0AAE0H193
Mantoniella_A0A7S0SF05
Volvox_A0A8J4FMK5
Trebouxia_A0A5J4XPB0
Vanrija_A0AAF0YCI7
Taxa
Cluster representative
Sar
Amoebozoa
Eukaryota
Fungi
Viridiplantae
Metazoa
Insecta
Collembola
Sciaridae
NA

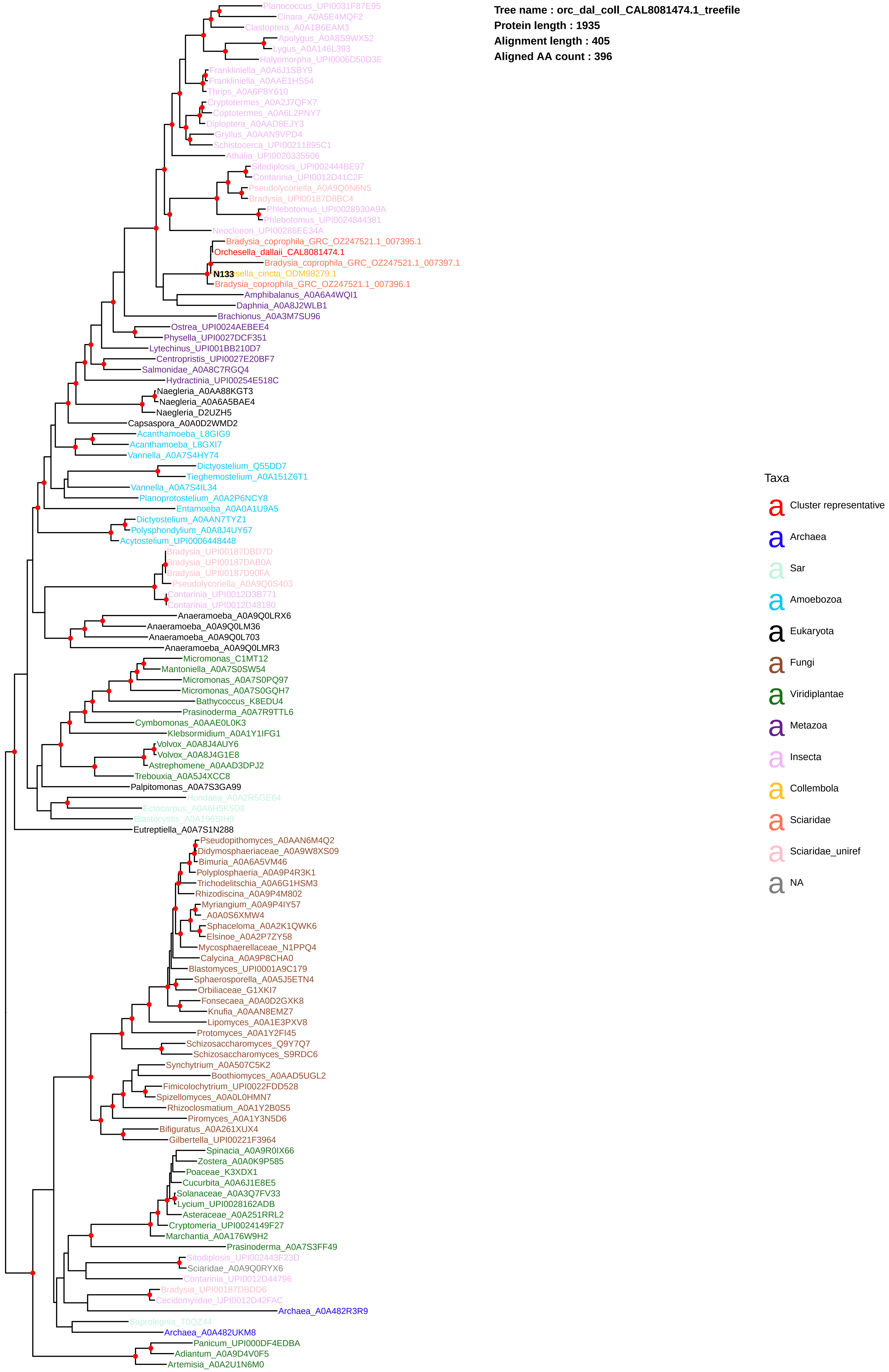

Tree name : orc_dal_coll_CAL8081474.1_treefile
Protein length : 1935
Alignment length : 405
Aligned AA count : 396
Planococcus_UPI0031F87E95
Cinara_A0A5E4MQF2
Clastoptera_A0A1B6EAM3
Apolygus_A0A8S9WX52
Lygus_A0A146L393
Halyomorpha_UPI0006D50D3E
Frankliniella_A0A6J1SBY9
Frankliniella_A0AAE1HS54
Thrips_A0A6P8Y610
Cryptotermes_A0A2J7QFX7
Coptotermes_A0A6L2PNY7
Diploptera_A0AAD8EJY3
Gryllus_A0AAN9VPD4
Schistocerca_UPI00211895C1
Athalia_UPI0020335506
Sitodiplosis_UPI002444BE97
Contarinia_UPI0012D41C2F
Pseudolycoriella_A0A9Q0N6N5
Bradysia_UPI00187D8BC4
Phlebotomus_UPI0028930A9A
Phlebotomus_UPI0024844381
Neocloeon_UPI00286EE34A
Bradysia_coprophila_GRC_OZ247521.1_007395.1
Orchesella_dallaii_CAL8081474.1
Bradysia_coprophila_GRC_OZ247521.1_007397.1
N133
sella_cincta_ODM98279.1
Bradysia_coprophila_GRC_OZ247521.1_007396.1
Amphibalanus_A0A6A4WQI1
Daphnia_A0A8J2WLB1
Brachionus_A0A3M7SU96
Ostrea_UPI0024AEBEE4
Physella_UPI0027DCF351
Lytechinus_UPI001BB210D7
Centropristis_UPI0027E20BF7
Salmonidae_A0A8C7RGQ4
Hydractinia_UPI00254E518C
Naegleria_A0AA88KGT3
Naegleria_A0A6A5BAE4
Naegleria_D2UZH5
Capsaspora_A0A0D2WMD2
Acanthamoeba_L8GIG9
Acanthamoeba_L8GXI7
Vannella_A0A7S4HY74
Dictyostelium_Q55DD7
Tieghemostelium_A0A151Z6T1
Vannella_A0A7S4IL34
Planoprotostelium_A0A2P6NCY8
Entamoeba_A0A0A1U9A5
Dictyostelium_A0AAN7TYZ1
Polysphondylium_A0A8J4UY67
Acytostelium_UPI0006448448
Bradysia_UPI00187DBD7D
Bradysia_UPI00187DAB0A
Bradysia_UPI00187D90FA
Pseudolycoriella_A0A9Q0S403
Contarinia_UPI0012D3B771
Contarinia_UPI0012D48180
Anaeramoeba_A0A9Q0LRX6
Anaeramoeba_A0A9Q0LM36
Anaeramoeba_A0A9Q0L703
Anaeramoeba_A0A9Q0LMR3
Micromonas_C1MT12
Mantoniella_A0A7S0SW54
Micromonas_A0A7S0PQ97
Micromonas_A0A7S0GQH7
Bathycoccus_K8EDU4
Prasinoderma_A0A7R9TTL6
Cymbomonas_A0AAE0L0K3
Klebsormidium_A0A1Y1IFG1
Volvox_A0A8J4AUY6
Volvox_A0A8J4G1E8
Astrephomene_A0AAD3DPJ2
Trebouxia_A0A5J4XCC8
Palpitomonas_A0A7S3GA99
Hondaea_A0A2R5GE64
Ectocarpus_A0A6H5K5D8
Blastocystis_A0A196SIH9
Eutreptiella_A0A7S1N288
Pseudopithomyces_A0AAN6M4Q2
Didymosphaeriaceae_A0A9W8XS09
Bimuria_A0A6A5VM46
Polyplosphaeria_A0A9P4R3K1
Trichodelitschia_A0A6G1HSM3
Rhizodiscina_A0A9P4M802
Myriangium_A0A9P4IY57
_A0A0S6XMW4
Sphaceloma_A0A2K1QWK6
Elsinoe_A0A2P7ZY58
Mycosphaerellaceae_N1PPQ4
Calycina_A0A9P8CHA0
Blastomyces_UPI0001A9C179
Sphaerosporella_A0A5J5ETN4
Orbiliaceae_G1XKI7
Fonsecaea_A0A0D2GXK8
Knufia_A0AAN8EMZ7
Lipomyces_A0A1E3PXV8
Protomyces_A0A1Y2FI45
Schizosaccharomyces_Q9Y7Q7
Schizosaccharomyces_S9RDC6
Synchytrium_A0A507C5K2
Boothiomyces_A0AAD5UGL2
Fimicolochytrium_UPI0022FDD528
Spizellomyces_A0A0L0HMN7
Rhizoclosmatium_A0A1Y2B0S5
Piromyces_A0A1Y3N5D6
Bifiguratus_A0A261XUX4
Gilbertella_UPI00221F3964
Spinacia_A0A9R0IX66
Zostera_A0A0K9P585
Poaceae_K3XDX1
Cucurbita_A0A6J1E8E5
Solanaceae_A0A3Q7FV33
Lycium_UPI0028162ADB
Asteraceae_A0A251RRL2
Cryptomeria_UPI0024149F27
Marchantia_A0A176W9H2
Prasinoderma_A0A7S3FF49
Sitodiplosis_UPI002443F23D
Sciaridae_A0A9Q0RYX6
Contarinia_UPI0012D44796
Bradysia_UPI00187DBDD6
Cecidomyiidae_UPI0012D42FAC
Archaea_A0A482R3R9
Saprolegnia_T0QZ44
Archaea_A0A482UKM8
Panicum_UPI000DF4EDBA
Adiantum_A0A9D4V0F5
Artemisia_A0A2U1N6M0
Taxa
Cluster representative
Archaea
Sar
Amoebozoa
Eukaryota
Fungi
Viridiplantae
Metazoa
Insecta
Collembola
Sciaridae
Sciaridae_uniref
NA

Tree name : orc_dal_coll_CAL8087782.1_treefile
Protein length : 1438
Alignment length : 1128
Aligned AA count : 1074

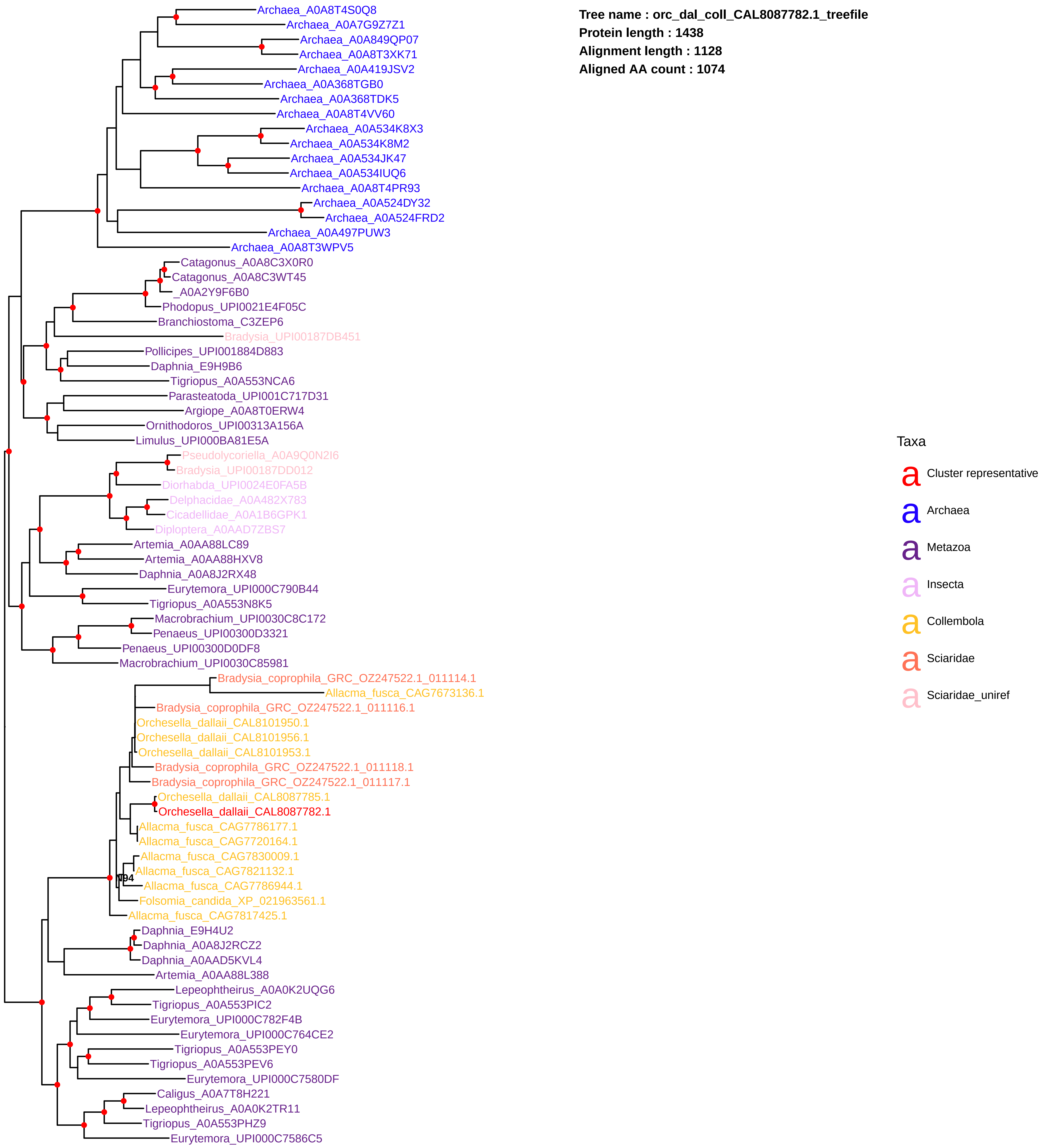

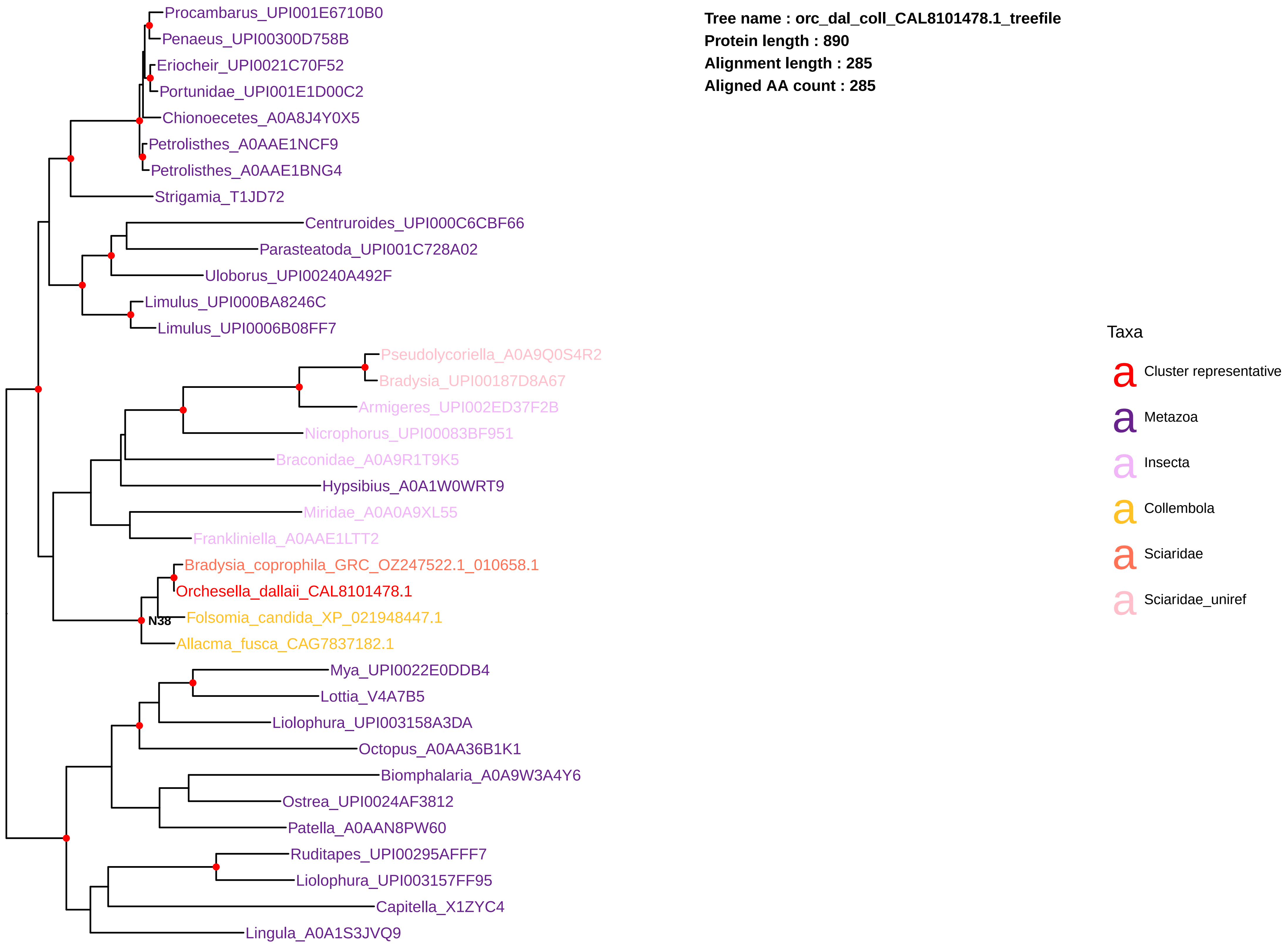
Tree name : orc_dal_coll_CAL8101478.1_treefile
Protein length : 890
Alignment length : 285
Aligned AA count : 285
Procambarus_UPI001E6710B0
Penaeus_UPI00300D758B
Eriocheir_UPI0021C70F52
Portunidae_UPI001E1D00C2
Chionoecetes_A0A8J4Y0X5
Petrolisthes_A0AAE1NCF9
Petrolisthes_A0AAE1BNG4
Strigamia_T1JD72
Centruroides_UPI000C6CBF66
Parasteatoda_UPI001C728A02
Uloborus_UPI00240A492F
Limulus_UPI000BA8246C
Limulus_UPI0006B08FF7
Pseudolycoriella_A0A9Q0S4R2
Bradysia_UPI00187D8A67
Armigeres_UPI002ED37F2B
Nicrophorus_UPI00083BF951
Braconidae_A0A9R1T9K5
Hypsibius_A0A1W0WRT9
Miridae_A0A0A9XL55
Frankliniella_A0AAE1LTT2
Bradysia_coprophila_GRC_OZ247522.1_010658.1
Orchesella_dallaii_CAL8101478.1
N38
Folsomia_candida_XP_021948447.1
Allacma_fusca_CAG7837182.1
Mya_UPI0022E0DDB4
Lottia_V4A7B5
Liolophura_UPI003158A3DA
Octopus_A0AA36B1K1
Biomphalaria_A0A9W3A4Y6
Ostrea_UPI0024AF3812
Patella_A0AAN8PW60
Ruditapes_UPI00295AFFF7
Liolophura_UPI003157FF95
Capitella_X1ZYC4
Lingula_A0A1S3JVQ9
Taxa
Cluster representative
Metazoa
Insecta
Collembola
Sciaridae
Sciaridae_uniref

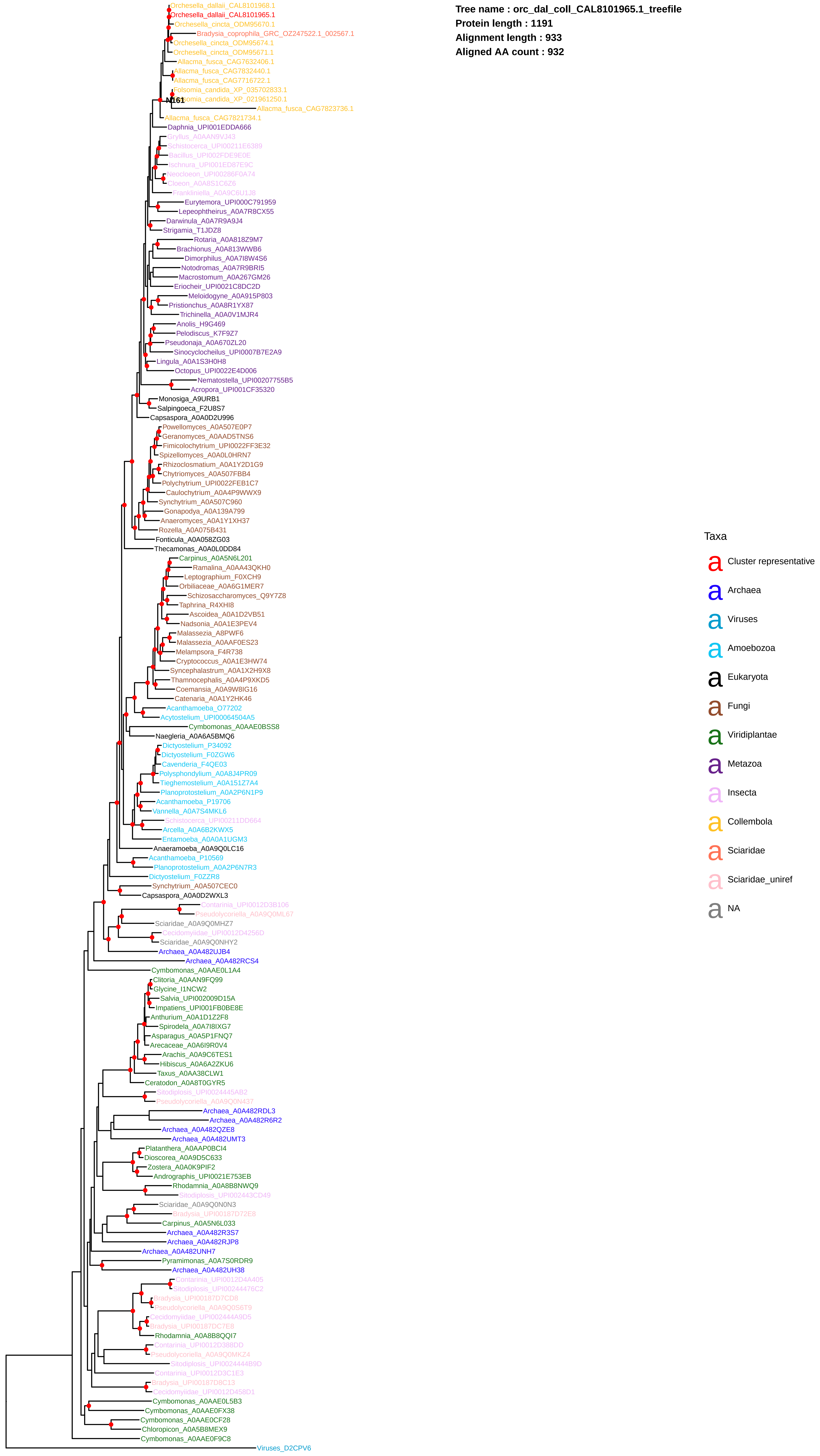

Tree name : orc_dal_coll_CAL8101965.1_treefile
Protein length : 1191
Alignment length : 933
Aligned AA count : 932
Orchesella_dallaii_CAL8101968.1
Orchesella_dallaii_CAL8101965.1
Orchesella_cincta_ODM95670.1
Bradysia_coprophila_GRC_OZ247522.1_002567.1
Orchesella_cincta_ODM95674.1
Orchesella_cincta_ODM95671.1
Allacma_fusca_CAG7632406.1
Allacma_fusca_CAG7832440.1
Allacma_fusca_CAG7716722.1
Folsomia_candida_XP_035702833.1
N161
omia_candida_XP_021961250.1
Allacma_fusca_CAG7823736.1
Allacma_fusca_CAG7821734.1
Daphnia_UPI001EDDA666
Gryllus_A0AAN9VJ43
Schistocerca_UPI00211E6389
Bacillus_UPI002FDE9E0E
Ischnura_UPI001ED87E9C
Neocloeon_UPI00286F0A74
Cloeon_A0A8S1C6Z6
Frankliniella_A0A9C6U1J8
Eurytemora_UPI000C791959
Lepeophtheirus_A0A7R8CX55
Darwinula_A0A7R9A9J4
Strigamia_T1JDZ8
Rotaria_A0A818Z9M7
Brachionus_A0A813WWB6
Dimorphilus_A0A7I8W4S6
Notodromas_A0A7R9BRI5
Macrostomum_A0A267GM26
Eriocheir_UPI0021C8DC2D
Meloidogyne_A0A915P803
Pristionchus_A0A8R1YX87
Trichinella_A0A0V1MJR4
Anolis_H9G469
Pelodiscus_K7F9Z7
Pseudonaja_A0A670ZL20
Sinocyclocheilus_UPI0007B7E2A9
Lingula_A0A1S3H0H8
Octopus_UPI0022E4D006
Nematostella_UPI00207755B5
Acropora_UPI001CF35320
Monosiga_A9URB1
Salpingoeca_F2U8S7
Capsaspora_A0A0D2U996
Powellomyces_A0A507E0P7
Geranomyces_A0AAD5TNS6
Fimicolochytrium_UPI0022FF3E32
Spizellomyces_A0A0L0HRN7
Rhizoclosmatium_A0A1Y2D1G9
Chytriomyces_A0A507FBB4
Polychytrium_UPI0022FEB1C7
Caulochytrium_A0A4P9WWX9
Synchytrium_A0A507C960
Gonapodya_A0A139A799
Anaeromyces_A0A1Y1XH37
Rozella_A0A075B431
Fonticula_A0A058ZG03
Thecamonas_A0A0L0DD84
Carpinus_A0A5N6L201
Ramalina_A0AA43QKH0
Leptographium_F0XCH9
Orbiliaceae_A0A6G1MER7
Schizosaccharomyces_Q9Y7Z8
Taphrina_R4XHI8
Ascoidea_A0A1D2VB51
Nadsonia_A0A1E3PEV4
Malassezia_A8PWF6
Malassezia_A0AAF0ES23
Melampsora_F4R738
Cryptococcus_A0A1E3HW74
Syncephalastrum_A0A1X2H9X8
Thamnocephalis_A0A4P9XKD5
Coemansia_A0A9W8IG16
Catenaria_A0A1Y2HK46
Acanthamoeba_O77202
Acytostelium_UPI00064504A5
Cymbomonas_A0AAE0BSS8
Naegleria_A0A6A5BMQ6
Dictyostelium_P34092
Dictyostelium_F0ZGW6
Cavenderia_F4QE03
Polysphondylium_A0A8J4PR09
Tieghemostelium_A0A151Z7A4
Planoprotostelium_A0A2P6N1P9
Acanthamoeba_P19706
Vannella_A0A7S4MKL6
Schistocerca_UPI00211DD664
Arcella_A0A6B2KWX5
Entamoeba_A0A0A1UGM3
Anaeramoeba_A0A9Q0LC16
Acanthamoeba_P10569
Planoprotostelium_A0A2P6N7R3
Dictyostelium_F0ZZR8
Synchytrium_A0A507CEC0
Capsaspora_A0A0D2WXL3
Contarinia_UPI0012D3B106
Pseudolycoriella_A0A9Q0ML67
Sciaridae_A0A9Q0MHZ7
Cecidomyiidae_UPI0012D4256D
Sciaridae_A0A9Q0NHY2
Archaea_A0A482UJB4
Archaea_A0A482RCS4
Cymbomonas_A0AAE0L1A4
Clitoria_A0AAN9FQ99
Glycine_I1NCW2
Salvia_UPI002009D15A
Impatiens_UPI001FB0BE8E
Anthurium_A0A1D1Z2F8
Spirodela_A0A7I8IXG7
Asparagus_A0A5P1FNQ7
Arecaceae_A0A6I9R0V4
Arachis_A0A9C6TES1
Hibiscus_A0A6A2ZKU6
Taxus_A0AA38CLW1
Ceratodon_A0A8T0GYR5
Sitodiplosis_UPI0024445AB2
Pseudolycoriella_A0A9Q0N437
Archaea_A0A482RDL3
Archaea_A0A482R6R2
Archaea_A0A482QZE8
Archaea_A0A482UMT3
Platanthera_A0AAP0BCI4
Dioscorea_A0A9D5C633
Zostera_A0A0K9PIF2
Andrographis_UPI0021E753EB
Rhodamnia_A0A8B8NWQ9
Sitodiplosis_UPI002443CD49
Sciaridae_A0A9Q0N0N3
Bradysia_UPI00187D72E8
Carpinus_A0A5N6L033
Archaea_A0A482R3S7
Archaea_A0A482RJP8
Archaea_A0A482UNH7
Pyramimonas_A0A7S0RDR9
Archaea_A0A482UH38
Contarinia_UPI0012D4A405
Sitodiplosis_UPI00244476C2
Bradysia_UPI00187D7CD8
Pseudolycoriella_A0A9Q0S6T9
Cecidomyiidae_UPI002444A9D5
Bradysia_UPI00187DC7E8
Rhodamnia_A0A8B8QQI7
Contarinia_UPI0012D388DD
Pseudolycoriella_A0A9Q0MKZ4
Sitodiplosis_UPI0024444B9D
Contarinia_UPI0012D3C1E3
Bradysia_UPI00187D8C13
Cecidomyiidae_UPI0012D458D1
Cymbomonas_A0AAE0L5B3
Cymbomonas_A0AAE0FX38
Cymbomonas_A0AAE0CF28
Chloropicon_A0A5B8MEX9
Cymbomonas_A0AAE0F9C8
Viruses_D2CPV6
Taxa
Cluster representative
Archaea
Viruses
Amoebozoa
Eukaryota
Fungi
Viridiplantae
Metazoa
Insecta
Collembola
Sciaridae
Sciaridae_uniref
NA

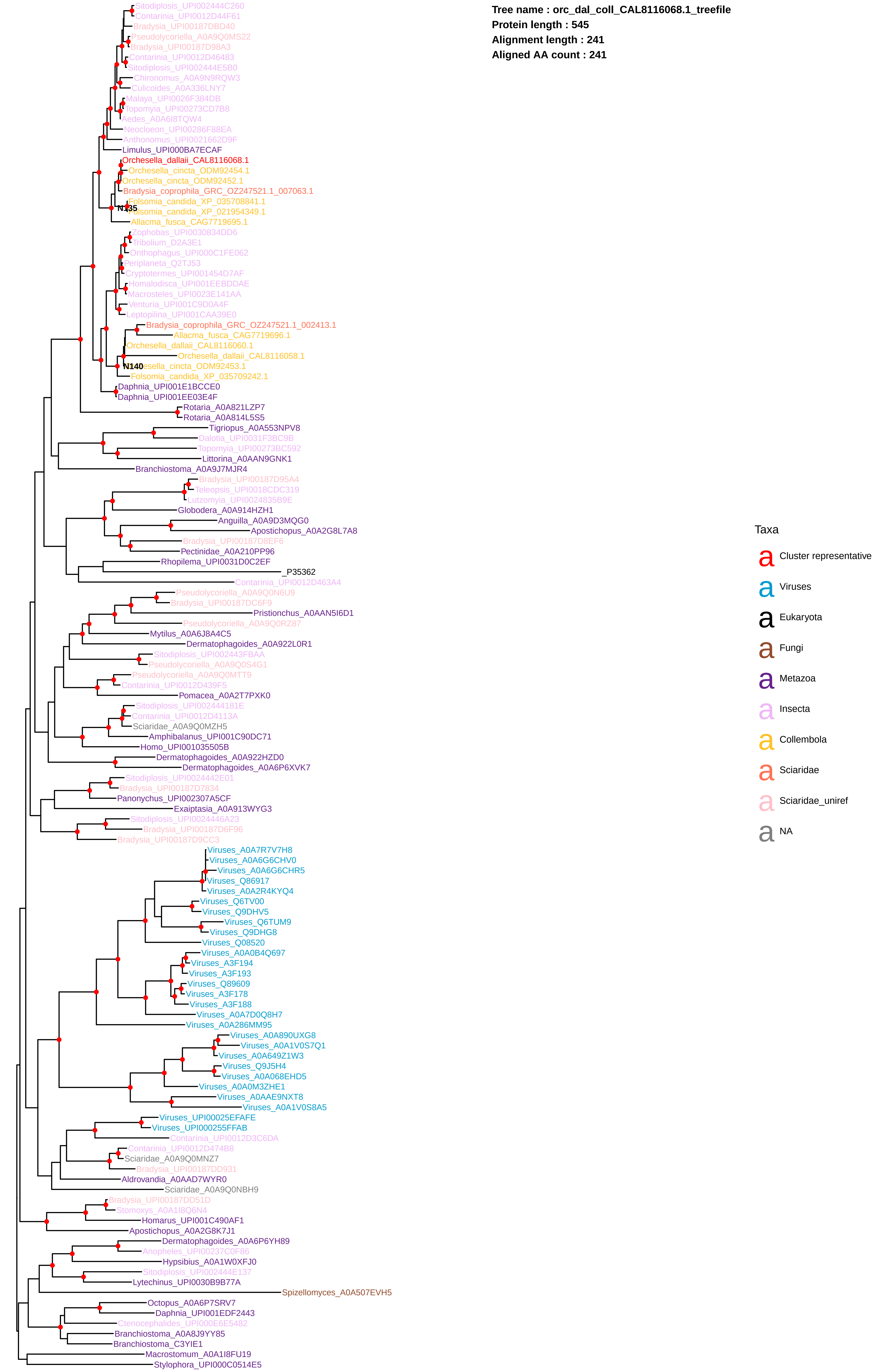
Tree name : orc_dal_coll_CAL8116068.1_treefile
Protein length : 545
Alignment length : 241
Aligned AA count : 241
Sitodiplosis_UPI002444C260
Contarinia_UPI0012D44F61
Bradysia_UPI00187DBD40
Pseudolycoriella_A0A9Q0MS22
Bradysia_UPI00187D98A3
Contarinia_UPI0012D46483
Sitodiplosis_UPI002444E5B0
Chironomus_A0A9N9RQW3
Culicoides_A0A336LNY7
Malaya_UPI0026F384DB
Topomyia_UPI00273CD7B8
Aedes_A0A6I8TQW4
Neocloeon_UPI00286F88EA
Anthonomus_UPI0021662D9F
Limulus_UPI000BA7ECAF
Orchesella_dallaii_CAL8116068.1
Orchesella_cincta_ODM92454.1
Orchesella_cincta_ODM92452.1
Bradysia_coprophila_GRC_OZ247521.1_007063.1
Folsomia_candida_XP_035708841.1
N135
Folsomia_candida_XP_021954349.1
Allacma_fusca_CAG7719695.1
Zophobas_UPI0030834DD6
Tribolium_D2A3E1
Onthophagus_UPI000C1FE062
Periplaneta_Q2TJ53
Cryptotermes_UPI001454D7AF
Homalodisca_UPI001EEBDDAE
Macrosteles_UPI0023E141AA
Venturia_UPI001C9D0A4F
Leptopilina_UPI001CAA39E0
Bradysia_coprophila_GRC_OZ247521.1_002413.1
Allacma_fusca_CAG7719696.1
Orchesella_dallaii_CAL8116060.1
Orchesella_dallaii_CAL8116058.1
N140
Orchesella_cincta_ODM92453.1
Folsomia_candida_XP_035709242.1
Daphnia_UPI001E1BCCE0
Daphnia_UPI001EE03E4F
Rotaria_A0A821LZP7
Rotaria_A0A814L5S5
Tigriopus_A0A553NPV8
Dalotia_UPI0031F3BC9B
Topomyia_UPI00273BC592
Littorina_A0AAN9GNK1
Branchiostoma_A0A9J7MJR4
Bradysia_UPI00187D95A4
Teleopsis_UPI0018CDC319
Lutzomyia_UPI0024835B9E
Globodera_A0A914HZH1
Anguilla_A0A9D3MQG0
Apostichopus_A0A2G8L7A8
Bradysia_UPI00187D8EF6
Pectinidae_A0A210PP96
Rhopilema_UPI0031D0C2EF
_P35362
Contarinia_UPI0012D463A4
Pseudolycoriella_A0A9Q0N6U9
Bradysia_UPI00187DC6F9
Pristionchus_A0AAN5I6D1
Pseudolycoriella_A0A9Q0RZ87
Mytilus_A0A6J8A4C5
Dermatophagoides_A0A922L0R1
Sitodiplosis_UPI002443FBAA
Pseudolycoriella_A0A9Q0S4G1
Pseudolycoriella_A0A9Q0MTT9
Contarinia_UPI0012D439F5
Pomacea_A0A2T7PXK0
Sitodiplosis_UPI002444181E
Contarinia_UPI0012D4113A
Sciaridae_A0A9Q0MZH5
Amphibalanus_UPI001C90DC71
Homo_UPI001035505B
Dermatophagoides_A0A922HZD0
Dermatophagoides_A0A6P6XVK7
Sitodiplosis_UPI0024442E01
Bradysia_UPI00187D7834
Panonychus_UPI002307A5CF
Exaiptasia_A0A913WYG3
Sitodiplosis_UPI0024446A23
Bradysia_UPI00187D6F96
Bradysia_UPI00187D9CC3
Viruses_A0A7R7V7H8
Viruses_A0A6G6CHV0
Viruses_A0A6G6CHR5
Viruses_Q86917
Viruses_A0A2R4KYQ4
Viruses_Q6TV00
Viruses_Q9DHV5
Viruses_Q6TUM9
Viruses_Q9DHG8
Viruses_Q08520
Viruses_A0A0B4Q697
Viruses_A3F194
Viruses_A3F193
Viruses_Q89609
Viruses_A3F178
Viruses_A3F188
Viruses_A0A7D0Q8H7
Viruses_A0A286MM95
Viruses_A0A890UXG8
Viruses_A0A1V0S7Q1
Viruses_A0A649Z1W3
Viruses_Q9J5H4
Viruses_A0A068EHD5
Viruses_A0A0M3ZHE1
Viruses_A0AAE9NXT8
Viruses_A0A1V0S8A5
Viruses_UPI00025EFAFE
Viruses_UPI000255FFAB
Contarinia_UPI0012D3C6DA
Contarinia_UPI0012D474B8
Sciaridae_A0A9Q0MNZ7
Bradysia_UPI00187DD931
Aldrovandia_A0AAD7WYR0
Sciaridae_A0A9Q0NBH9
Bradysia_UPI00187DD51D
Stomoxys_A0A1I8Q6N4
Homarus_UPI001C490AF1
Apostichopus_A0A2G8K7J1
Dermatophagoides_A0A6P6YH89
Anopheles_UPI00237C0F86
Hypsibius_A0A1W0XFJ0
Sitodiplosis_UPI002444E137
Lytechinus_UPI0030B9B77A
Spizellomyces_A0A507EVH5
Octopus_A0A6P7SRV7
Daphnia_UPI001EDF2443
Ctenocephalides_UPI000E6E5482
Branchiostoma_A0A8J9YY85
Branchiostoma_C3YIE1
Macrostomum_A0A1I8FU19
Stylophora_UPI000C0514E5
Taxa
Cluster representative
Viruses
Eukaryota
Fungi
Metazoa
Insecta
Collembola
Sciaridae
Sciaridae_uniref
NA

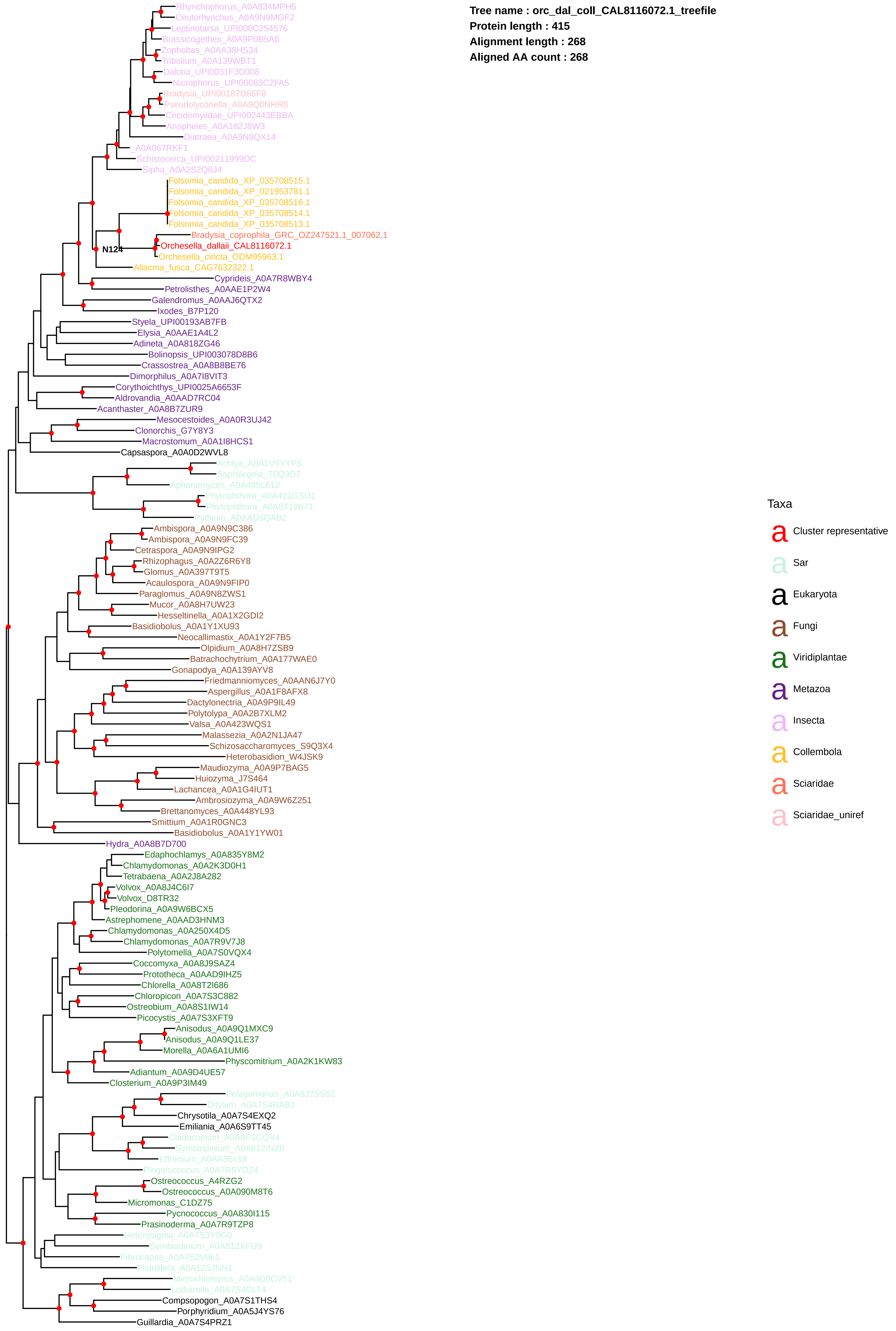

Tree name : orc_dal_coll_CAL8116072.1_treefile
Protein length : 415
Alignment length : 268
Aligned AA count : 268
Rhynchophorus_A0A834MPH5
Ceutorhynchus_A0A9N9MGF2
Leptinotarsa_UPI000C254576
Brassicogethes_A0A9P0B5A6
Zophobas_A0A38HS34
Tribolium_A0A139WBT1
Dalotia_UPI0031F3D008
Nicrophorus_UPI00083C2FA5
Bradysia_UPI00187D86F8
Pseudolycoriella_A0A9Q0NHR8
Cecidomyiidae_UPI002443EBBA
Anopheles_A0A182J8W3
Diatraea_A0A9N9QX14
_A0A067RKF1
Schistocerca_UPI00211999DC
Sipha_A0A2S2Q6J4
Folsomia_candida_XP_035708515.1
Folsomia_candida_XP_021953781.1
Folsomia_candida_XP_035708516.1
Folsomia_candida_XP_035708514.1
Folsomia_candida_XP_035708513.1
Bradysia_coprophila_GRC_OZ247521.1_007062.1
N124
Orchesella_dallaii_CAL8116072.1
Orchesella_cincta_ODM95963.1
Allacma_fusca_CAG7632322.1
Cyprideis_A0A7R8WBY4
Petrolisthes_A0AAE1P2W4
Galendromus_A0AAJ6QTX2
Ixodes_B7P120
Styela_UPI00193AB7FB
Elysia_A0AAE1A4L2
Adineta_A0A818ZG46
Bolinopsis_UPI003078D8B6
Crassostrea_A0A8B8BE76
Dimorphilus_A0A7I8VIT3
Corythoichthys_UPI0025A6653F
Aldrovandia_A0AAD7RC04
Acanthaster_A0A8B7ZUR9
Mesocestoides_A0A0R3UJ42
Clonorchis_G7Y8Y3
Macrostomum_A0A1I8HCS1
Capsaspora_A0A0D2WVL8
Achlya_A0A1V9YYP5
Saprolegnia_T0Q9D7
Aphanomyces_A0A485L612
Phytophthora_A0A421GSU1
Phytophthora_A0A8T1WI71
Pythium_A0AAD5QAB2
Ambispora_A0A9N9C386
Ambispora_A0A9N9FC39
Cetraspora_A0A9N9IPG2
Rhizophagus_A0A2Z6R6Y8
Glomus_A0A397T9T5
Acaulospora_A0A9N9FIP0
Paraglomus_A0A9N8ZWS1
Mucor_A0A8H7UW23
Hesseltinella_A0A1X2GDI2
Basidiobolus_A0A1Y1XU93
Neocallimastix_A0A1Y2F7B5
Olpidium_A0A8H7ZSB9
Batrachochytrium_A0A177WAE0
Gonapodya_A0A139AYV8
Friedmanniomyces_A0AAN6J7Y0
Aspergillus_A0A1F8AFX8
Dactylonectria_A0A9P9IL49
Polytolypa_A0A2B7XLM2
Valsa_A0A423WQS1
Malassezia_A0A2N1JA47
Schizosaccharomyces_S9Q3X4
Heterobasidion_W4JSK9
Maudiozyma_A0A9P7BAG5
Huiozyma_J7S464
Lachancea_A0A1G4IUT1
Ambrosiozyma_A0A9W6Z251
Brettanomyces_A0A448YL93
Smittium_A0A1R0GNC3
Basidiobolus_A0A1Y1YW01
Hydra_A0A8B7D700
Edaphochlamys_A0A835Y8M2
Chlamydomonas_A0A2K3D0H1
Tetrabaena_A0A2J8A282
Volvox_A0A8J4C6I7
Volvox_D8TR32
Pleodorina_A0A9W6BCX5
Astrephomene_A0AAD3HNM3
Chlamydomonas_A0A250X4D5
Chlamydomonas_A0A7R9V7J8
Polytomella_A0A7S0VQX4
Coccomyxa_A0A8J9SAZ4
Prototheca_A0AAD9IHZ5
Chlorella_A0A8T2I686
Chloropicon_A0A7S3C882
Ostreobium_A0A8S1IW14
Picocystis_A0A7S3XFT9
Anisodus_A0A9Q1MXC9
Anisodus_A0A9Q1LE37
Morella_A0A6A1UMI6
Physcomitrium_A0A2K1KW83
Adiantum_A0A9D4UE57
Closterium_A0A9P3IM49
Pelagomonas_A0A8J2SS62
Ditylum_A0A7S4RAB1
Chrysotila_A0A7S4EXQ2
Emiliania_A0A6S9TT45
Cladocopium_A0A9P1GQV4
Symbiodinium_A0A812INZ6
Effrenium_A0AA36II18
Pinguiococcus_A0A7R9YDZ4
Ostreococcus_A4RZG2
Ostreococcus_A0A090M8T6
Micromonas_C1DZ75
Pycnococcus_A0A830I115
Prasinoderma_A0A7R9TZP8
Heterosigma_A0A7S3Y0G0
Symbiodinium_A0A812XFU9
Fibrocapsa_A0A7S2V0L1
Fistulifera_A0A1Z5JNN1
Microchloropsis_A0A4D9CV51
Lotharella_A0A7S4DLT4
Compsopogon_A0A7S1THS4
Porphyridium_A0A5J4YS76
Guillardia_A0A7S4PRZ1
Taxa
Cluster representative
Sar
Eukaryota
Fungi
Viridiplantae
Metazoa
Insecta
Collembola
Sciaridae
Sciaridae_uniref

**Tree name : orc_dal_coll_CAL8129638.1_treefile**
**Protein length : 502**
**Alignment length : 423**
**Aligned AA count : 423**

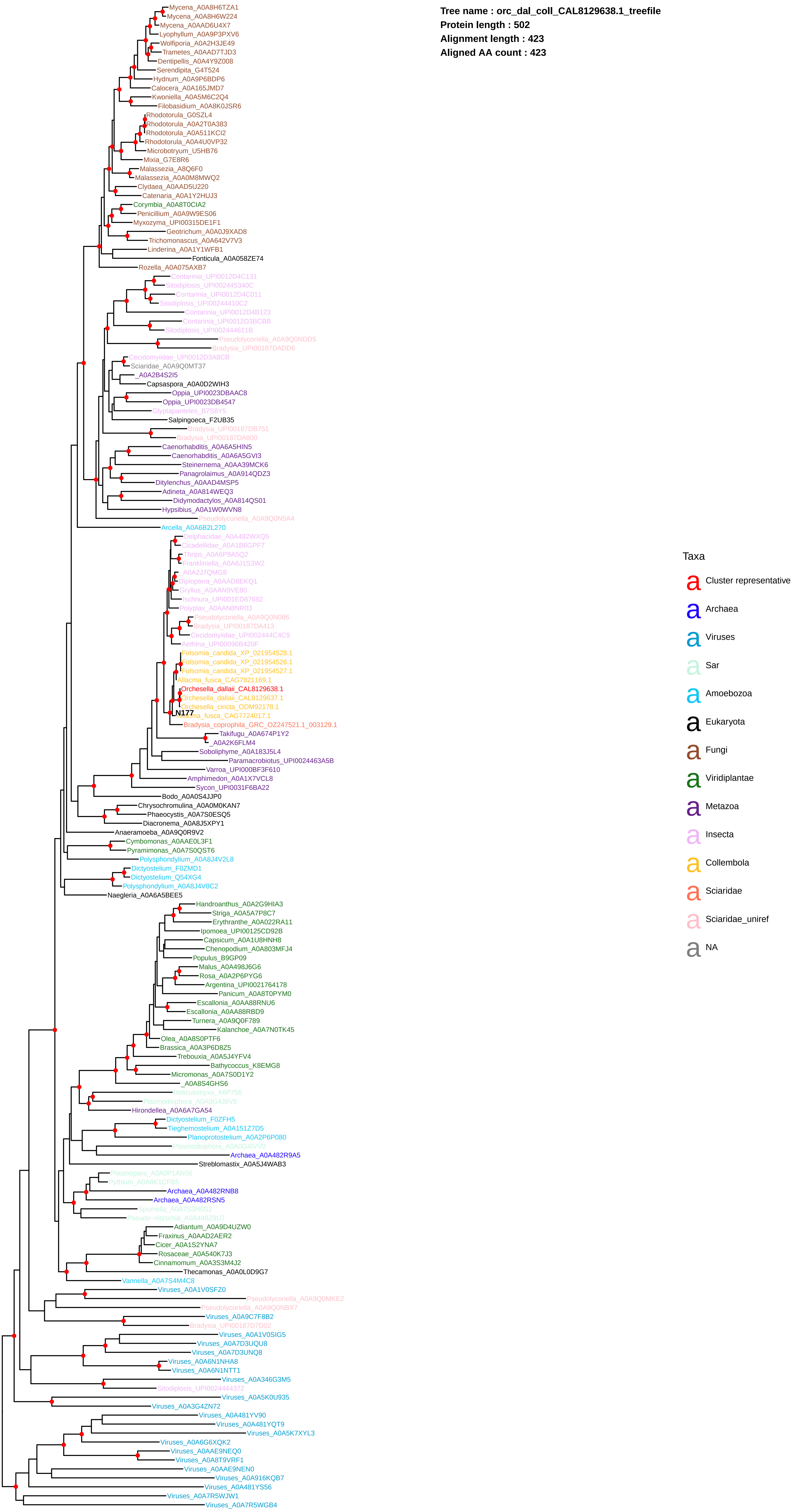

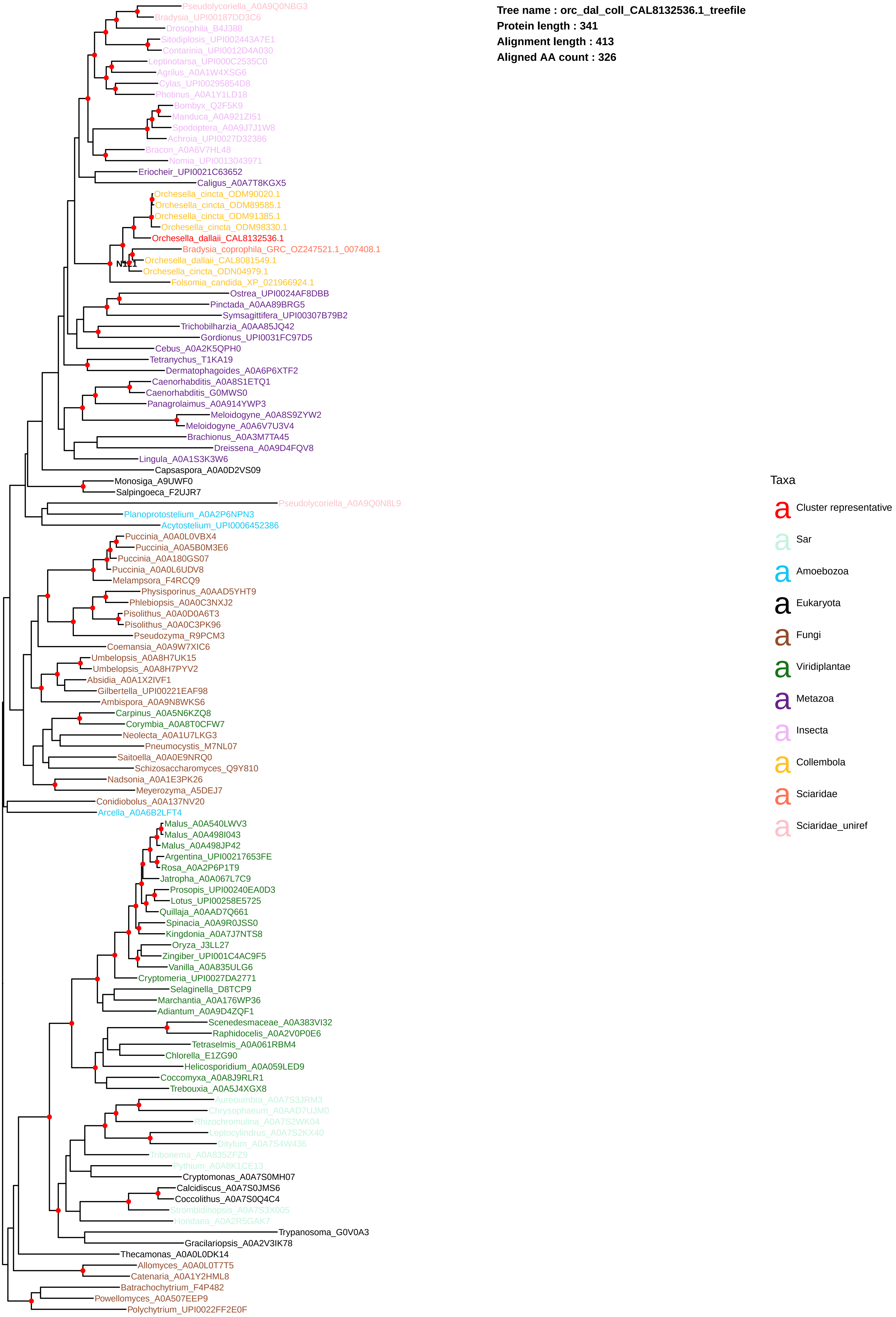

Tree name : orc_dal_coll_CAL8132536.1_treefile
Protein length : 341
Alignment length : 413
Aligned AA count : 326
Pseudolycoriella_A0A9Q0NBG3
Bradysia_UPI00187DD3C6
Drosophila_B4J388
Sitodiplosis_UPI002443A7E1
Contarinia_UPI0012D4A030
Leptinotarsa_UPI000C2535C0
Agrilus_A0A1W4XSG6
Cylas_UPI00295854D8
Photinus_A0A1Y1LD18
Bombyx_Q2F5K9
Manduca_A0A921ZI51
Spodoptera_A0A9J7J1W8
Achroia_UPI0027D32386
Bracon_A0A6V7HL48
Nomia_UPI0013043971
Eriocheir_UPI0021C63652
Caligus_A0A7T8KGX5
Orchesella_cincta_ODM90020.1
Orchesella_cincta_ODM89585.1
Orchesella_cincta_ODM91385.1
Orchesella_cincta_ODM98330.1
Orchesella_dallaii_CAL8132536.1
Bradysia_coprophila_GRC_OZ247521.1_007408.1
N1.1
Orchesella_dallaii_CAL8081549.1
Orchesella_cincta_ODN04979.1
Folsomia_candida_XP_021966924.1
Ostrea_UPI0024AF8DBB
Pinctada_A0AA89BRG5
Symsagittifera_UPI00307B79B2
Trichobilharzia_A0AA85JQ42
Gordionus_UPI0031FC97D5
Cebus_A0A2K5QPH0
Tetranychus_T1KA19
Dermatophagoides_A0A6P6XTF2
Caenorhabditis_A0A8S1ETQ1
Caenorhabditis_G0MWS0
Panagrolaimus_A0A914YWP3
Meloidogyne_A0A8S9ZYW2
Meloidogyne_A0A6V7U3V4
Brachionus_A0A3M7TA45
Dreissena_A0A9D4FQV8
Lingula_A0A1S3K3W6
Capsaspora_A0A0D2VS09
Monosiga_A9UWF0
Salpingoeca_F2UJR7
Pseudolycoriella_A0A9Q0N8L9
Planoprotostelium_A0A2P6NPN3
Acytostelium_UPI0006452386
Puccinia_A0A0L0VBX4
Puccinia_A0A5B0M3E6
Puccinia_A0A180GS07
Puccinia_A0A0L6UDV8
Melampsora_F4RCQ9
Physisporinus_A0AAD5YHT9
Phlebiopsis_A0A0C3NXJ2
Pisolithus_A0A0D0A6T3
Pisolithus_A0A0C3PK96
Pseudozyma_R9PCM3
Coemansia_A0A9W7XIC6
Umbelopsis_A0A8H7UK15
Umbelopsis_A0A8H7PYV2
Absidia_A0A1X2IVF1
Gilbertella_UPI00221EAF98
Ambispora_A0A9N8WKS6
Carpinus_A0A5N6KZQ8
Corymbia_A0A8T0CFW7
Neolecta_A0A1U7LKG3
Pneumocystis_M7NL07
Saitoella_A0A0E9NRQ0
Schizosaccharomyces_Q9Y810
Nadsonia_A0A1E3PK26
Meyerozyma_A5DEJ7
Conidiobolus_A0A137NV20
Arcella_A0A6B2LFT4
Malus_A0A540LWV3
Malus_A0A498I043
Malus_A0A498JP42
Argentina_UPI00217653FE
Rosa_A0A2P6P1T9
Jatropha_A0A067L7C9
Prosopis_UPI00240EA0D3
Lotus_UPI00258E5725
Quillaja_A0AAD7Q661
Spinacia_A0A9R0JSS0
Kingdonia_A0A7J7NTS8
Oryza_J3LL27
Zingiber_UPI001C4AC9F5
Vanilla_A0A835ULG6
Cryptomeria_UPI0027DA2771
Selaginella_D8TCP9
Marchantia_A0A176WP36
Adiantum_A0A9D4ZQF1
Scenedesmaceae_A0A383VI32
Raphidocelis_A0A2V0P0E6
Tetraselmis_A0A061RBM4
Chlorella_E1ZG90
Helicosporidium_A0A059LED9
Coccomyxa_A0A8J9RLR1
Trebouxia_A0A5J4XGX8
Aureoumbra_A0A7S3JRM3
Chrysophaeum_A0AAD7UJM0
Rhizochromulina_A0A7S2WK04
Leptocylindrus_A0A7S2KX40
Ditylum_A0A7S4W436
Tribonema_A0A835ZFZ9
Pythium_A0A8K1CE13
Cryptomonas_A0A7S0MH07
Calcidiscus_A0A7S0JMS6
Coccolithus_A0A7S0Q4C4
Strombidinopsis_A0A7S3X005
Hondaea_A0A2R5GAK7
Trypanosoma_G0V0A3
Gracilariopsis_A0A2V3IK78
Thecamonas_A0A0L0DK14
Allomyces_A0A0L0T7T5
Catenaria_A0A1Y2HML8
Batrachochytrium_F4P482
Powellomyces_A0A507EEP9
Polychytrium_UPI0022FF2E0F
Taxa
a Cluster representative
a Sar
a Amoebozoa
a Eukaryota
a Fungi
a Viridiplantae
a Metazoa
a Insecta
a Collembola
a Sciaridae
a Sciaridae_uniref

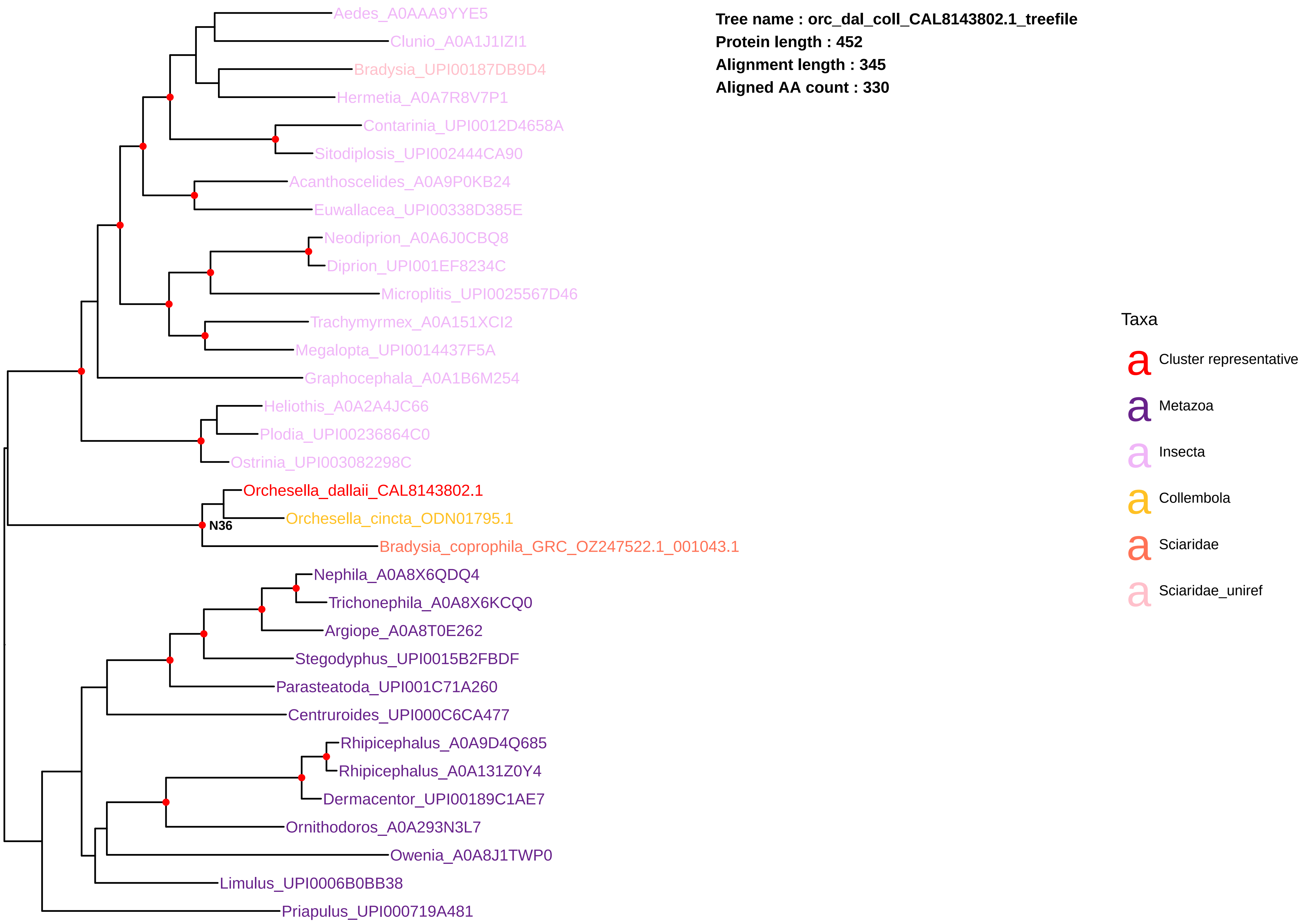
Tree name : orc_dal_coll_CAL8143802.1_treefile
Protein length : 452
Alignment length : 345
Aligned AA count : 330
Aedes_A0AAA9YYE5
Clunio_A0A1J1IZI1
Bradysia_UPI00187DB9D4
Hermetia_A0A7R8V7P1
Contarinia_UPI0012D4658A
Sitodiplosis_UPI002444CA90
Acanthoscelides_A0A9P0KB24
Euwallacea_UPI00338D385E
Neodiprion_A0A6J0CBQ8
Diprion_UPI001EF8234C
Microplitis_UPI0025567D46
Trachymyrmex_A0A151XCI2
Megalopta_UPI0014437F5A
Graphocephala_A0A1B6M254
Heliothis_A0A2A4JC66
Plodia_UPI00236864C0
Ostrinia_UPI003082298C
Orchesella_dallaii_CAL8143802.1
Orchesella_cincta_ODN01795.1
N36
Bradysia_coprophila_GRC_OZ247522.1_001043.1
Nephila_A0A8X6QDQ4
Trichonephila_A0A8X6KCQ0
Argiope_A0A8T0E262
Stegodyphus_UPI0015B2FBDF
Parasteatoda_UPI001C71A260
Centruroides_UPI000C6CA477
Rhipicephalus_A0A9D4Q685
Rhipicephalus_A0A131Z0Y4
Dermacentor_UPI00189C1AE7
Ornithodoros_A0A293N3L7
Owenia_A0A8J1TWP0
Limulus_UPI0006B0BB38
Priapulus_UPI000719A481
Taxa
Cluster representative
Metazoa
Insecta
Collembola
Sciaridae
Sciaridae_uniref

**Tree name : orc_dal_coll_CAL8143809.1_treefile**
**Protein length : 468**
**Alignment length : 225**
**Aligned AA count : 222**

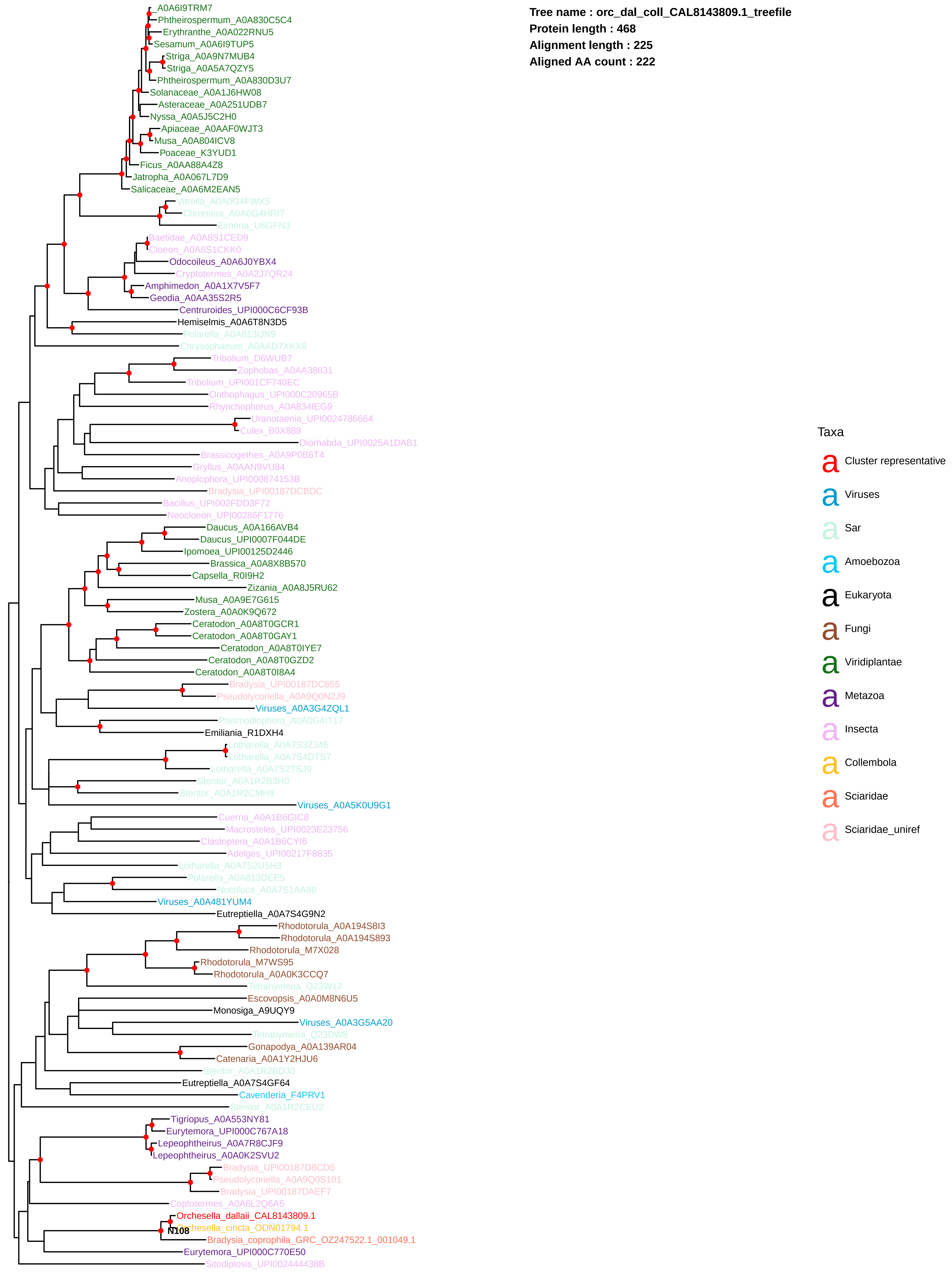

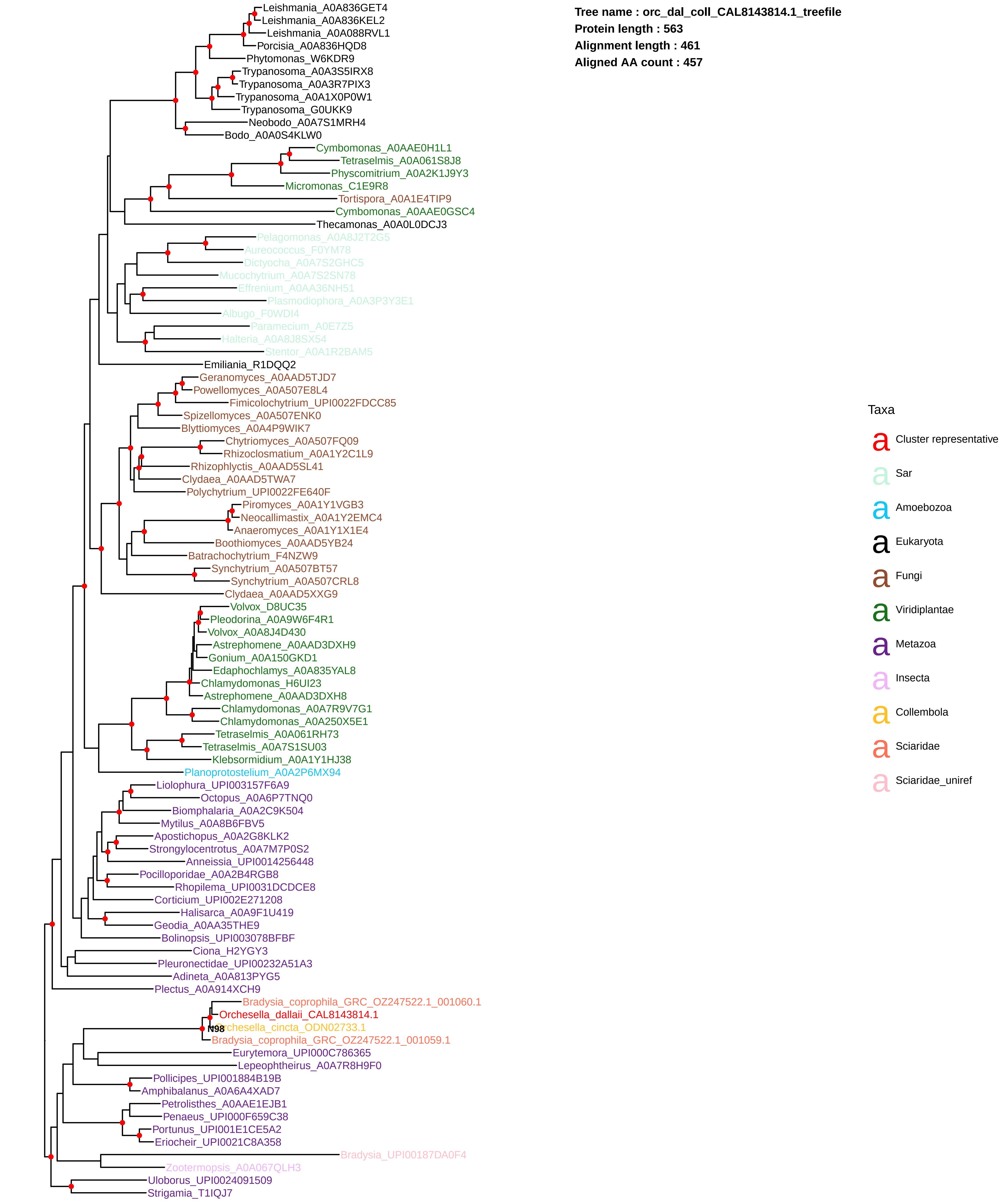
Tree name : orc_dal_coll_CAL8143814.1_treefile
Protein length : 563
Alignment length : 461
Aligned AA count : 457
Leishmania_A0A836GET4
Leishmania_A0A836KEL2
Leishmania_A0A088RVL1
Porcisia_A0A836HQD8
Phytomonas_W6KDR9
Trypanosoma_A0A3S5IRX8
Trypanosoma_A0A3R7PIX3
Trypanosoma_A0A1X0P0W1
Trypanosoma_G0UKK9
Neobodo_A0A7S1MRH4
Bodo_A0A0S4KLW0
Cymbomonas_A0AAE0H1L1
Tetraselmis_A0A061S8J8
Physcomitrium_A0A2K1J9Y3
Micromonas_C1E9R8
Tortispora_A0A1E4TIP9
Cymbomonas_A0AAE0GSC4
Thecamonas_A0A0L0DCJ3
Pelagomonas_A0A8J2T2G5
Aureococcus_F0YM78
Dictyocha_A0A7S2GHC5
Mucochytrium_A0A7S2SN78
Effrenium_A0AA36NH51
Plasmodiophora_A0A3P3Y3E1
Albugo_F0WDI4
Paramecium_A0E7Z5
Halteria_A0A8J8SX54
Stentor_A0A1R2BAM5
Emiliania_R1DQQ2
Geranomyces_A0AAD5TJD7
Powellomyces_A0A507E8L4
Fimicolochytrium_UPI0022FDCC85
Spizellomyces_A0A507ENK0
Blyttiomyces_A0A4P9WIK7
Chytriomyces_A0A507FQ09
Rhizoclosmatium_A0A1Y2C1L9
Rhizophlyctis_A0AAD5SL41
Clydaea_A0AAD5TWA7
Polychytrium_UPI0022FE640F
Piromyces_A0A1Y1VGB3
Neocallimastix_A0A1Y2EMC4
Anaeromyces_A0A1Y1X1E4
Boothiomyces_A0AAD5YB24
Batrachochytrium_F4NZW9
Synchytrium_A0A507BT57
Synchytrium_A0A507CRL8
Clydaea_A0AAD5XXG9
Volvox_D8UC35
Pleodorina_A0A9W6F4R1
Volvox_A0A8J4D430
Astrephomene_A0AAD3DXH9
Gonium_A0A150GKD1
Edaphochlamys_A0A835YAL8
Chlamydomonas_H6UI23
Astrephomene_A0AAD3DXH8
Chlamydomonas_A0A7R9V7G1
Chlamydomonas_A0A250X5E1
Tetraselmis_A0A061RH73
Tetraselmis_A0A7S1SU03
Klebsormidium_A0A1Y1HJ38
Planoprotostelium_A0A2P6MX94
Liolophura_UPI003157F6A9
Octopus_A0A6P7TNQ0
Biomphalaria_A0A2C9K504
Mytilus_A0A8B6FBV5
Apostichopus_A0A2G8KLK2
Strongylocentrotus_A0A7M7P0S2
Anneissia_UPI0014256448
Pocilloporidae_A0A2B4RGB8
Rhopilema_UPI0031DCDCE8
Corticium_UPI002E271208
Halisarca_A0A9F1U419
Geodia_A0AA35THE9
Bolinopsis_UPI003078BFBF
Ciona_H2YGY3
Pleuronectidae_UPI00232A51A3
Adineta_A0A813PYG5
Plectus_A0A914XCH9
Bradysia_coprophila_GRC_OZ247522.1_001060.1
Orchesella_dallaii_CAL8143814.1
N98
Orchesella_cincta_ODN02733.1
Bradysia_coprophila_GRC_OZ247522.1_001059.1
Eurytemora_UPI000C786365
Lepeophtheirus_A0A7R8H9F0
Pollicipes_UPI001884B19B
Amphibalanus_A0A6A4XAD7
Petrolisthes_A0AAE1EJB1
Penaeus_UPI000F659C38
Portunus_UPI001E1CE5A2
Eriocheir_UPI0021C8A358
Bradysia_UPI00187DA0F4
Zootermopsis_A0A067QLH3
Uloborus_UPI0024091509
Strigamia_T1IQJ7
Taxa
a Cluster representative
a Sar
a Amoebozoa
a Eukaryota
a Fungi
a Viridiplantae
a Metazoa
a Insecta
a Collembola
a Sciaridae
a Sciaridae_uniref

**Tree name : orc_dal_coll_CAL8147246.1_treefile**
**Protein length : 476**
**Alignment length : 217**
**Aligned AA count : 217**

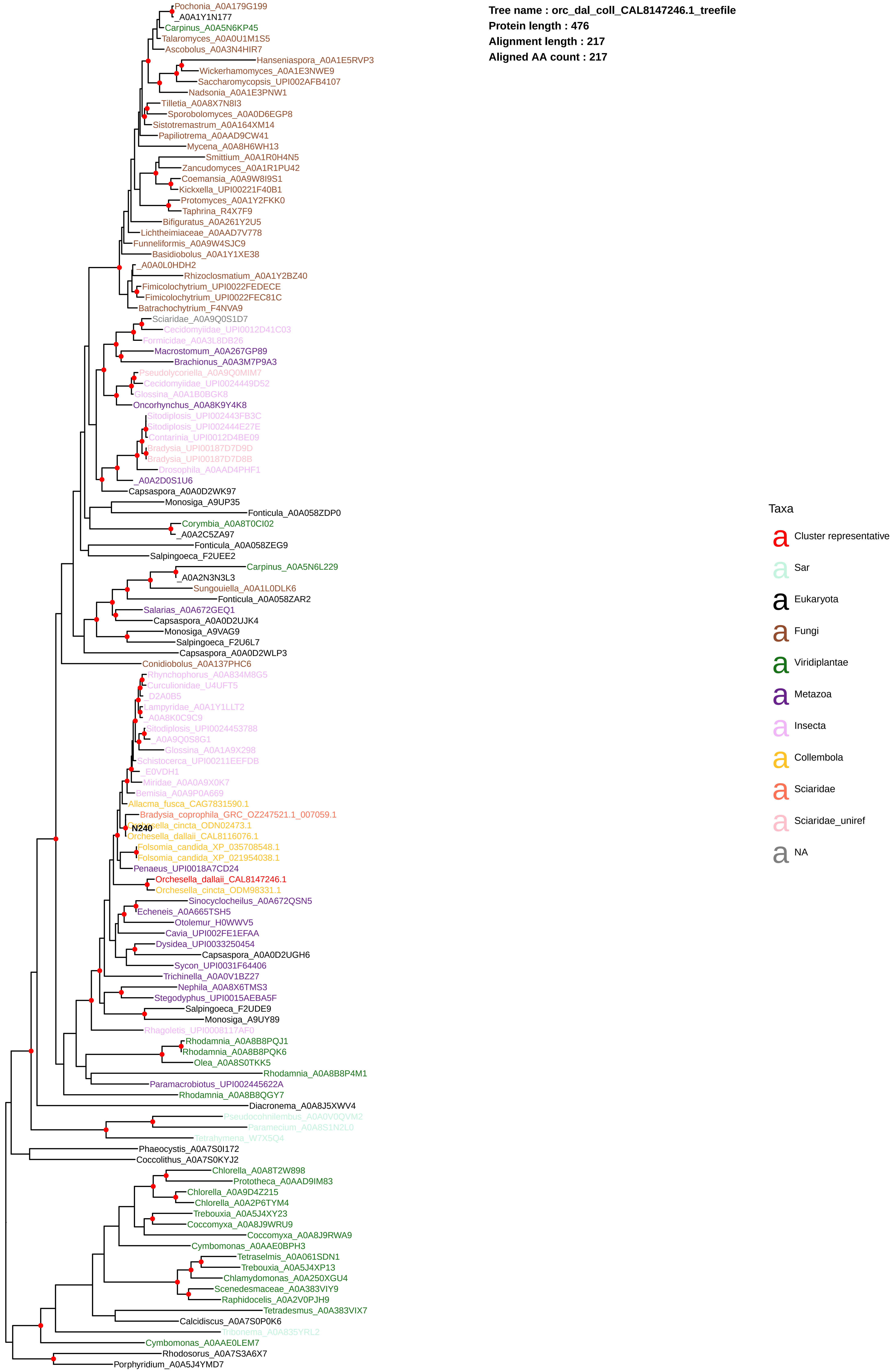

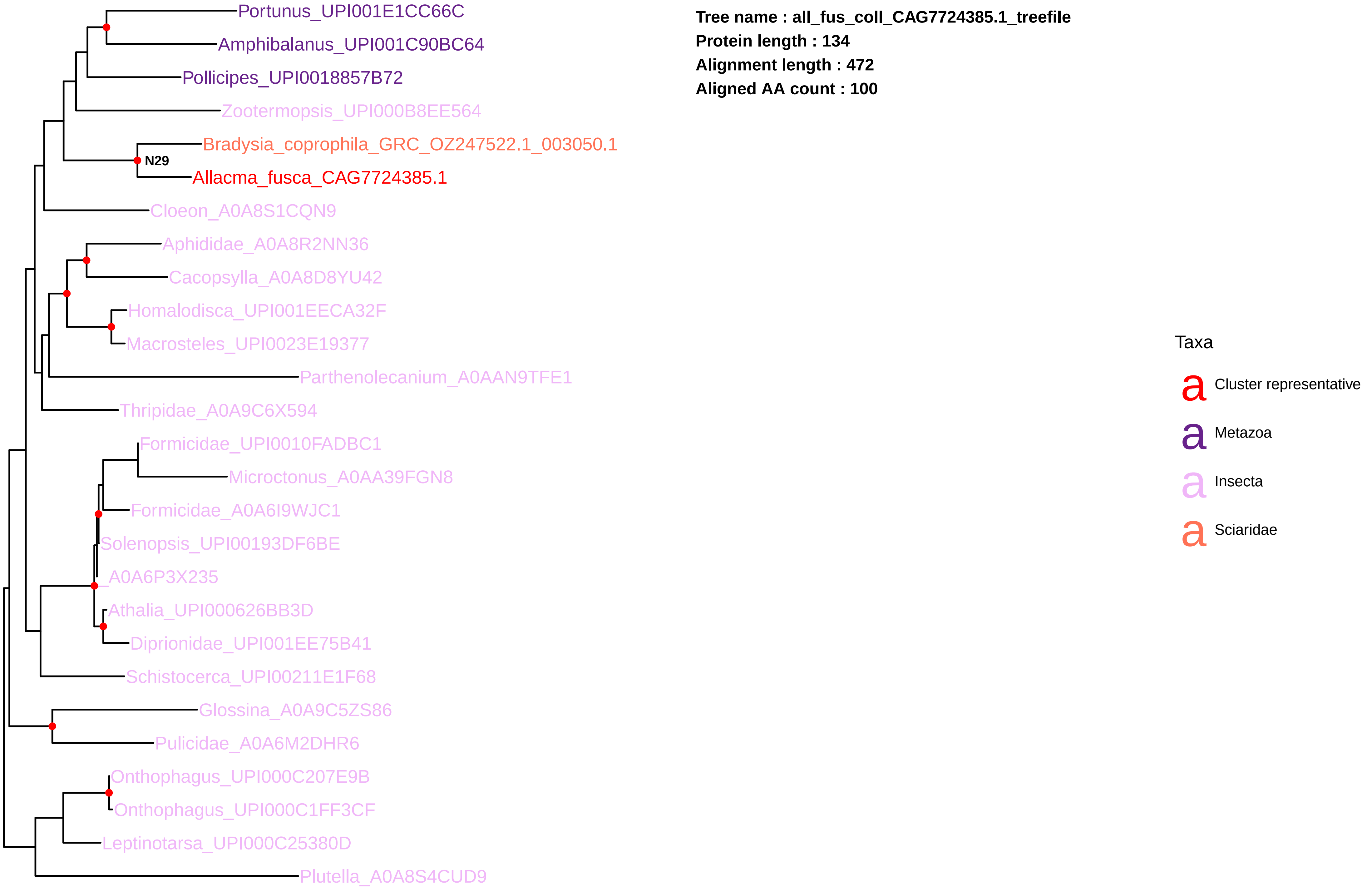
Portunus_UPI001E1CC66C
Amphibalanus_UPI001C90BC64
Pollicipes_UPI0018857B72
Zootermopsis_UPI000B8EE564
Bradysia_coprophila_GRC_OZ247522.1_003050.1
N29
Allacma_fusca_CAG7724385.1
Cloeon_A0A8S1CQN9
Aphididae_A0A8R2NN36
Cacopsylla_A0A8D8YU42
Homalodisca_UPI001EECA32F
Macrosteles_UPI0023E19377
Parthenolecanium_A0AAN9TFE1
Thripidae_A0A9C6X594
Formicidae_UPI0010FADBC1
Microctonus_A0AA39FGN8
Formicidae_A0A6I9WJC1
Solenopsis_UPI00193DF6BE
_A0A6P3X235
Athalia_UPI000626BB3D
Diprionidae_UPI001EE75B41
Schistocerca_UPI00211E1F68
Glossina_A0A9C5ZS86
Pulicidae_A0A6M2DHR6
Onthophagus_UPI000C207E9B
Onthophagus_UPI000C1FF3CF
Leptinotarsa_UPI000C25380D
Plutella_A0A8S4CUD9
Tree name : all_fus_coll_CAG7724385.1_treefile
Protein length : 134
Alignment length : 472
Aligned AA count : 100
Taxa
Cluster representative
Metazoa
Insecta
Sciaridae

**Tree name : all_fus_coll_CAG7727266.1_treefile**
**Protein length : 2055**
**Alignment length : 603**
**Aligned AA count : 603**

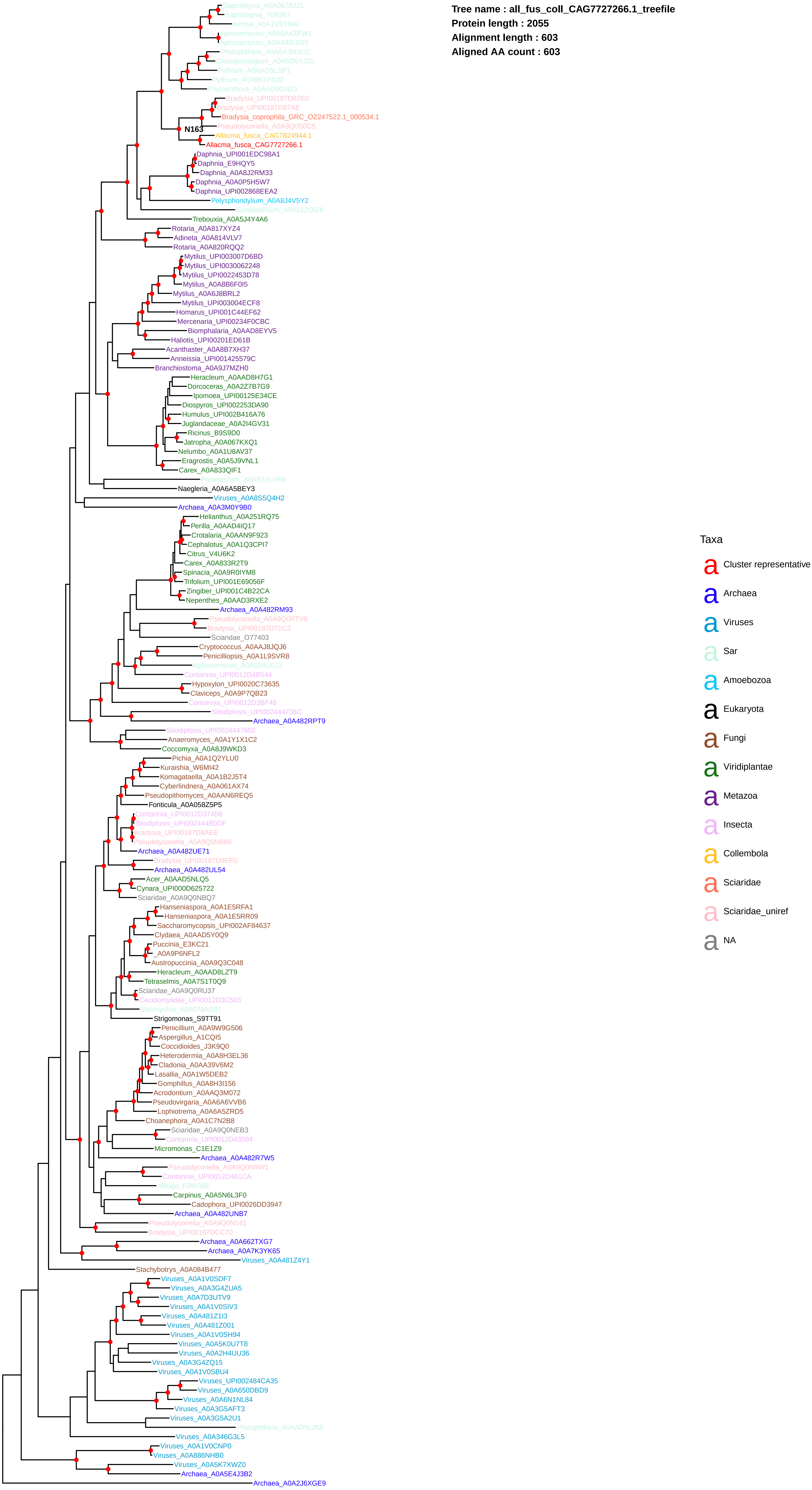

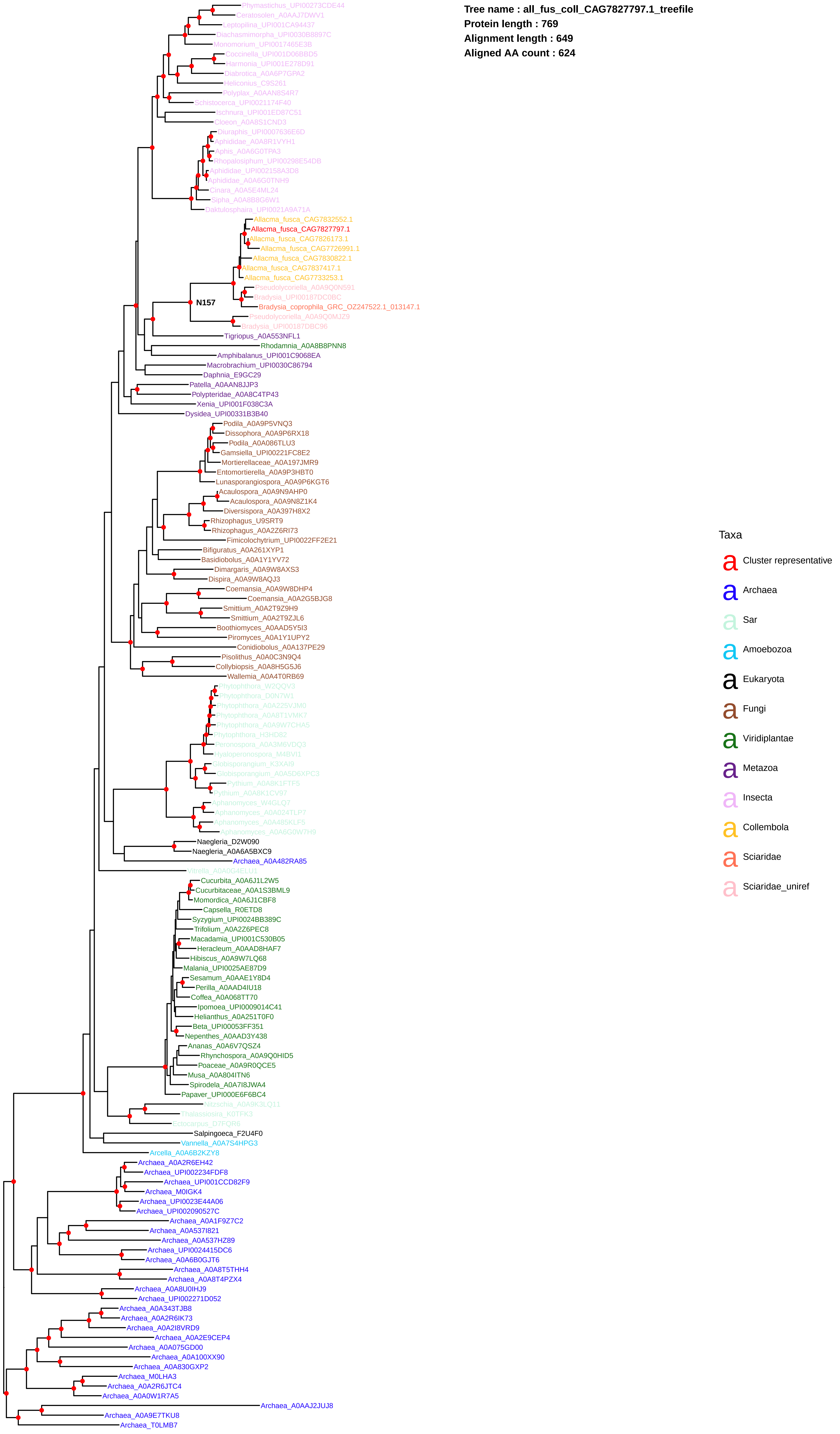
Tree name : all_fus_coll_CAG7827797.1_treefile
Protein length : 769
Alignment length : 649
Aligned AA count : 624
Phymastichus_UPI00273CDE44
Ceratosolen_A0AAJ7DWV1
Leptopilina_UPI001CA94437
Diachasmimorpha_UPI0030B8897C
Monomorium_UPI0017465E3B
Coccinella_UPI001D06BBD5
Harmonia_UPI001E278D91
Diabrotica_A0A6P7GPA2
Heliconius_C9S261
Polyplax_A0AAN8S4R7
Schistocerca_UPI0021174F40
Ischnura_UPI001ED87C51
Cloeon_A0A8S1CND3
Diuraphis_UPI0007636E6D
Aphididae_A0A8R1VYH1
Aphis_A0A6G0TPA3
Rhopalosiphum_UPI00298E54DB
Aphididae_UPI002158A3D8
Aphididae_A0A6G0TNH9
Cinara_A0A5E4ML24
Sipha_A0A8B8G6W1
Daktulosphaira_UPI0021A9A71A
Allacma_fusca_CAG7832552.1
Allacma_fusca_CAG7827797.1
Allacma_fusca_CAG7826173.1
Allacma_fusca_CAG7726991.1
Allacma_fusca_CAG7830822.1
Allacma_fusca_CAG7837417.1
Allacma_fusca_CAG7733253.1
N157
Pseudolycoriella_A0A9Q0N591
Bradysia_UPI00187DC0BC
Bradysia_coprophila_GRC_OZ247522.1_013147.1
Pseudolycoriella_A0A9Q0MJZ9
Bradysia_UPI00187DBC96
Tigriopus_A0A553NFL1
Rhodamnia_A0A8B8PNN8
Amphibalanus_UPI001C9068EA
Macrobrachium_UPI0030C86794
Daphnia_E9GC29
Patella_A0AAN8JJP3
Polypteridae_A0A8C4TP43
Xenia_UPI001F038C3A
Dysidea_UPI00331B3B40
Podila_A0A9P5VNQ3
Dissophora_A0A9P6RX18
Podila_A0A086TLU3
Gamsiella_UPI00221FC8E2
Mortierellaceae_A0A197JMR9
Entomortierella_A0A9P3HBT0
Lunasporangiospora_A0A9P6KGT6
Acaulospora_A0A9N9AHP0
Acaulospora_A0A9N8Z1K4
Diversispora_A0A397H8X2
Rhizophagus_U9SRT9
Rhizophagus_A0A2Z6RI73
Fimicolochytrium_UPI0022FF2E21
Bifiguratus_A0A261XYP1
Basidiobolus_A0A1Y1YV72
Dimargaris_A0A9W8AXS3
Dispira_A0A9W8AQJ3
Coemansia_A0A9W8DHP4
Coemansia_A0A2G5BJG8
Smittium_A0A2T9Z9H9
Smittium_A0A2T9ZJL6
Boothiomyces_A0AAD5Y5I3
Piromyces_A0A1Y1UPY2
Conidiobolus_A0A137PE29
Pisolithus_A0A0C3N9Q4
Collybiopsis_A0A8H5G5J6
Wallemia_A0A4T0RB69
Phytophthora_W2QQV3
Phytophthora_D0N7W1
Phytophthora_A0A225VJM0
Phytophthora_A0A8T1VMK7
Phytophthora_A0A9W7CHA5
Phytophthora_H3HD82
Peronospora_A0A3M6VDQ3
Hyaloperonospora_M4BVI1
Globisporangium_K3XAI9
Globisporangium_A0A5D6XPC3
Pythium_A0A8K1FTF5
Pythium_A0A8K1CV97
Aphanomyces_W4GLQ7
Aphanomyces_A0A024TLP7
Aphanomyces_A0A485KLF5
Aphanomyces_A0A6G0W7H9
Naegleria_D2W090
Naegleria_A0A6A5BXC9
Archaea_A0A482RA85
Vitrella_A0A0G4ELU1
Cucurbita_A0A6J1L2W5
Cucurbitaceae_A0A1S3BML9
Momordica_A0A6J1CBF8
Capsella_R0ETD8
Syzygium_UPI0024BB389C
Trifolium_A0A2Z6PEC8
Macadamia_UPI001C530B05
Heracleum_A0AAD8HAF7
Hibiscus_A0A9W7LQ68
Malania_UPI0025AE87D9
Sesamum_A0AAE1Y8D4
Perilla_A0AAD4IU18
Coffea_A0A068TT70
Ipomoea_UPI0009014C41
Helianthus_A0A251T0F0
Beta_UPI00053FF351
Nepenthes_A0AAD3Y438
Ananas_A0A6V7QSZ4
Rhynchospora_A0A9Q0HID5
Poaceae_A0A9R0QCE5
Musa_A0A804ITN6
Spirodela_A0A7I8JWA4
Papaver_UPI000E6F6BC4
Nitzschia_A0A9K3LQ11
Thalassiosira_K0TFK3
Ectocarpus_D7FQR6
Salpingoeca_F2U4F0
Vannella_A0A7S4HPG3
Arcella_A0A6B2KZY8
Archaea_A0A2R6EH42
Archaea_UPI002234FDF8
Archaea_UPI001CCD82F9
Archaea_M0IGK4
Archaea_UPI0023E44A06
Archaea_UPI002090527C
Archaea_A0A1F9Z7C2
Archaea_A0A537I821
Archaea_A0A537HZ89
Archaea_UPI0024415DC6
Archaea_A0A6B0GJT6
Archaea_A0A8T5THH4
Archaea_A0A8T4PZX4
Archaea_A0A8U0IHJ9
Archaea_UPI002271D052
Archaea_A0A343TJB8
Archaea_A0A2R6IK73
Archaea_A0A2I8VRD9
Archaea_A0A2E9CEP4
Archaea_A0A075GD00
Archaea_A0A100XX90
Archaea_A0A830GXP2
Archaea_M0LHA3
Archaea_A0A2R6JTC4
Archaea_A0A0W1R7A5
Archaea_A0AAJ2JUJ8
Archaea_A0A9E7TKU8
Archaea_T0LMB7
Taxa
Cluster representative
Archaea
Sar
Amoebozoa
Eukaryota
Fungi
Viridiplantae
Metazoa
Insecta
Collembola
Sciaridae
Sciaridae_uniref

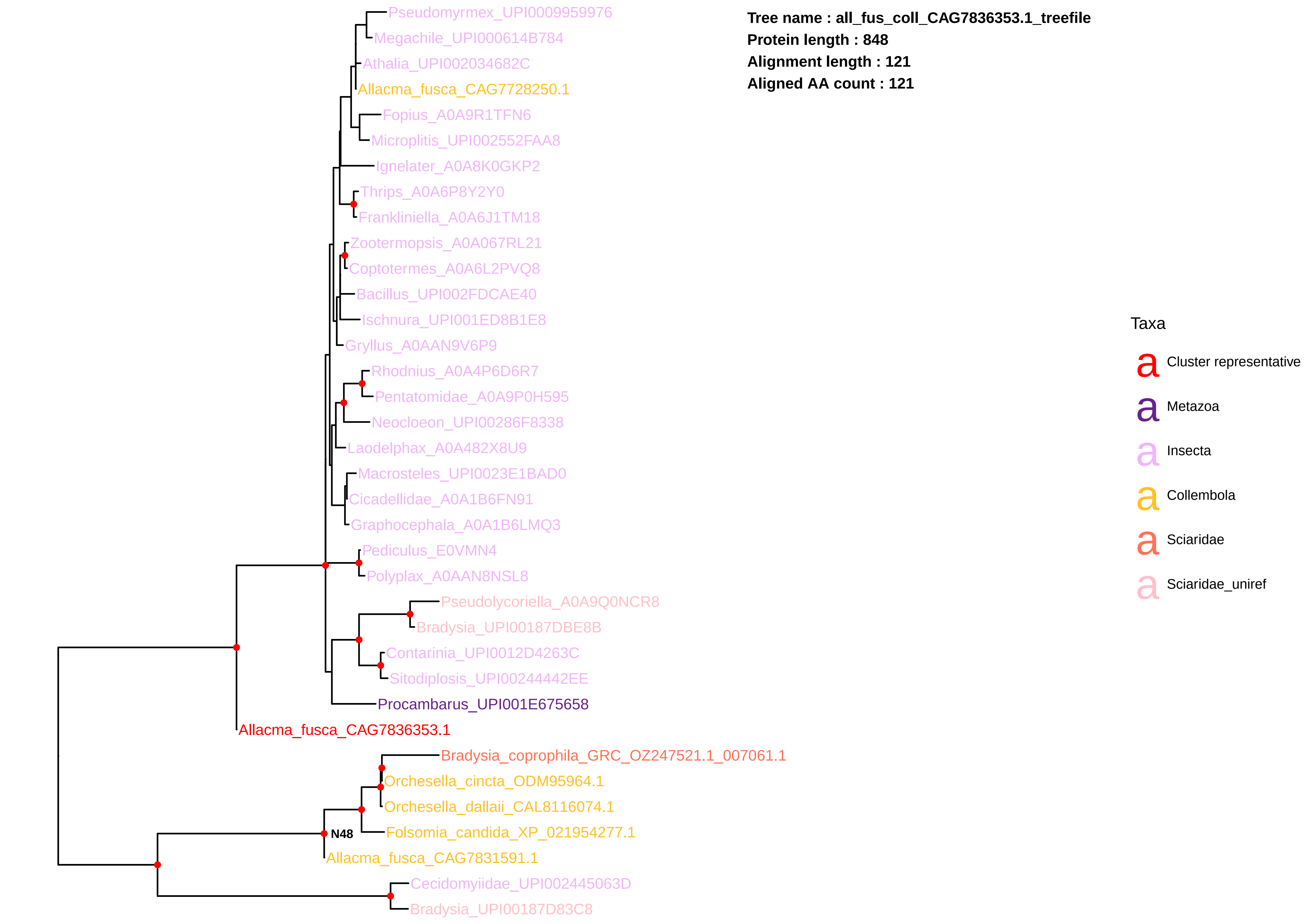
Tree name : all_fus_coll_CAG7836353.1_treefile
Protein length : 848
Alignment length : 121
Aligned AA count : 121
Pseudomyrmex_UPI0009959976
Megachile_UPI000614B784
Athalia_UPI002034682C
Allacma_fusca_CAG7728250.1
Fopius_A0A9R1TFN6
Microplitis_UPI002552FAA8
Ignelater_A0A8K0GKP2
Thrips_A0A6P8Y2Y0
Frankliniella_A0A6J1TM18
Zootermopsis_A0A067RL21
Coptotermes_A0A6L2PVQ8
Bacillus_UPI002FDCAE40
Ischnura_UPI001ED8B1E8
Gryllus_A0AAN9V6P9
Rhodnius_A0A4P6D6R7
Pentatomidae_A0A9P0H595
Neocloeon_UPI00286F8338
Laodelphax_A0A482X8U9
Macrosteles_UPI0023E1BAD0
Cicadellidae_A0A1B6FN91
Graphocephala_A0A1B6LMQ3
Pediculus_E0VMN4
Polyplax_A0AAN8NSL8
Pseudolycoriella_A0A9Q0NCR8
Bradysia_UPI00187DBE8B
Contarinia_UPI0012D4263C
Sitodiplosis_UPI00244442EE
Procambarus_UPI001E675658
Allacma_fusca_CAG7836353.1
Bradysia_coprophila_GRC_OZ247521.1_007061.1
Orchesella_cincta_ODM95964.1
Orchesella_dallaii_CAL8116074.1
Folsomia_candida_XP_021954277.1
N48
Allacma_fusca_CAG7831591.1
Cecidomyiidae_UPI002445063D
Bradysia_UPI00187D83C8
Taxa
Cluster representative
Metazoa
Insecta
Collembola
Sciaridae
Sciaridae_uniref

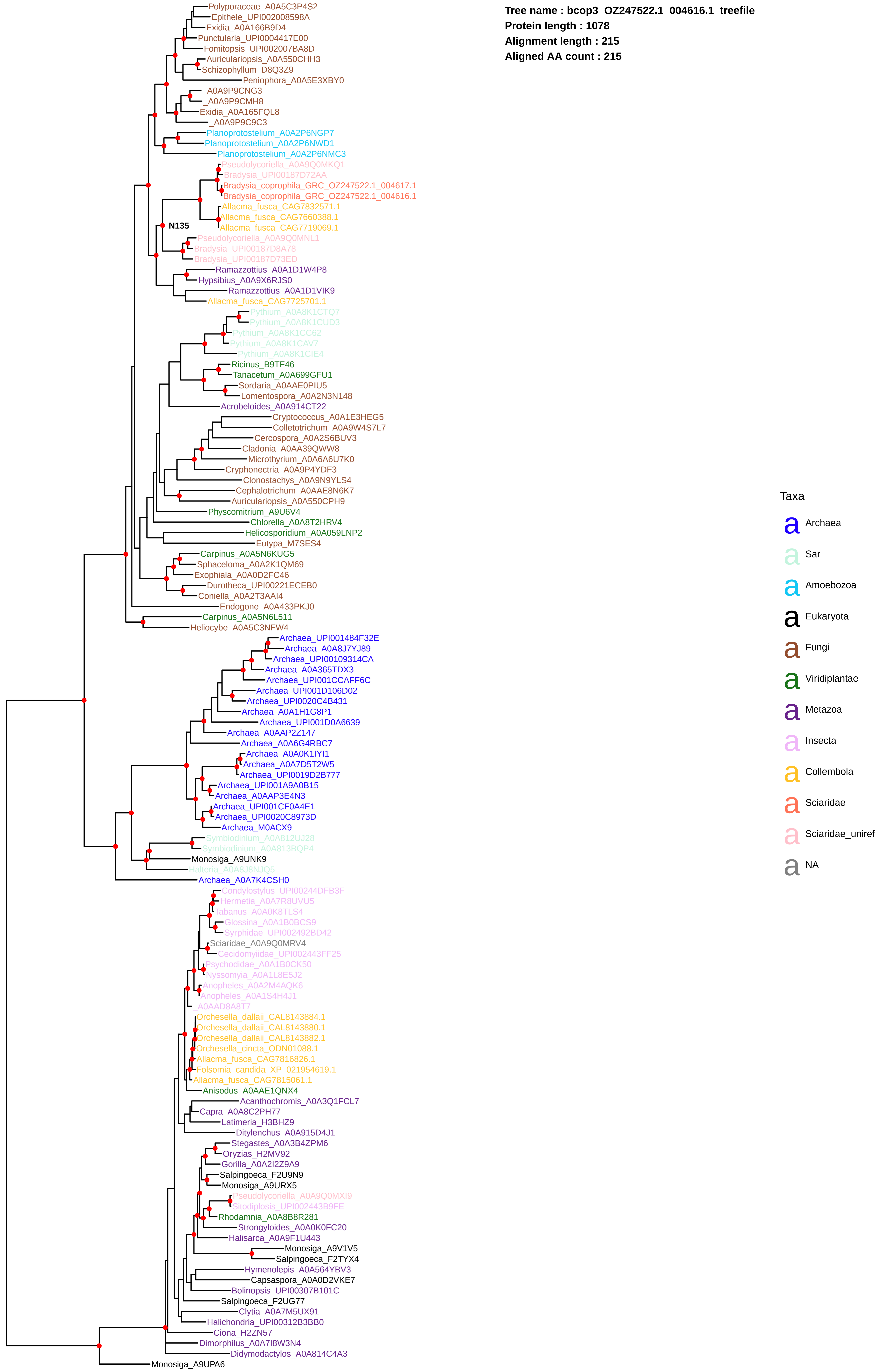
Tree name : bcop3_OZ247522.1_004616.1_treefile
Protein length : 1078
Alignment length : 215
Aligned AA count : 215
Polyporaceae_A0A5C3P4S2
Epithele_UPI002008598A
Exidia_A0A166B9D4
Punctularia_UPI0004417E00
Fomitopsis_UPI002007BA8D
Auriculariopsis_A0A550CHH3
Schizophyllum_D8Q3Z9
Peniophora_A0A5E3XBY0
_A0A9P9CNG3
_A0A9P9CMH8
Exidia_A0A165FQL8
_A0A9P9C9C3
Planoprotostelium_A0A2P6NGP7
Planoprotostelium_A0A2P6NWD1
Planoprotostelium_A0A2P6NMC3
Pseudolycoriella_A0A9Q0MKQ1
Bradysia_UPI00187D72AA
Bradysia_coprophila_GRC_OZ247522.1_004617.1
Bradysia_coprophila_GRC_OZ247522.1_004616.1
Allacma_fusca_CAG7832571.1
Allacma_fusca_CAG7660388.1
Allacma_fusca_CAG7719069.1
N135
Pseudolycoriella_A0A9Q0MNL1
Bradysia_UPI00187D8A78
Bradysia_UPI00187D73ED
Ramazzottius_A0A1D1W4P8
Hypsibius_A0A9X6RJS0
Ramazzottius_A0A1D1VIK9
Allacma_fusca_CAG7725701.1
Pythium_A0A8K1CTQ7
Pythium_A0A8K1CUD3
Pythium_A0A8K1CC62
Pythium_A0A8K1CAV7
Pythium_A0A8K1CIE4
Ricinus_B9TF46
Tanacetum_A0A699GFU1
Sordaria_A0AAE0PIU5
Lomentospora_A0A2N3N148
Acrobeloides_A0A914CT22
Cryptococcus_A0A1E3HEG5
Colletotrichum_A0A9W4S7L7
Cercospora_A0A2S6BUV3
Cladonia_A0AA39QWW8
Microthyrium_A0A6A6U7K0
Cryphonectria_A0A9P4YDF3
Clonostachys_A0A9N9YLS4
Cephalotrichum_A0AAE8N6K7
Auriculariopsis_A0A550CPH9
Physcomitrium_A9U6V4
Chlorella_A0A8T2HRV4
Helicosporidium_A0A059LNP2
Eutypa_M7SES4
Carpinus_A0A5N6KUG5
Sphaceloma_A0A2K1QM69
Exophiala_A0A0D2FC46
Durotheca_UPI00221ECEB0
Coniella_A0A2T3AAI4
Endogone_A0A433PKJ0
Carpinus_A0A5N6L511
Heliocybe_A0A5C3NFW4
Archaea_UPI001484F32E
Archaea_A0A8J7YJ89
Archaea_UPI00109314CA
Archaea_A0A365TDX3
Archaea_UPI001CCAFF6C
Archaea_UPI001D106D02
Archaea_UPI0020C4B431
Archaea_A0A1H1G8P1
Archaea_UPI001D0A6639
Archaea_A0AAP2Z147
Archaea_A0A6G4RBC7
Archaea_A0A0K1IYI1
Archaea_A0A7D5T2W5
Archaea_UPI0019D2B777
Archaea_UPI001A9A0B15
Archaea_A0AAP3E4N3
Archaea_UPI001CF0A4E1
Archaea_UPI0020C8973D
Archaea_M0ACX9
Symbiodinium_A0A812UJ28
Symbiodinium_A0A813BQP4
Monosiga_A9UNK9
Halteria_A0A8J8NJQ5
Archaea_A0A7K4CSH0
Condylostylus_UPI00244DFB3F
Hermetia_A0A7R8UVU5
Tabanus_A0A0K8TLS4
Glossina_A0A1B0BCS9
Syrphidae_UPI002492BD42
Sciaridae_A0A9Q0MRV4
Cecidomyiidae_UPI002443FF25
Psychodidae_A0A1B0CK50
Nyssomyia_A0A1L8E5J2
Anopheles_A0A2M4AQK6
Anopheles_A0A1S4H4J1
_A0AAD8A8T7
Orchesella_dallaii_CAL8143884.1
Orchesella_dallaii_CAL8143880.1
Orchesella_dallaii_CAL8143882.1
Orchesella_cincta_ODN01088.1
Allacma_fusca_CAG7816826.1
Folsomia_candida_XP_021954619.1
Allacma_fusca_CAG7815061.1
Anisodus_A0AAE1QNX4
Acanthochromis_A0A3Q1FCL7
Capra_A0A8C2PH77
Latimeria_H3BHZ9
Ditylenchus_A0A915D4J1
Stegastes_A0A3B4ZPM6
Oryzias_H2MV92
Gorilla_A0A2I2Z9A9
Salpingoeca_F2U9N9
Monosiga_A9URX5
Pseudolycoriella_A0A9Q0MXI9
Sitodiplosis_UPI002443B9FE
Rhodamnia_A0A8B8R281
Strongyloides_A0A0K0FC20
Halisarca_A0A9F1U443
Monosiga_A9V1V5
Salpingoeca_F2TYX4
Hymenolepis_A0A564YBV3
Capsaspora_A0A0D2VKE7
Bolinopsis_UPI00307B101C
Salpingoeca_F2UG77
Clytia_A0A7M5UX91
Halichondria_UPI00312B3BB0
Ciona_H2ZN57
Dimorphilus_A0A7I8W3N4
Didymodactylos_A0A814C4A3
Monosiga_A9UPA6
Taxa
Archaea
Sar
Amoebozoa
Eukaryota
Fungi
Viridiplantae
Metazoa
Insecta
Collembola
Sciaridae
Sciaridae_uniref
NA

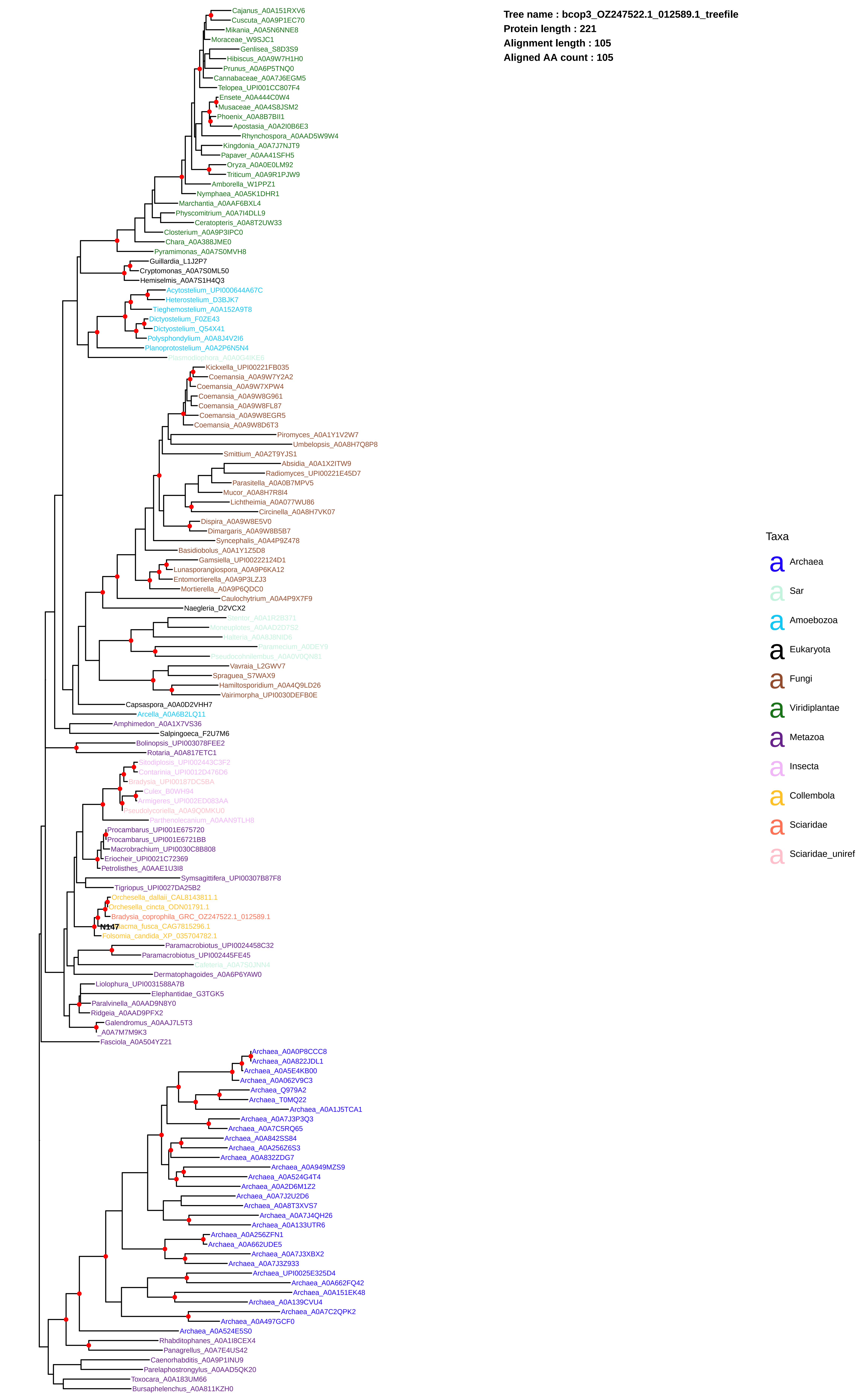
Tree name : bcop3_OZ247522.1_012589.1_treefile
Protein length : 221
Alignment length : 105
Aligned AA count : 105
Cajanus_A0A151RXV6
Cuscuta_A0A9P1EC70
Mikania_A0A5N6NNE8
Moraceae_W9SJC1
Genlisea_S8D3S9
Hibiscus_A0A9W7H1H0
Prunus_A0A6P5TNQ0
Cannabaceae_A0A7J6EGM5
Telopea_UPI001CC807F4
Ensete_A0A444C0W4
Musaceae_A0A4S8JSM2
Phoenix_A0A8B7BII1
Apostasia_A0A2I0B6E3
Rhynchospora_A0AAD5W9W4
Kingdonia_A0A7J7NJT9
Papaver_A0AA41SFH5
Oryza_A0A0E0LM92
Triticum_A0A9R1PJW9
Amborella_W1PPZ1
Nymphaea_A0A5K1DHR1
Marchantia_A0AAF6BXL4
Physcomitrium_A0A7I4DLL9
Ceratopteris_A0A8T2UW33
Closterium_A0A9P3IPC0
Chara_A0A388JME0
Pyramimonas_A0A7S0MVH8
Guillardia_L1J2P7
Cryptomonas_A0A7S0ML50
Hemiselmis_A0A7S1H4Q3
Acytostelium_UPI000644A67C
Heterostelium_D3BJK7
Tieghemostelium_A0A152A9T8
Dictyostelium_F0ZE43
Dictyostelium_Q54X41
Polysphondylium_A0A8J4V2I6
Planoprotostelium_A0A2P6N5N4
Plasmodiophora_A0A0G4IKE6
Kickxella_UPI00221FB035
Coemansia_A0A9W7Y2A2
Coemansia_A0A9W7XPW4
Coemansia_A0A9W8G961
Coemansia_A0A9W8FL87
Coemansia_A0A9W8EGR5
Coemansia_A0A9W8D6T3
Piromyces_A0A1Y1V2W7
Umbelopsis_A0A8H7Q8P8
Smittium_A0A2T9YJS1
Absidia_A0A1X2ITW9
Radiomyces_UPI00221E45D7
Parasitella_A0A0B7MPV5
Mucor_A0A8H7R8I4
Lichtheimia_A0A077WU86
Circinella_A0A8H7VK07
Dispira_A0A9W8E5V0
Dimargaris_A0A9W8B5B7
Syncephalis_A0A4P9Z478
Basidiobolus_A0A1Y1Z5D8
Gamsiella_UPI00222124D1
Lunasporangiospora_A0A9P6KA12
Entomortierella_A0A9P3LZJ3
Mortierella_A0A9P6QDC0
Caulochytrium_A0A4P9X7F9
Naegleria_D2VCX2
Stentor_A0A1R2B371
Moneuplotes_A0AAD2D7S2
Halteria_A0A8J8NID6
Paramecium_A0DEY9
Pseudocohnilembus_A0A0V0QN81
Vavraia_L2GWV7
Spraguea_S7WAX9
Hamiltosporidium_A0A4Q9LD26
Vairimorpha_UPI0030DEFB0E
Capsaspora_A0A0D2VHH7
Arcella_A0A6B2LQ11
Amphimedon_A0A1X7VS36
Salpingoeca_F2U7M6
Bolinopsis_UPI003078FEE2
Rotaria_A0A817ETC1
Sitodiplosis_UPI002443C3F2
Contarinia_UPI0012D476D6
Bradysia_UPI00187DC5BA
Culex_B0WH94
Armigeres_UPI002ED083AA
Pseudolycoriella_A0A9Q0MKU0
Parthenolecanium_A0AAN9TLH8
Procambarus_UPI001E675720
Procambarus_UPI001E6721BB
Macrobrachium_UPI0030C8B808
Eriocheir_UPI0021C72369
Petrolisthes_A0AAE1U3I8
Symsagittifera_UPI00307B87F8
Tigriopus_UPI0027DA25B2
Orchesella_dallaii_CAL8143811.1
Orchesella_cincta_ODN01791.1
Bradysia_coprophila_GRC_OZ247522.1_012589.1
N147
acma_fusca_CAG7815296.1
Folsomia_candida_XP_035704782.1
Paramacrobiotus_UPI0024458C32
Paramacrobiotus_UPI002445FE45
Cafeteria_A0A7S0JNN4
Dermatophagoides_A0A6P6YAW0
Liolophura_UPI0031588A7B
Elephantidae_G3TGK5
Paralvinella_A0AAD9N8Y0
Ridgeia_A0AAD9PFX2
Galendromus_A0AAJ7L5T3
_A0A7M7M9K3
Fasciola_A0A504YZ21
Archaea_A0A0P8CCC8
Archaea_A0A822JDL1
Archaea_A0A5E4KB00
Archaea_A0A062V9C3
Archaea_Q979A2
Archaea_T0MQ22
Archaea_A0A1J5TCA1
Archaea_A0A7J3P3Q3
Archaea_A0A7C5RQ65
Archaea_A0A842SS84
Archaea_A0A256Z6S3
Archaea_A0A832ZDG7
Archaea_A0A949MZS9
Archaea_A0A524G4T4
Archaea_A0A2D6M1Z2
Archaea_A0A7J2U2D6
Archaea_A0A8T3XVS7
Archaea_A0A7J4QH26
Archaea_A0A133UTR6
Archaea_A0A256ZFN1
Archaea_A0A662UDE5
Archaea_A0A7J3XBX2
Archaea_A0A7J3Z933
Archaea_UPI0025E325D4
Archaea_A0A662FQ42
Archaea_A0A151EK48
Archaea_A0A139CVU4
Archaea_A0A7C2QPK2
Archaea_A0A497GCF0
Archaea_A0A524E5S0
Rhabditophanes_A0A1I8CEX4
Panagrellus_A0A7E4US42
Caenorhabditis_A0A9P1INU9
Parelaphostrongylus_A0AAD5QK20
Toxocara_A0A183UM66
Bursaphelenchus_A0A811KZH0
Taxa
a Archaea
a Sar
a Amoebozoa
a Eukaryota
a Fungi
a Viridiplantae
a Metazoa
a Insecta
a Collembola
a Sciaridae
a Sciaridae_uniref

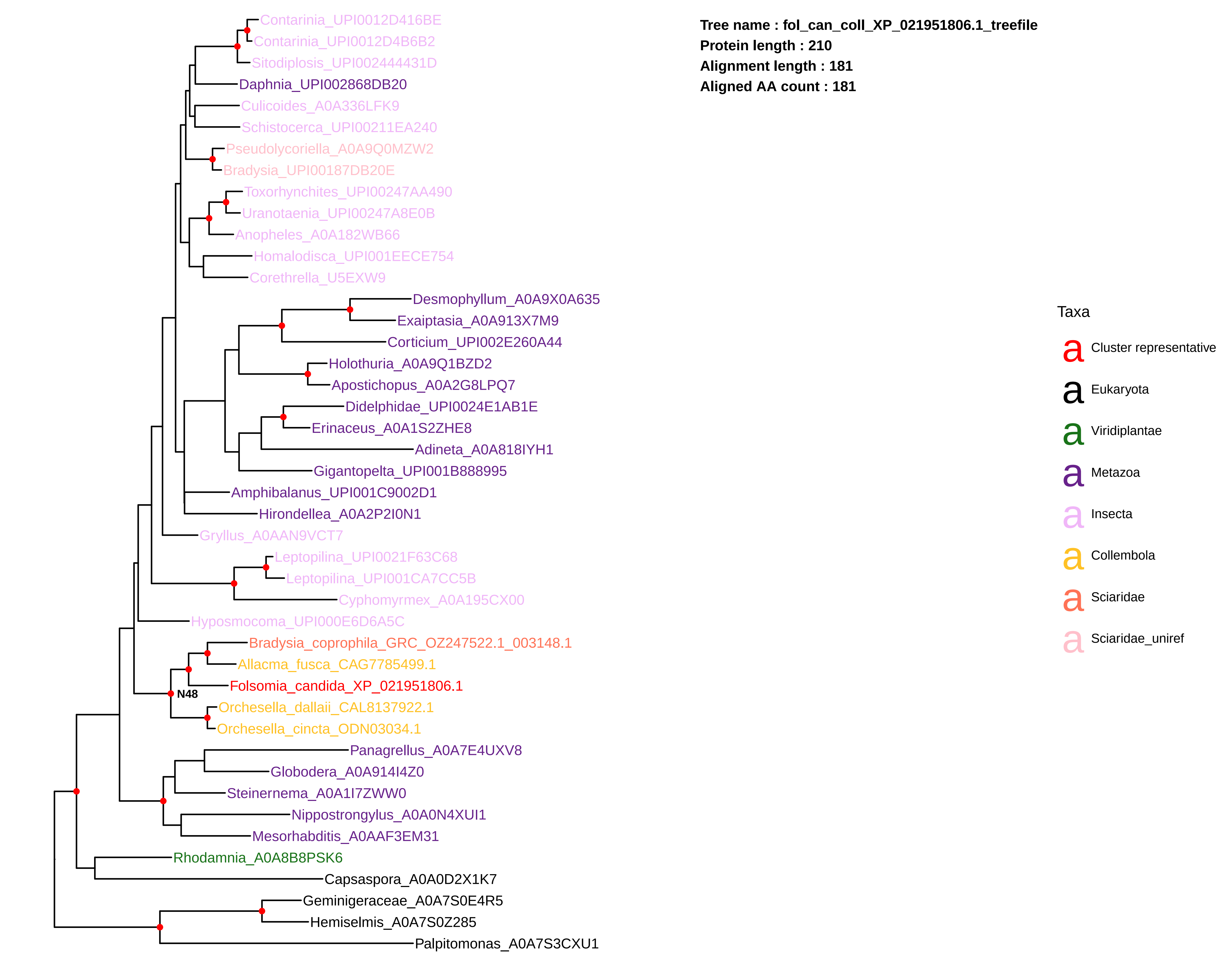

Tree name : fol_can_coll_XP_021951806.1_treefile
Protein length : 210
Alignment length : 181
Aligned AA count : 181
Contarinia_UPI0012D416BE
Contarinia_UPI0012D4B6B2
Sitodiplosis_UPI002444431D
Daphnia_UPI002868DB20
Culicoides_A0A336LFK9
Schistocerca_UPI00211EA240
Pseudolycoriella_A0A9Q0MZW2
Bradysia_UPI00187DB20E
Toxorhynchites_UPI00247AA490
Uranotaenia_UPI00247A8E0B
Anopheles_A0A182WB66
Homalodisca_UPI001EECE754
Corethrella_U5EXW9
Desmophyllum_A0A9X0A635
Exaiptasia_A0A913X7M9
Corticium_UPI002E260A44
Holothuria_A0A9Q1BZD2
Apostichopus_A0A2G8LPQ7
Didelphidae_UPI0024E1AB1E
Erinaceus_A0A1S2ZHE8
Adineta_A0A818IYH1
Gigantopelta_UPI001B888995
Amphibalanus_UPI001C9002D1
Hirondellea_A0A2P2I0N1
Gryllus_A0AAN9VCT7
Leptopilina_UPI0021F63C68
Leptopilina_UPI001CA7CC5B
Cyphomyrmex_A0A195CX00
Hyposmocoma_UPI000E6D6A5C
Bradysia_coprophila_GRC_OZ247522.1_003148.1
Allacma_fusca_CAG7785499.1
Folsomia_candida_XP_021951806.1
N48
Orchesella_dallaii_CAL8137922.1
Orchesella_cincta_ODN03034.1
Panagrellus_A0A7E4UXV8
Globodera_A0A914I4Z0
Steinernema_A0A1I7ZWW0
Nippostrongylus_A0A0N4XUI1
Mesorhabditis_A0AAF3EM31
Rhodamnia_A0A8B8PSK6
Capsaspora_A0A0D2X1K7
Geminigeraceae_A0A7S0E4R5
Hemiselmis_A0A7S0Z285
Palpitomonas_A0A7S3CXU1
Taxa
Cluster representative
Eukaryota
Viridiplantae
Metazoa
Insecta
Collembola
Sciaridae
Sciaridae_uniref

**Tree name : fol_can_coll_XP_021964531.1_treefile**
**Protein length : 198**
**Alignment length : 159**
**Aligned AA count : 159**

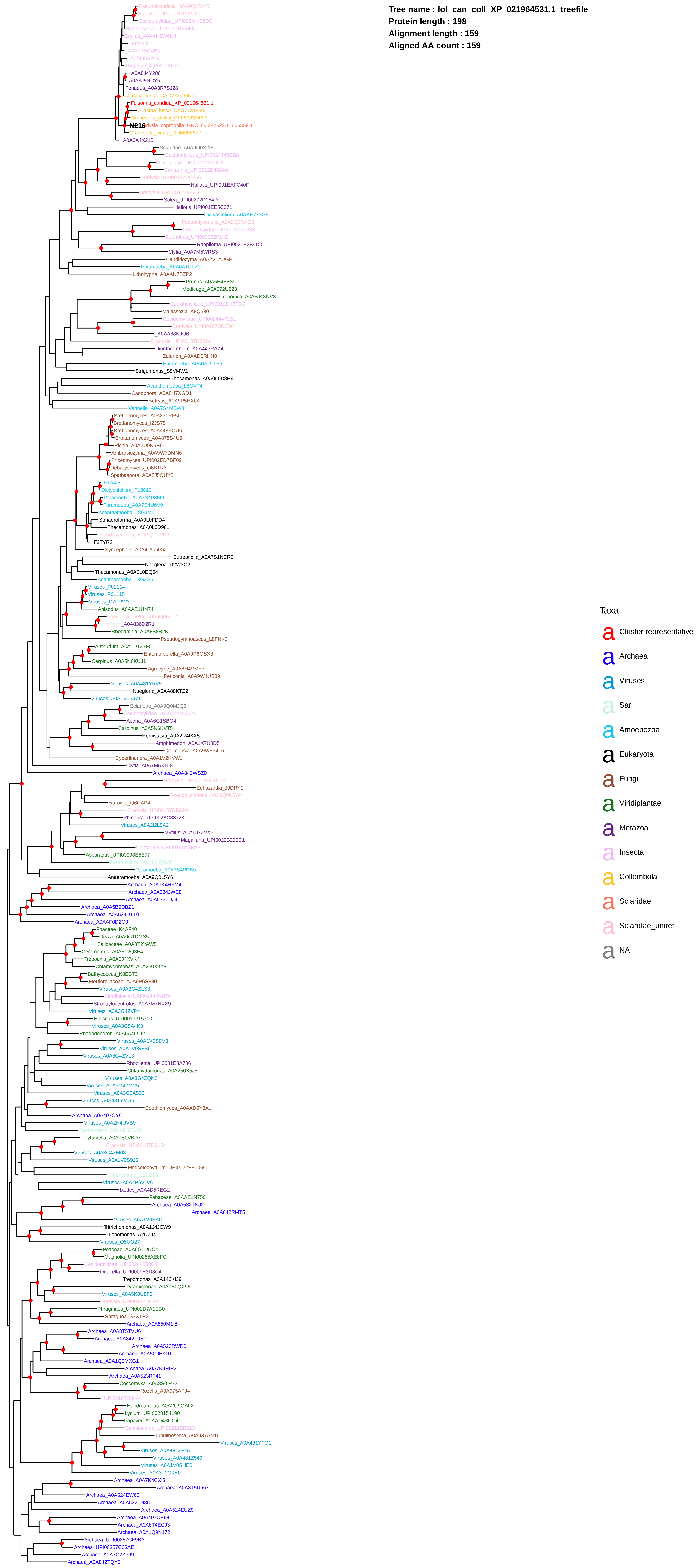

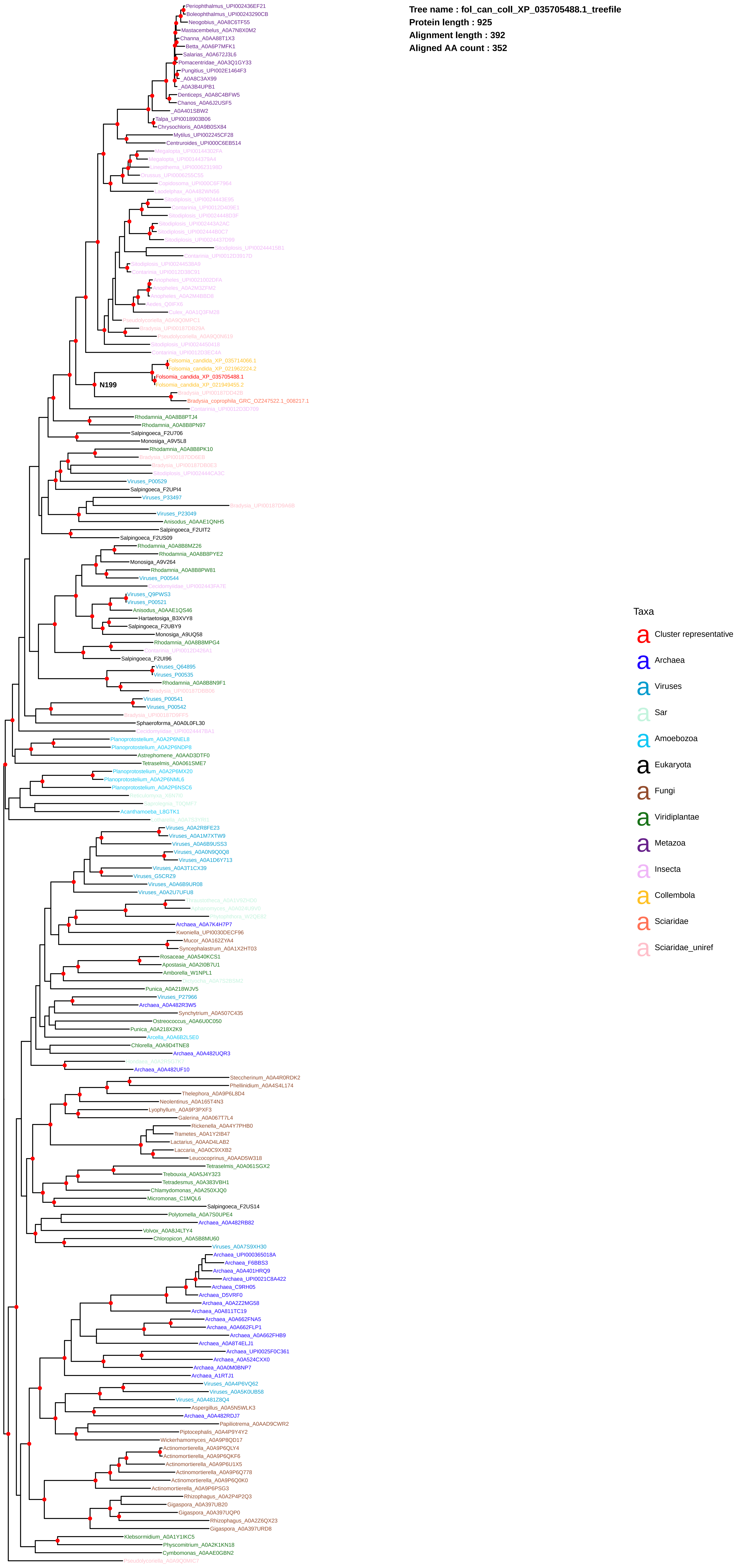

Tree name : fol_can_coll_XP_035705488.1_treefile
Protein length : 925
Alignment length : 392
Aligned AA count : 352
Periophthalmus_UPI002436EF21
Boleophthalmus_UPI00243290CB
Neogobius_A0A8C6T5F55
Mastacembelus_A0A7N8X0M2
Channa_A0AA88T1X3
Betta_A0A6P7MFK1
Salarias_A0A672J3L6
Pomacentridae_A0A3Q1GY33
Pungitius_UPI002E1464F3
_A0A8C3AX99
_A0A3B4UPB1
Denticeps_A0A8C4BFW5
Chanos_A0A6J2USF5
_A0A401SBW2
Talpa_UPI0018903B06
Chrysochloris_A0A9B0SX84
Mytilus_UPI002245CF28
Centruroides_UPI000C6EB514
N199
Folsomia_candida_XP_035714066.1
Folsomia_candida_XP_021962224.2
Folsomia_candida_XP_035705488.1
Folsomia_candida_XP_021949455.2
Bradysia_coprophila_GRC_OZ247522.1_008217.1
Rhodamnia_A0A8B8PTJ4
Rhodamnia_A0A8B8PN97
Salpingoeca_F2U706
Monosiga_A9V5L8
Rhodamnia_A0A8B8PK10
Viruses_P00529
Salpingoeca_F2UPI4
Viruses_P33497
Viruses_P23049
Anisodus_A0AAE1QNH5
Salpingoeca_F2UIT2
Salpingoeca_F2US09
Rhodamnia_A0A8B8MZ26
Rhodamnia_A0A8B8PYE2
Monosiga_A9V264
Rhodamnia_A0A8B8PW81
Viruses_P00544
Viruses_Q9PWS3
Viruses_P00521
Anisodus_A0AAE1QS46
Hartaetosiga_B3XVY8
Salpingoeca_F2UBY9
Monosiga_A9UQ58
Rhodamnia_A0A8B8MPG4
Salpingoeca_F2UI96
Viruses_Q64895
Viruses_P00535
Rhodamnia_A0A8B8N9F1
Viruses_P00541
Viruses_P00542
Sphaeroforma_A0A0L0FL30
Planoprotostelium_A0A2P6NEL8
Planoprotostelium_A0A2P6NDP8
Astrephomene_A0AAD3DTF0
Tetraselmis_A0A061SME7
Planoprotostelium_A0A2P6MX20
Planoprotostelium_A0A2P6NML6
Planoprotostelium_A0A2P6NSC6
Acanthamoeba_L8GTK1
Viruses_A0A2R8FE23
Viruses_A0A1M7XTW9
Viruses_A0A6B9USS3
Viruses_A0A0N9Q0Q8
Viruses_A0A1D6Y713
Viruses_A0A3T1CX39
Viruses_G5CRZ9
Viruses_A0A6B9UR08
Viruses_A0A2U7UFU8
Archaea_A0A7K4H7P7
Kwoniella_UPI0030DECF96
Mucor_A0A162ZYA4
Syncephalastrum_A0A1X2HT03
Rosaceae_A0A540KCS1
Apostasia_A0A2I0B7U1
Amborella_W1NPL1
Punica_A0A218WJV5
Viruses_P27966
Archaea_A0A482R3W5
Synchytrium_A0A507C435
Ostreococcus_A0A6U0C050
Punica_A0A218X2K9
Arcella_A0A6B2L5E0
Chlorella_A0A9D4TNE8
Archaea_A0A482UQR3
Archaea_A0A482UF10
Steccherinum_A0A4R0RDK2
Phellinidium_A0A4S4L174
Thelephora_A0A9P6L8D4
Neolentinus_A0A165T4N3
Lyophyllum_A0A9P3PXF3
Galerina_A0A067T7L4
Rickenella_A0A4Y7PHB0
Trametes_A0A1Y2IB47
Lactarius_A0AAD4LAB2
Laccaria_A0A0C9XXB2
Leucocoprinus_A0AAD5W318
Tetraselmis_A0A061SGX2
Trebouxia_A0A5J4Y323
Tetradesmus_A0A383VBH1
Chlamydomonas_A0A250XJQ0
Micromonas_C1MQL6
Salpingoeca_F2US14
Polytomella_A0A7S0UPE4
Archaea_A0A482RB82
Volvox_A0A8J4LTY4
Chloropicon_A0A5B8MU60
Viruses_A0A7S9XH30
Archaea_UPI000365018A
Archaea_F6BBS3
Archaea_A0A401HRQ9
Archaea_UPI0021C8A422
Archaea_C9RH05
Archaea_D5VRF0
Archaea_A0A2Z2MG58
Archaea_A0A811TC19
Archaea_A0A662FNA5
Archaea_A0A662FLP1
Archaea_A0A662FHB9
Archaea_A0A8T4ELJ1
Archaea_UPI0025F0C361
Archaea_A0A524CXX0
Archaea_A0A0M0BNP7
Archaea_A1RTJ1
Viruses_A0A4P6VQ62
Viruses_A0A5K0UB58
Viruses_A0A4B1Z8Q4
Aspergillus_A0A5N5WLK3
Archaea_A0A482RDJ7
Papiliotrema_A0AAD9CWR2
Piptocephalis_A0A4P9Y4Y2
Wickerhamomyces_A0A9P8QD17
Actinomortierella_A0A9P6QLY4
Actinomortierella_A0A9P6QKF6
Actinomortierella_A0A9P6U1X5
Actinomortierella_A0A9P6Q778
Actinomortierella_A0A9P6Q0K0
Actinomortierella_A0A9P6PSG3
Rhizophagus_A0A2P4P2Q3
Gigaspora_A0A397UB20
Gigaspora_A0A397UQP0
Rhizophagus_A0A2Z6QX23
Gigaspora_A0A397URD8
Klebsormidium_A0A1Y1IKC5
Physcomitrium_A0A2K1KN18
Cymbomonas_A0AAE0GBN2
Taxa
Cluster representative
Archaea
Viruses
Sar
Amoebozoa
Eukaryota
Fungi
Viridiplantae
Metazoa
Insecta
Collembola
Sciaridae
Sciaridae_uniref

**Tree name : orc_cin_coll_ODM86514.1_treefile**
**Protein length : 668**
**Alignment length : 597**
**Aligned AA count : 575**

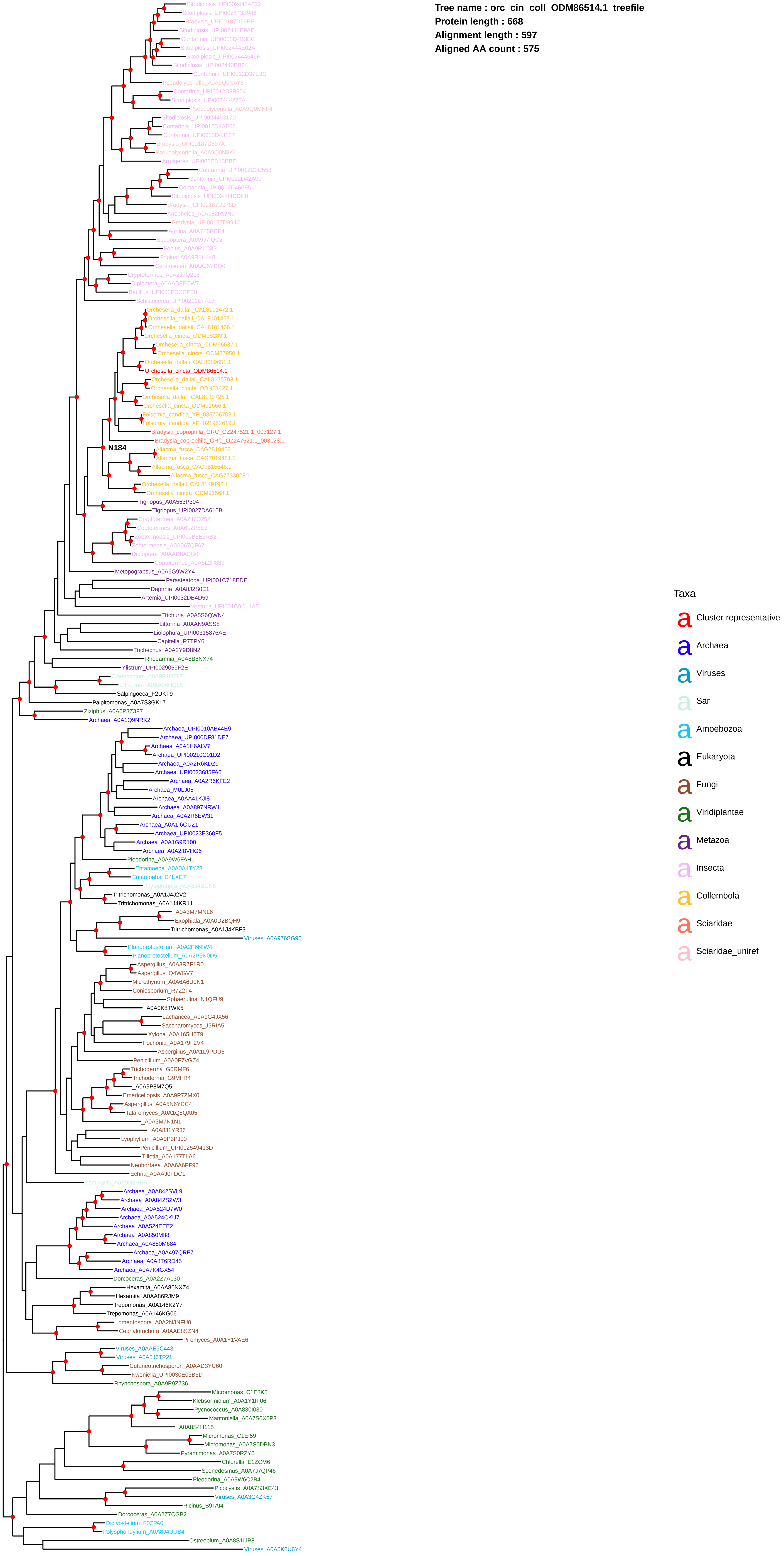

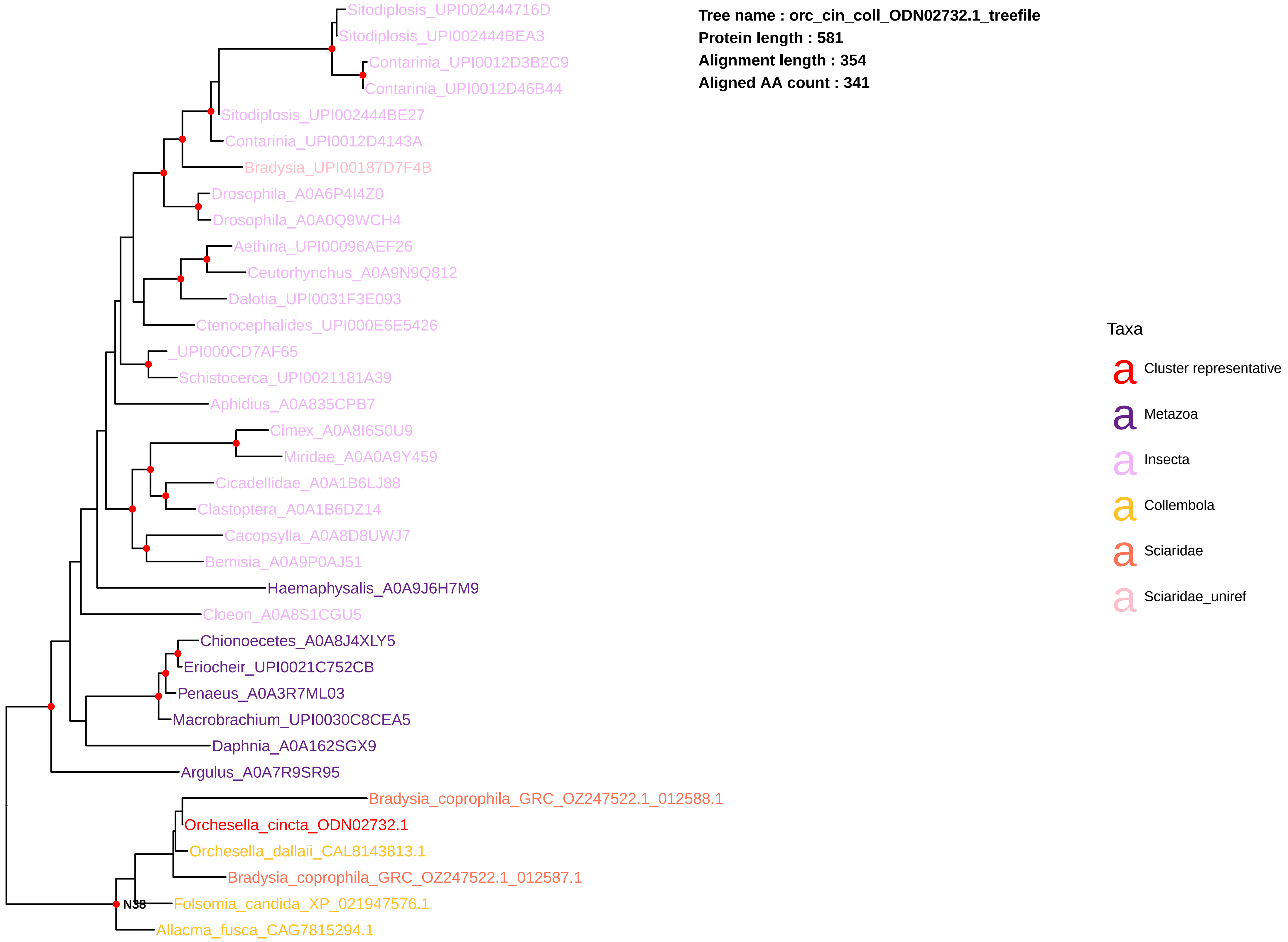
Tree name : orc_cin_coll_ODN02732.1_treefile
Protein length : 581
Alignment length : 354
Aligned AA count : 341
Sitodiplosis_UPI002444716D
Sitodiplosis_UPI002444BEA3
Contarinia_UPI0012D3B2C9
Contarinia_UPI0012D46B44
Sitodiplosis_UPI002444BE27
Contarinia_UPI0012D4143A
Bradysia_UPI00187D7F4B
Drosophila_A0A6P4I4Z0
Drosophila_A0A0Q9WCH4
Aethina_UPI00096AEF26
Ceutorhynchus_A0A9N9Q812
Dalotia_UPI0031F3E093
Ctenocephalides_UPI000E6E5426
_UPI000CD7AF65
Schistocerca_UPI0021181A39
Aphidius_A0A835CPB7
Cimex_A0A8I6S0U9
Miridae_A0A0A9Y459
Cicadellidae_A0A1B6LJ88
Clastoptera_A0A1B6DZ14
Cacopsylla_A0A8D8UWJ7
Bemisia_A0A9P0AJ51
Haemaphysalis_A0A9J6H7M9
Cloeon_A0A8S1CGU5
Chionoecetes_A0A8J4XLY5
Eriocheir_UPI0021C752CB
Penaeus_A0A3R7ML03
Macrobrachium_UPI0030C8CEA5
Daphnia_A0A162SGX9
Argulus_A0A7R9SR95
Bradysia_coprophila_GRC_OZ247522.1_012588.1
Orchesella_cincta_ODN02732.1
Orchesella_dallaii_CAL8143813.1
Bradysia_coprophila_GRC_OZ247522.1_012587.1
N38
Folsomia_candida_XP_021947576.1
Allacma_fusca_CAG7815294.1
Taxa
Cluster representative
Metazoa
Insecta
Collembola
Sciaridae
Sciaridae_uniref

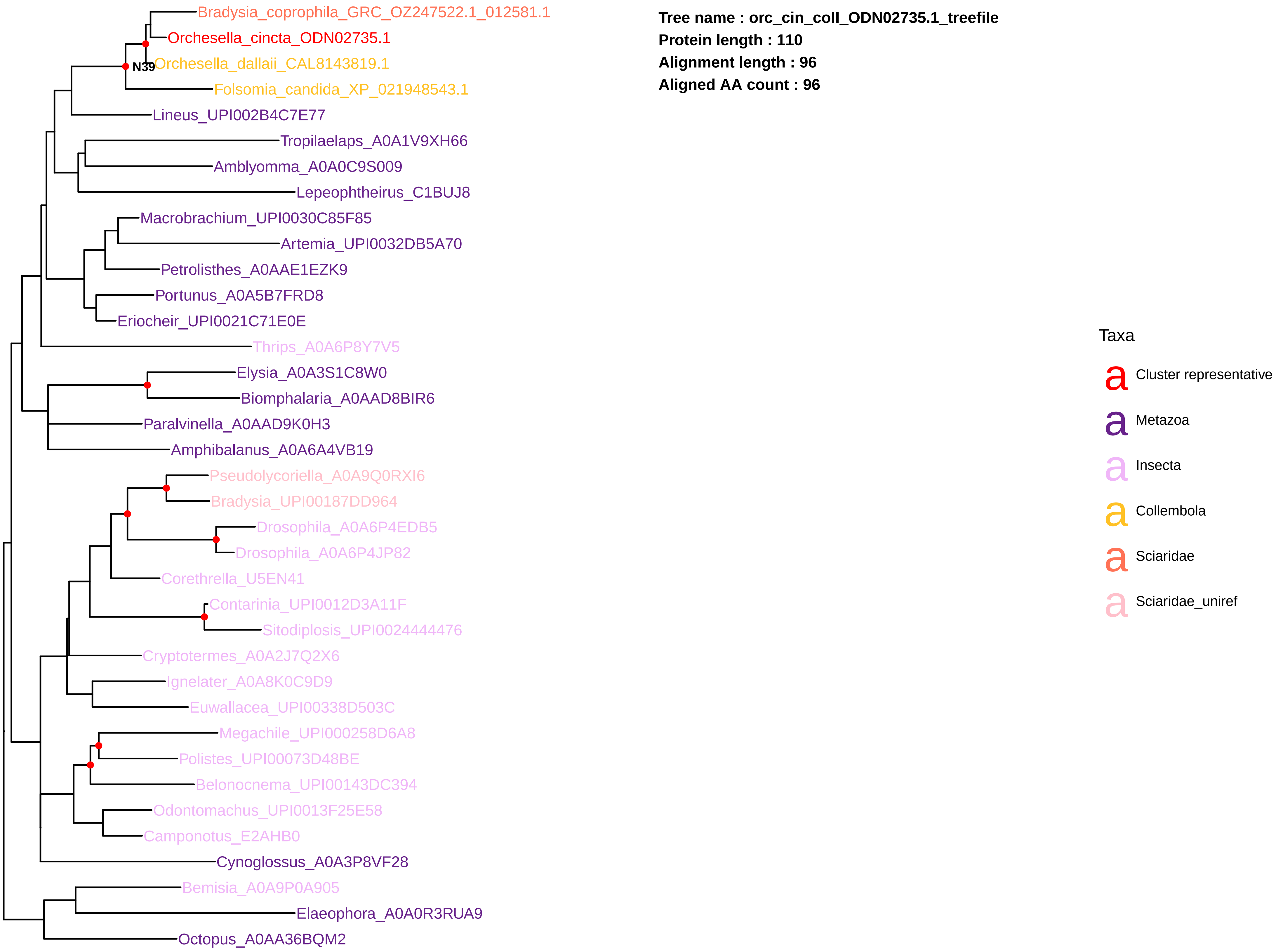


**Tree name : orc_cin_coll_ODN02735.1_treefile**
**Protein length : 110**
**Alignment length : 96**
**Aligned AA count : 96**

**Tree name : orc_cin_coll_ODN04165.1_treefile**
**Protein length : 568**
**Alignment length : 391**
**Aligned AA count : 390**

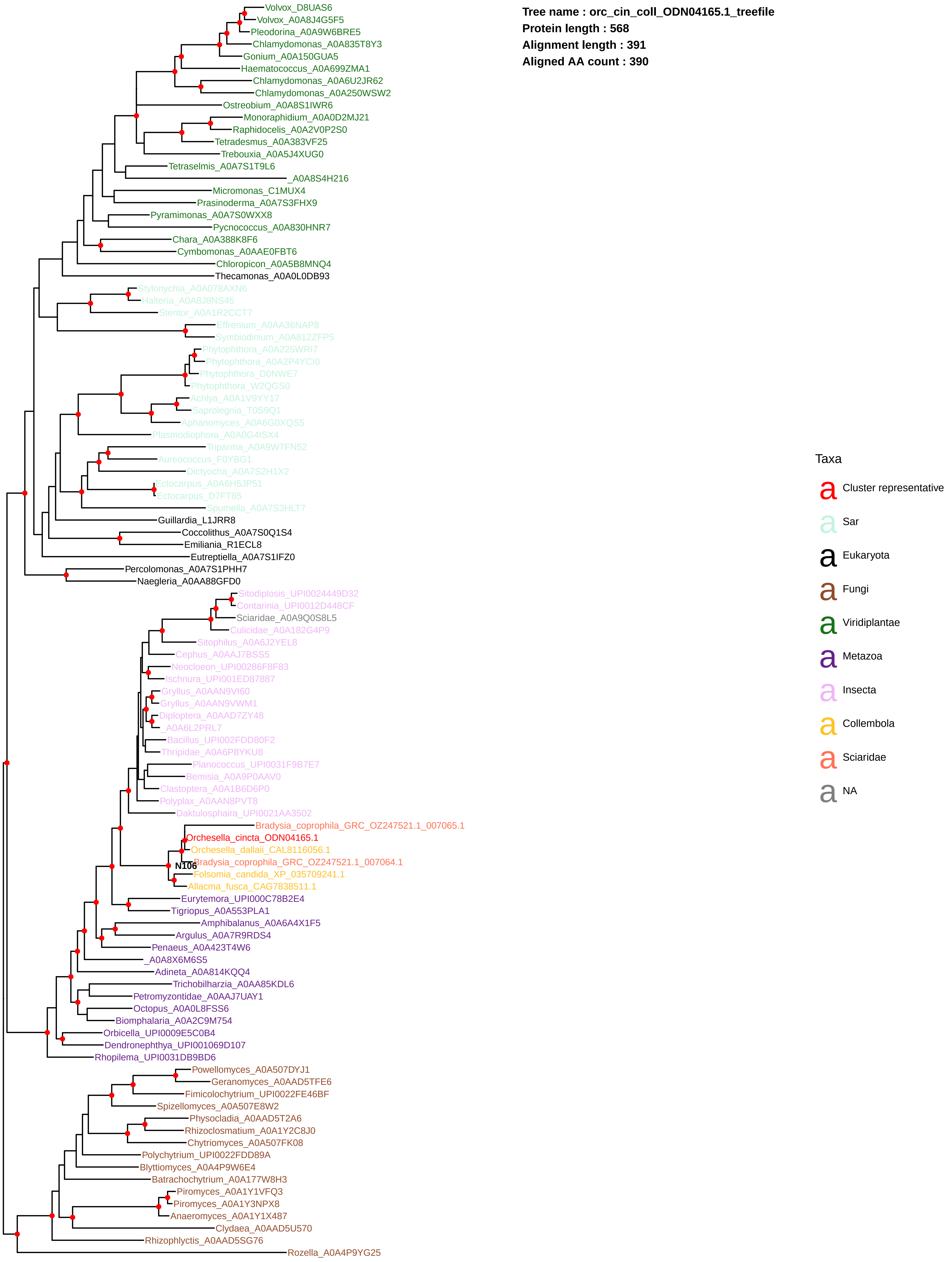

**Tree name : orc_dal_coll_CAL8081574.1_treefile**
**Protein length : 765**
**Alignment length : 247**
**Aligned AA count : 247**

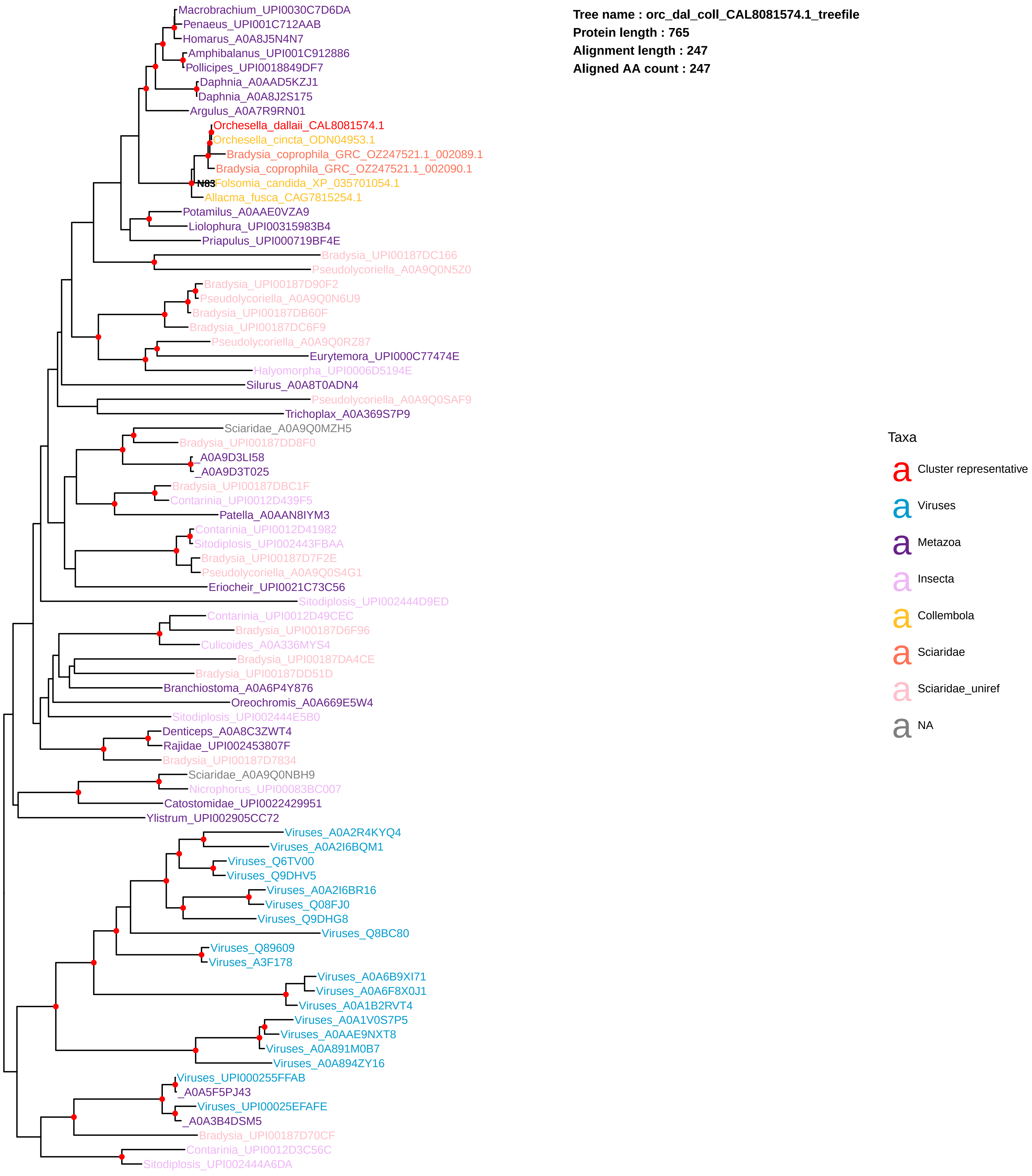

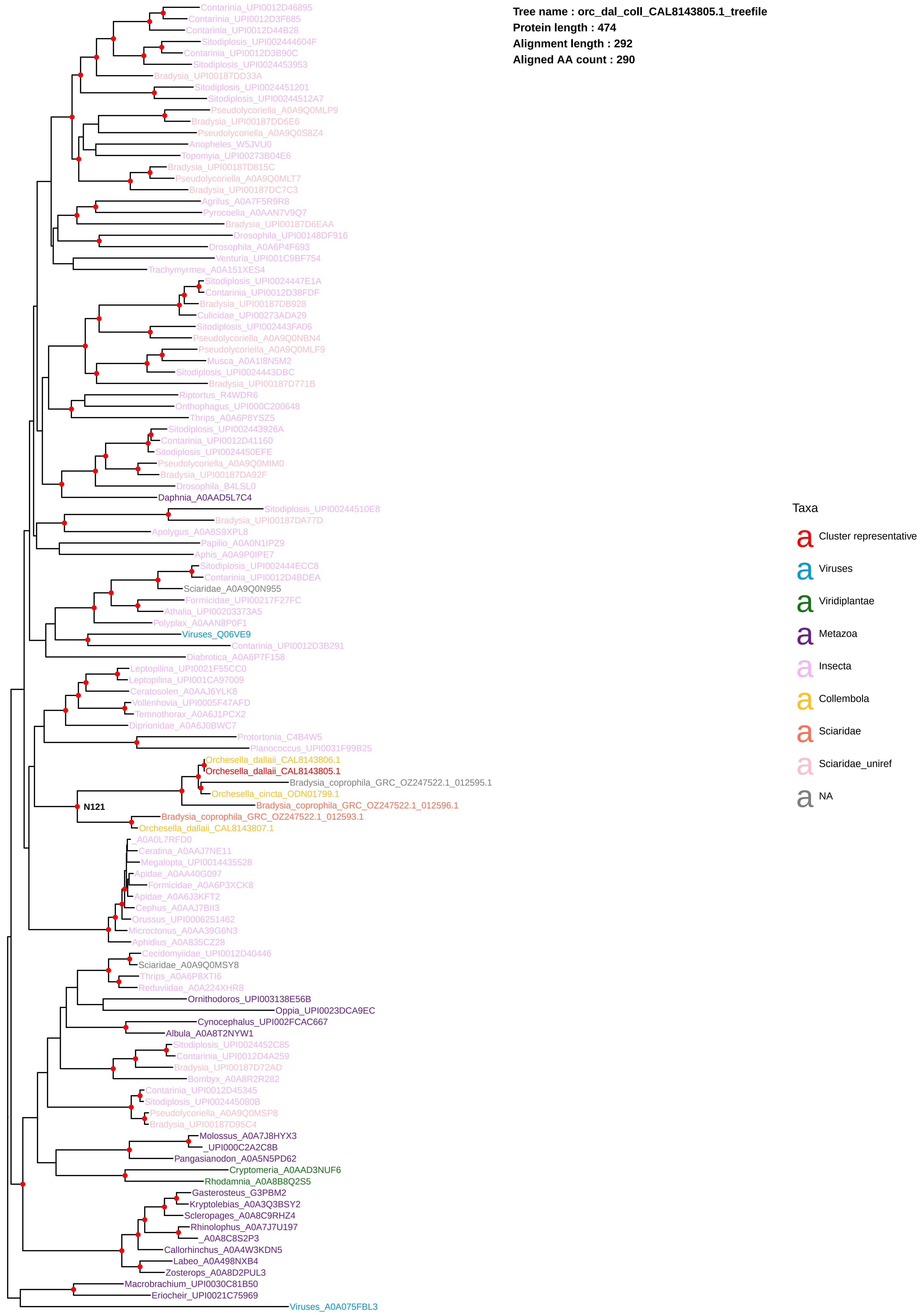

Tree name : orc_dal_coll_CAL8143805.1_treefile
Protein length : 474
Alignment length : 292
Aligned AA count : 290
Contarinia_UPI0012D46895
Contarinia_UPI0012D3F685
Contarinia_UPI0012D44B28
Sitodiplosis_UPI002444604F
Contarinia_UPI0012D3B90C
Sitodiplosis_UPI0024453953
Bradysia_UPI00187DD33A
Sitodiplosis_UPI0024451201
Sitodiplosis_UPI00244512A7
Pseudolycoriella_A0A9Q0MLP9
Bradysia_UPI00187DD6E6
Pseudolycoriella_A0A9Q0S8Z4
Anopheles_W5JVU0
Topomyia_UPI00273B04E6
Bradysia_UPI00187D815C
Pseudolycoriella_A0A9Q0MLT7
Bradysia_UPI00187DC7C3
Agrilus_A0A7F5R9R8
Pyrocoelia_A0AAN7V9Q7
Bradysia_UPI00187D6EAA
Drosophila_UPI00148DF916
Drosophila_A0A6P4F693
Venturia_UPI001C9BF754
Trachymyrmex_A0A151XES4
Sitodiplosis_UPI0024447E1A
Contarinia_UPI0012D38FDF
Bradysia_UPI00187DB928
Culicidae_UPI00273ADA29
Sitodiplosis_UPI002443FA06
Pseudolycoriella_A0A9Q0NBN4
Pseudolycoriella_A0A9Q0MLF9
Musca_A0A1I8N5M2
Sitodiplosis_UPI0024443DBC
Bradysia_UPI00187D771B
Riptortus_R4WDR6
Onthophagus_UPI000C200648
Thrips_A0A6P8YSZ5
Sitodiplosis_UPI002443926A
Contarinia_UPI0012D41160
Sitodiplosis_UPI0024450EFE
Pseudolycoriella_A0A9Q0MIM0
Bradysia_UPI00187DA92F
Drosophila_B4LSL0
Daphnia_A0AAD5L7C4
Sitodiplosis_UPI00244510E8
Bradysia_UPI00187DA77D
Apolygus_A0A8S9XPL8
Papilio_A0A0N1IPZ9
Aphis_A0A9P0IPE7
Sitodiplosis_UPI002444ECC8
Contarinia_UPI0012D4BDEA
Sciaridae_A0A9Q0N955
Formicidae_UPI00217F27FC
Athalia_UPI00203373A5
Polyplax_A0AAN8P0F1
Viruses_Q06VE9
Contarinia_UPI0012D3B291
Diabrotica_A0A6P7F158
Leptopilina_UPI0021F55CC0
Leptopilina_UPI001CA97009
Ceratosolen_A0AAJ6YLK8
Vollenhovia_UPI0005F47AFD
Temnothorax_A0A6J1PCX2
Diprionidae_A0A6J0BWC7
Protortonia_C4B4W5
Planococcus_UPI0031F99B25
Orchesella_dallaii_CAL8143806.1
Orchesella_dallaii_CAL8143805.1
Bradysia_coprophila_GRC_OZ247522.1_012595.1
Orchesella_cincta_ODN01799.1
N121
Bradysia_coprophila_GRC_OZ247522.1_012596.1
Bradysia_coprophila_GRC_OZ247522.1_012593.1
Orchesella_dallaii_CAL8143807.1
_A0A0L7RFD0
Ceratina_A0AAJ7NE11
Megalopta_UPI0014435528
Apidae_A0AA40G097
Formicidae_A0A6P3XCK8
Apidae_A0A6J3KFT2
Cephus_A0AAJ7BII3
Orussus_UPI0006251462
Microctonus_A0AA39G6N3
Aphidius_A0A835CZ28
Cecidomyiidae_UPI0012D40446
Sciaridae_A0A9Q0MSY8
Thrips_A0A6P8XTI6
Reduviidae_A0A224XHR8
Ornithodoros_UPI003138E56B
Oppia_UPI0023DCA9EC
Cynocephalus_UPI002FCAC667
Albula_A0A8T2NYW1
Sitodiplosis_UPI0024452C85
Contarinia_UPI0012D4A259
Bradysia_UPI00187D72AD
Bombyx_A0A8R2R282
Contarinia_UPI0012D45345
Sitodiplosis_UPI002445080B
Pseudolycoriella_A0A9Q0MSP8
Bradysia_UPI00187D95C4
Molossus_A0A7J8HYX3
_UPI000C2A2C8B
Pangasianodon_A0A5N5PD62
Cryptomeria_A0AAD3NUF6
Rhodamnia_A0A8B8Q2S5
Gasterosteus_G3PBM2
Kryptolebias_A0A3Q3BSY2
Scleropages_A0A8C9RHZ4
Rhinolophus_A0A7J7U197
_A0A8C8S2P3
Callorhinchus_A0A4W3KDN5
Labeo_A0A498NXB4
Zosterops_A0A8D2PUL3
Macrobrachium_UPI0030C81B50
Eriocheir_UPI0021C75969
Viruses_A0A075FBL3
Taxa
Cluster representative
Viruses
Viridiplantae
Metazoa
Insecta
Collembola
Sciaridae
Sciaridae_uniref
NA

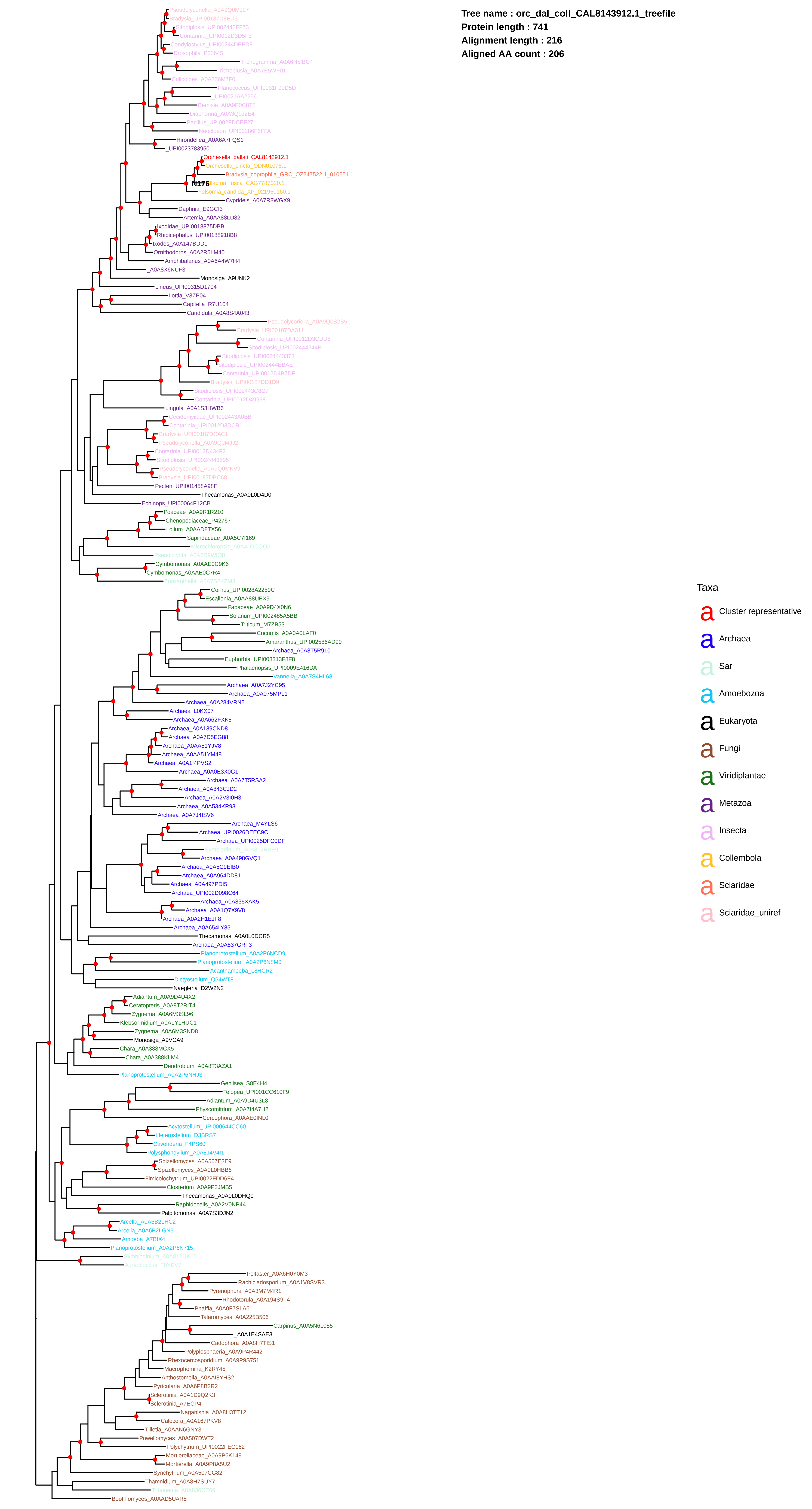
Tree name : orc_dal_coll_CAL8143912.1_treefile
Protein length : 741
Alignment length : 216
Aligned AA count : 206
Pseudolycoriella_A0A9Q0MJ27
Bradysia_UPI00187D8ED3
Sitodiplosis_UPI002443FF73
Contarinia_UPI0012D3D5F3
Condylostylus_UPI00244DEED8
Drosophila_P23645
Trichogramma_A0A6H5IBC4
Trichoplusia_A0A7E5WFO1
Culicoides_A0A336M7F0
Planococcus_UPI0031F90D5D
_UPI0021AA2256
Bemisia_A0A9P0C8T8
Diaphorina_A0A3Q0J2E4
Bacillus_UPI002FDCEF27
Neocloeon_UPI00286F6FFA
Hirondella_A0A6A7FQS1
_UPI0023783950
Orchesella_dallaii_CAL8143912.1
Orchesella_cincta_ODN01078.1
Bradysia_coprophila_GRC_OZ247522.1_010551.1
N176
Allacma_fusca_CAG7787020.1
Folsomia_candida_XP_021950160.1
Cyprideis_A0A7R8WGX9
Daphnia_E9GCI3
Artemia_A0AA88LD82
Ixodidae_UPI0018875DBB
Rhipicephalus_UPI00188918B8
Ixodes_A0A147BDD1
Ornithodoros_A0A2R5LM40
Amphibalanus_A0A6A4W7H4
_A0A8X6NUF3
Monosiga_A9UNK2
Lineus_UPI00315D1704
Lottia_V3ZP04
Capitella_R7U104
Candidula_A0A8S4A043
Pseudolycoriella_A0A9Q0S2S5
Bradysia_UPI00187DA311
Contarinia_UPI0012D3CDD8
Sitodiplosis_UPI002444244E
Sitodiplosis_UPI0024443373
Sitodiplosis_UPI002444EBAE
Contarinia_UPI0012D4B7DF
Bradysia_UPI00187DD1D9
Sitodiplosis_UPI002443C8C7
Contarinia_UPI0012D49998
Lingula_A0A1S3HWB6
Cecidomyiidae_UPI002443A0BB
Contarinia_UPI0012D3DCB1
Bradysia_UPI00187DCAC1
Pseudolycoriella_A0A9Q0MJJ2
Contarinia_UPI0012D434F2
Sitodiplosis_UPI0024443595
Pseudolycoriella_A0A9Q0MKV9
Bradysia_UPI00187DBC68
Pecten_UPI001458A98F
Thecamonas_A0A0L0D4D0
Echinops_UPI00064F12CB
Poaceae_A0A9R1R210
Chenopodiaceae_P42767
Lolium_A0AAD8TX56
Sapindaceae_A0A5C7I169
Microchloropsis_A0A4D0CQQ6
Pseudictyota_A0A7R9WIQ8
Cymbomonas_A0AAE0C9K6
Cymbomonas_A0AAE0C7R4
Zooxanthella_A0A7S2KZM3
Cornus_UPI0028A2259C
Escallonia_A0AA88UEX9
Fabaceae_A0A9D4X0N6
Solanum_UPI002485A5BB
Triticum_M7ZB53
Cucumis_A0A0A0LAF0
Amaranthus_UPI002586AD99
Archaea_A0A8T5R910
Euphorbia_UPI003313F8F8
Phalaenopsis_UPI0009E416DA
Vannella_A0A7S4HL68
Archaea_A0A7J2YC95
Archaea_A0A075MPL1
Archaea_A0A284VRN5
Archaea_L0KX07
Archaea_A0A662FXK5
Archaea_A0A139CND8
Archaea_A0A7D5EG88
Archaea_A0AA51YJV8
Archaea_A0AA51YM48
Archaea_A0A1I4PVS2
Archaea_A0A0E3X0G1
Archaea_A0A7T5RSA2
Archaea_A0A843CJD2
Archaea_A0A2V3I0H3
Archaea_A0A534KR93
Archaea_A0A7J4ISV6
Archaea_M4YLS6
Archaea_UPI0026DEEC9C
Archaea_UPI0025DFC0DF
Symbiodinium_A0A812RXE9
Archaea_A0A498GVQ1
Archaea_A0A5C9EIB0
Archaea_A0A964DD81
Archaea_A0A497PDI5
Archaea_UPI002D098C64
Archaea_A0A835XAK5
Archaea_A0A1Q7X9V8
Archaea_A0A2H1EJF8
Archaea_A0A654LY85
Thecamonas_A0A0L0DCR5
Archaea_A0A537GRT3
Planoprotostelium_A0A2P6NCD9
Planoprotostelium_A0A2P6N8M0
Acanthamoeba_L8HCR2
Dictyostelium_Q54WT8
Naegleria_D2W2N2
Adiantum_A0A9D4U4X2
Ceratopteris_A0A8T2RIT4
Zygnema_A0A6M3SL96
Klebsormidium_A0A1Y1HUC1
Zygnema_A0A6M3SND8
Monosiga_A9VCA9
Chara_A0A388MCX5
Chara_A0A388KLM4
Dendrobium_A0A8T3AZA1
Planoprotostelium_A0A2P6NHJ3
Genlisea_S8E4H4
Telopea_UPI001CC610F9
Adiantum_A0A9D4U3L8
Physcomitrium_A0A7I4A7H2
Cercophora_A0AAE0INL0
Acytostelium_UPI000644CC60
Heterostelium_D3BRS7
Cavenderia_F4PS60
Polysphondylium_A0A8J4V4I1
Spizellomyces_A0A507E3E9
Spizellomyces_A0A0L0HBB6
Fimicolochytrium_UPI0022FDD6F4
Closterium_A0A9P3JMB5
Thecamonas_A0A0L0DHQ0
Raphidocelis_A0A2V0NP44
Palpitomonas_A0A7S3DJN2
Arcella_A0A6B2LHC2
Arcella_A0A6B2LGN5
Amoeba_A7BIX4
Planoprotostelium_A0A2P6N715
Symbiodinium_A0A812UKL0
Aureococcus_F0YEV7
Peltaster_A0A6H0Y0M3
Rachicladosporium_A0A1V8SVR3
Pyrenophora_A0A3M7M4R1
Rhodotorula_A0A194S9T4
Phaffia_A0A0F7SLA6
Talaromyces_A0A225B506
Carpinus_A0A5N6L055
_A0A1E4SAE3
Cadophora_A0A8H7TIS1
Polyplosphaeria_A0A9P4R442
Rhexocercosporidium_A0A9P9S751
Macrophomina_K2RY45
Anthostomella_A0AAI8YHS2
Pyricularia_A0A6P8B2R2
Sclerotinia_A0A1D9Q2K3
Sclerotinia_A7ECP4
Naganishia_A0A8H3TT12
Calocera_A0A167PKV8
Tilletia_A0AAN6GNY3
Powellomyces_A0A507DWT2
Polychytrium_UPI0022FEC162
Mortierellaceae_A0A9P6K149
Mortierella_A0A9P8A5U2
Synchytrium_A0A507CG82
Thamnidium_A0A8H7SUY7
Tribonema_A0A836CE69
Boothiomyces_A0AAD5UAR5
Taxa
Cluster representative
Archaea
Sar
Amoebozoa
Eukaryota
Fungi
Viridiplantae
Metazoa
Insecta
Collembola
Sciaridae
Sciaridae_uniref
